# Maintenance of chronic neuroinflammation in multiple sclerosis via interferon signaling and CD8 T cell-mediated cytotoxicity

**DOI:** 10.1101/2025.06.09.658729

**Authors:** Syed Ali Raza, Yoshimi Enose-Akahata, Anna Blazier, Lauren M. Guerra, Erin S. Beck, María I. Gaítan, Nyater Ngouth, Jefte M. Drijvers, Eric B. Dammer, Devan Moodley, Kory R. Johnson, Martina Absinta, Timothy J. Turner, Richard M. Ransohoff, Steven Jacobson, Dimitry Ofengeim, Ellen Cahir-McFarland, Daniel S. Reich

**Author notes:** Correspondence to Daniel S. Reich.

## Abstract

Chronic neuroinflammation and neurodegeneration are critical but unresolved drivers of disability accumulation in progressive multiple sclerosis (MS). Chronic active white matter lesions (CAL), identifiable radiologically as paramagnetic rim lesions (PRL), indicate progression-relevant chronic neuroinflammation. Using single-cell transcriptomics (scRNAseq) and T-cell receptor sequencing (scTCR-seq), we profiled cerebrospinal fluid (CSF) and blood immune cells of 34 radiologically characterized adults with MS (17 untreated, 6 treated with B-cell-depletion) and 5 healthy controls. Coupled with proteomics, we found PRL-associated enrichment of interferon (IFN) signaling and upregulation of TCR signaling in CSF and blood. This was accompanied by clonal expansion of CD8^+^ T effector memory (TEM) cells, with the highly expanded clonal cells exhibiting T helper type 1 (T_H_1) and cytotoxic profiles. Validating the cytotoxic immune profile in blood using flow cytometry, we identified a cellular correlate of PRL exhibiting features of CD8^+^ T_EMRA_ cells. Despite B-cell depletion, PRL-associated neuroinflammation, driven by myeloid activation and CD8^+^ T-cell cytotoxicity, persisted. Serum and CSF proteomic networks showed PRL-pertinent signatures, including networks unaffected by B-cell depletion. Using *in silico* perturbation, we nominated therapeutic targets, including *MYD88*, *TNF*, *MYC*, *TYK2*, *JAK2,* and *BTK*, for alleviating chronic neuroinflammation in MS. Our findings highlight mechanisms of chronic neuroinflammation in MS and point to potential biomarkers for monitoring disease progression.

## Main

Recent advances provide a framework for redefining MS as an inflammation-mediated neurodegenerative disorder, with chronic neuroinflammation being central to progressive disability accumulation^1,2^. Understanding these processes from a mechanistic standpoint is imperative to address unmet medical needs in progressive MS, where effective treatments are very limited. CAL detected at autopsy are strongly associated with adverse clinical course during life^1,3^ and are mechanistically associated with progression independent of relapse activity (PIRA)^4^. Identification of CAL by their characteristic paramagnetic rims on advanced magnetic resonance imaging (MRI), in more than half of MS cases^5^, has provided new opportunities to probe mechanisms underlying pathophysiology^4–6^. To this end, MRI-informed single-nucleus RNA sequencing (snRNAseq) profiling of cells at the edge of demyelinated lesions at various stages of inflammation has been performed^7^, but the blood and cerebrospinal fluid (CSF) immune correlates of CAL remain undetermined, despite the fact that multiple studies have previously investigated the cellular composition of CSF, contrasting MS-affected and healthy controls^8–14^.

Here, we investigated the CSF and peripheral immune cell correlates of PRL, studying untreated individuals and exploring the effect of B-cell depletion on the transcriptome and proteome. We profiled matched CSF cells and peripheral blood mononuclear cells (PBMC) from 39 adults. Among these, we performed three primary comparisons (Supplementary File 1, Table 1) involving untreated, radiologically and clinically inactive MS cases (those without a gadolinium-enhancing or new lesion and without a clinical relapse within 6 months from the time of sample acquisition, referred to hereafter as “untreated and inactive MS”), with and without PRL, as well as cases with PRL despite treatment with anti-CD20-antibody-mediated B-cell depletion.

**Table 1.** Cohort demographics. Times are measured from the point of sample acquisition (CSF and blood), except for the time between except for the “time between MRI and LP,” where the date of LP was subtracted from that of the MRI. Abbreviations – RIS: Radiologically Isolated Syndrome; RRMS: Relapsing Remitting Multiple Sclerosis; SPMS: Secondary Progressive Multiple Sclerosis; PPMS: Primary Progressive Multiple Sclerosis; DMT: Disease Modifying Therapy; SD: Standard Deviation; M: Male; F: Female

|  | Untreated and radiologically inactive |  | Anti-CD20-treated cases |  |
| --- | --- | --- | --- | --- |
|  | PRL-positive | PRL-negative | Anti-CD20-monoclonal antibody treated (ocrelizumab) | Healthy volunteer |
| Number of participants – No | 10 | 7 | 6 | 5 |
| Age at LP (Mean ± SD years) | 56 ± 11 | 63 ± 12 | 47 ± 10 | 57.2 ± 10.6 |
| Sex (M/F, F%) | 5/5, 50 | 4/3, 43 | 3/3, 50 | 3/2, 40 |
| Age at symptom onset (Mean ± SD years) | 39 ± 10 | 50 ± 9 | 34 ± 10 | |
| Disease duration (Mean ± SD years) | 17 ± 13 | 13 ± 13 | 13 ± 9 | |
| EDSS (Mean; Range) | 2 (0 – 7) | 2 (1 – 4) | 3 (1 – 6) |  |
| Clinical course at the time of LP: |  |  |  |  |
| RIS | 5 |  |  |  |
| RRMS | 1 | 4 | 3 |  |
| SPMS | 3 |  | 2 |  |
| PPMS | 1 | 3 | 1 |  |
| Time since last DMT (Mean ± SD years) | 2.7 ± 2.0 | 6.8 ± 4.6 | | |
| Time since last lesion and/or relapse (Mean ± SD years) | 7.0 ± 6.1 | 1.9 ± 5.7 | 4.4 ± 2.0 | |
| Time since start of Anti-CD20-monoclonal antibody (Mean ± SD years) | | | 2.8 ± 1.0 | |
| Time between MRI and LP (Mean ± SD days; [MRI date – LP date]) | -42 ± 48 | -32 ± 36 | -65 ± 108 | |

**Table 2.**
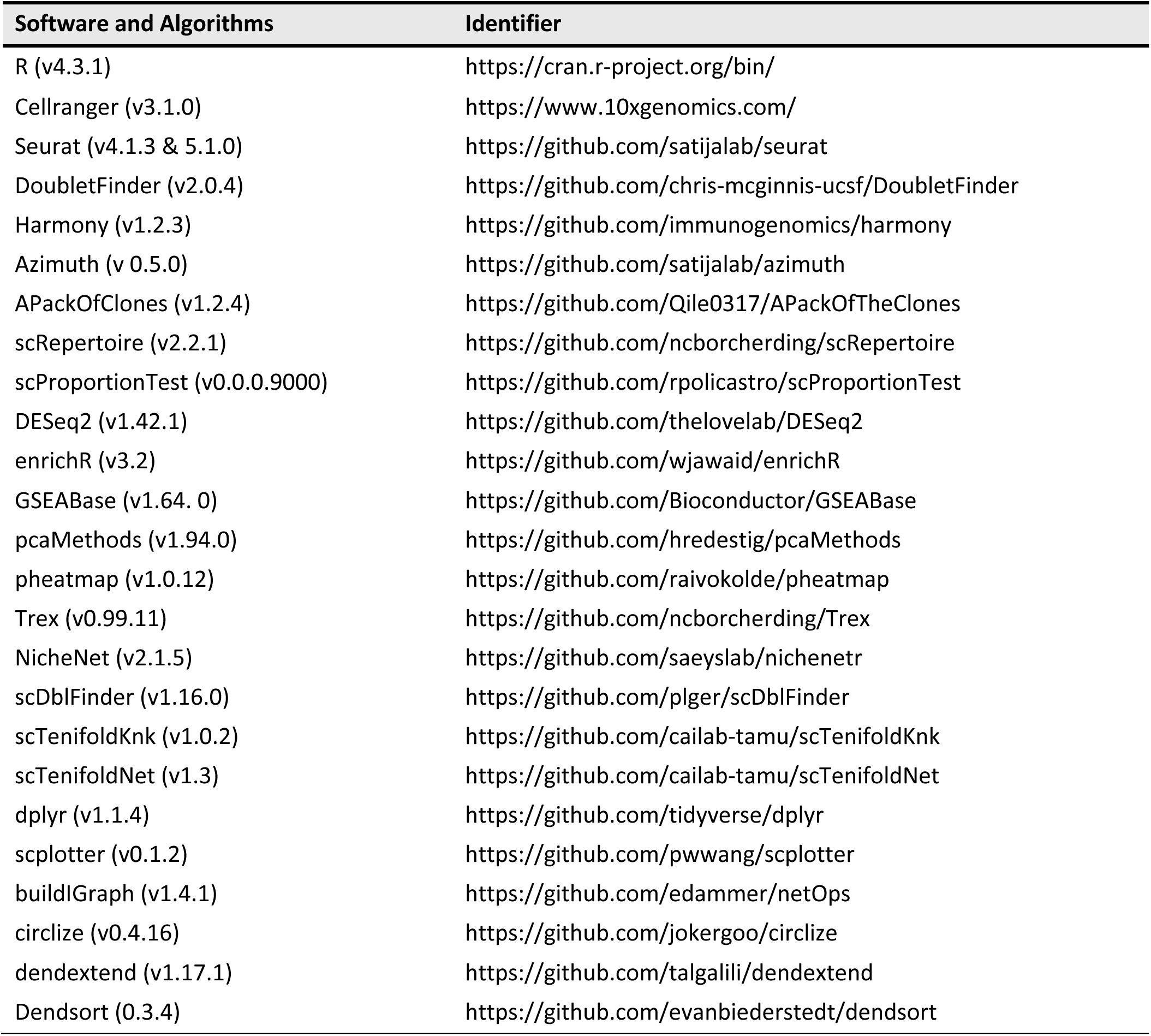
Key resources.

### Overview of multiomics analysis

To decipher cellular and molecular mechanisms pertinent to PRL-pathology and B-cell depletion in PRL-positive and PRL-negative cases, we leveraged scRNAseq for profiling CSF cells and CSF-matched PBMC from 39 adults and performed scTCR-seq in a subset (13 PBMC and 11 CSF samples, Figure 1A, Supplementary File 1). Following filtering and quality control measures, we profiled 74,501 CSF cellular and 177,556 PBMC transcriptomes. Based on known lineage markers and a reference tool^15^, we annotated level 1 (L1) clusters in the CSF and PBMC (Figure 1B, C). For better characterization of the myeloid, T-lymphoid and NK, and B-lymphoid cells, we used unsupervised clustering to define level 2 clusters (L2) (Extended Figure 1A-D), following erythrocyte and platelet removal from CSF samples (see Methods; Figure S2). Note that in general, CSF samples were not contaminated with blood considering the negligible numbers of erythrocytes or platelets in 11 CSF samples, with only one healthy sample containing 114 erythrocytes (Supplementary File 1). Detailed annotations and the description of all clusters are provided in a <u>Supplementary information document</u> (Figure S3-7), including the lists of differentially expressed cluster-defining genes (Supplementary Files 2-4).

**Figure 1:**
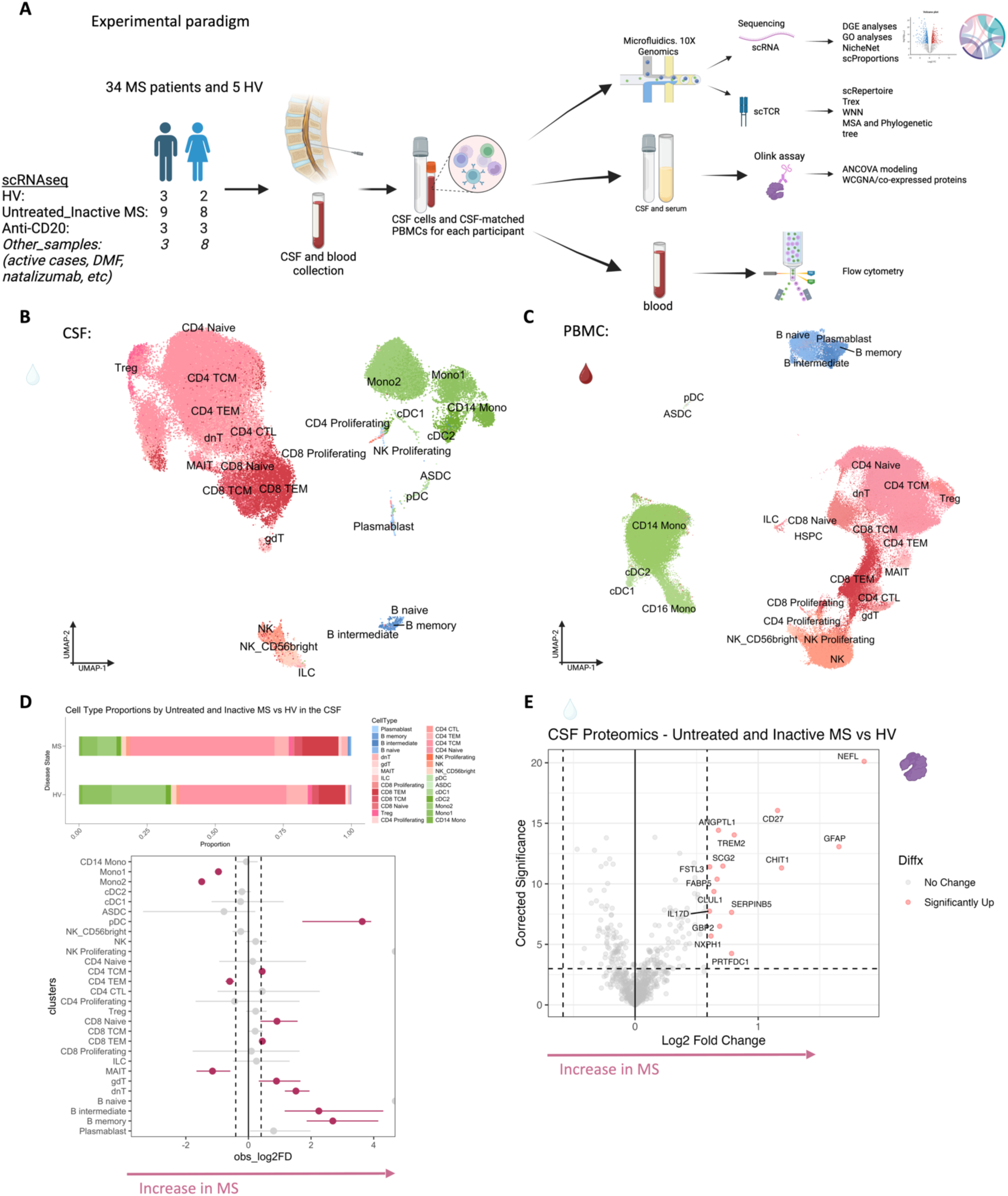
Multimodal analyses reveal a unique inflammatory profile of the CSF of untreated and inactive MS participants. a) Experimental paradigm of a multimodal approach involving single-cell transcriptomics of the CSF cells and CSF-matched PBMCs from 39 adults (34 MS cases and 5 healthy volunteers, HV) using 10X Genomics platform, coupled with proteomics of the serum and CSF performed using Olink Explorer proximity extension assay. Flow cytometric validation in the blood was also performed. This figure was created using Biorender.com. b) UMAP of the level 1 (L1) scRNA-seq analyses of the cellular compartment in CSF representing 74,501 single-cell transcriptomes with a total of 28 cell clusters. c) UMAP scatter plot of the L1 analyses of PBMCs measuring 177,556 single-cell transcriptomes and illustrating 28 clusters. d) Stacked column graph summarizing proportions of L1 annotations in untreated and inactive MS patients compared to HV (upper panel) in the CSF. Lineplot demonstrating significant differences in the CSF cellular composition between MS cases and HV (lower panel). Single-cell proportion test was used to perform permutation tests, randomly segregating cells into the two conditions while maintaining the original sample size and subsequently calculating proportional difference between the two conditions, followed by comparison with the observed difference (see Methods). Notable are the increased proportions of Bmemory, B-intermediate, and dnT cells in the CSF of untreated and inactive MS cases versus HV. e) Volcano plot of differentially expressed proteins (DEP) contrasting the CSF of inactive and untreated MS cases and HV. The purple-colored globular cartoon on the upper-right of the panel represents proteomic data, and we use this throughout the figures to differentiate from the transcriptomic data, which is shown as a wiggly line. The maroon-colored drop represents blood-derived samples, while the clear drop represents CSF-derived samples. (Biorender.com was used for the globular cartoon, wiggle, and two drops.)

Of note, we analyzed the CSF and PBMC L1 objects separately considering the difference in composition of both tissues^16–18^ and to reduce the chance of regressing out important biological variation by integrating samples across covariates^19^. This is emphasized by the difference in elbow plots for CSF and PBMC in explaining the variance of expression (Figure S8A). However, we also downsized each sample at random to nearly 1000 cells, forming a combined object and contrasting immune cells across the CSF and blood compartments (Figure S8 and <u>Supplementary</u> <u>Information</u>).

### Inflammatory profile of the CSF of untreated and inactive MS patients

Before dissecting immune correlates of PRL-pathology, we contrasted the CSF and PBMC of untreated and inactive MS cases — referred to hereafter as MS in this section unless stated otherwise — versus healthy volunteers (HV). We observed enrichment of B-intermediate and B-memory cells (Figure 1D, Extended Figure 2A) in the CSF of MS, consistent with prior reports^9,12,14^. In addition, we observed expansion of double-negative (dn) T-cells in MS CSF, which has been reported previously^20^ and could be related to T-helper (T_H_) like or regulatory functions^21,22^. We also observed decreased proportions of monocyte (Mono1 and Mono2) clusters in MS CSF, which may reflect increased recruitment into the CNS parenchyma or aberrant myeloid composition considering the increased frequencies of B-cell or T-cell subsets^14,23,24^. The decreased proportion of myeloid cells in the CSF of MS versus HV was also a finding in a recent study^14^. We also found enrichment of plasmacytoid dendritic cells (pDC) in CSF of MS relative to HV. There was also a trend toward increased frequency of CD8-TEM and CD4 T-central-memory (TCM) in MS, which is consistent with prior report^12^.

**Figure 2:**
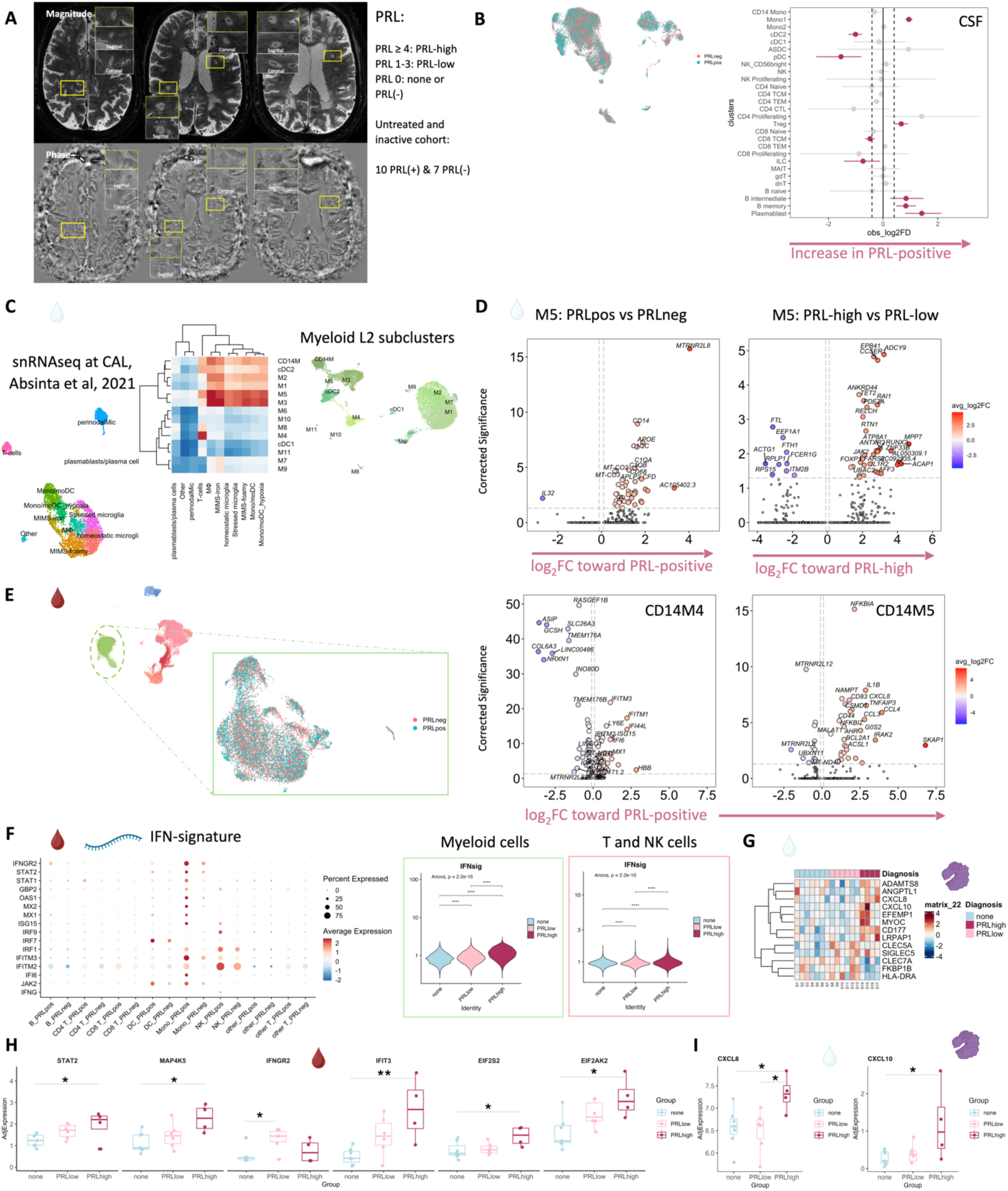
IFN-signature enrichment in MS cases with PRL. a) In vivo advanced 7-tesla MRI demonstrating three PRL in a 70-year old man with primary progressive MS and untreated with any disease-modifying therapy for approximately 6 months. Axial T2*-weighted motion- and B0-corrected images at 0.5 mm isotropic resolution, magnitude and unwrapped filtered phase images. Images are observed in three planes. A hypointense rim surrounds an isointense core in phase images (bottom row); a central vein sign is also observed through the lesions. On the right of the panel is the PRL categorization^5^. b) CSF L1 UMAP for PRL-positive versus PRL-negative comparison with total cell population down-sampled equally per condition to 12,500 cells. Right panel is a lineplot showing increased proportion of B-intermediate, B memory, plasmablast, T-regulatory, and Mono1 clusters in PRL-positive cases. cDC2 is decreased in proportion in PRL-positive cases. c) Correlation heatmap comparing average transcriptome similarity of CSF myeloid L2 subclusters vs. the immune-cell populations from a previously published snRNAseq dataset in different cases^7^. Subclusters M3 and M5 show the strongest transcriptional similarity to previously identified MIMS-iron and MIMS-foamy microglia at the CAL edge. d) Volcano plots of DEG in CSF M5 myeloid subcluster comparing PRL-positive versus PRL-negative (left panel) and PRL-high versus PRL-low cases (right panel). Corrected significance on y-axes represents -log_10_(adjusted p-value). e) PBMC myeloid L2 UMAP shown for PRL-positive versus PRL-negative comparison with total cell population randomly down-sampled equally per condition to 7000 cells (left panel). Volcano plots of DEG in the L2 CD14M4 and CD14M5 myeloid clusters comparing PRL-positive versus PRL-negative cells. CD14M4 shows upregulation of IFN-related genes while CD14M5 demonstrates upregulation of chemokines, including *CCL3, CCL4, CXCL8*, and pro-inflammatory cytokines, including *IL1B*. f) Transcriptional abundance of genes involved in the IFN pathway in the PBMC represented in a dot plot comparing cells from PRL-positive versus PRL-negative cases (left panel). On the right are violin plots showing enrichment of the average IFN module score (“IFN-signature”) across PRL categories in the myeloid and T/NK cells (right panel) in blood. g) Heatmap demonstrating differentially expressed proteins in CSF across PRL categories in patients. These are the sex- and age-adjusted abundances with abs[(log_2_(Fold Change (FC))] > 0.58 and *p* < 0.05. h) Boxplots showing abundance of IFN-related protein assays across patients with 0, 1-3, and ≥4 PRL (PRL categories: none, PRL-low, and PRL-high, respectively) in the serum (*\* p* ≤ *0.05*, *\*\* p* ≤ *0.01)*. i) Differential abundance of CXCL8 and CXCL10 in the CSF of patients across PRL categories (*\* p* ≤ *0.05)*.

Consistent with these results were the differentially abundant levels of soluble CD27 in CSF as measured by Olink (Figure 1E), considering that CD27 (a member of the TNF receptor superfamily) is a costimulatory molecule needed for long-term T-cell immune responses and is found on T, B, and NK cells^25–27^. Also elevated were the levels of neurofilament light-chain (NEFL), reflecting neuroaxonal damage, and glial fibrillary acidic protein (GFAP), suggestive of astrocytic activation and gliosis. Both have been suggested as biomarkers for MS^28–30^ in CSF, with serum NEFL and GFAP also being associated with chronic neuroinflammation in MS^30–32^. In addition, CSF GFAP has been associated with progression in MS^33^. We also found abundance of myeloid activation markers^34–36^, specifically CHIT1 (chitotriosidase) and TREM2 (triggering receptor expressed on myeloid cells 2), in MS CSF, consistent with prior studies showing enrichment of CHIT1 and TREM2 in CSF of patients with a chronic neuroinflammatory and neurodegenerative disease state^37,38^. Recent evidence also suggests that CHIT1 levels at the time of diagnosis predict disability progression in MS^39^. In addition, we found increased levels of GPB2 (an IFN-γ-inducible GTPase^40^) and IL17D (a pro-inflammatory cytokine associated with dysfunctional T_H_17 cells in autoimmune processes^41–44^), and enrichment of processes like metabolic dysregulation and cellular stress (PRTFDC1, CLUL1) and vascular or endothelial dysfunction (ANGPTL1)^45^. There was also upregulation of SERPINB5, a member of the serpin family^46^, which could be involved in the regulation of serine endopeptidases and hence immune responses.

In blood, we observed increased proportions of B-naïve, B-memory, and dnT-cells in MS^47,48^ (Extended Figure 2B, C). In addition, CD4-TCM cells were increased in frequency in MS. This is expected considering the immune dysregulation involved in MS, with antigen-exposed CD4 T-cells circulating between blood and lymphoid organs^47^. Interestingly, we also found decreased proportions of CD4-TEM and CD8-TEM cells in blood, which might be explained by effector cells trafficking into tissue and perhaps centrally into the CNS^47,49,50^. It is pertinent to note that both CSF and blood are highly dynamic rather than closed environments, making it difficult to fully account for the cell proportions in a cross-sectional sampling. Overall, even in untreated, inactive MS patients, CSF and blood exhibited altered cellular compositions relative to HV, with the CSF proteome affirming the presence of an inflammatory profile. Our findings largely parallel prior reports, validating our methodological approach and quality controls.

### Altered cellular composition of the CSF and blood immune compartments in patients with PRL

Considering that we observed CSF inflammation, with evidence of myeloid and adaptive immune activation, even in untreated and inactive MS, we next contrasted untreated and inactive cases with and without PRL. We used *in vivo* advanced MRI to detect PRL (Figure 2A), a radiological correlate of CAL, within 6 months of the acquisition of the analyzed blood and CSF samples (Figure 1A, 2A; Table 1). In the CSF, we discovered increased frequency of plasmablasts, B-intermediate, B-memory, Mono1, and Treg clusters in PRL-positive relative to PRL-negative cases, with decreased frequency of cDC2 and pDC (Figure 2B, Extended Figure 3A-B). The enrichment of B-cell and Mono1 clusters in the PRL-positive state could relate to the association between meningeal inflammation and CAL/PRL^4,51^.

**Figure 3:**
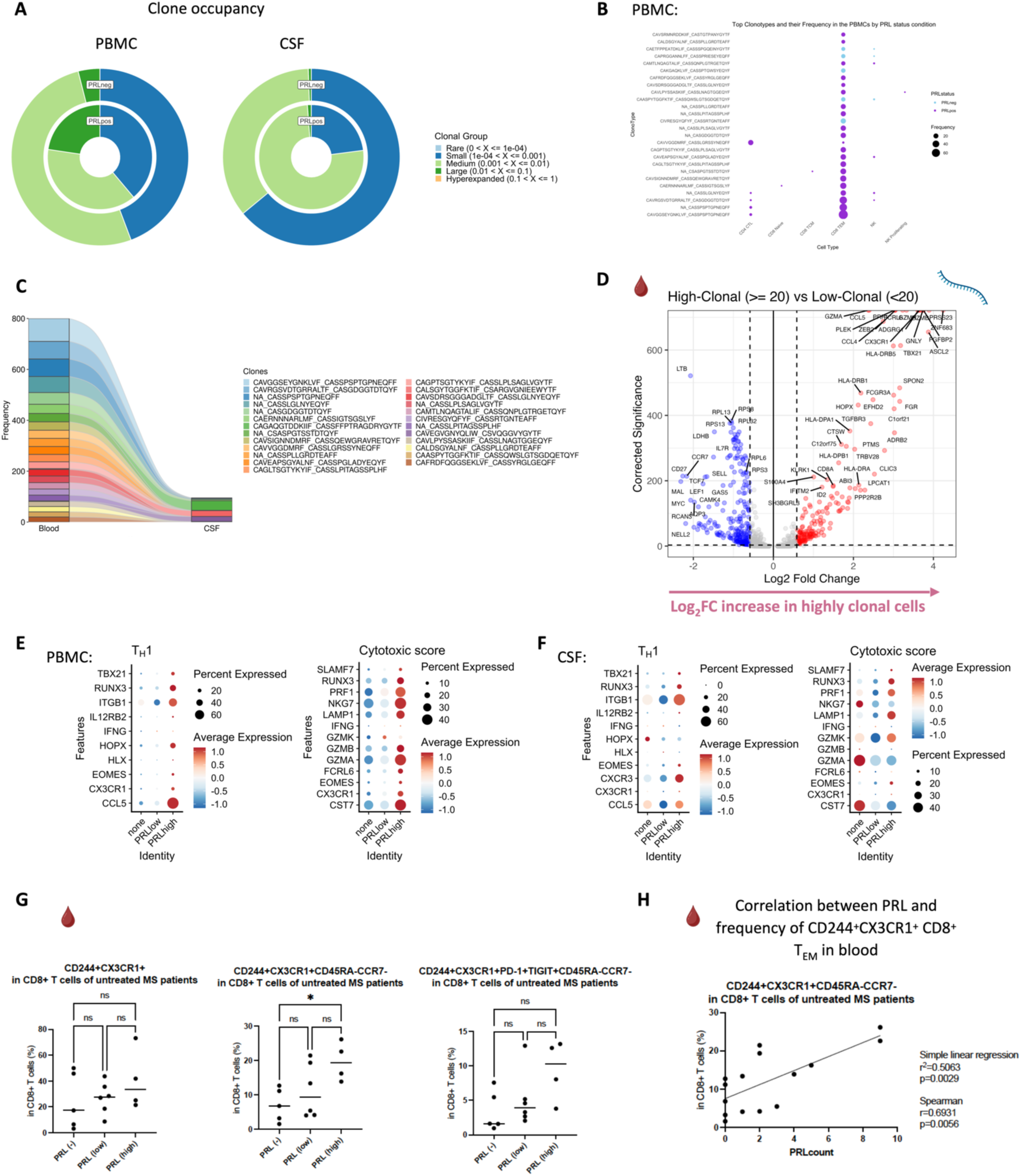
Clonally expanded CD8-TEM cells with T_H_1 effector and cytotoxic profiles mark chronic active MS. a) Donut plots representing the relative space occupied by clones at specific proportions in the blood and CSF, further split by PRL-positive versus PRL-negative status. b) Top 15 clonotypes with the CDR3-aa (complementarity determining regions 3 – amino acid) sequences shown for both the α- and β-chains, in the blood. The highly frequent clonotypes were largely found in CD8-TEM and CD4-CTL cells in PRL-positive cases. c) Top expanded clones in the cohort plotted across the blood and CSF compartments. d) Volcano plot showing DEG across high-clonal (clone size ≥20) versus low-clonal (<20) cells in the blood. e) T_H_1 and cytotoxicity-pertinent genes, and their expression shown as a dotplot in the T-lymphoid and NK cells in the blood across PRL categories: no PRL, 1-3 PRL (PRL low), and ≥ 4 PRL (PRL high). f) Expression of T_H_1 and cytotoxicity related genes illustrated by a dotplot in the T-lymphoid and NK cells in CSF across PRL categories. g) Percentage of CX3CR1^+^CD244^+^ (left panel), CD244^+^CX3CR1^+^CD45RA^-^CCR7^-^ (middle panel), and CD244^+^CX3CR1^+^PD-1^+^TIGIT^+^CD45RA^-^CCR7^-^ (right panel) cells relative to CD8^+^ T-cells in the blood across PRL-negative, PRL-low, and PRL-high patients. PRL-high cases have a higher frequency of CD244^+^CX3CR1^+^CD45RA^-^CCR7^-^ CD8^+^ T_EM_ cells relative to PRL-negative cases. There is a trend towards increasing frequency of CD244^+^CX3CR1^+^PD-1^+^TIGIT^+^CD45RA^-^CCR7^-^ CD8^+^ T subset with higher PRL burden. Statistical significance was assessed using Kruskal-Wallis’s test (* *p < 0.05*). h) Correlation of PRL burden with CD244^+^CX3CR1^+^PD-1^+^TIGIT^+^CD45RA^-^CCR7^-^ subset in CD8^+^ T-cells in the blood.

Since prior work had shown greater disability accumulation in patients with 4 or more PRL^5^, we stratified the PRL-positive cases into PRL-high (PRL ≥4) and PRL-low (PRL 1-3), contrasting the transcriptome and proteome. In CSF, PRL-high showed increased proportions of B-lineage clusters, plasmablasts, Treg, and CD8-TCM relative to PRL-low cases. Interestingly, we found both a relative and an absolute decrease in myeloid cells in PRL-high cases (Extended Figure 4A, B), potentially explainable by increased trafficking into CNS parenchyma^24^. The enrichment of B-cell clusters could contribute to the previously reported meningeal lymphoid aggregation associated with subpial inflammation and CAL^51^. In blood, we found increased frequency of CD4-CTL, CD8-TEM, and B-lineage cells in PRL-high (Extended Figure 4C), raising the possibility of increased lymphocyte trafficking into the CNS in these cases.

**Figure 4:**
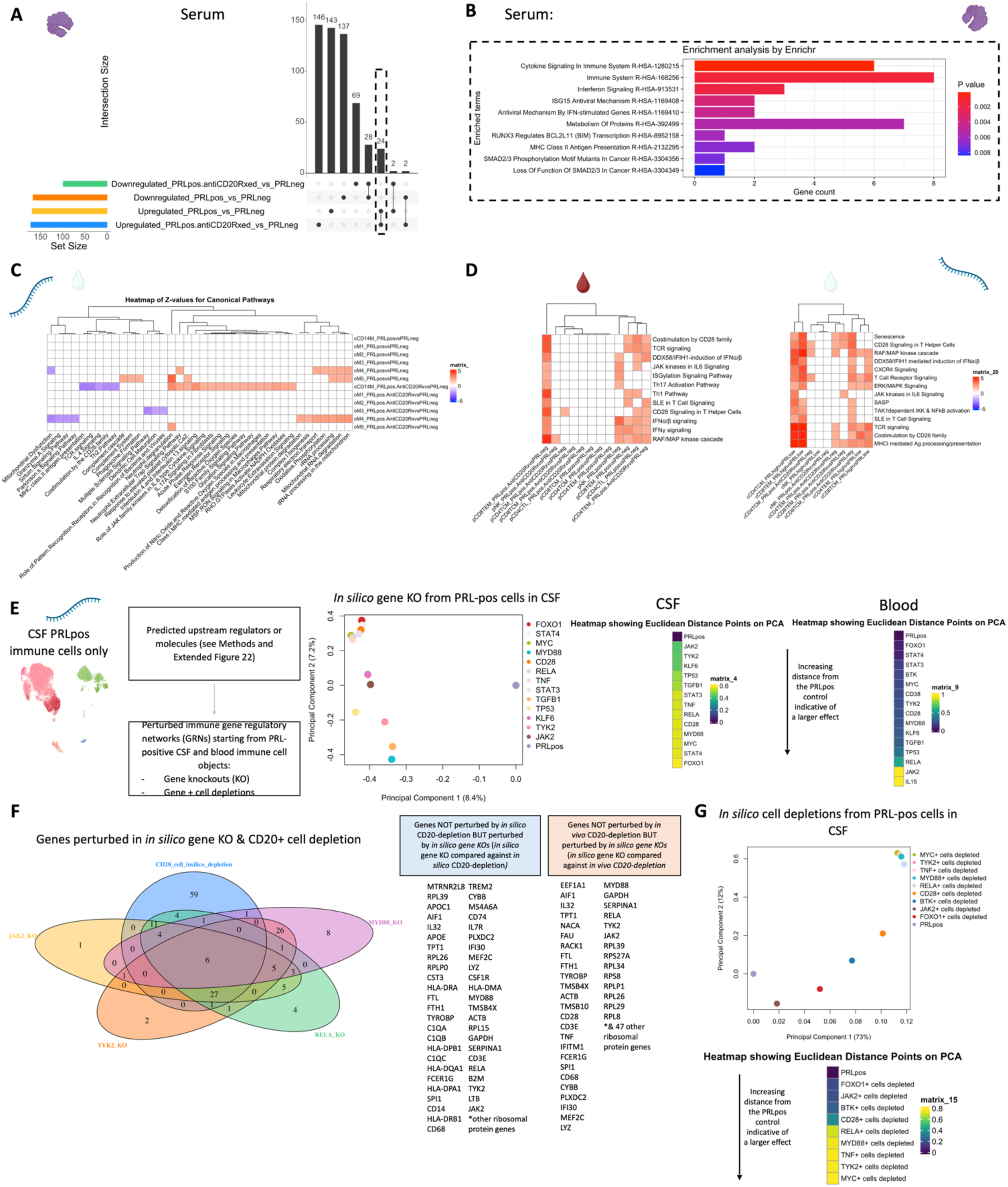
B-cell depletion fails to abrogate myeloid activation and CD8^+^ T-cell related cytotoxicity in PRL-positive cases, and new predicted targets. a) Upset plot demonstrating protein changes in the serum across B-cell-depleted versus PRL-negative and PRL-positive versus PRL-negative comparisons. The concordantly enriched or depleted proteins across the comparisons represent PRL-related pathology that remains unaddressed by B-cell depletion. b) Gene ontology (GO) terms for the concordantly enriched proteins in serum across the two comparisons: (1) B-cell-depleted versus PRL-negative; and (2) PRL-positive versus PRL-negative cases. c) Heatmap demonstrating Z-score enrichment of canonical pathways across CSF myeloid subclusters for DEG across B-cell-depleted (all PRL-positive) vs. PRL-negative and PRL-positive vs. PRL-negative comparisons. The heatmaps show concordant and discordant pathways between the two comparisons. Concordant pathways may reflect PRL-related pathology unaccounted for by B-cell depletion. See Extended Figure 17 for Z-score enrichment of pathways in CSF and peripheral lymphoid and myeloid cells for similar comparisons. d) Summary of Z-score enrichment of T-cell-related pathways across blood (left panel) and CSF lymphoid (right panel) cell clusters for the DEG across the comparisons shown (refer to Extended Figure 17D-G) e) From left to right: (1) Schematic showing the application of *in silico* perturbation strategy involving predicted upstream regulators driving PRL-pertinent immune responses (see Methods and Extended Figure 18), starting from the PRL-positive immune cells; (2) PCA plot and corresponding heatmap of the simulated distances from comparing the gene regulatory network (GRN) of the original PRL-positive state in CSF versus the GRN resulting from *in-silico* KO of the predicted regulator; and (3) heatmap of the simulated distances comparing the GRN of PRL-positive immune cells in blood vs. the same dataset following the gene knock-out (Extended Figure 19A shows the corresponding PCA plot). f) Venn diagram showing the significantly altered genes by *in silico* CD20-depletion, *in silico JAK2*, *MYD88*, *TYK2*, and *RELA* KOs. Accompanying are the list of genes that remain unaffected by *in silico* and *in vivo* anti-CD20-mediated B-cell depletion strategies, are contributory toward PRL-pathology, and might potentially be perturbed by the gene knock-out. Gene-set enrichment analysis (GSEA) of the significantly altered and ranked genes was performed (Supplementary File 7). g) *In silico* cell depletions from PRL-positive immune network of CSF, showing *TYK2-*, *MYD88-*, *RELA-*, and *JAK2-*positive cell-depletion strategies for abrogating PRL-related chronic neuroinflammation.

### Diverse PRL-associated myeloid activation signatures in CSF

We next performed differential expression analyses in various cell clusters to define PRL-pertinent changes. Since myeloid cells are prevalent in CAL^7^, we assessed if similar activation states existed in the myeloid cells of CSF. In the CSF M5 subcluster, differentially expressed genes (DEG) included those involved in classical and alternative complement pathways (*CFD*, *C1QA*, *C1QB*, *C1QC*, *C3AR1)^7^*, innate immune activation and response to lipopolysaccharide (*GRN*, *CLEC12A, GAA, SLC11A1, TNFRSF1B*, *CTSL*, *CD68*, *CTSB*), and VEGFA-VEGFR2 mediated signaling and vascular permeability, in cells from PRL-positive cases (Figure 2D, left panel). Contrasting cells from PRL-high versus PRL-low cases, M5-cluster upregulated genes are involved in IFN-γ and IL1 signaling (*JAK2*, *IL1R2*), myeloid regulation (*RUNX2*, *FOXP1, TET2*, *ZNF33B*), and G-protein signal transduction (*PDE7A*, *ADCY9*, *FARS2*) (Figure 2D, right panel).

In the M3 subcluster of CSF (Extended Figure 1A), we found upregulation of innate immune (*CLECL12A*, *CTSL*) and lysosomal-related genes (*ASAH1*, *LIPA*, *CTSL*) in cells from PRL-positive relative to PRL-negative cases (Extended Figure 5B). Another gene that was upregulated was *MTRNR2L8*, which was initially considered to be a pseudogene but more recently has been suggested to be a bioenergetic stress marker and negative regulator of apoptosis^52,53^. There was also upregulation of *SLC7A7* (a cationic amino acid transporter), which has been implicated in macrophages giving rise to microglia in development and in macrophages maintaining efferocytosis^54^. CSF myeloid subcluster transcriptome similarity analyses against AD brain microglia revealed M3 as the second most similar to microglia, after M5 (Extended Figure 3D). M3 and M5 also resembled the MIMS phenotype^7^, previously identified at the edge of CAL (Figure 2C). Pseudo-bulking myeloid cells to model for PRL categories, we found DEG across categories in a dose-response fashion, with the DEG across PRL-high relative to no-PRL significantly enriched for IFN and MAPK signaling, and DEG across PRL-high versus PRL-low cases enriched for transcription, PIP3/AKT activation, and intracellular signaling by second messengers (Extended Figure 5C).

### Synergistic proinflammatory Type-I and Type-II interferon signaling in chronic active MS

In blood myeloid cells (Figure 2E, left panel), contrasting PRL-positive versus PRL-negative, CD14M_4 subcluster upregulated genes including type-I IFN stimulated genes (ISGs) (*IFITM2*, *IFITM1*, *IFITM2*, *OAS1*, *MX2*, *MX1*, *IFI6*, *IRF7*, *ISG15*, *IFI44L*), and those involved in type-II IFN (*IFI6*, *ISG15*), and MAPK signaling (Figure 2E, middle panel)^55–58^. Concordantly, subcluster CD14M_5 demonstrated upregulation of genes involved in NFκB activation^59^, response to LPS and IL1 (*CXCL8*, *IRAK2*, *NLRP3*, *NFKB1, CCL4*, *CCL3*), and regulation of inflammatory response (*NFKBIA*, *IL1B*, *NFKBIZ*) (Figure 2E, right panel). Prediction of upstream regulators driving the DEG in myeloid cells of blood across PRL-positive versus PRL-negative yielded IFNA2, IFNG, TNF, and IL1B as top hits (Extended Figure 5D). Among other myeloid subclusters, similar pathways involving TLR-signaling, positive regulation of NFκB transcription factors (*PRKCB*), and regulation of MAPK signaling (*LYN*, *LRRK2*) were enriched in PRL-positive cases (Extended Figure 6A-B). Contrasting PRL-high versus PRL-low cases, myeloid cells in blood showed enrichment of similar pathways, including type-I and type-II IFN, strongly suggesting a pro-inflammatory environment (Extended Figure 7)^55,56,58^

Next, we plotted transcript abundances of IFN-pathway genes in the blood comparing PRL-positive to PRL-negative (Figure 2F, left panel; Extended Figure 8). We created an average module score comprising of *IFNG*, *JAK2*, *IFI6*, *IFITM2*, *IFITM3*, *IRF1*, *IRF9*, *ISG15*, *MX1*, *MX2*, *OAS1*, *STAT1*, and *IFNGR2* genes (“IFN-signature score”). We observed progressive enrichment of the IFN-signature score, highest in cells from PRL-high cases, in both myeloid and T/NK cells (Figure 2F, middle and right panels respectively). There was also increased IFN-signature in various myeloid and T/NK subclusters (Extended Figure 8A, top and bottom panels respectively) with higher PRL burden. In CSF myeloid cells, overall, there was no difference in the enrichment of IFN-signature, but T and NK cells in CSF showed increased average score at higher PRL burden (Extended Figure 8B). However, CSF myeloid subclusters M2, M3, and M4 demonstrated enrichment of the IFN score with higher PRL burden, as did CD4-TCM, CD4-TEM, and Treg among lymphoid cells (Extended Figure 8C). Plotting this score in a previously published postmortem tissue-based dataset^7^ showed clear enrichment of IFN-signature at the CAL edge versus other regions (Extended Figure 9A).

In parallel with the transcriptomic differences observed, we found increased soluble IFNGR2 protein in the serum of PRL-low relative to no-PRL patients (*p = 0.*02, Figure 2H). The IFN-γ-receptor is composed of two subunits, IFNGR1 and IFNGR2, which associate with JAK1 and JAK2, respectively^55^. We did not find a significant difference in the IFNGR1 levels across PRL-categories. Downstream of IFNα/β- and IFNγ-signaling, IFIT3 (*p = 0.006*) and EIF2AK2/PKR *(p = 0.01)* proteins were also more abundant in the serum of PRL-high versus no-PRL cases^58,60^. Consistent with transcriptional enrichment of MAPK signaling observed in blood, we found increased MAP4K5 in the serum of PRL-high versus PRL-low cases (*p = 0.03)*.

In CSF (Figure 2G, I), CXCL10, a chemokine produced by astrocytes^61^ and microglia^62^, and downstream of IFNγ-signaling, was significantly increased in PRL-high burden cases relative to those with none (*p* = 0.04), with a trend toward difference in PRL-high versus PRL-low cases (PRL 1-3; *p* = 0.09)^63,64^. Consistent with the transcriptional enrichment of p38/MAPK and PI3K/Akt signaling, CXCL8 was higher in the CSF of PRL-high versus PRL-low cases (*p* = 0.04) and those without PRL (*p* = 0.05)^65^. CXCL8 is a known potent pro-inflammatory chemokine with specificity toward neutrophils that is involved in trafficking as well as phagocytosis and other innate immune responses^66^. Additionally, we found increased abundance of proteins involved in extracellular matrix (ECM) proteolysis (ADAMTS8)^67^, neutrophil activation and migration (CD177)^68^, leukocyte adhesion to endothelium (MYOC)^69^, and previously defined biomarkers of neurodegeneration and ECM maintenance (EFEMP1)^70,71^ in high PRL burden cases (Figure 2G).

Interestingly, LRPAP1 was elevated in the CSF of high PRL burden (Figure 2G), with roles in regulating microglial activation and phagocytosis^72^ and in impacting type-I IFN responses^73^. The innate immune activation evident transcriptionally in PRL translated to increased levels of CLEC5A protein in CSF of PRL-positive cases. Prior evidence has shown involvement of CLEC5A in innate immune responses to viral/bacterial infections, with its activation leading to TNF and IL-6 production by myeloid cells^74^. We also found abundant SIGLEC5/CD170 in CSF of PRL cases; this has been shown to be an inhibitory receptor on myeloid cells, dampening innate immune responses and maintaining immune tolerance^75–77^. Collectively, these data show IFN-pathway enrichment across both blood and CSF, suggesting a proinflammatory state in communication across brain barriers.

### T-cell activation with dysregulated metabolism in PRL

Since we found evidence of IFN-signaling in PRL-positive cases, we next compared the adaptive immune cell clusters between PRL-positive versus PRL-negative cases. In CSF, CD8-TCM upregulated genes involved in translation and TCR signaling, while CD4-TCM cells showed upregulation of genes involved in phosphatidylinositol phosphate biosynthesis and negative regulation of insulin receptor signaling (*SOCS1*, *CISH*, *PIP4K2A*) in PRL-positive cases (Extended Figure 9B, C), reflecting a shift toward an effector phenotype by enabling a transition from glycolysis to fatty-acid oxidation and oxidative phosphorylation (OXPHOS)^78–81^. CD4-TEM cells upregulated genes involved in translation and cell cycle regulation (Extended Figure 9D). Contrasting PRL-high to PRL-low cases, CD4-TCM and CD8-TEM clusters shared 384 upregulated genes (Extended Figure 10A-B) enriched in TCR activation, OXPHOS, STING, and IFN signaling (*STAT1*, *JAK1*, *IFNAR1*, *IFNAR2*, *ST13*) (Extended Figure 10C). CD8-TEM cells upregulated genes involved in glycolysis (*PGK1*, *PFKL*), insulin/IGF and PI3K pathways (*GSK3B*, *PIK3CD*, *IGF2R*, *PIK3R5, PIK3R1, AKT3*) (Extended Figure 10C, D). These changes align with increased TCR signaling^82^ and metabolic programming^80,83^, favoring aerobic glycolysis – apart from OXPHOS genes (Warburg effect)^79,84^ – and PI3K/Akt/mTOR signaling^80,83,85–88^.

In blood, cells from PRL-positive compared to PRL-negative cases showed upregulation of IFN-stimulated and effector genes across cell types. CD4-TEM increased nitric oxide synthesis^89,90^ (Extended Figure 11). CD8-TEM upregulated T-cell differentiation (*ZNF683*, *TBX21*), and NK cells showed robust IFN responses (*IFNAR2*, *BST2*, *ISG20, IFITM2*, *STAT1*, *IRF7*, *ISG15*, *JAK1*) ^91,92^, supporting an effector and cytotoxic phenotype^55,89,90,93–95^ (Extended Figure 11). Note, *TBX21* is a key regulator of T_H_1 programming of CD4^+^ and CD8^+^ T-cells^93,94^. Similar transcriptional patterns were observed in all cell clusters, including type-I and type-II IFN responses in CD4-naïve cells (Extended Figure 12). Overall, we observed enhanced TCR activation, IFN signaling and increased anabolic state of adaptive T cells and cytotoxic NK cells in CSF and blood, associated with PRL^80,82,83^.

### Clonal expansion of CD8-TEM cells in cases with PRL

Since we found enrichment of IFN and TCR signaling in cases with PRL burden, we performed TCR sequencing in blood and CSF. For most of the analyses, we defined a clone as a group of cells that shares identical nucleotide sequences in both the α- and β-chains of their TCR, except when we used amino-acid sequences for the chains. We discovered large clonal groups to be occupying greater proportions in the “TCR space” in PRL-positive versus PRL-negative cases in both blood and CSF (Figure 3A). To integrate gene expression and TCR data, we created weighted-nearest-neighbor (WNN)-based UMAPs, depicting only those lymphoid cells that had a corresponding TCR sequence. We found PRL-positive specific cells that corresponded to CD8-TEM cells and were clonally expanded in both blood and CSF (Figure 3B; Extended Figure 13A-D). Notably, the most highly expanded clones were present in both blood and CSF, suggesting a continuum (Figure 3C). The top 15 frequent clonotypes and their CDR3-aa sequences mapped back onto CD8-TEM and CD4-CTL cells from the PRL-positive cases in blood (Figure 3B), and onto CD8-TEM cells from the PRL-positive cases in CSF (Extended Figure 13E). The top expanded clones in blood could be found in CSF for both PRL-positive and PRL-negative cases, and the converse was also true (Extended Figure 13F, 14A). There also appeared to be similar clones in CSF and blood for the same individual (Extended Figure 14B). Interestingly, there was little or no overlap of clonotypes across any pair of blood samples (Extended Figure 14C, left panel) or any pair of CSF samples (Extended Figure 14C, right panel) from different cases.

Using the annotateDB function, which is a collation of multiple databases, we attempted to use the sequences in our data to search for the likely epitopes associated with the TCR sequences (Extended Figure 14D, E) but could not determine the epitope association of the top clonotype using this approach alone. Among the most expanded clones in PRL-positive cases were the ones associated with epitopes like UL29 (CMV) and M1 (Influenza A). In the PRL-negative cases, there were clones in blood and CSF that were associated with EB4 (EBV) and BZLF1/NP177 (EBV and Influenza A) antigens, respectively. The TCR specificities for some EBV antigens have been characterized previously^96^. Importantly, most of the expanded clones were not found to be associated with any antigens predicted using annotateDB algorithm, so these additional antigenic targets remain unknown, but it remains possible, especially considering our finding of increased IFN signaling in both blood and CSF, that the target epitopes have a viral origin.

### Clonally expanded cytotoxic CD8^+^ T_EM_ cells as a correlate of PRL

We next stratified cells based on the clone size, with clonotype frequency ≥20 termed “high-clonal” and <20 “low-clonal.” We found upregulation of genes like *CCL4*, *CCL5*, *TBX21*, *GZMA*, *GZMB*, *HLA-DRB5*, *HLA-DRB1, HLA-DPB1*, *CX3CR1*, and *FCGR3A* in high-clonal cells in blood (Figure 3D), accompanied by downregulation of a naïve-like T-cell signature (*LEF1*, *CCR7*, *SELL*, *MYC*, *TCF7*). In CSF, high-clonal cells upregulated genes like *GZMH*, *CCL5*, *CXCR6*, *LGALS1*, *LAG3*, and TCR-α/β chain variable genes (Extended Figure 15A). This corresponds to enrichment of T_H_1, IFN-γ, TCR, MHC-II-restricted antigen presentation, and T-cell activation responses in the clonally expanded cells. Recently, a study in monozygotic twins discordant for MS identified dysregulated pro-inflammatory peripheral CD8^+^ T-cells in cotwins affected by MS when compared to healthy ones, which is consistent with our results^97^.

We next employed canonical gene expression programs^98^ to determine the likely phenotype of the highly expanded cells. We found overall enrichment of T_H_1 (*CCL5*, *CX3CR1*, *CXCR3*, *EOMES*, *HLX*, *HOPX*, *IFNG*, *IL12RB2*, *ITGB1*, *RUNX3*, *TBX21*) and cytotoxicity (*CST7*, *CX3CR1*, *EOMES*, *FCRL6*, *GZMA*, *GZMB*, *GZMK*, *IFNG*, *LAMP1*, *NKG7, PRF1, SLAMF7*) scores in high-clonal versus low-clonal cells in both blood and CSF (Extended Figure 15B, C). T_H_1 and cytotoxic score was also enriched in PRL-high versus PRL-low and no-PRL cases (note, all T lymphoid and NK cells from PBMC PRL object were used for this analysis) in blood (Figure 3E). In CSF (T lymphoid and NK cells from CSF PRL object were used), while T_H_1 score seemed overall enriched in PRL-high versus PRL-low and no-PRL cases, *GZMK* was enriched in PRL-high, and *GZMA* and *NKG7* genes were more abundant in no-PRL cases (Figure 3F).

Additionally, while effector memory score (*CD44*, *CD82*, *ITBG1*, *ITGB7*, *LGALS1*, *S100A10, S100A11*, *S1PR1, TCF7*) and exhaustion markers (*TOX, PDCD1, LAG3, CD200*) were enriched in all T lymphoid and NK cells from PRL-high cases (Extended Figure 15D, E) in CSF and blood, high-clonal cells were enriched in *TOX* and depleted in *TCF7* relative to low-clonal cells. This suggests that a high PRL burden, in addition to having highly expanded clonal cells, is also characterized by a reservoir of *TCF7^+^/TCF1^+^* cells that maintain formation of memory CD8^+^ T-cells. This is in line with prior reports^99^ suggesting that TCF1^-^ cells carry a cytotoxic signature.

To confirm these cytotoxic CD8^+^ T-cells (with exhaustion markers), we performed flow cytometry on PBMC. We discovered increased frequency of CD244^+^CX3CR1^+^CD45RA^-^CCR7^-^ CD8^+^ T cells in PRL-high relative to no-PRL cases (Figure 3G, middle panel), and there was an increasing trend in frequencies of CD244^+^CX3CR1^+^PD-1^+^TIGIT^+^CD45RA^-^ CCR7^-^ CD8^+^ T-cell subset with higher PRL burden (Figure 3G, right panel). We also found a strong correlation of PRL burden with CD244^+^CX3CR1^+^CD45RA^-^CCR7^-^ CD8^+^ T-cell subset in the blood (Figure 3H). This is suggestive of the presence of CD8 T_EM_ cells in high PRL burden cases, which adopt highly differentiated, cytotoxic profiles and acquire exhaustion markers, with some overlap with CD8 T_EMRA_ cells. We also found no significant difference in the frequency of this subset of cells in B-cell-depleted (all PRL-positive) relative to untreated, inactive MS cases (Extended Figure 15F). This is consistent with our transcriptomic and proteomic findings, wherein depletion of CD20^+^ B-cells did not abrogate CD8^+^ T-cell-related cytotoxicity in PRL-positive individuals. Therefore, we suggest that the blood CD244^+^CX3CR1^+^CD45RA^-^CCR7^-^ CD8^+^ T_EM_ subset is a cellular correlate of PRL burden, which should be investigated as a tool to measure response to therapeutic interventions for ameliorating progression in MS.

### Serum proteome is reflective of type-I IFN-mediated responses

In parallel with the transcriptional changes in the blood of PRL-positive cases, serum proteomics of PRL-positive versus PRL-negative cases identified 40 upregulated and 45 downregulated proteins (Extended Figure 16A). Pathway analyses showed enrichment of ISG15 and IFN-stimulated gene (ISG)-related immunomodulatory mechanisms in PRL-positive cases (Extended Figure 15B). Mirroring the transcriptomic findings in the blood, ISG15 is a ubiquitin-like protein induced by type-I IFN-signaling^100^ that leads to ISGylation of proteins^100–102^, conferring its antiviral properties^100^ but also acting as an extracellular cytokine modulating immune responses^103^. It has been proposed to induce NK proliferation and stimulate IFN-γ production^100,104,105^. Paralleling the transcript, EIF2AK2 was also differentially abundant in the serum of PRL-positive cases^91,92,106^. EIF2AK2 is a kinase that phosphorylates a serine residue (Ser 51) on eukaryotic initiation factor 2 α-subunit (eIF2α), resulting in transient suppression of protein translation^91,106^ and an antiviral effect. It is pertinent to note that EIF2AK2/PKR is activated by pro-inflammatory signaling (TNF-α, IL-1, IFN-γ)^107^ and then activates inflammatory pathways, including NFκB signaling^106^, as evident transcriptionally in our study. Additionally, EIF2AK2/PKR has been implicated in inflammasome activation^108^. IFIT3, induced by type-I IFN signaling, was also abundant in the serum of PRL-positive cases and contributes to immune modulation^109^ and antiviral defenses^110^. Predicted molecules upstream of the upregulated proteins in the serum of PRL-positive cases included EIF2AK2 and IL15 (Extended Figure 16B). Overall, these findings suggest IFN signaling with subsequent immunomodulation evident on both transcriptional and proteomic levels.

### Serum and CSF proteomic networks reveal PRL signatures that remain unaffected by CD20 depletion

As B-cell depletion does not resolve PRL over a ∼2-year period^111^, we reasoned that comparing untreated and inactive PRL-positive and PRL-negative cases with PRL-positive CD20-depleted cases would allow discovery of genes, proteins, and pathways that may be critical for CAL persistence. A serum proteomic network analysis (Extended Figure 16C) uncovered modules with concordant and discordant correlations. Of the modules uncovered by this analysis, sM11, sM15, and sM26 were associated with PRL presence regardless of treatment status. Module sM11 (*r = 0.55* with PRL, *p = 0.01*) included proteins involved in apoptosis (TNFRSF10A, BL2L1), ERAD pathway (DNAJB2, TRIM25), and regulation of proteosome-dependent catabolism (DNAJB2, PARK7). Module sM15 (*r* = *0.51*, *p* = *0.03*) comprised proteins involved in mTOR signaling (EIF4B, EIF4E, AKT1S1), cellular response to stress (DNAJB1, DNAJA4, TXNRD1, HSPA1A), and MAPK1/MAPK3 signaling (PSMD9, MAP2K1) responses. Module sM26 also correlated positively with PRL presence (*r = 0.58*; *p = 0.01*), with module hub members, including STAT2, STAT5B, HARS1, and PQBP1 (Supplementary File 5, Extended Figure 16C), being involved in responses to IFN-γ (IFNGR2, GBP1), tRNA-aminoacylation for protein translation (YARS1, FARSA, HARS1), and double-stranded DNA break repair (SETMAR, MORF4L1, PAXX).

Interestingly, module sM18 concordantly correlated and modeled positively and significantly with both CD20-depletion (*r = 0.68*, *p* = *0.02*; see Extended Figure 13C and Methods for details regarding EP) and PRL (*r = 0.69*, *p = 0.001*). This could be inferred as a set of proteins that remain unmodulated despite CD20-depletion in PRL-positive cases. Module members of sM18 include proteins induced by IFN-γ and involved in antigen processing (IFI30), oxidative stress (IMMT), ubiquitin-mediated proteolysis (UBE2B, CDC26), and iron transport (SCARA5). Previously, SCARA5 was found to be an antigenic target of intrathecal IgM in MS^112^.

In the CSF, contrasting cell proportions across CD20-depleted (all PRL-positive) versus PRL-positive cases, we found depletion of B-memory and B-intermediate cells in CSF, which is expected considering the effect of B-cell depletion (Extended Figure 17A). Consistent with recent results^113^, we also found relative enrichment of CD14-Mono and Mono2 clusters, and relative enrichment of CD4-naïve cells, in the CSF of B-cell depleted cases. Using the CSF proteome, we created a network involving the PRL-positive, PRL-negative, and CD20-depleted (all PRL-positive) cases (Extended Figure 17B). Applying a similar approach as we did for the serum proteome, we found a module (cM7) that correlated positively with PRL presence, with proteins involved in receptor-mediated endocytosis (MSR1, LDLR) and myeloid differentiation and activation (CXCL8, IL15, LGALS9). In addition, module cM8 correlated positively with PRL presence and B-cell depletion, suggestive of proteins unaffected by B-cell depletion (Extended Figure 17B, Supplementary File 6). This module enriched for positive regulation of protein phosphorylation (CD40, IL6, CSF1, PLAUR, LIF, CHI3L1) and MAPK cascade (TDGF1, CRKL, LIF), among others.

Overall, these analyses, which are consistent with our transcriptomic data, strongly suggest that IFN-mediated inflammation, which is associated with PRL and chronic neuroinflammation, persists even in the setting of a treatment that nearly completely abolishes new inflammatory lesions and relapses in MS but has only limited impact on clinical progression. These analyses also point to serum and CSF biomarkers that could potentially be used to track resolution of chronic active lesions, as the paramagnetic rim only reports on the presence or absence of iron within microglia at the lesion rim.

### Depletion of CD20+ B cells fails to address PRL pathology

Applying a similar strategy as above, comparing B-cell depleted (all PRL-positive) versus PRL-negative and PRL-positive versus PRL-negative cases yielded 24 and 28 proteins in the serum that were concordantly enriched and depleted across both comparisons, respectively (Figure 4A, dashed-lines show the proteins enriched across both comparisons; Extended Figure 17C; note the fold-change for this particular analysis was set as 1.25 instead of the usual 1.5). The enriched proteins are involved in cytokine signaling (SMAD3, ANXA2, UBE2L6, IFI30, RELB), IFN and ISG15-related pathways (TPR, UBE2L6, IFI30), and MHC-II antigen presentation (KIF22, IFI30) among others (Figure 4B). Among the depleted proteins were those involved in Wnt-signaling (CDH15, SMARCA2) and cell-cell adhesion (CDH15, CXADR, ALCAM). In the CSF, we found one protein, ULBP2, which was depleted across both comparisons; no proteins were enriched across both comparisons.

Transcriptionally, concordant pathways across B-cell-depleted (all PRL-positive) versus PRL-negative and PRL-positive versus PRL-negative comparisons identified additional PRL-related pathways that remain unaddressed with B-cell depletion. In the myeloid compartment of CSF, we observed concordant enrichment of OXPHOS (*MT-CO1, MT-ND1, MT-ND2, MT-CYB*), ETC, and ATP synthesis mediated by the M4 subcluster (Figure 4C). CD14M and M5 subclusters enriched in neutrophilic degranulation across the above-mentioned comparisons, representing a possible pathway pertaining to PRL-positive pathology that remains uncorrected with CD20-depletion, despite the fact that neutrophils are rarely observed in the context of MS (Figure 4C).

We undertook a similar approach in the transcriptomic dissection of peripheral lymphoid and NK cells, finding a multitude of pathways in the overall peripheral cell paradigm (Extended Figure 17D). Filtering for the concordant pathways across the two comparisons (Extended Figure 17E), we observed CD8-TEM cells as the primary correlate of PRL-related pathology that remains unaddressed with B-cell-depletion (Extended Figure 17E). Related pathways included JAK/STAT, IFN-α/β, and IFN-γ (*JAK1*, *JUN*, *STATs*, *IRF1*), T_H_1 (*TBX21*, *RUNX3*, *STAT3/4*), TCR (*CD3E*, *CD3G, ITK, ZAP70*), cGAS-STING (*IRF1, STING1*), ISGylation, PI3K/AKT (*PIK3K1*, *ITGB1*/2, *ITGAL*), EKR/MAPK (*MAP2K1/2*, *ERK1/2*), TNFR1 and TNFR2 signaling, and neddylation, among others. Neddylation has been previously suggested as a possible therapeutic target based on a study in EAE^114^. There was also enrichment of pathways involved in OXPHOS, iron transport, and senescence.

In the CSF, since PRL-positive versus PRL-negative showed relatively a smaller number of DEG in lymphoid subsets, we compared PRL-high versus PRL-low and B-cell depleted (all PRL-positive) versus PRL-negative cases. We identified concordant pathways likely driven by CD8-TCM and CD8-TEM cells, with similar mechanisms as above, including CD28 signaling in lymphocytes, IFN, IL7, ERK/MAPK, PIP3/AKT signaling, IL15 production, and RUNX3-related transcriptional regulation (Extended Figure 17G). Overall, these analyses suggest the critical role played by myeloid and CD8^+^ T-cells (Figure 4C, D) in mediating chronic neuroinflammation that remains unaddressed with B-cell depletion.

### *In silico* deletions to help predict potential therapeutic targets for ameliorating PRL-related inflammation

Since we found PRL-relevant pathways that remain unperturbed with *in vivo* B-cell depletion, we next screened and filtered for the top upstream regulatory molecules driving PRL-related pathways (Extended Figure 18A-E). Using a machine learning (ML)-based tool, we computed a gene regulatory network (GRN) initially made from PRL-positive cells in CSF or blood, comparing it with GRN versions following depletion of a gene of interest^115^ (top “regulators”) or of specific gene-positive cell populations^116^ (Figure 4E-G, Extended Figure 18A). Comparing the virtual deletions with one another and to the control PRL-positive condition (Figure 4E; Extended Figure 19A), we shortlisted virtual deletions of *JAK2*, *MYD88*, *TYK2*, and *RELA* to be strongly affecting the PRL-positive GRN in both CSF and blood. In all virtual deletions of the regulators, the most significantly perturbed genes were involved in IFN-γ and IFN-α response, complement, PI3K/Akt/mTOR, IL6-JAK-STAT3, and TNF signaling via NFκB (Supplementary File 7).

Subsequently, we assessed how the significantly impacted genes resulting from virtual deletions compared with *in vivo* and *in silico* CD20-depletion, identifying gene lists not perturbed by anti-CD20-mediated B-cell depletion but predicted to be affected by targeting PRL-relevant regulatory genes (Figure 4F; Extended Figure 19B). These lists included genes involved in microglia/macrophage activation, complement, IFN pathway, iron metabolism, MHC-II antigen presentation, and ribosomal biogenesis (Figure 4F, right panel). Virtual depletion of cells expressing *TYK2*, *MYD88*, *TNF*, and *MYC* in the CSF most affected the GRN of PRL-positive cells, with smaller effects predicted by depletion of *JAK2* and *BTK* (Figure 4G). Employing this machine-learning framework in PRL-positive cells, we were thus able to identify putative targets, several of which are associated with either approved therapeutics or therapeutics in development, for attenuating PRL-related inflammation.

## Discussion

Impeding or halting progression in MS remains a pressing problem. Here, we used a combination of scRNAseq, scTCRseq, and proteomics to identify the CSF and blood immune correlates of chronic neuroinflammation in MS, defined by the presence of PRL as a radiological biomarker associated with disability accumulation and progression. In elucidating the immune correlates of PRL, we used freshly isolated samples from a cohort of patients who were clinically and radiologically inactive (no new lesions or clinical relapses) and untreated for at least 6 months (Table 1), providing an opportunity to probe into chronic active MS-related pathology in an unperturbed setting. Our data therefore represent a unique single-cell transcriptomic resource of CSF and CSF-matched PBMC in radiologically well characterized MS cases representative of those with a currently unmet therapeutic need. To begin to address this need, we leveraged data from B-cell-depleted patients, all of whom had PRL, to inform PRL-related mechanisms that remain unaddressed with B-cell depletion. Currently, no disease modifying therapies — including Bruton’s tyrosine kinase (BTK) inhibitors — are known to resolve PRL over a ∼2 year period^111,117^. Using an *in silico* approach, we identified putative regulatory pathways that, when modulated, are predicted to attenuate PRL-related pathology.

Via transcriptome analysis, we discovered enrichment of IFN-signature – type-I and type-II, with the prior being predominant – and JAK/STAT signaling in the blood and CSF of PRL-positive relative to PRL-negative patients. We also found concomitant PI3K-AKT, MAPK, and NFκB signaling in myeloid and adaptive immune cells, as well as a complement signature in CSF myeloid cells, explicating prior results^7^. This signaling was accompanied by TCR signaling and activation of adaptive immune cells in the blood and CSF of patients with PRL. Consistent with the increased metabolic demands requiring adaptive immune activation, we observed metabolic reprogramming toward enrichment of glycolytic and oxidative phosphorylation gene programs^79,80,83,84,118,119^. Our proteomic data corroborated the transcriptomic findings.

Aligning with TCR signaling, we discovered clonal expansion of CD8-TEM and CD4-CTL cells in the blood, and CD8-TEM cells in the CSF, in PRL-positive cases. Dichotomizing the clonally expanded cells based on clone size, we found that highly expanded cells were enriched in T_H_1 and cytotoxicity-pertinent signatures relative to those with no to minimal expansion. This T_H_1 and cytotoxicity signature was also enriched in the lymphoid cells of both CSF and blood in PRL-high patients. Together, these results suggest that IFN-driven cytotoxic CD8^+^ T-effector responses and clonal expansion are important for maintaining PRL-related inflammation, and that this pathway may be targetable therapeutically.

In addition to the above analyses, we also incorporated data from individuals treated with anti-CD20 antibody-mediated B-cell depletion, all of whom had PRL, to dissect pathology pertinent to PRL that remains unabrogated with this treatment modality, which is generally extremely effective in preventing new CNS inflammation in MS but ineffective in resolving PRL^111^ and limiting PRL-associated disability progression. Using both transcriptomic and proteomic network approaches, we identified persistent myeloid activation and CD8^+^ T-effector cytotoxicity mediated by several key pathways, including type-I and type-II IFN, JAK/STAT, ISGylation, EIF2, cGAS-STING, PI3K-Akt-mTOR signaling, senescence, autophagy, and MHC-II antigen presentation, as being unaddressed by B-cell depletion.

Based on the highly expanded cells of PRL-positive patients, we identified a peripheral cellular correlate of PRL, identifiable as the CD244^+^CX3CR1^+^CD45RA^-^CCR7^-^CD8^+^ T_EM_ cells. These highly clonal cells exhibited T_H_1 and cytotoxic features. In addition, this subset harbored high expression of GZMK. Pending validation in a larger cohort, we propose this as a possible immune cellular correlate of chronic inflammation-mediated progression in MS. Prior reports have identified such cytotoxic CD8^+^ T effector cells in the context of neurodegenerative diseases including Alzheimer’s disease (AD)^120^ and amyotrophic lateral sclerosis^121^. Importantly, we did not find this subset of CD8^+^ T effectors to be diminished in frequency with CD20-depletion (Extended Figure 15F).

It is important to note that our study involves immune cells within the CSF and blood compartments, and while immune cells play a significant role in MS progression, as suggested previously^2,7^ and by our data, the role of CNS-resident glial cells in progression could not be considered here. However, in work involving tauopathy mice, microglial and T cell mediated responses were implicated in tau-mediated neurodegeneration^122^. Additionally, recent work in an AD mouse model (5xFAD) showed evidence of CD8-mediated microglial activation via IFN-γ, resulting in myelin damage^123,124^. Considering the enrichment of IFN-γ-signature and upregulation of JAK2 in the M5 cluster of CSF myeloid cells, in parallel with increased abundance of proteins downstream of type-I and type-II IFN-signaling in chronic active MS (CXCL10 in CSF; IFIT3, EIF2AK2, MAP4K5, and STAT2 in serum), it is plausible that the CD8-TEM cells described in our data traffic to the CNS and influence resident glial cells to propagate neuroinflammation and neurodegeneration. This is particularly relevant since the identified CD8^+^ T-cell subset in our study carries a T_H_1 phenotype and highly expresses GZMK, which was recently reported to be involved in directly activating the complement system, distinct from the usual known activation pathways^125^. This is also in line with the observation of infiltration of T-cells impacting neurogenic niches via IFN-γ^126^. Thus, our data provide rationale for therapeutic targeting of IFN and JAK1/2, and potentially GZMK, for amelioration of PRL-associated inflammation in MS. In the *in silico* study, although the simulated effect of JAK2^+^ cell depletion was smaller than depletion of some other cell types (Figure 4G), it is worth noting that this was only considered in CSF, and given that T-cells can traffic into the CNS, the impact could be larger when both CSF and blood are impacted.

It is important to note that while we identified clonotypes enriched in PRL-positive and PRL-negative cases, listing the top frequent clonotypes, and predicted epitope associations, we did not perform experiments screening for antigen specificity. Importantly, we found no to minimal overlap in clonotypes across PRL-positive and PRL-negative cases in CSF or blood (Extended Figure 14B). Although some of the predicted antigen specificities were shared, this suggests an element of redundancy in CDR3-defining regions of TCRs, a suggestion not probed here. One implication is that PRL presence could be linked to an increased variety of epitope exposures with aging. This follows from the association of PRL and non-relapse-related progression in MS with age^2^ and the presence of clonally expanded T-cells in aged brains^123,126^. Along these lines, we found senescence-related modules as contributing to PRL pathology but unaddressed by B-cell depletion. It is important to note, though, that while we provide a resource of newly identified clonotypes in MS, some of which may be related to progression, antigen specificities for these clonotypes remain to be determined.

In summary, our work highlights the significance of myeloid and CD8^+^ T-cell activation programs in maintaining and possibly propagating chronic neuroinflammation in progressive MS. This inflammatory signature is enriched even in untreated and clinically/radiologically inactive patients with PRL. We highlight the role of IFN-mediated signaling (both type-I and type-II), along-with MAPK and TCR immune signaling cascades, providing mechanistic insights that could be leveraged for therapeutic purposes. Although our data suggest a framework (Extended Figure 19C) with a continuum of immune dysregulation between blood and CSF, there is nonetheless evidence of compartmentalization of the CSF. We have also deciphered the TCR immune repertoire of chronic active MS, and it will be interesting to know the antigen-specificities of the TCRs identified. Finally, we propose using CD244^+^CX3CR1^+^CD45RA^-^CCR7^-^CD8^+^ T or CD244^+^CX3CR1^+^CD8^+^ T_EM_ cells as a cellular correlate of PRL and in measuring response to therapies targeting PRL.

## Methods

### Cohort and metadata

The transcriptomic, proteomic, imaging, clinical and demographic data for this cohort was collected following the approval of the institutional review board (IRB) and after written, informed consent was obtained as part of the National Institute of Neurological Disorders and Stroke’s “Evaluation of Progression in Multiple Sclerosis by Magnetic Resonance imaging” protocol (NCT00001248). In this protocol, participants with MS or MS mimics are seen in the clinic and undergo imaging and sample collection, including CSF and blood. The advanced imaging is performed using either 7T MRI, or both. Experienced MS clinicians collected the clinical history and determined EDSS.

We performed single-cell transcriptomics (scRNAseq) on CSF cells and CSF-matched PBMCs from 34 MS cases and 5 healthy volunteers (HV). This included 6 MS cases who were treated with CD20-depleting agent (ocrelizumab) and all had paramagnetic rim lesions (PRL-positive) at the time of sample acquisition. Seventeen MS patients were untreated and radiologically inactive for at least 6 months (but in most cases longer than that; see Table 1) from the timepoint of sample acquisition and MRI, with 10 being PRL-positive and 7 PRL-negative (Figure 2A). Our primary comparisons were: (1) PRL-positive vs, PRL-negative patients; (2) CD20-depleted versus PRL-positive vs. PRL-negative patients; and (3) untreated and inactive MS patients vs. HV. Additionally, gene expression data from 11 MS patients who received other treatments and/or were radiologically active were used for creating the overall CSF, PBMC, and combined (CSF+PBMC) objects, but their metadata are not analyzed in this paper. Relevant subsets for these primary and other comparisons were derived (subsetted) from the main Seurat objects. Demographics and metadata for subjects, including those with Olink proteomic data and TCR sequencing, are summarized and shown in Table 1 and Supplementary File 1, respectively. The metadata for the Olink analysis involving untreated and inactive MS vs. HV is shown in the second sheet in Supplementary File 1.

### In vivo MRI acquisition and PRL determination

The in vivo MRI acquisition for participants involved using a Siemens Magnetom 7T scanner equipped with a birdcage-type transmit coil and a 32-channel receive coil. The imaging protocol included a pre-contrast, high resolution 3-dimensional (3D) gradient dual-echo sequence for acquiring T2*-weighted (T2*) and phase contrast images (repetition time [TR]: 74 milliseconds; echo times [TE]: 18, 29.5, 50.0 and 52.4 milliseconds; flip angle [FA]: 10°; acquisition time [AT]: 11 minutes 50 seconds per 30 mm slab; 0.5 mm isotropic resolution). Axial T2*-weighted images were motion- and B0-corrected, and magnitude and unwrapped filtered phase images were obtained. Additionally, MP2RAGE was acquired and resampled to 0.5 mm, and a T2-weighted fluid-attenuated inversion recovery (FLAIR) sequence was also obtained for visualization of white matter lesions in some cases.

PRL were determined by an experienced neuroradiologist based on the NAIMS consensus criteria^127^. In summary, the lesion must be identifiable on T2* or FLAIR and must have a discrete rim with a paramagnetic signal on susceptibility-sensitive sequences including filtered phase. This paramagnetic signal is observed as a hypointensity, which should be continuous through at least two-thirds of the outer edge of the white matter portion of the lesion (excluding any cortical or ependymal border), and the lesion should have a relatively isointense center. DSR rated all the PRL in our cohort.

### Sample acquisition, processing and loading onto Chromium platform

CSF and CSF-matched blood samples were obtained from each participant enrolled in the study. A lumbar puncture (LP) was performed either under fluoroscopy or at the bedside, and approximately 20-30 ml of CSF was obtained. CSF cells were subsequently collected within an hour by centrifugation at 400 x g for 10 min at 4 °C, and the supernatant was stored for Olink-based proteomics. PBMC were isolated from heparinized blood samples from each participant using lymphocyte separation medium and density gradient centrifugation, as described previously^128^. Fresh CSF cells and PBMC were suspended in PBS with 0.04% BSA, and viable cells were quantified on a Muse® cell analyzer (Millipore). Immediately after isolation, CSF cells and PBMC were then freshly loaded onto a Single Cell G Chip and run using Next GEM Single Cell 5’ Library and Gel bead kit v1.1 for cell capture and barcoding, targeting 5000 cells according to manufacturer’s instructions. Reverse transcription, cDNA amplification, and library construction were performed per protocol. For analyses of T-cell receptors (TCR), full-length V(D)J segments were enriched from cDNA by PCR amplification with primers specific to the TCR region, using single-cell V(D)J enrichment kits per instructions. Sequencing of both the gene expression and TCR-libraries was performed using Illumina’s NovaSeq sequencer. Serum was isolated after centrifugation of blood at 2000 rpm for 10 minutes, then collected and aliquoted into cryotubes and stored at -80 °C until use for proteomics.

### Single-cell gene expression data analyses pipeline

For each CSF and PBMC sample, library demultiplexing, barcode processing, FASTQ file generation, and gene alignment to the human genome (GRCh38 – 2020 – A) was performed using the CellRanger v3.1.0 software. This generated a cell barcode-to-gene matrix, which was used for further downstream analyses. Seurat v4.1.3 was used to create a Seurat object for individual samples using the CreateSeuratObject() function in R v4.3.1.

Following Seurat guidelines, preprocessing and quality control (QC) involved generating metrics using the PercentageFeatureSet() function and keeping cells with 200 – 5000 genes and <10% of the total counts associated with mitochondrial genes. The filtered object was then log-normalized using NormalizeData() function (LogNormalized method – normalizing the gene expression for each cell by the total expression, scaling it by a factor of 10,000, and log-transforming the results). The top 2000 variable genes were calculated using FindVariableFeatures(), and the object was then subjected to scaling using ScaleData() prior to linear dimension reduction (principal component analysis (PCA)). PCA was then performed on the object using RunPCA() function, and the top 20 principal components (PCs) were implemented for unsupervised clustering. Selection of the 20 PCs for each sample was based on elbow plot inspection, created using ElbowPlot(). FindNeighbors() function was used to construct the k-nearest neighbors graph based on the Euclidean distance in PCA space, and then FindClusters() function applied the Louvain algorithm to iteratively group cells at a resolution of 0.5. Resolutions were inspected over a range via clustree(), and 0.5 was deemed the ideal resolution, avoiding over-clustering for each individual sample. We subsequently implemented the UMAP for cluster visualization for each object using RunUMAP(). DoubletFinder v3 was then implemented for each object to estimate and remove putative doublets from each sample. The function find.pK() was used to select the optimal pK threshold for doublet detection by maximizing the bimodality coefficient metric, which optimally separates singlets from doublets^129^.

### Clustering and visualization

A total of 78 individual Seurat objects were created from the 39 CSF and 39 PBMC samples. These were then merged, integrated, and harmonized to create an overall CSF (only CSF samples), PBMC (only PBMC samples), and a combined (CSF+PBMC) object. For the combined object, all samples were randomly down-sampled to 1000 cells each if the sample had greater than 1000 cells, otherwise the original number of cells were retained. Similar processing pipelines were implemented for all objects, and hierarchical level-1 (L1) and level-2 (L2) analyses were employed for the scRNAseq dataset to help identify subclusters and result interpretations for downstream analyses. To create the object(s) identified above, the pertinent individual Seurat objects were merged using merge() function and the QC metrics observed (Figure S1A-J); this included filtering out any cells with: (1) number of unique genes identified greater than 2 standard-deviations (SD) above the mean and fewer than 250; (2) total UMI (unique molecular identifier) counts greater than 2 SD above the mean and less than 500; and (3) greater than 2% mitochondrial counts. This was followed by yet another putative removal of doublet cells per sample using computeDoubletDensity() function of the scran package, with the threshold here set at 2 SD above the mean of doublet score values within each sample^130^.

The merged object was then log-normalized and scaled. The top 3000 variable genes calculated by Seurat were used in the PCA. Initially, 200 PCs were computed. An elbow plot was then generated, as above, and inspected. Based on the elbow plot, 75 PCs were used for Harmony v1.2 to integrate the different samples over the orig.ident variable since there was not much variance explained in the expression data by more than 75 PCs^131^. The Harmony embeddings were used for Seurat to learn the UMAP and find clusters at multiple resolutions. The clustering at incremental resolutions ranging from 0.04 – 3.5 were inspected along-with clustree(), to decide a reasonable resolution for the initial clustering, such that over- or under-clustering was avoided. After choosing an appropriate resolution for initial clustering, doublet clusters were identified to be removed using the findDoubletClusters() function. Here, the doublet clusters were identified using isOutlier(), with parameters “type” and “log” set as lower and TRUE respectively. In all our cases (CSF, PBMC, and the combined object), we found no doublet clusters. However, computeDoubletDensity() function was again used to compute a “doublet score,” and cells with a doublet score of >2 SD above the mean score were removed as doublets. We then applied the FindNeighbors() function with 75 PCs, and Harmony reduction, followed by clustering at multiple resolutions for the final clustering. The clustering was evaluated at multiple incremental resolutions ranging from 0.04 – 3.5 along-with clustree(). The final resolutions chosen for CSF L1 and PBMC L1 objects were 0.9 and 0.4, respectively.

Following this, we performed cluster pruning, using Z-score-based outlier threshold selection, to form clean clusters. In cluster pruning, cells were subsetted by cluster and Z-scores calculated using the UMAP1 and UMAP2 dimension values separately. Next, cells observed to be outliers with Z-score greater than the thresholds defined by the outermost limits of the main cluster were discarded. The resulting Seurat objects were saved as L1 objects for CSF and PBMC (Extended Figure 1A, B) and combined CSF and PBMC samples (Figure S8A-D). The L1 PBMC object was annotated using Azimuth by executing RunAzimuth() function^15^. For the L1 CSF object, the cluster annotation was a combination of Azimuth and manual annotation based on prior literature, considering that CSF and PBMC are different tissues with varying immune cell types^132^. Manual assignments of cell-type identity to clusters were based on canonical cell-lineage markers. For the combined object, we show L2 annotations mapped onto the L1 object (Figure S8C).

FindAllMarkers() was used to find the markers specific for the annotated clusters. A cluster-defining gene was defined as a gene significantly expressed (p-value adjusted for multiple comparisons, adj_p < 0.05) with an average >0.7-fold difference (log-scale) of expression between the cluster of interest (ident.1) vs. the rest of the cells, while the average expression in the rest of the cells (ident.2) is required to be ≤30%. DotPlot() was then subsequently used to visualize the top 2 cell markers for each L1 annotation (Extended Figure 1C) and the top 5 markers for each L1 (Figure S3B,F) and L2 annotation mapped onto L1 object (Figure S3D,H).

### Level 2 subclustering (L2) analyses

Based on the L1 cell annotations, subclusterings were performed on myeloid lineage cells, T/NK-related cells, and B-lineage cells (dashed green, pink, and blue lines respectively) for both CSF and PBMC. This was performed using the same functions, methods, and unsupervised algorithm as described above, except that the number of PCs used for Harmony integration over samples, and subsequent clustering of CSF L2 myeloid, lymphoid (T-cell and NK-related lineages), and B-lymphoid objects (Extended Figure 1A), were 60, 75, and 30, respectively, while 75, 75, and 20 PCs were used for PBMC L2 objects (Extended Figure 1B). There was no cluster pruning performed for the L2 Seurat objects. Subclusters for the myeloid and B-lineage L2 objects were manually annotated for both CSF cells and PBMC, while the lymphoid (T- and NK-related) L2 objects were annotated using Azimuth (Extended Figure 1A, B). FindAllMarkers() function was used as stated above to find the canonical markers for the subclusters with each of the L2 objects, and dot plots were used to describe them (Figure S4-7).

### Erythrocyte and platelet correction

To account for the contribution of erythrocytes in CSF samples, we performed a correction despite the very low numbers of erythrocytes and platelets relative to the total number of cells per sample (Supplementary File 1). We calculated the erythrocyte proportion per sample, postulating that this proportion represents the “peripheral contribution” in the CSF. For each cell type in each sample, we applied a corrective factor by multiplying the sample-specific erythrocyte proportion by the cell type proportion within the same sample. This process was repeated for all cell types across all samples. The corrective counts — attributed to peripheral contribution — were then determined by multiplying the corrective factor for each cell type by its original counts per sample. The corrected counts were obtained by subtracting the corrective counts from the original counts for each cell type per sample. We then plotted cell proportions per sample as stacked bar plots before and after the correction, both for the overall CSF object and for the pertinent comparisons (Figures S2C, E, G). For PBMC, we just removed the erythrocytes and platelets, demonstrating no change in cell type proportions per sample following the removal (Figures S2D, F, H). Since we did not find much difference in cell type proportions per sample following these corrections, we removed the erythrocytes and platelets from all the main Seurat objects.

### Single-cell proportion test

For comparing the cell proportions between different conditions, we used the single-cell proportion test^133^. Specifically, a permutation test was used to calculate a p-value for each cluster, and a confidence interval for the magnitude difference was returned through bootstrapping. Line plots were then generated to visualize the difference of proportion of cell populations across different conditions with abs(log_2_FD) >0.41 and FDR <0.05.

### Differentially expressed genes (DEG), gene enrichment, and pathway analyses

DEG for a cluster or subcluster across the various conditions stated above were found using FindMarkers() function with the significant gene(s) identified as having abs(log_2_FC) >0.58 and adjusted p-value of <0.05. AddModuleScore() function was used for the average expression of a gene program (IFN-γ-signature genes in our case) across various clusters. For gene ontology (GO) terms pertinent to cluster-defining genes, to slightly widen the bracket of genes captured, while the average >-0.7-fold difference (log-scale) of expression between the cluster of interest (ident.1) vs. the rest of the cells was kept the same, the average expression in the rest of the cells (ident.2) was instead defined as ≤40% (instead of ≤30% as above for the Dotplot). The top 30 genes from this list were then used for gene enrichment analyses using EnrichR v3.2^134^. For differentially upregulated or downregulated genes across the various comparisons (abs(avg_log_2_FC) >0.58 and adjusted p-value <0.05), the lists were used as input for EnrichR for enrichment analyses. The databases used for human gene enrichment analyses included Gene Ontology (GO; 2023) – Biological Process (BP), Cellular Component (CC), and Molecular Function (MF) as well as pathway databases: KEGG (2021, human), Reactome (2022), BioCarta (2016), Panther (2016), and WikiPathways (2021, human).

Ingenuity was used for pathway analyses. For a list of DEG across comparisons for subclusters, the DEG were coded +100 or -100 when the avg_log_2_FC was >0.58 or <0.58, respectively. This was then input into Ingenuity/IPA^135^ for the classical pathway analyses. Comparison analyses were also done, and the Z-scored pathways and regulators were exported as a data-frame to be used for heatmap generation (as in Figures 4C, Extended Figure 17-18). Note that for heatmaps representing pathways across B-cell-depleted (all PRL-positive) vs. PRL-negative and PRL-positive vs. PRL-negative (or PRL-high versus PRL-low) comparisons, Z-score thresholds of abs(±2) were used (Figure 4C, Extended Figure 17).

### NicheNet ligand-receptor-target analysis

NicheNet was implemented to model intercellular interactions among immune cells in the CSF and blood^136,137^. The receivers (target genes) in various immune cells were set as the DEG across PRL-positive versus PRL-negative cases, while the senders were also set as the immune cells. Note, we employed crude cell annotations (CD4 T, CD8 T, B-cells, Mono (myeloid cells), NK, and cDC) for these analyses. We used a sender-agnostic and a sender-focused approach, using the previously subsetted Seurat object for the PRL-positive versus PRL-negative comparison according to detailed instructions (https://github.com/saeyslab/nichenetr?tab=readme-ov-file). Prioritized ligandreceptor (LR) and ligand-target gene interactions were plotted as Circos plots (Figure S9-10). The list of DEG across PRL-positive versus PRL-negative cases (expressed in ≥5% of cells in one cluster, adjusted p-value <0.05, and abs(log_2_FC ≥ 0.25)) was used as the input gene set.

### Comparison with immune-cell subset and microglial dataset from previous single-nucleus RNA-seq datasets

We downloaded the processed microglial Seurat object associated with the reference^138^ from Synapse. In this paper, microglial nuclei were enriched from 12 AD and 10 control human dorsolateral prefrontal cortices before being subjected to snRNA-seq. We then compared the transcriptome similarity on a cluster level between the processed Seurat object of the microglia^138^ against the CSF myeloid L2 object (query dataset). Briefly, we calculated the average transcriptome profiles of every annotated cell type in both the objects. Following that, we obtained the genes to represent the transcriptome and calculated pairwise Spearman and Pearson correlation across the genes’ expression between reference cell types and query clusters. Subsequently, we used a heatmap to represent the correlation matrices, showing that the two correlation approaches did not differ significantly for our cases (Extended Figure 3D). We performed a similar correlation analysis — employing Spearman correlation since it is less sensitive to outliers — to measure cluster level similarity of the transcriptome between the CSF L2 myeloid clusters and immune cell subset from Absinta et al., 2021^7^ (Figure 2C).

### Pseudo-bulking and DESeq2

We performed pseudo-bulking and DESeq2 analyses per the instructions provided on their GitHub webpage (https://github.com/hbctraining/scRNA-seq_online). Note that we used the crudest level annotations for performing pseudo-bulking with clusters defined as myeloid, T, B, NK, and other-T cells (Extended Figure 5C and 10D). The p-value cutoff was set at 0.05.

### T-cell receptor analysis and visualization

Following sequencing of the TCR libraries on Illumina’s NovaSeq platform, CellRanger v3.1.0 was used for demultiplexing, barcode processing, FASTQ file generation and alignment. Human VDJ reference (GRCh38/Ensembl/10x) was used for alignment. The filtered_contig_annotations.csv files for each of the CSF cell and PBMC samples were used to load the contigs into scRepertoire^139,140^ for further downstream analyses including basic clonal analyses, visualizing clonal dynamics and summarizing repertoires. The function combineExpression() was used for combining clones and single-cell objects and to visualize clone size on a gene expression UMAP for both the CSF cell and PBMC Seurat objects. Visualization for single cell objects was performed using scRepertoire package as detailed on the webpage (https://www.borch.dev/uploads/screpertoire/).

### Trex: combining deep learning and TCRs

For the purposes of combining gene expression and TCR profiling at the single cell level, we employed Trex^140^ according to detailed instructions (https://www.borch.dev/uploads/screpertoire/articles/trex). Briefly, TCRbased metrics (TRB and TRA chains) were used in the algorithm to return autoencoded values to be used for dimensional reduction. Firstly, maTrex() function was employed on the list output of combineTCR() function of the scRepertoire: TRA chains were used initially with variational autoencoder (VAE) as the model and Atchley factors used as the encoder.input. Note, maTrex() does not filter the input for only T-cells with attached TCR data. This was followed by running runTrex() using TRB chains, VAE model, and Kidera factors as the encoder.input; runTrex() does filter for only T-cells with attached TCR data. The resulting Trex reduction stored in the Seurat object was then subsequently projected onto the UMAP. We implemented weighted nearest neighbors (WNN) to use the Trex embedding information and gene expression PCA for integration of gene expression and TCR modalities^15^. The function FindMultiModalNeighbors() was used on Trex embedding and gene expression PCA with 30 dimensions for each. Subsequently, a wnnUMAP was generated and clusters determined at a resolution of 0.6. For finding possible relationships epitope relationships with the CDR3-sequences, annotateDB() was used with defined edit distances from the reference set as 2.

### Protein measurement by Olink proximity extension assay and analyses

Proteins were quantified by Olink proximity extension assay as previously described^141^. CSF and serum aliquots were analyzed using the Cardiometabolic, Inflammation, Neurology, and Oncology Olink Explore 384 panels (∼750 and 1463 analytes in CSF and serum, respectively). All samples passed quality control measures. Results were reported as Normalized Protein eXpression (NPX) values in log_2_ scale for quantification of protein abundances.

#### Differential expression analyses and ANCOVA

For all the differential expression analyses of the three primary comparisons in CSF and serum, relevant samples for each of the three comparisons mentioned above were selected. Cross-sample normalization was performed by implementing cyclic LOESS normalization using the normalizeBetweenArrays() function. This was followed by replacing the negative values in the expression matrix by zero. We then plotted pre- and post-normalization boxplots. We computed the mean and coefficient of variation for each assay, applying lowess() smoothing for visualization. Subsequently, we removed proteins below the noise cutoff, defined to be the lowest mean value at which the linear relationship with coefficient of variation is grossly lost. Thus, we filtered out noisy or low-signal proteins, and the filtered data was then used for downstream analyses.

ANCOVA was performed to model for the group-pertinent comparisons, adjusting for age and sex as follows:

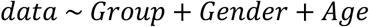

Adjusted data matrices were then used for downstream analyses including differential expression of significant proteins (defined as log_2_(fold difference) >0.58 and p-value/post-hoc p-value <0.05), and generation of heatmaps, volcano plots, and boxplots.

#### Protein Co-expression Network Analyses

We used WGCNA (Weighted Gene Correlation Network Analysis; version 1.72-5) for making a CSF and serum proteome network, using 22 and 21 serum samples respectively^142^. For the input expression matrix, we used ANCOVA adjusted for covariates, including age and sex, not modeling for any specific comparison. This was followed by covariate inspection and plotting the variance explained by the covariates as a heatmap to confirm if the adjustment minimized the contributions of the covariates to the expression matrix. WGCNA was then implemented on the CSF and serum protein abundances separately with the following parameters: power = 14, deepSplit = 4, minModuleSize = 10, mergeCutHeight = 0.07, TOMdenom = “mean”, bicor correlation, signed network type, PAM staging and PAM respects dendro as TRUE, and a maxBlockSize larger than the total number of protein assays. Module members were then iteratively reassigned to enforce kME table consistency, as previously described^143,144^. The modules were then visualized using iGraph (v1.4.1) package using a custom implementation available as the buildIgraphs() function available at https://github.com/edammer/netOps. Module eigenprotein correlations and significance were visualized in circular heatmap using circlize, dendextend (v1.17.1), and dendsort (0.3.4) R packages.

#### Interpreting iGraphs made for the protein modules

Protein modules and their networks shown as iGraphs. Each graph represents approximately top 100 proteins (or isoforms). They are ranked started in the inner circle at 3 o’clock, going counterclockwise, then continuing at 3 o’clock in the next outer concentric circle of nodes, and so forth. Ranking is by kME, bicor rho-ranked correlation to the module eigenprotein, which is the 1^st^ principal component (PC) of the module’s abundance variance. The node size is proportional to intramodular kME values (bicor calculation). Detailed networks are present as a Supplementary File 5 and 6 for both the serum and the CSF protein networks respectively. To the right of every iGraph (network) for each module, there is a decomposition of the iGraph into nodes connected by the strongest edges which are ranked by pairwise variance similarity (topology overlap matrix, TOM calculation). The degree (number of strongest connections) to each node kept in the graph is a superscript after the gene symbol.

### *In silico* knock out (KO) of selected genes (‘regulators’) and gene-positive cell population depletion from PRL-positive objects in CSF and PBMC

The gene regulatory network (GRN) was generated starting from PRL-positive immune cells only from CSF or blood. The simulated effect of gene knock-out on the original GRN was measured using scTenifoldKnk^115^, while the simulated effect of gene-positive cell population depletion was gauged using scTenifoldNet^116^. These are machine-learning-based frameworks that take two single-cell matrices as inputs, with the analysis identifying genes whose transcriptional regulation is shifted between the two conditions (GRN from PRL-positive cells as the control condition, with the other being the altered GRN computed from dataset with gene knock-out or cell population depletion). In the case of control, both the input matrices were PRL-positive matrices. The workflow, as presented in the original papers, is summarized as follows: (1) cell subsampling; (2) network construction; (3) network denoising; (4) manifold alignment; and (5) module detection. The final output is a table comprising of genes and their associated Euclidean distances computed as the distance between the coordinates of the same gene in the two conditions.

We also performed PCA (principal component analysis) of the simulated distances from comparing the original GRN and the same GRN following the *in silico* perturbation. The Euclidean distances between the different comparisons were plotted as a color-coded heatmap, although only the first 2-dimensional space was used. Significantly altered genes (*p < 0.05*) were used for functional enrichment analyses using GSEA (gene set enrichment analysis). The DEG in the B-cell-depleted versus PRL-positive comparison were used as the *in vivo* B-cell-depletion gene lists, to compare against other gene lists (Figure 4E).

### Flow Cytometry

Cryopreserved PBMC were used for immunophenotyping by flow cytometric analysis. PBMC were stained with specific antibodies specific for cell-surface antigens (Supplementary File 8). For intracellular or intranuclear staining, PBMC were fixed and permeabilized with Fixation/Permeabilization solution (BD Biosciences) or Foxp3/Transcription factor staining buffer (eBioscience), respectively, after surface staining. The fixed/permeabilized cells were intracellularly or intranuclearly stained with specific antibodies (Supplementary File 8). All the stained cells were analyzed using spectral flow cytometry Cytek Aurora (Cytek Biosciences). The data were analyzed using FlowJo 10.10 software (FlowJo LLC).

## Data availability

Raw and processed datasets generated in this study will be deposited to Gene Expression Omnibus (GEO) upon acceptance. Source data are provided with this paper.

## Acknowledgments

The Intramural Research Program of NINDS (ZIA NS003119, DSR), Abata Therapeutics, Sanofi, and the Dr. Miriam and Sheldon G. Adelson Medical Research Foundation provided funding toward this work. We acknowledge contributions of the NHLBI DNA sequencing and genomics core, the NIH High-Performance Computing Biowulf cluster, and the NIMH Functional Magnetic Resonance Imaging Facility. We thank all the members of the NINDS Neuroimmunology Clinic (NIC) and the outpatient (OP5) clinic for expert care and assessment of the patient participants who contributed to this study, and we are especially grateful to those participants. We also acknowledge and thank Prater et al^138^ for allowing us to download and use their transcriptomic dataset from Synapse. In addition, we thank Edoardo Pedrini for introducing scTenifoldKnk and scTenifoldNet to us, and Jing-Ping Lin for helping troubleshoot bioinformatic analyses and for discussions, and the NINDS Advanced MRI Section (Jeff Duyn, Peter van Gelderen, Jacco De Zwart, and Jiaen Liu) for expertise involving advanced MRI.

## Author contributions

SAR and DSR designed the study, interpreted the results, and prepared the manuscript. SAR acquired and processed libraries and analyzed the scRNAseq and scTCR-seq data. YEA and ESB acquired and processed some libraries for scRNAseq and scTCR-seq. SAR, YEA, and NN obtained samples for proteomics. SAR and YEA performed and analyzed the flow cytometry data. SAR, AB, and LMG analyzed the proteomic data. KRJ and EBD helped with bioinformatic analyses. SAR and MIG acquired MRI data. SAR cleaned and processed published datasets. SAR, MIG, TJT, DO, RMR, SJ, ECM, and DSR developed protocols for the study. SAR, YEA, AB, ESB, MIG, JMD, DM, EBD, KRJ, MA, TJT, DO, RMR, SJ, ECM, and DSR provided critical feedback on the manuscript, helping with the review and edits. DSR supervised the study.

## Corresponding author

Correspondence to DSR.

## Declaration of Interests

DSR has received research funding from Abata and Sanofi, related to the current study.

JMD, DM, ECF were employees of Abata at the time this study was performed and may hold shares and/or stocks in the company.

RMR is a venture partner at Third Rock Ventures, is a cofounder of Abata, and may hold shares and/or stocks in the company.

ASB, TJT, and DO are employees of Sanofi and may hold shares and/or stocks in the company.

---

**Extended Figure 1:**
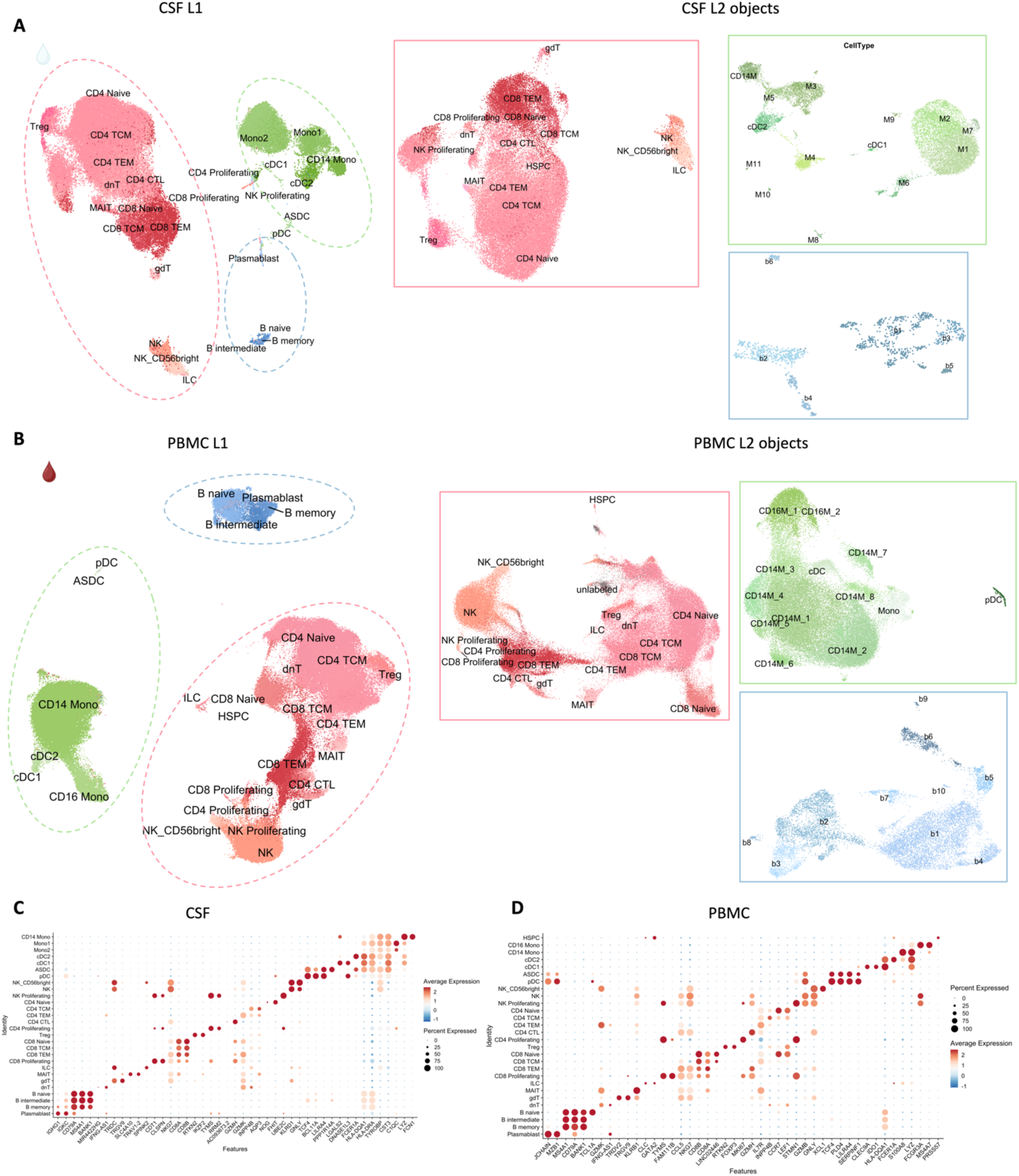
Single-cell transcriptomics reveal compartment-specific cell populations in the CSF and blood. a) Level 2 (L2) analyses of the myeloid, B-lymphoid, and T-lymphoid and NK cell populations of the CSF (enclosed by green, blue, and pink dashed lines, respectively). The L2 subclusters were derived from L1 clusters enclosed within similarly colored dashed lines (right panel). Myeloid CSF cells subclustered into 14 populations with cell markers of the individual cell clusters depicted in Figure S4. L2 analysis of the Blymphoid cells of the CSF clustered into 6 populations (Figure S5). T-lymphoid and NK cells at L2 are also shown (box enveloped by the pink line). b) L2 analyses of the myeloid, B-lymphoid, and T-lymphoid and NK cell populations of PBMC (enclosed by green, blue, and pink dashed lines, respectively). Peripheral myeloid cells subclustered into 13 populations with the subcluster-defining genes represented in Figure S6. L2 analyses of the peripheral B-lymphoid lineage cells showed 10 populations (markers shown in Figure S7). Note, a combined UMAP of both CSF and PBMC samples is shown in Figure S8. c) Dotplot representing the top 2 canonical cluster-defining genes of the cell populations in the CSF. d) Dotplot representing the canonical marker genes of the cell populations in the PBMCs.

**Extended Figure 2:**
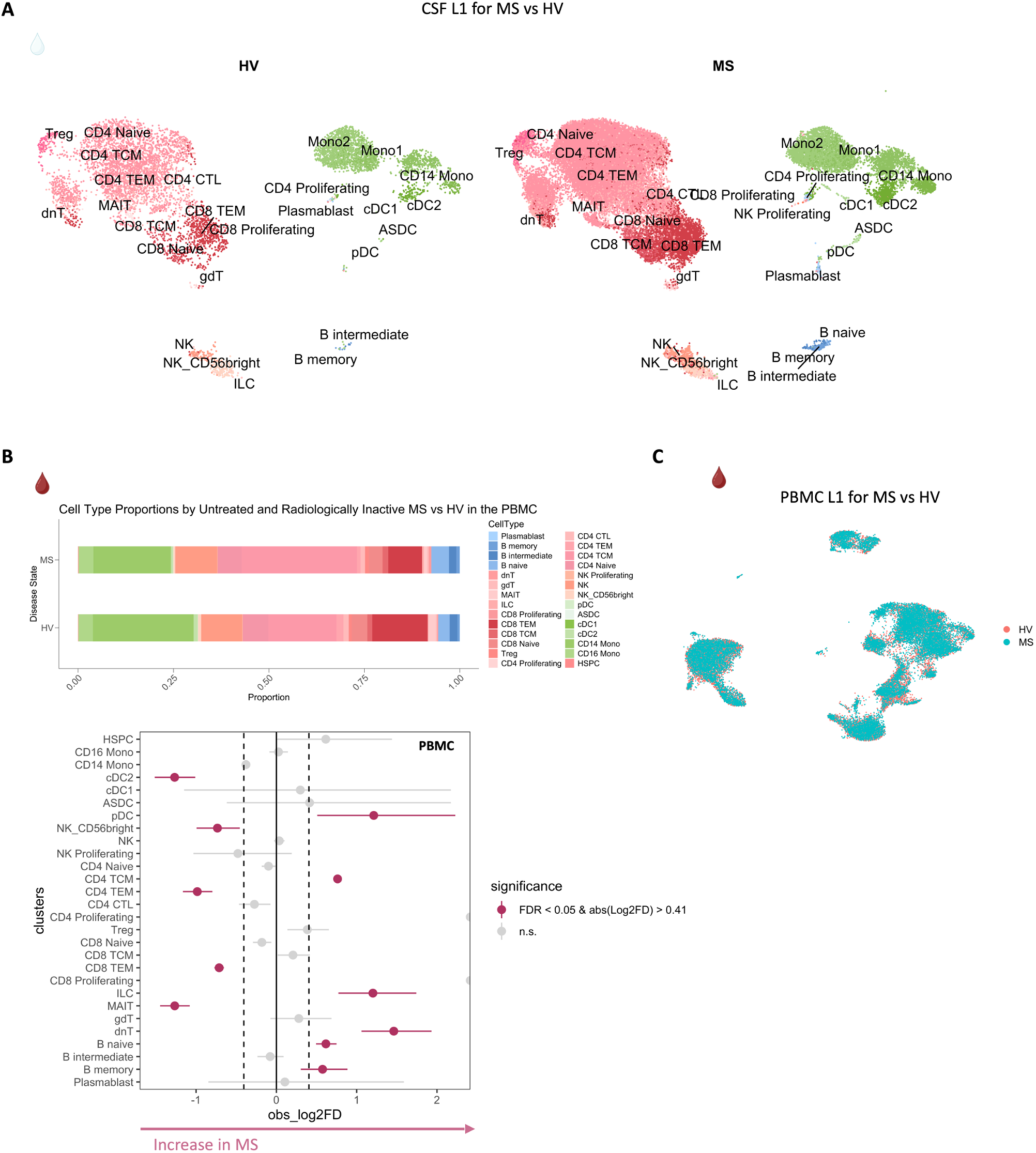
Altered CSF and PBMC cellular composition in untreated and inactive MS. a) CSF L1 UMAP for untreated and inactive MS vs. HV. Evident is the enrichment of B-lineage cells in the CSF of patients with MS (see also Fig. 1). b) Stacked column graph summarizing proportions of L1 PBMC annotations in MS and HV (upper panel). Bottom panel shows a lineplot with significant differences in the PBMC cellular composition between untreated and inactive MS cases and HV. There are increased proportions of B-memory, B-naïve, dnT, and CD4 TCM cells in MS cases. CD4 TEM and CD8 TEM cell proportions were decreased in the PBMC of MS cases. c) PBMC L1 UMAP comparing MS and HV with random sampling of 15,000 cells per condition.

**Extended Figure 3:**
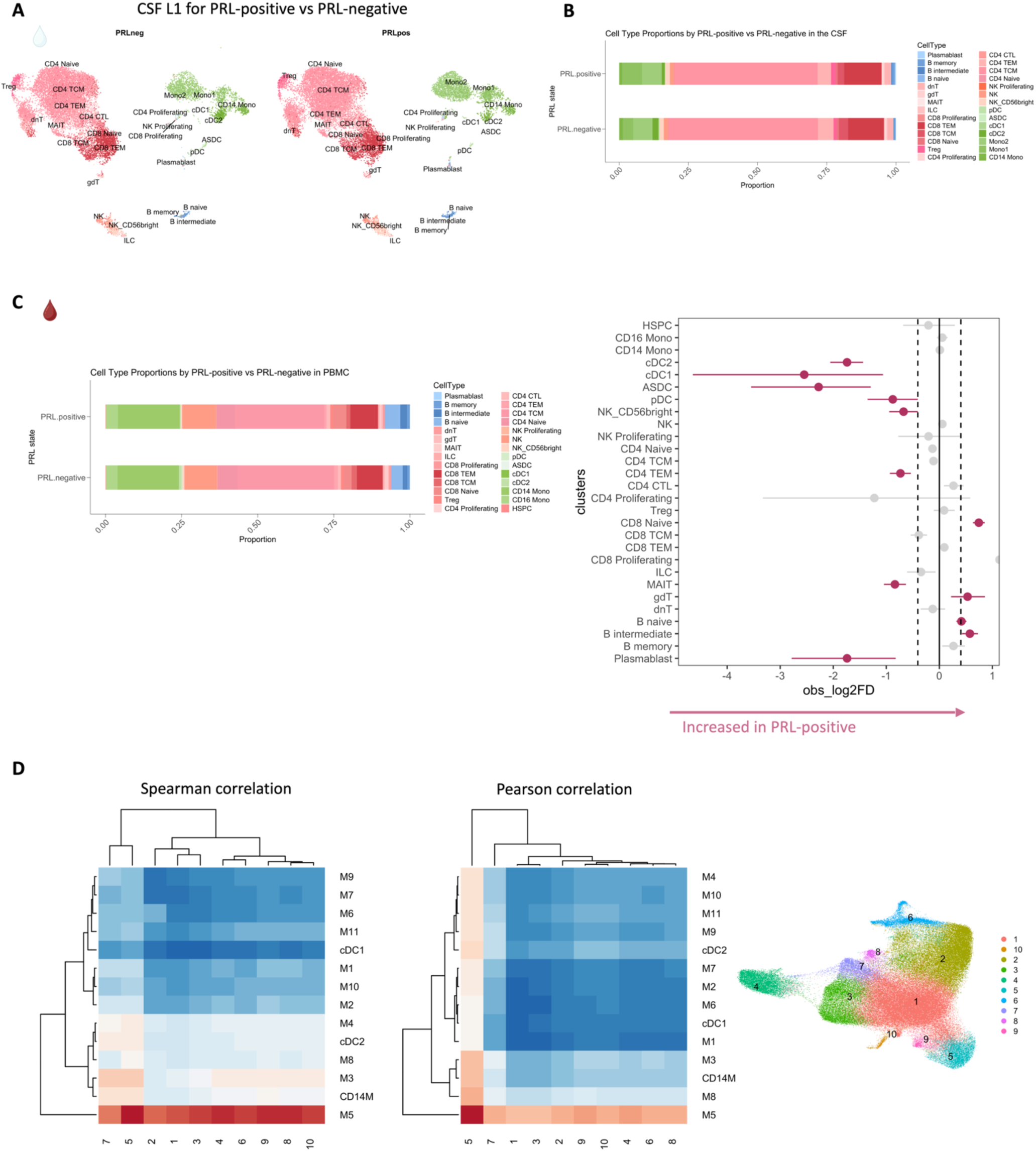
B-lineage cells are increased in proportion in PBMC of PRL-positive relative to PRL-negative cases. a) CSF UMAP for the PRL-positive versus PRL-negative samples (subset from the main object). b) Stacked bar graph illustrating the L1 CSF cell proportions contrasting PRL-positive versus PRL-negative cases. c) Comparison of the PBMC L1 cell proportions for PRL-positive versus PRL-negative cases shown in stacked bar plots. The panel on the right shows a lineplot contrasting peripheral immune cell compartment of PRL-positive versus PRL-negative cases, showing slightly increased proportion of CD8-naïve and B-intermediate cells in PRL-positive cases. d) Correlation heatmaps (Spearman and Pearson correlations) comparing average transcriptome similarity of L2 annotations of the CSF myeloid cells versus microglia from the dorsolateral prefrontal cortices of an Alzheimer’s disease cohort^138^, showing that M5 myeloid subcluster in the MS CSF most closely resemble microglia in the brain parenchyma.

**Extended Figure 4:**
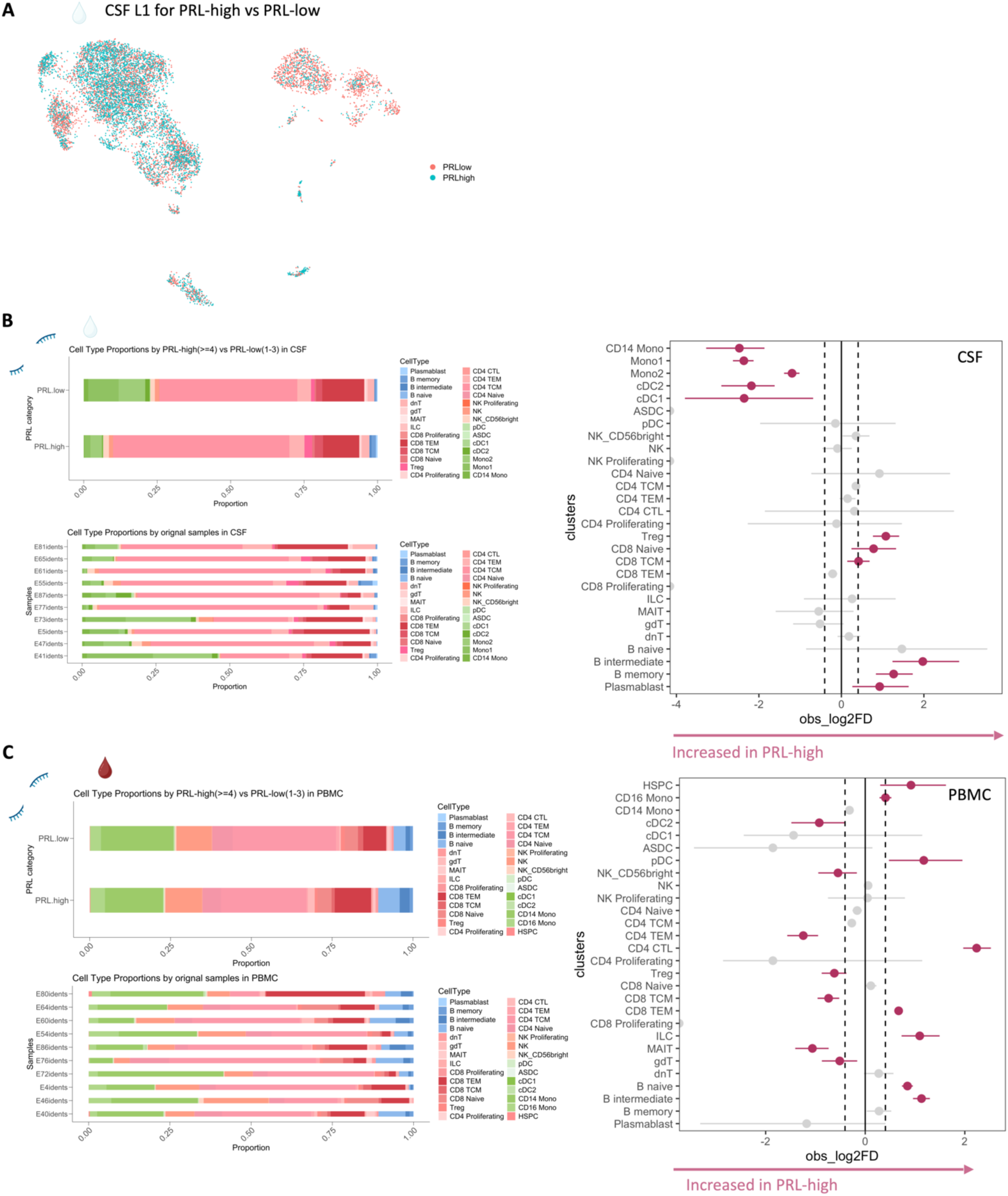
Relative to low PRL burden (PRL 1-3), high PRL burden (PRL ≥ 4) patients demonstrate increased proportions of B-lineage cells and Tregs in the CSF, and increased proportion of CD4 CTL cells in the blood, of PRL-high cases. a) CSF L1 UMAP for PRL-high (PRL ≥4) versus PRL-low (PRL 1-3) comparison with the total cell population down-sampled equally per condition to 5000 cells. b) Comparison of CSF L1 cell proportions in PRL-high vs. PRL-low cases shown in a stacked bar chart. Bottom panel shows the L1 cell proportions in a stacked bar chart per sample. The line plot on the right contrasts the difference in cell proportions in the sample space for the PRL-high versus PRL-low cases and shows that the CSF of PRL-high cases is enriched with B-intermediate, B-memory, and Treg cells and possibly CD8-naïve, CD8-TCM, and plasmablast cell populations. c) Stacked bar plots summarizing the PBMC L1 cell proportions across the PRL-high vs. PRL-low comparison, and the L1 cell proportions per sample for the same comparison (bottom panel). The line plot illustrates increased proportion of CD4-CTL, CD8-TEM, ILC, B-intermediate, and possibly pDCs in the PBMC of PRL-high cases relative to PRL-low.

**Extended Figure. 5:**
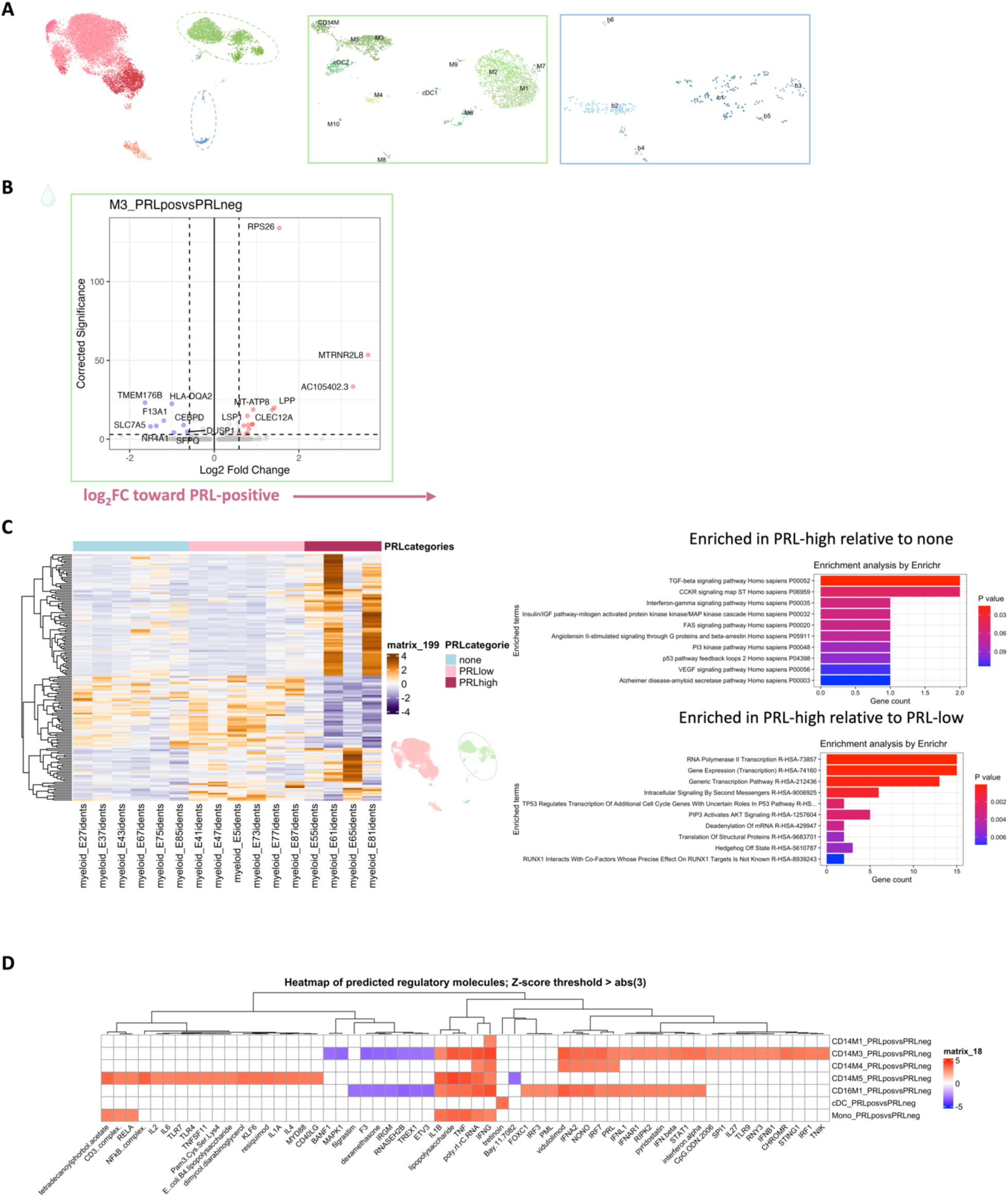
IFN-mediated regulation of myeloid cells in both CSF and blood in chronic active MS. a) CSF L1 UMAP and the myeloid and B-lineage cell subsets shown on the right of the figure. b) Volcano plot of DEG in CSF myeloid L2 M3 subcluster comparing cells from PRL-positive versus PRL-negative cases. c) Heatmap showing the DEG in CSF myeloid cells across PRL categories. Pseudo-bulk data is shown, using DESeq2 to model for PRL categories. The panels on the right represent GO terms for: (1) genes up-regulated in cases with PRL-high burden vs. none, including terms such as IFN-γ, MAPK cascade; and (2) terms enriched for PRL-high vs. PRL-low cases, including transcription, intracellular signaling by second messengers, or PIP3/Akt activation. The fold-change for this analysis was set at 1.25 and p-value < 0.05. d) Heatmap showing Z-score enrichment of predicted regulatory molecules with a Z-score threshold ≥ abs(±3). Among the enriched predicted regulatory molecules across PRL-positive vs. PRL-negative comparison in the myeloid subsets of blood are IFNG, IFNA2, TNF, and IL1B.

**Extended Figure 6:**
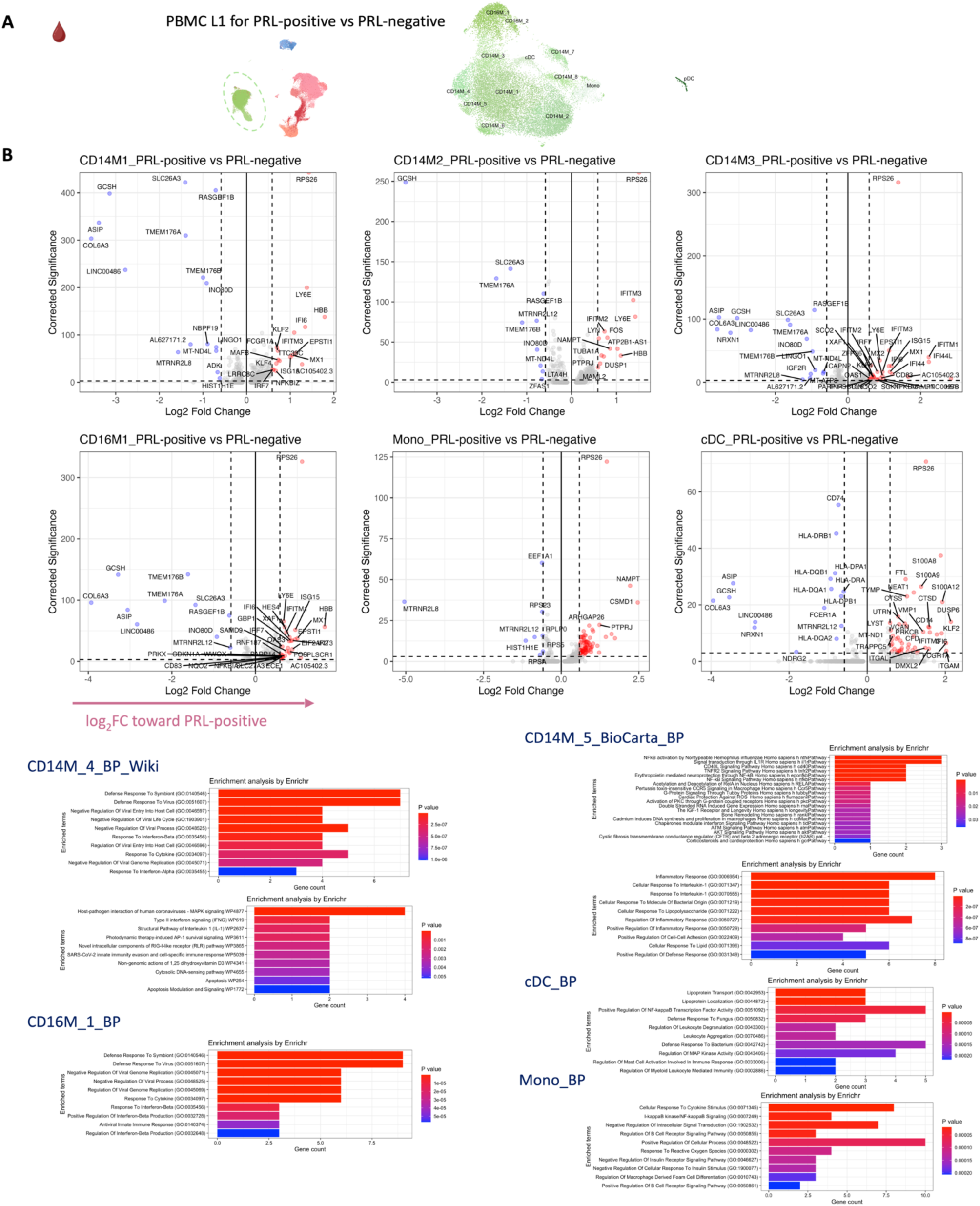
Blood CD14+ and CD16+ monocytes demonstrate an inflammatory phenotype in PRL-positive cases. a) PBMC L1 UMAP (left panel) with the peripheral myeloid cells encircled in green and the L2 myeloid subclusters shown in the right panel. b) Volcano plots comparing PRL-positive versus PRL-negative in the peripheral myeloid subsets. c) GO terms for the upregulated genes in PRL-positive for the corresponding myeloid subclusters reveals enrichment of IFN-mediated responses, TNFR2-signaling pathways, response to IL-1, and involvement of canonical NF-κB inflammatory signaling.

**Extended Figure 7:**
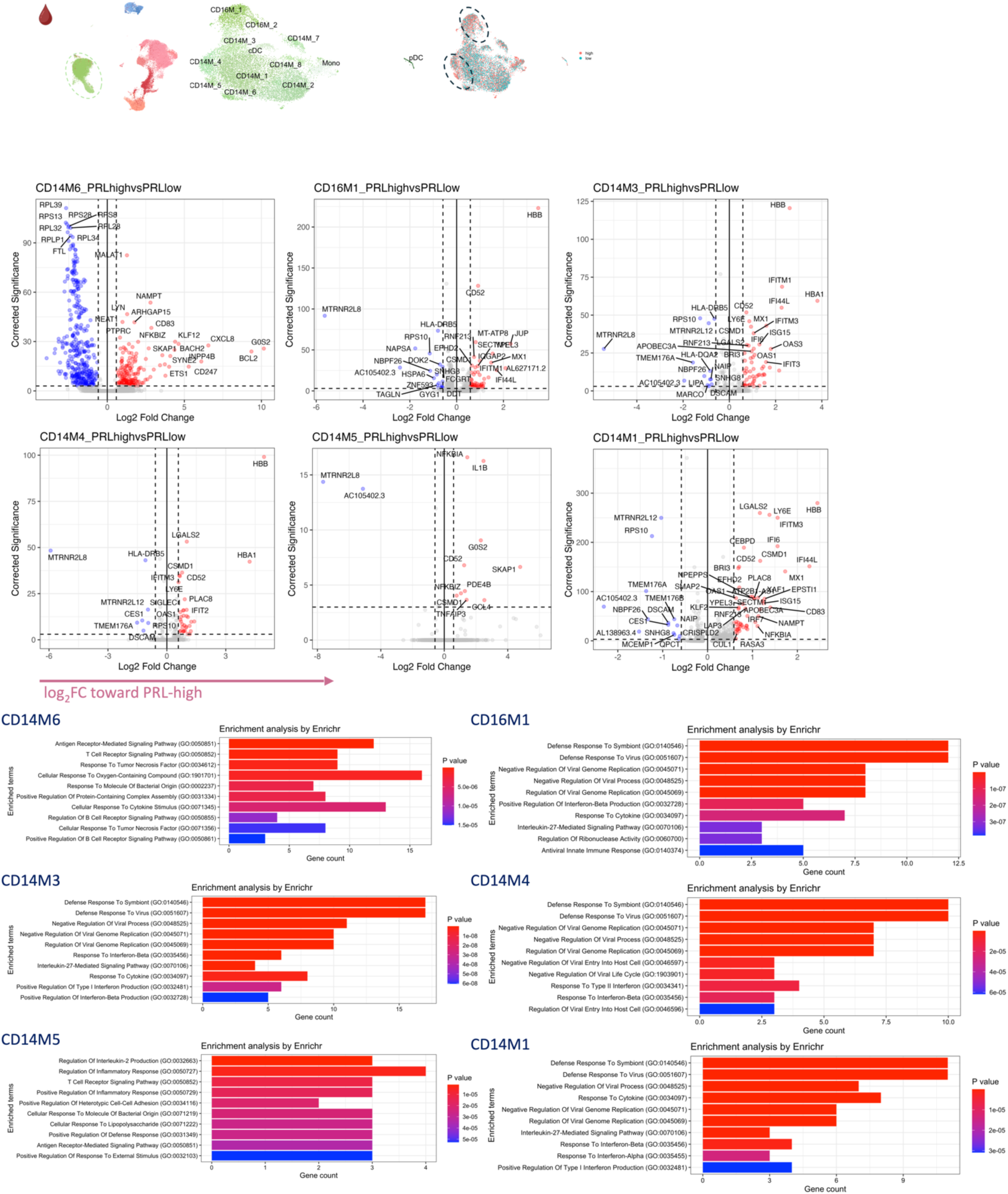
High PRL burden state correlates with increased myeloid activation involving IFN and NF-κB signaling. PBMC L2 myeloid subclusters and the volcano plots for the DEG contrasting PRL-high to PRL-low cases are shown with the corresponding GO terms. For the upregulated genes, there is enrichment of terms positive regulation of type-I IFN production, response to type-II IFN, response to LPS, regulation of IL-2 production, and IL27-mediated signaling. For the CD14M_6 subcluster, there was also significant downregulation of genes involved in translation and oxidative phosphorylation across cells from PRL-high vs. PRL-low cases.

**Extended Figure 8:**
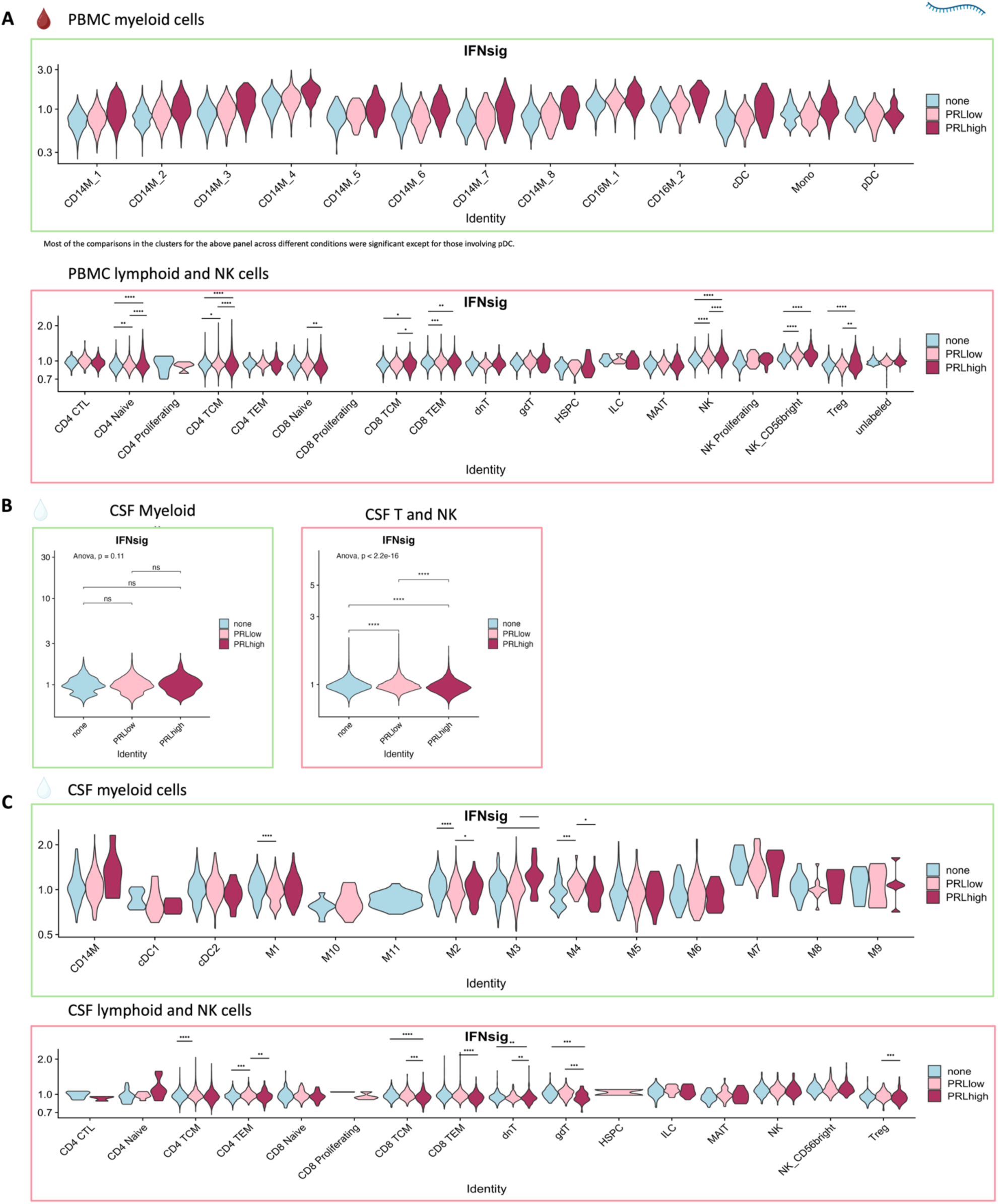
Transcriptional abundances of the IFN pathway genes in the immune cells of blood and CSF. a) Violin plots comparing the expression of IFN-signature score (average module score, defined in the main text) across PRL categories within the peripheral myeloid subclusters (top panel). Violin plots comparing the expression of IFN-signature in the T-lymphoid and NK cells of the blood across PRL categories (none=0, low=1-3, high=≥4) (bottom panel). b) Average expression of the IFN-signature in CSF myeloid (left panel) and T/NK cells (right panel) across PRL categories. c) Average expression of the IFN-signature in various CSF myeloid subclusters (top panel) and T/NK cell subclusters (bottom panel) across PRL categories.

**Extended Figure 9:**
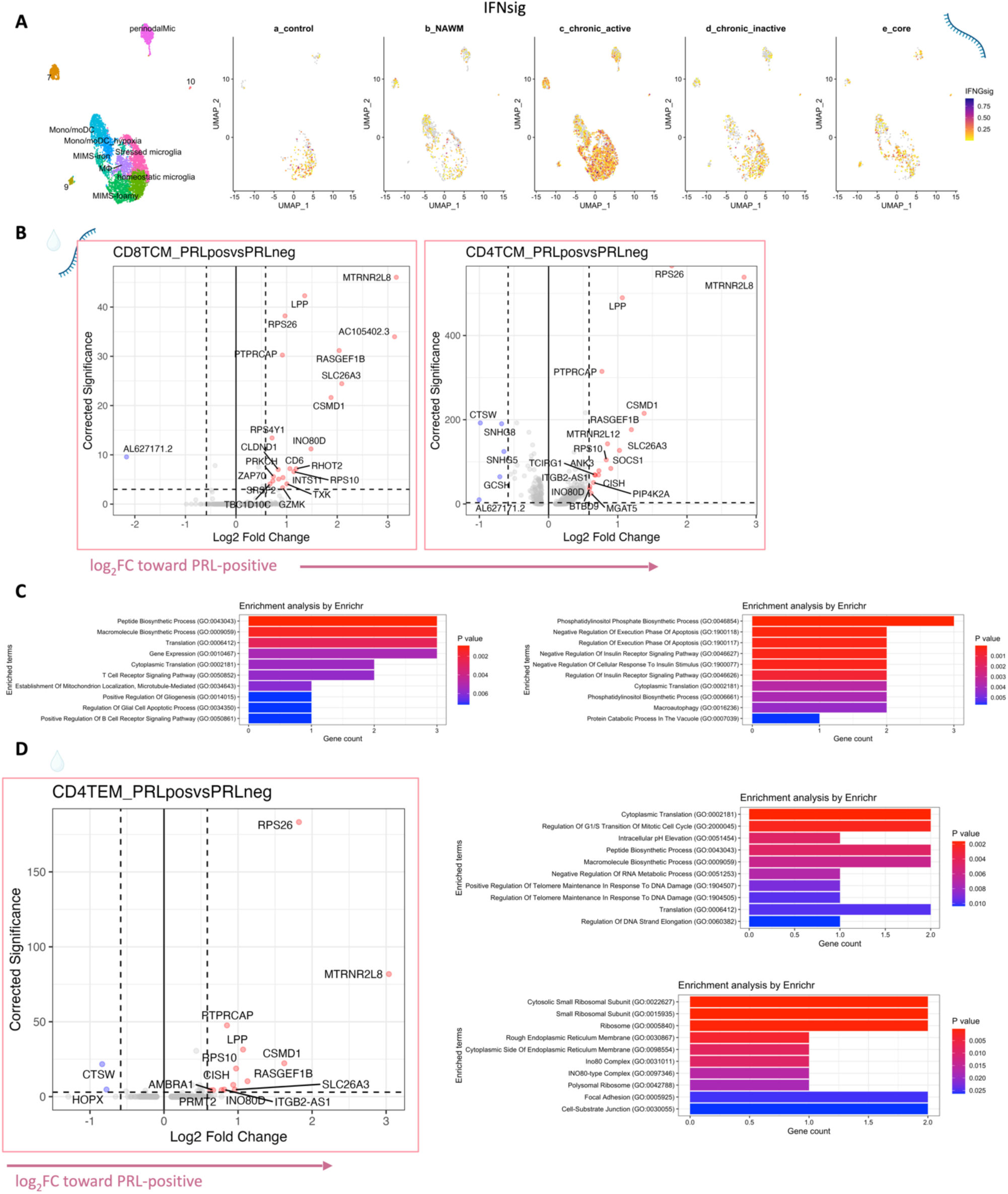
Chronic active lesions are associated with peptide biosynthesis, translation, anti-apoptotic mechanisms, and TCR-signaling in the T-cell compartment of the CSF. a) Enrichment of IFN-signature at chronic active lesion edge versus other locations using a previously published dataset^7^. b) Volcano plots for CD8-TCM and CD4-TCM cells in the CSF across PRL-positive vs. PRL-negative showing upregulation of genes like *MTRNR2L8, MTRNR2L12, SLC26A3*, and *PTPRCAP*. c) The corresponding GO terms for the upregulated genes in CD8- and CD4-TCM clusters of CSF in PRL-positive relative to PRL-negative (Extended Figure 9C) are shown. Note terms such as peptide biosynthetic process, translation, and negative regulation of apoptotic process. d) Volcano plot for CD4-TEM cell population comparing PRL-positive vs. PRL-negative in the CSF and the corresponding GO terms on the right of the panel.

**Extended Figure 10:**
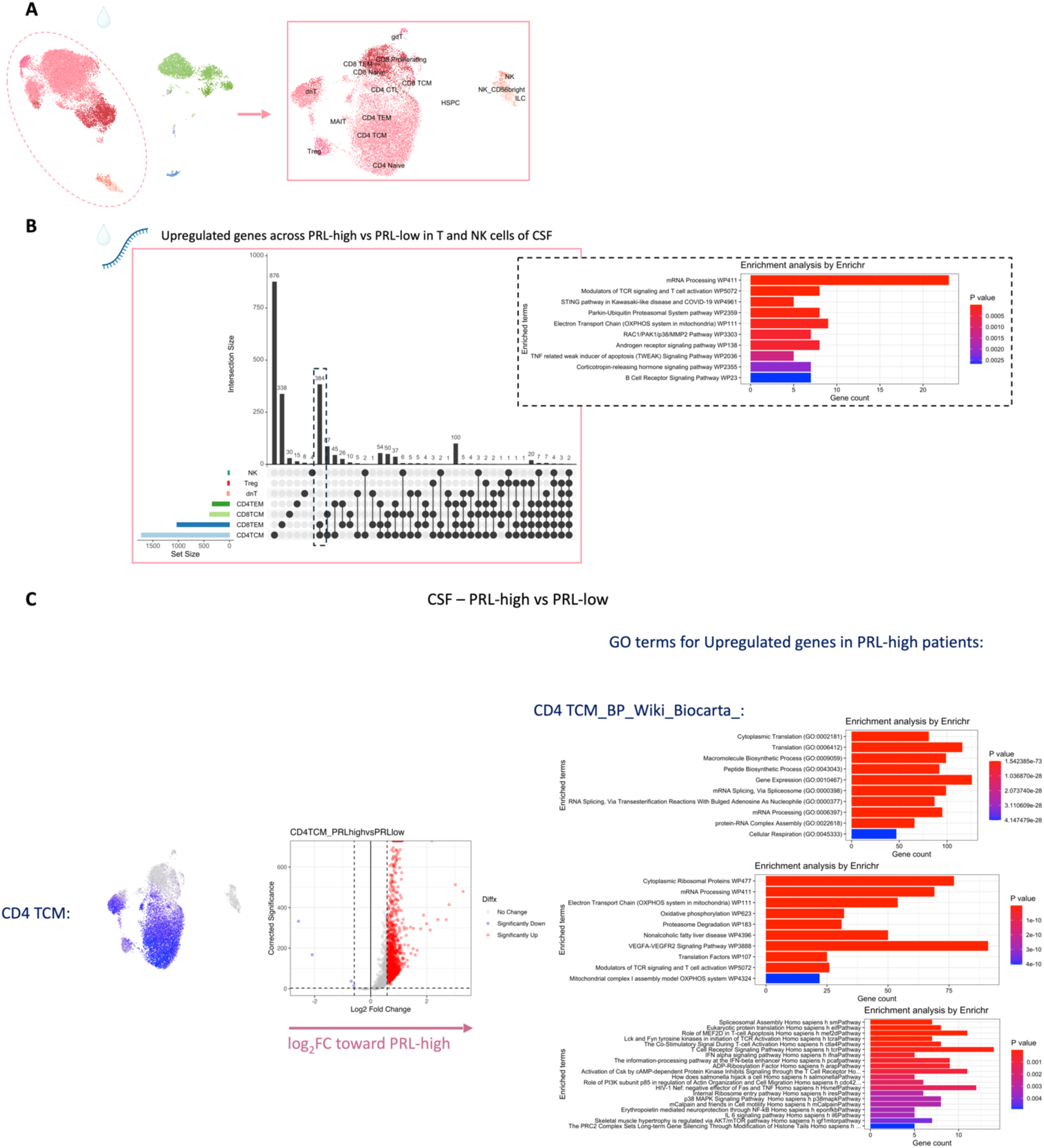
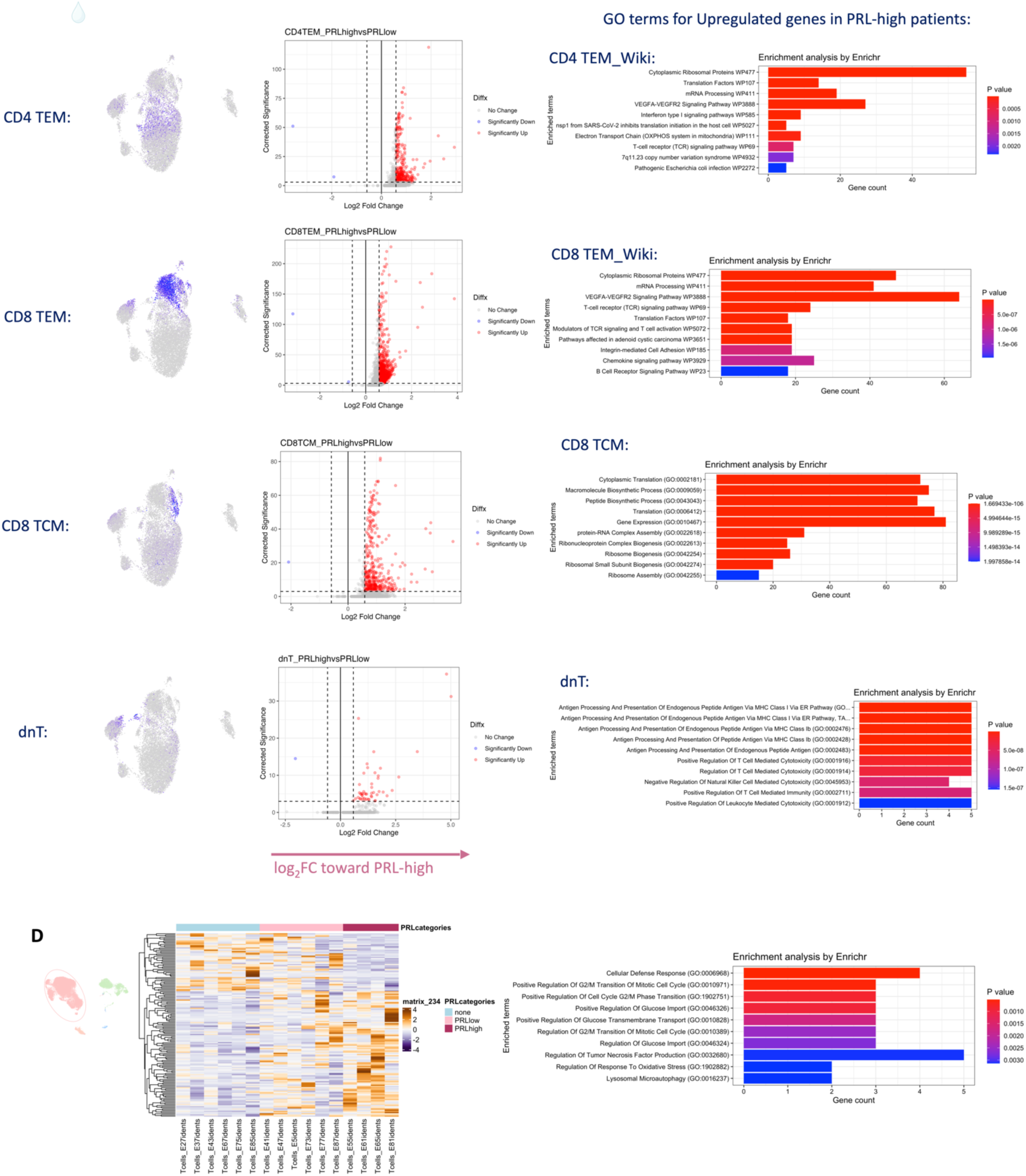
CSF T cells massively upregulate TCR-signaling, oxidative phosphorylation, electron transport chain (ETC) pathways in cases with high PRL burden (PRL ≥ 4) a) CSF L1 UMAP for the PRL-high vs. PRL-low comparison with L2 T-lymphoid and NK cells in the right panel. b) Upset plot showing the differentially upregulated genes in various T-cell subsets and NK-cells of the CSF in PRL-high vs. PRL-low states. The vertical dashed-line box represents the shared up-regulated genes amongst CD4-TCM and CD8-TEM clusters. Shown also are the GO terms of the shared upregulated genes in the PRL-high group. Note the enrichment of mRNA processing, TCR signaling and activation, OXPHOS/ETC pathway, and STING pathway. Similar transcriptional changes involve the blood adaptive immune cells as well (see Extended Figures 11-12). c) CSF CD4 and CD8 T-cell immune subsets are shown in the L2 UMAPs for lymphoid cell populations on the left. dnT cells are also shown, since they were found in high proportions in untreated and inactive MS cases relative to HV. The volcano plots comparing PRL-high (PRL ≥ 4) vs. PRL-low (PRL 1-3) show differential expression of genes in the various cell populations. Note the remarkable upregulation of genes in PRL-high relative to PRL-low, particularly in the CD4-TCM, CD8-TCM, CD4-TEM and CD8-TEM cells. The GO terms (shown on the right in bar plots) common for the upregulated genes in these cell populations included translation, peptide biosynthesis, mRNA splicing and processing, electron transport chain (OXPHOS in mitochondria), TCR signaling, costimulatory signals for T-cell activation, IFN type-I signaling, and VEGFAVEGFR2 signaling pathway. The dnT cells were significant for antigen processing and presentation via MHC-I pathway and positive regulation of T cell mediated cytotoxicity. d) Heatmap representation of significant genes in CSF T-cells across PRL categories per pseudo-bulking and DESeq2 (left panel) and the corresponding enrichment terms (right panel). There is similarity in GO terms for genes enriched across PRL categories and those across PRL-high (PRL ≥ 4) relative to PRL-low (PRL 1-3) cases.

**Extended Figure 11:**
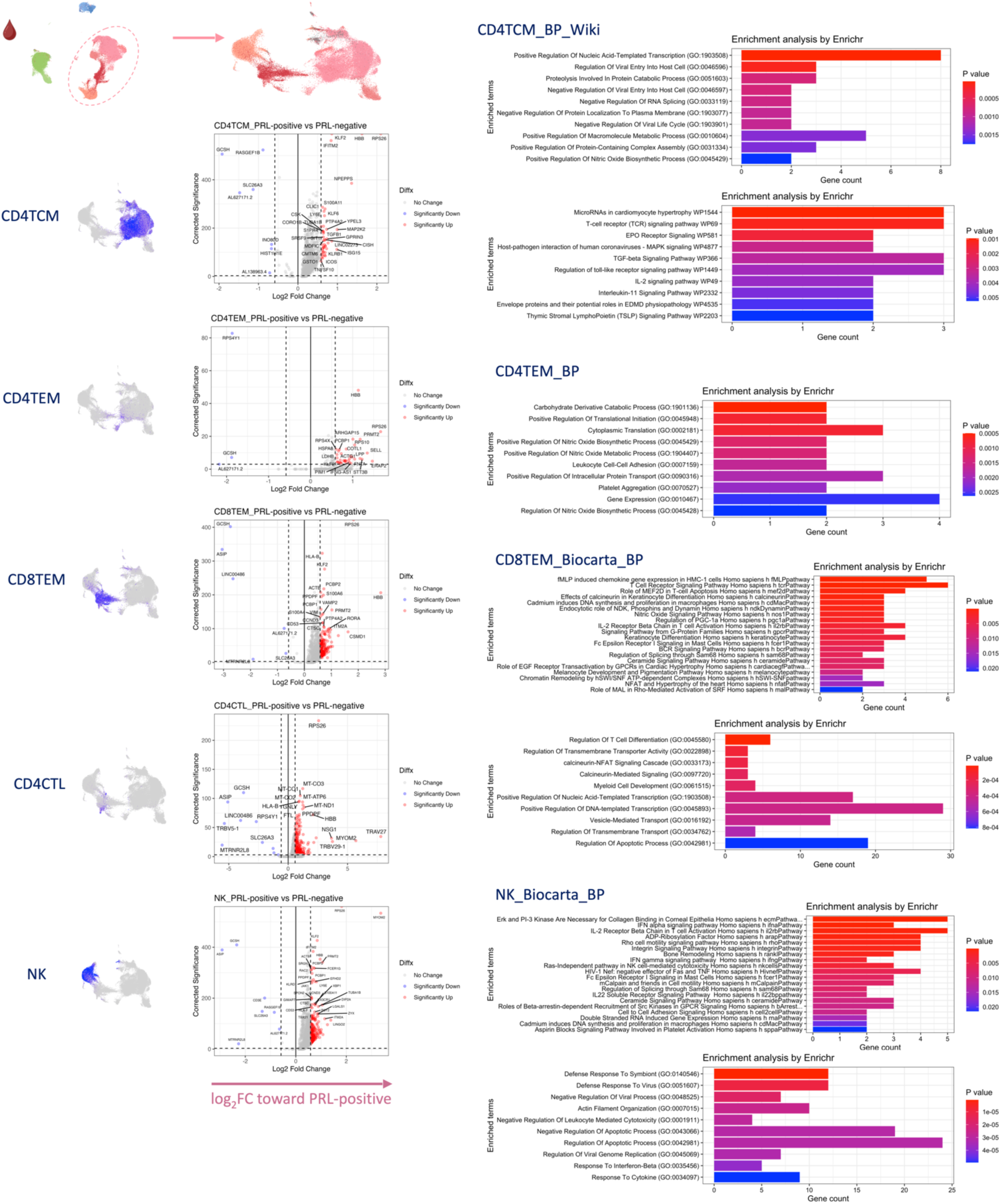
Peripheral CD4 and CD8 T cell subsets and NK cells upregulate TCR- and cytotoxicity-related responses in PRL-positive cases. PBMC L2 UMAP lymphoid cell subsets and NK cells are represented on the left. Volcano plots comparing PRL-positive versus PRL-negative in peripheral lymphoid and NK cells. Upregulated genes and their corresponding GO terms are shown on the right, with predominant terms including positive regulation of DNA-templated transcription, T-cell differentiation, TCR-signaling pathway, interferon-pertinent signaling, IL2-R β-chain in T-cell activation, regulation of apoptotic process, and negative regulation of viral processes.

**Extended Figure. 12:**
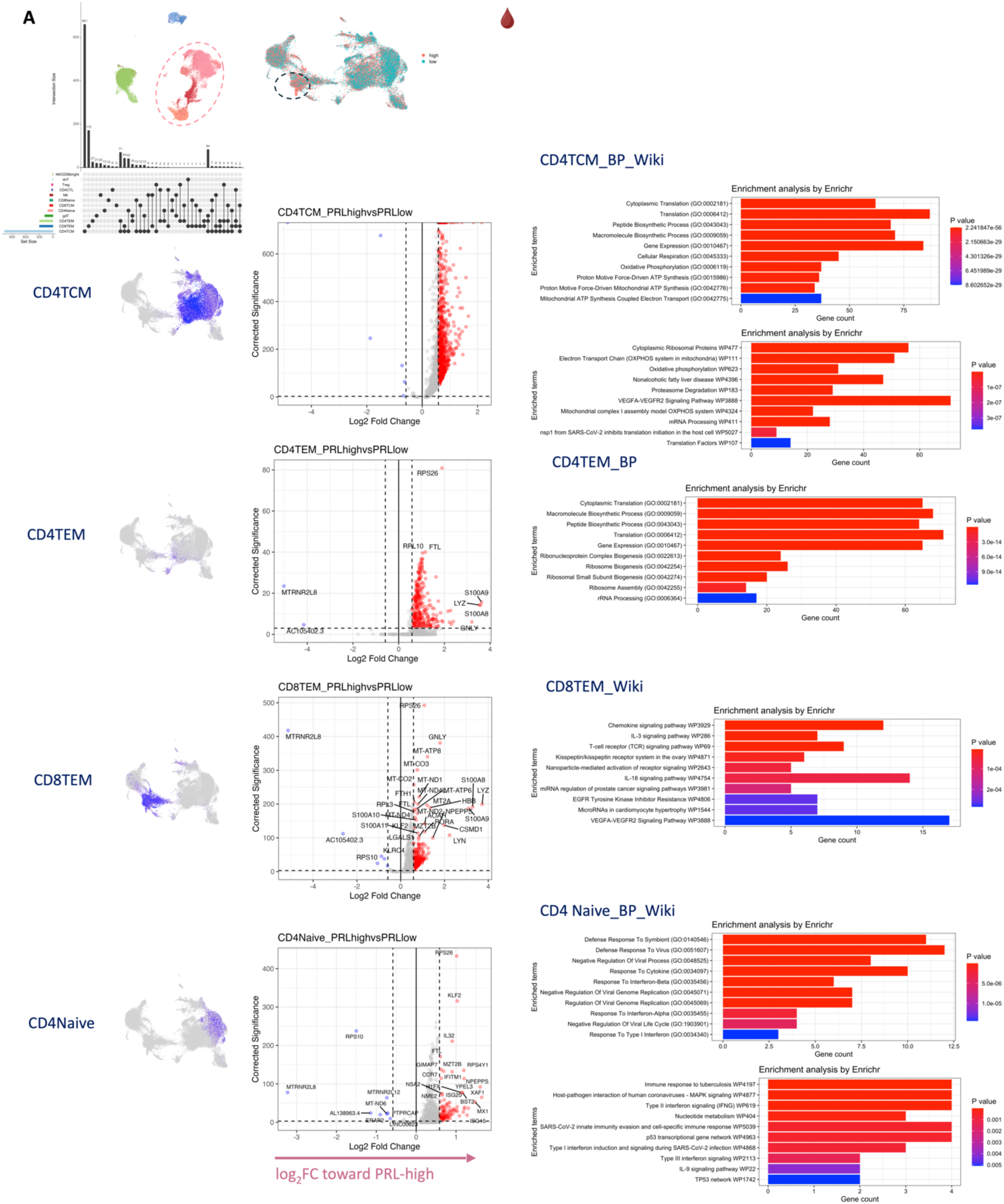
High PRL burden state correlates with increased peripheral TCR-signaling in the lymphoid compartment. PBMC lymphoid cells shown with the L2 UMAP and the Upset plot, showing the overlap between upregulated genes across the PRL-high versus PRL-low comparison. Volcano plots for the CD4-TCM, CD4-TEM, CD8-TEM, and CD4-naïve cell populations are shown with corresponding GO terms for the upregulated genes. The GO terms for the upregulated genes are essentially like those for the upregulated genes in the CSF lymphoid clusters when comparing PRL-high vs. PRL-low. These included translation, peptide biosynthesis, electron transport chain, (OXPHOS), TCR-signaling, MAPK-signaling, type-I and type-II interferon signaling. Note enrichment of VEGFA-VEGFR2 signaling.

**Extended Figure 13:**
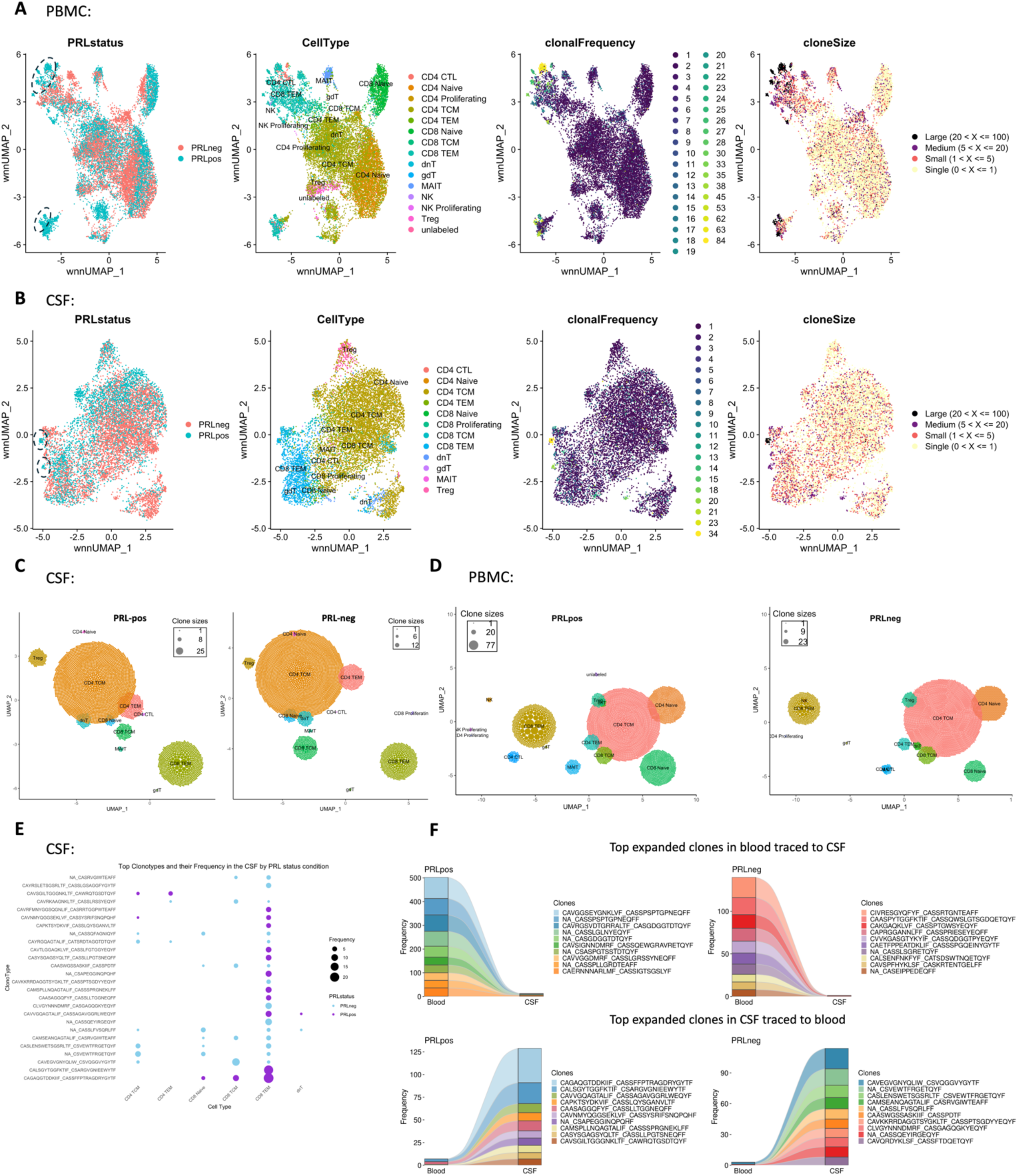
Clonal expansion of CD8-TEM in the blood and CSF of PRL-positive relative to PRL-negative cases, and clonal “continuity” across compartments. a) Weighted neighbor network (WNN) integration of RNA and TCR modalities in PBMC for creating WNN-based UMAPs with only those lymphoid cells depicted with a corresponding TCR-sequence. This follows from Trex^139,140^ based pipeline. Left panel illustrates a wnnUMAP with lymphoid cells specific to PRL-positive state (marked by dashed ovals). Cell annotations and clonal frequency plots reveal CD8-TEM and CD4-CTL cells as being clonally expanded with the largest clone sizes. b) Depiction of wnnUMAP integration of CSF RNA and TCR modalities, with cell types and clonal size representations. c) Ball-packing plots depicting the size of the clones within each cluster with the left panel representing data from the CSF of PRL-positive cases and right panel showing PRL-negative cases. Each ball is one clone, and the clone size represents the number of cells belonging to that individual clone. A clone was defined as a group of cells that share identical nucleotide sequences in both the α- and β-chains of their TCRs. Relative to PRL-negative cases, the CSF of PRL-positive cases was clonally expanded, particularly the CD8-TEM cells. d) Ball-packing plots depicting the size of the clones within each cluster with the left panel representing data from the PBMC of PRL-positive cases and right panel showing PRL-negative cases. Each ball is one clone, and the clone size represents the number of cells belonging to that individual clone. Note the increased sizes of each clone in the CD8-TEM and CD4-CTL cell populations in PRL-positive versus PRL-negative conditions. e) Dotplot summarizing the top 15 clonotypes with the CDR3-aa (complementarity-determining region 3– amino acid) sequences shown for both the α- and β-chains in PRL-positive versus PRL-negative samples. Again, evident is the CD8-TEM clonal expansion in the CSF of PRL-positive patients. f) Flow diagram showing the top expanded clones in the blood traced to CSF (top panel) in PRL-positive and PRL-negative cases (left and right respectively). Bottom panel shows the top expanded clones in the CSF traced to blood in the PRL-positive and PRL-negative cases (left and right respectively).

**Extended Figure 14:**
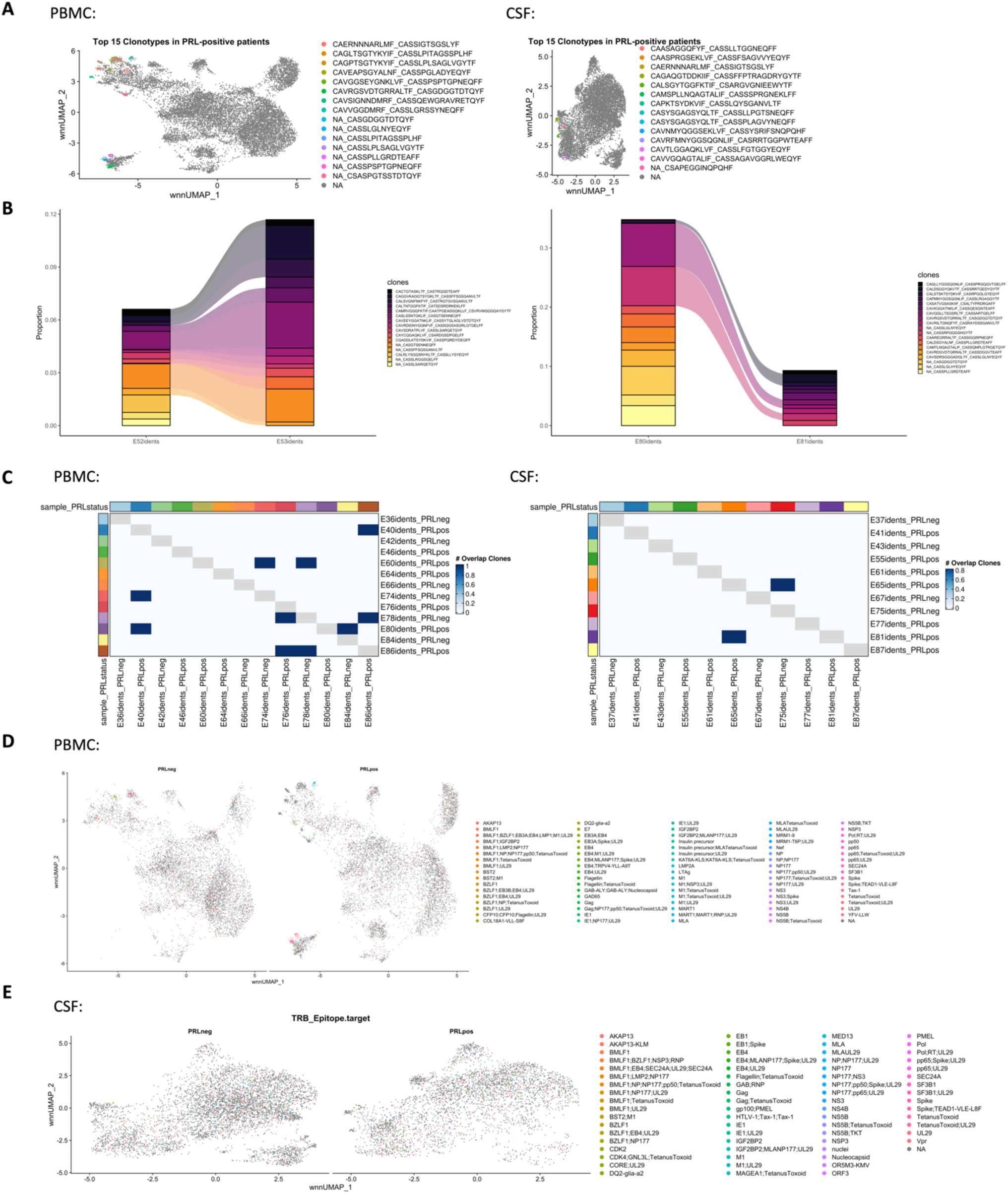
Prediction of TCR-specificities and clonal expansion of CD8-TEM cells in chronic active MS. a) Top 15 in blood clonotypes mapped back onto the wnnUMAP of PBMC (left panel) and top 15 clonotypes mapped back onto the wnnUMAP of CSF (right panel). b) Similar clones are seen in PBMC and CSF cells derived from the same patient as seen in the panels on the left (E52/E53) and right (E80/E81). c) Heatmaps showing clonal overlap across samples in the blood (left panel) and CSF (right panel). d) wnnUMAP of PBMC split by PRL status representing the TCR-specificities (epitope predictions) and the list of epitopes. e) Epitope predictions (TCR specificities) shown for the CSF cells in PRL-positive versus PRL-negative cases.

**Extended Figure 15:**
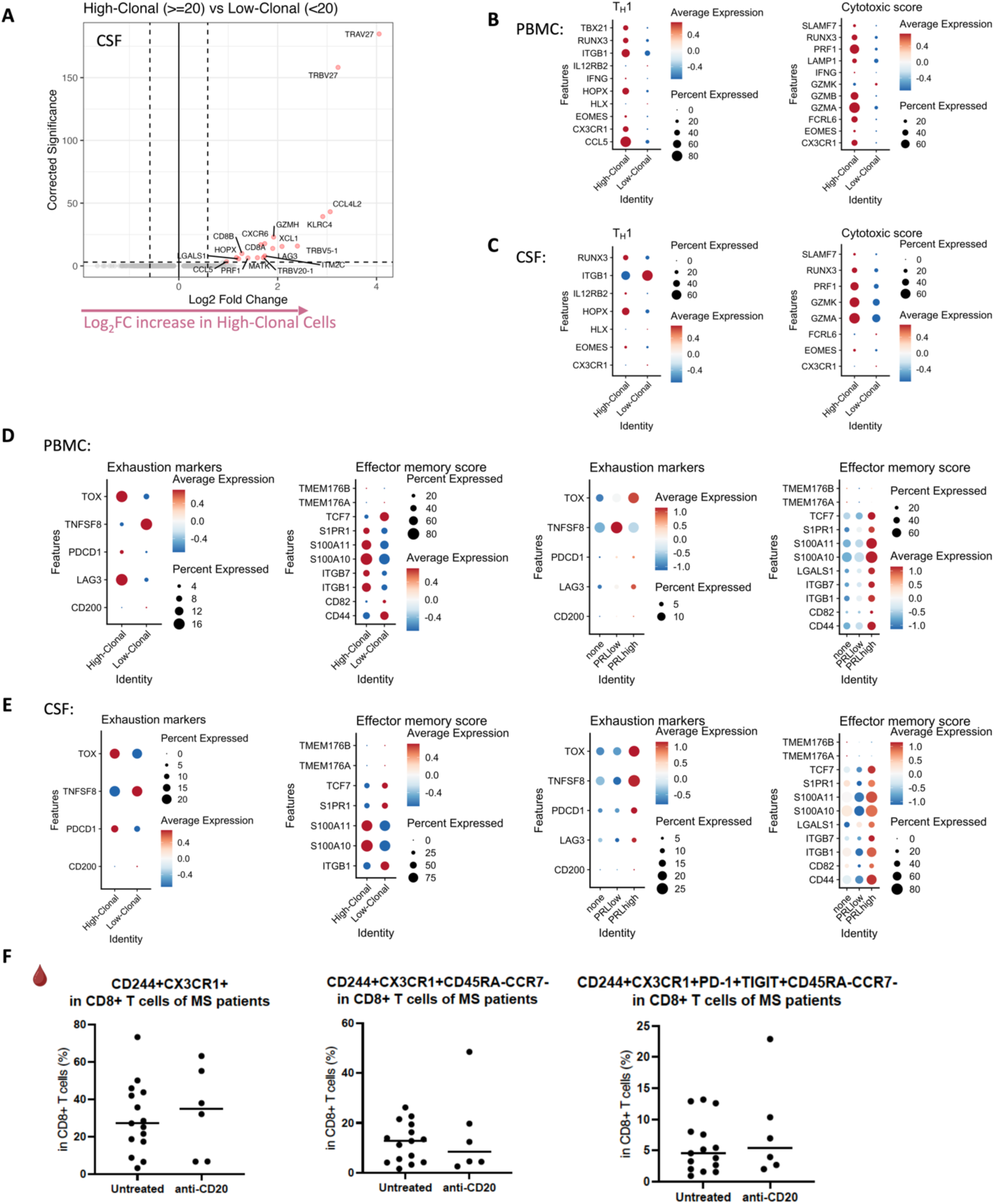
Exhaustion and effector memory states of lymphoid cells in the blood and CSF of cases with PRL. a) Volcano plot showing DEG across high-clonal (clone size ≥ 20) versus low-clonal (< 20) cells in CSF. b) Dotplot showing average expression of T_H_1 and cytotoxicity pertinent genes in high-clonal versus low-clonal cells in blood. c) Dotplot representing T_H_1 and cytotoxicity signature genes in high-clonal versus low-clonal cells in CSF. d) Dotplot showing expression of genes pertinent to exhaustion and effector memory state in the blood across: (1) highvs. low-clonal; and (2) PRL categories. e) Dotplot showing expression of genes pertinent to exhaustion and effector memory state in the CSF across: (1) highvs. low-Clonal; and (2) PRL categories. All panels are related to Figure 3. f) Frequency of CD244^+^CX3CR1^+^ (left panel), CD244^+^CX3CR1^+^CD45RA^-^CCR7^-^ (middle panel) and CD244^+^CX3CR1^+^PD-1^+^TIGIT^+^CD45RA^-^CCR7^-^ (right panel) subsets relative to CD8^+^ T-cells in the blood across untreated and inactive MS vs. B-cell depleted cases.

**Extended Figure. 16:**
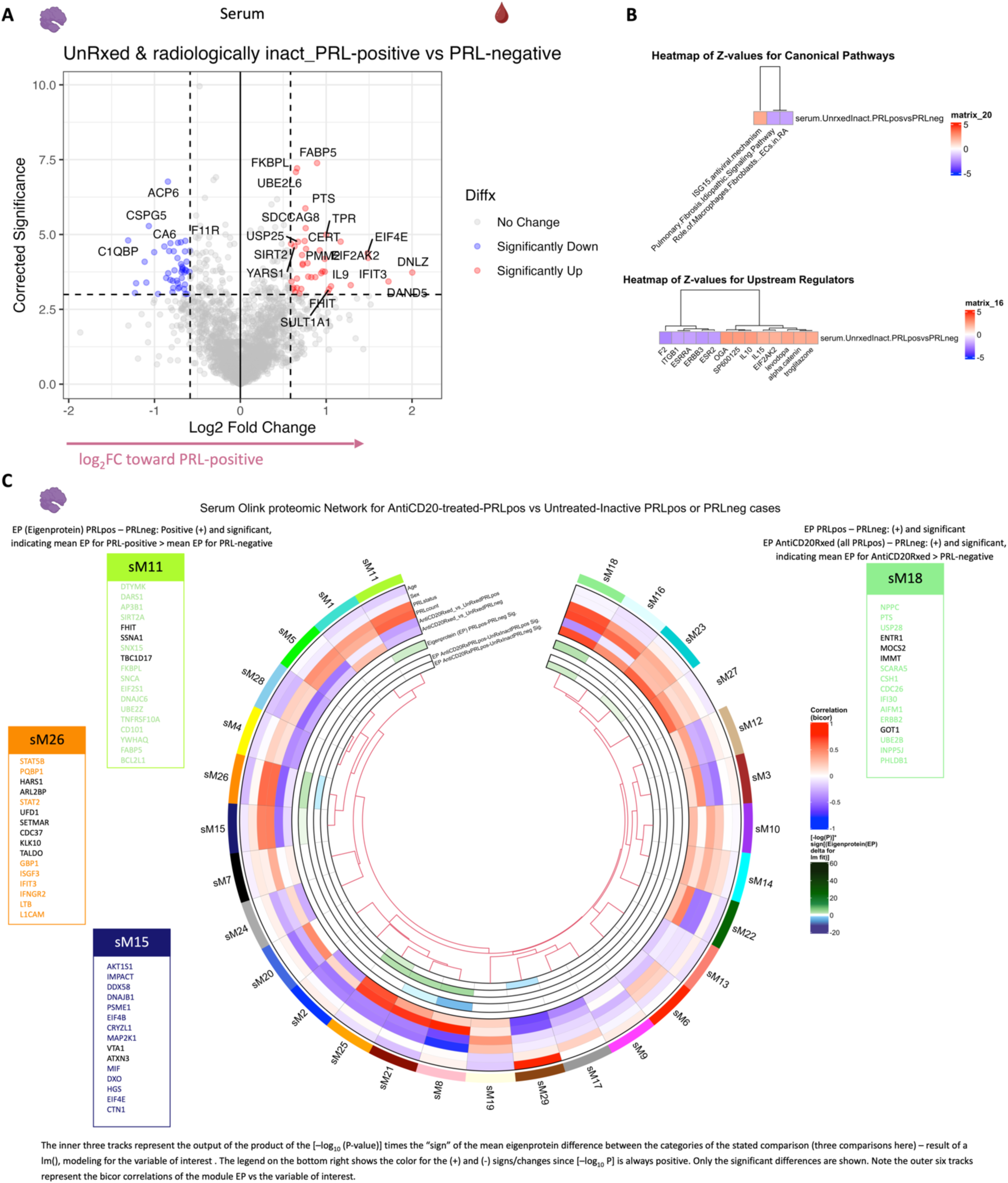
Serum proteomic network reveals signatures associated with the PRL-state. a) Differential abundance of proteins in serum are depicted in a volcano plot comparing PRL-positive vs. PRL-negative cases. b) Z-score enrichment of pathways (top panel) and upstream regulators (bottom panel) for differentially expressed proteins in blood across PRL-positive vs. PRL-negative cases. c) A serum protein co-expression network of 1463 protein assays from Olink measured across 21 individuals (10 untreated and inactive PRL-positive, 7 untreated and inactive PRL-negative, and 4 anti-CD20-antibody treated PRL-positive MS cases). The module eigenprotein (EP) or the first principal component of module expression was correlated with age, sex, PRL status (PRL-positive vs. PRL-negative), PRL count (total number of PRL on the MRI at the time of sample acquisition), anti-CD20-treated against PRL-positive (anti-CD20-treated vs. PRL-positive *only*), and anti-CD20-treated against PRL-negative (anti-CD20-treated vs. PRL-negative *only*) status. These correlations are shown in the outer six tracks with the corresponding bar on the upper right (“Correlation (bicor)”). A linear model was used to assess the effect of the variable of interest (PRL-positive vs. PRL-negative, AntiCD20-treated vs. PRL-neg, and AntiCD20-treated vs. PRL-positive) on EP while adjusting for covariates. The inner three tracks show the signed [-log_10_(one-way ANOVA p-value)] for each comparison (EP PRLpos-PRLneg Sig.; EP AntiCD20RxPRLpos-UnRxInactPRLpos Sig.; and AntiCD20RxPRLpos-UnRxInactPRLneg Sig.), with effect size reflecting the magnitude of EP differences across groups. The lower bar on the right represents this, where color indicates directionality. We used this to infer significant PRL-pertinent modules, which might not be modulated with B-cell depletion.

**Extended Figure 17:**
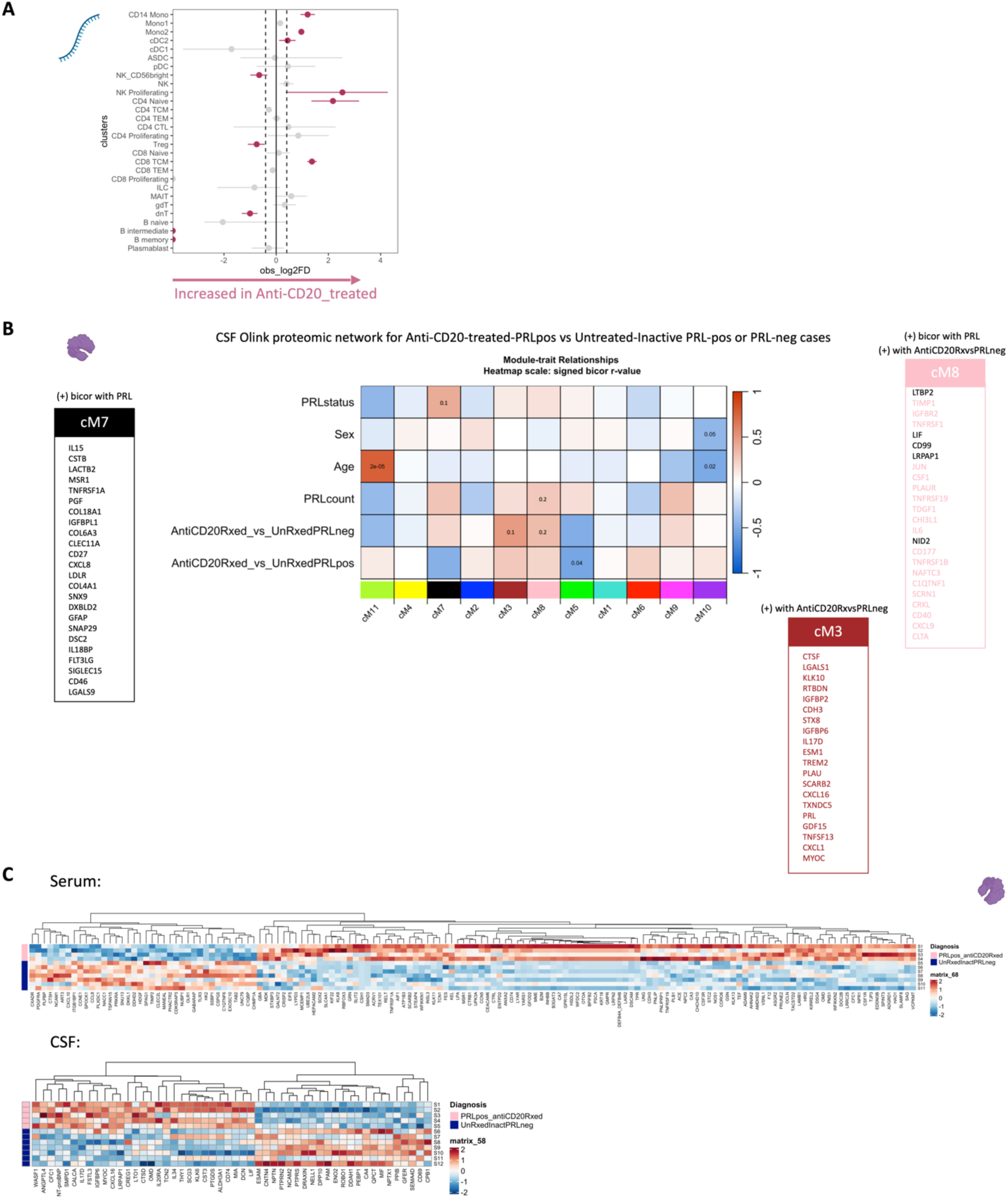
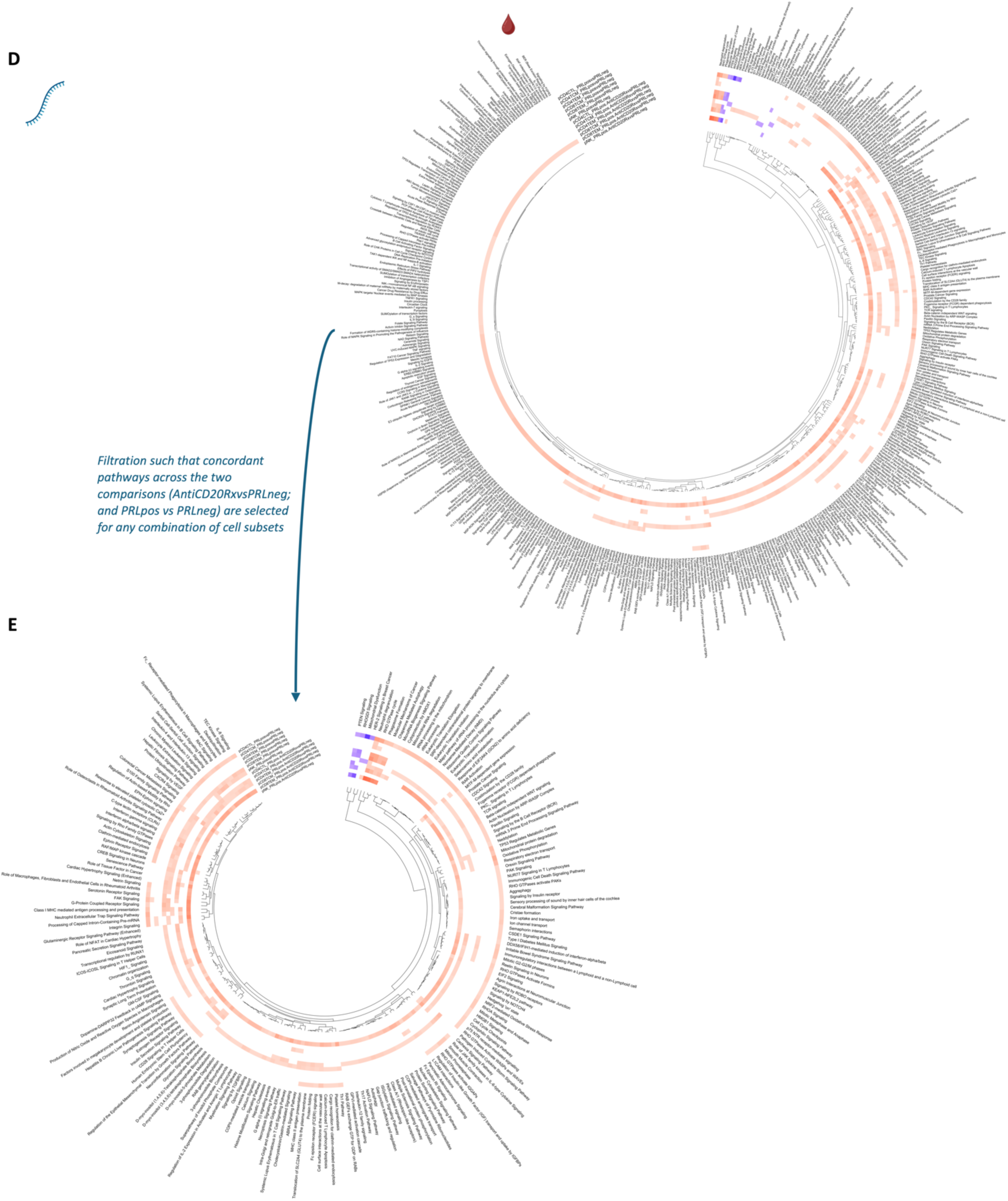
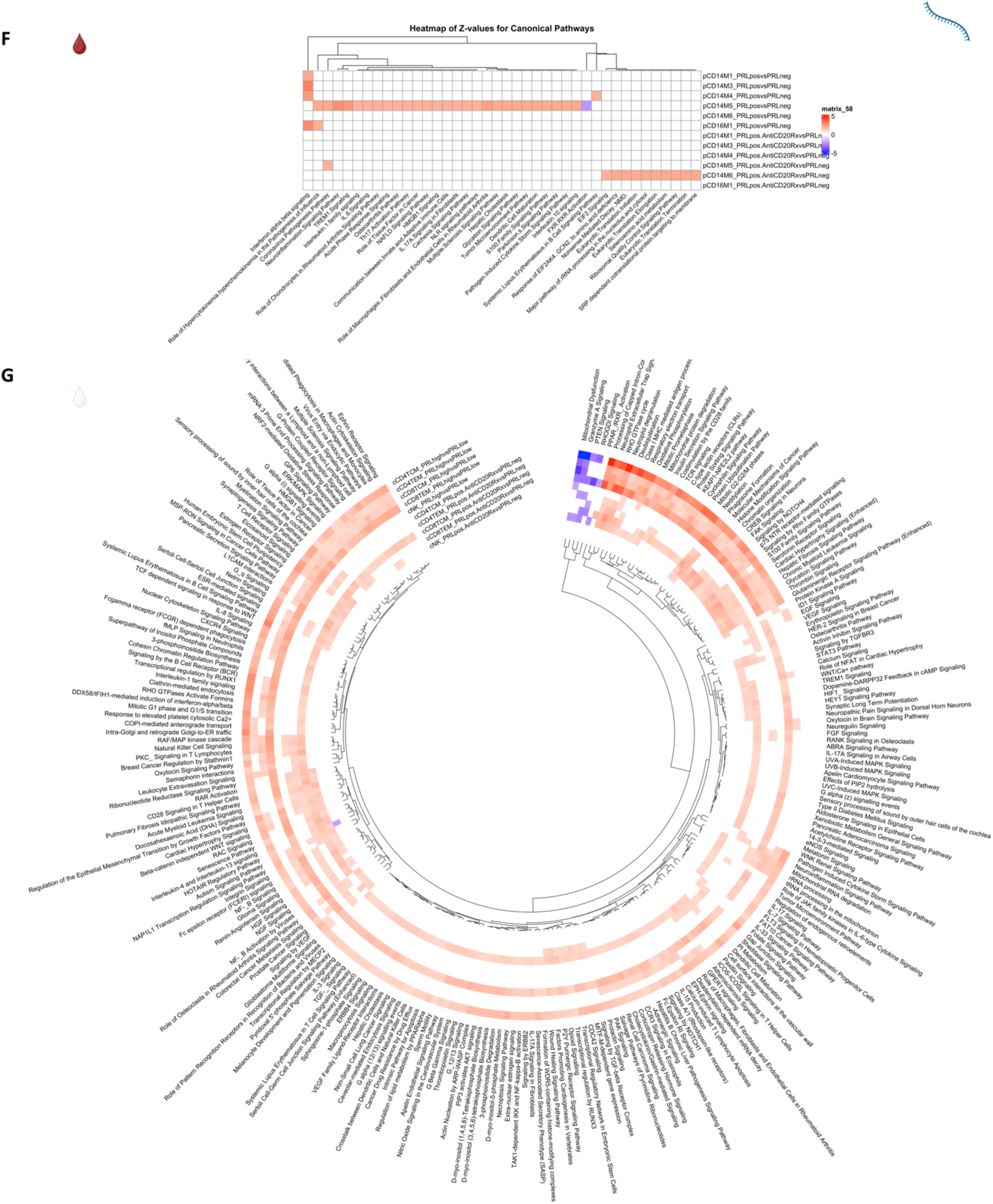
Pathway analyses to dissect correlates of PRL-related pathology which remain unaddressed with B-cell depletion; CD8-TEM cells emerge as central players. a) Lineplot showing increased proportion of CD14 Mono, Mono2, and CD8-TCM cells in the CSF of B-cell-depleted (*all PRL-positive*) versus PRL-positive cases. b) Heatmap showing module-trait correlation for a CSF protein co-expression network of 758 protein assays from Olink platform measured across 22 samples (10 untreated and inactive PRL-positive, 7 untreated and inactive PRL-negative, and 5 anti-CD20-antibody treated MS patients (all PRL-positive)). The module eigenprotein (first principal component of module expression) was correlated with PRL status (PRL-positive vs PRL-negative), PRL count (total number of PRL on MRI at the time the samples and data were curated), anti-CD20-treated against PRL-negative and anti-CD20-treated against PRL-positive. Despite age- and sex-adjustment of proteins prior to network construction, there were correlations of the modules with age. Note, all anti-CD20-treated patients in the cohort were PRL-positive at the time of sample and data acquisition. c) Heatmap showing proteins enriched in the serum (top panel) and CSF (bottom panel) in CD20-depleted cases (*all PRL-positive*) vs. untreated and inactive PRL-negative cases. This is related to Figure 4A (note the fold-change in this analysis was 1.25). d) Circular heatmap demonstrating Z-score enrichment of pathways in peripheral lymphoid cells for DEG across B-cell-depleted (*all PRL-positive*) vs. PRL-negative and PRL-positive vs. PRL-negative comparisons. Enriched pathways in B-cell-depleted (*all PRL-positive*) vs. PRL-negative, which are simultaneously concordant with the other comparison (PRL-positive vs. PRL-negative), likely reflect the PRL pathology unaccounted for by B-cell depletion. e) Circular heatmap of peripheral lymphoid cells illustrating Z-scores of canonical pathways for DEG concordantly enriched or depleted across B-cell depleted (*all PRL-*positive) vs. PRL-negative and PRL-positive vs. PRL-negative comparisons. This resulted from filtration of pathways in panel d. Note the concordant terms including IFN-α/β, IFN-γ, ISGylation, TCR, EIF2 signaling, T_H_1 pathway, neddylation, and MHC-II antigen presentation, which are all enriched in B-cell-depleted (*all PRL-positive*) vs. PRL-negative and the PRL-positive vs. PRL-negative comparisons. f) Heatmap showing Z-score enrichment of pathways for DEG across B-cell-depleted (*all PRL-positive*) vs. PRL-negative and PRL-positive vs. PRL-negative cases in peripheral myeloid cells. g) Circular heatmap of the CSF lymphoid subclusters illustrating Z-score enrichment of canonical pathways for DEG across B-cell-depleted (*all PRL-positive*) vs. PRL-negative and PRL-high vs. PRL-low comparisons. The concordant terms map onto CD8^+^ T cells, primarily including IFN-γ, IFN-α/β signaling, NF-κB signaling, ERK/MAPK signaling, JAK family kinases in IL6, and TCR signaling among others. CD4-TCM and CD8-TCM subclusters also showed enrichment of pathways suggestive of persistence of “memory” in the CSF in chronic active MS.

**Extended Figure 18:**
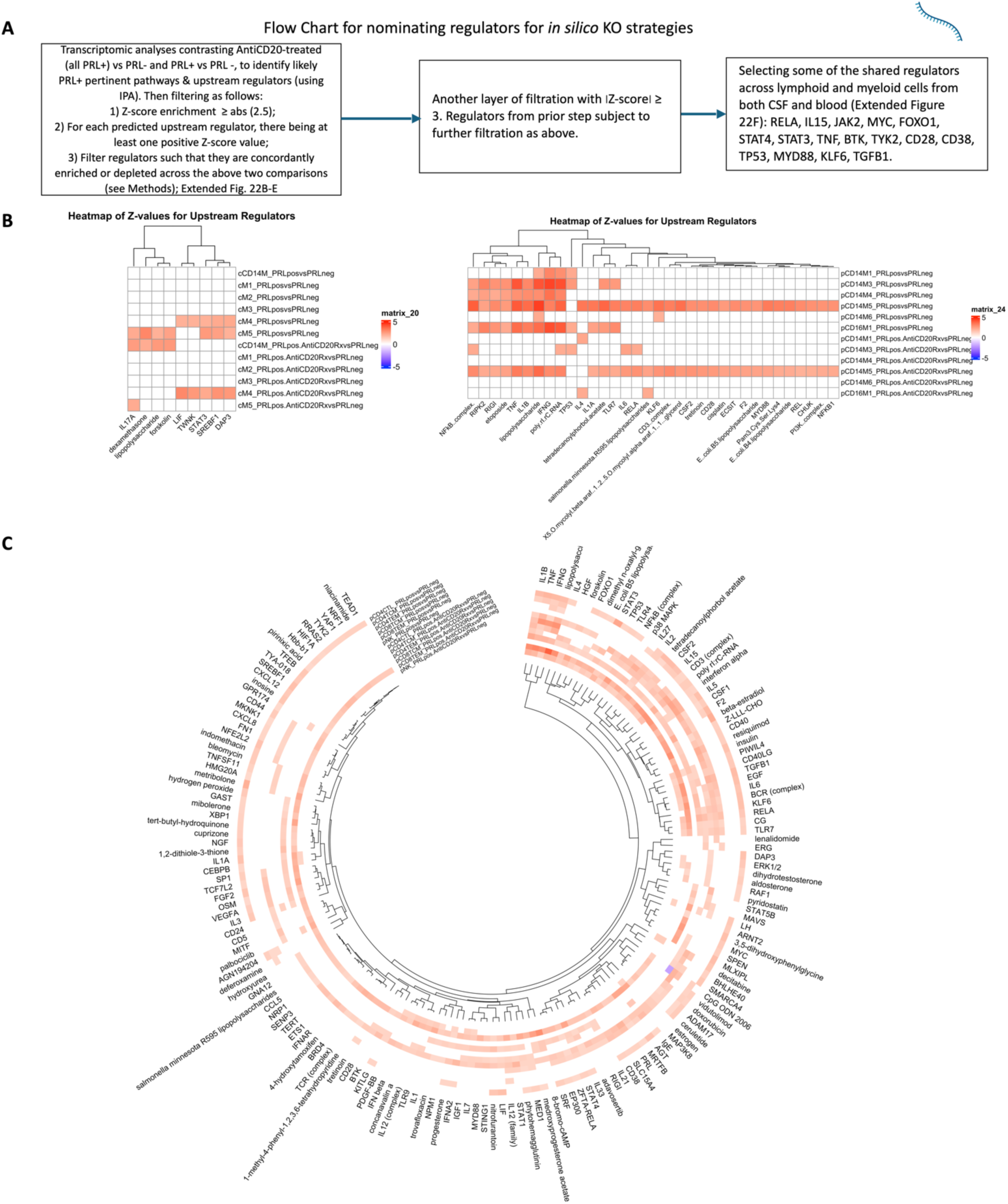
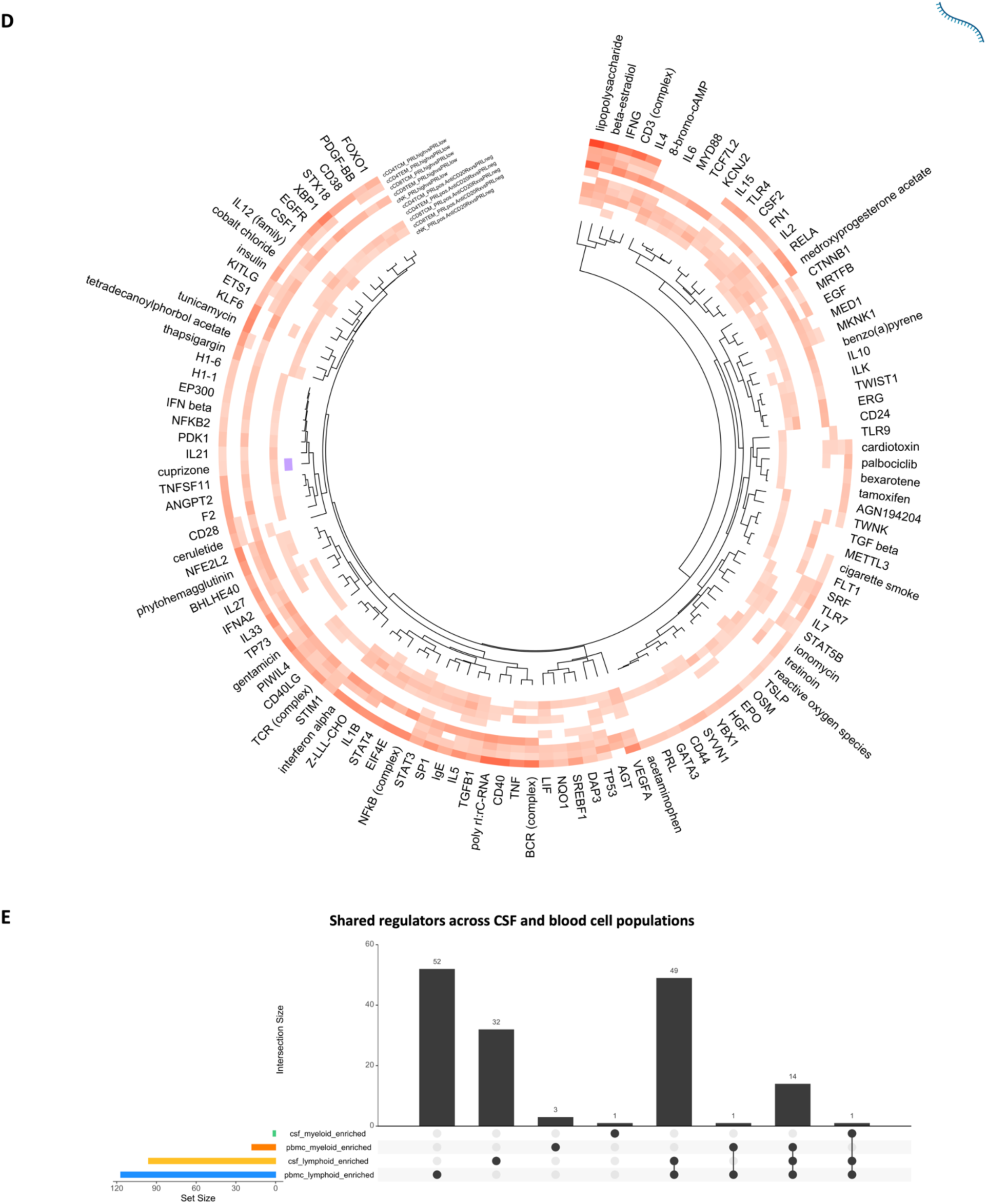
Nominating new targets for ameliorating chronic neuroinflammation. a) Flow chart schematic illustrating the method used for nominating the upstream molecules regulating PRL-positive pertinent pathology. These nominated upstream regulators were subsequently used for *in silico* deletion strategies (see Figure 4E-G and Methods) to curate genes lists involved in PRL-pathology that remain unaffected by *in vivo* and *in silico* B-cell depletion but might be modulated by the predicted regulatory molecules, as implied by their *in-silico* deletion. Hence, possible new targets for chronic neuroin-flammation in MS. b) Heatmap showing Z-scores of candidate regulators in CSF (left panel) and blood (right panel) myeloid cells across the B-cell depleted vs. PRL-negative and PRL-positive versus PRL-negative comparisons. c) Circular heatmap illustrating top candidate regulatory molecules driving likely PRL-pertinent pathology unperturbed by B-cell depletion, in the lymphoid cells of blood. d) Heatmap representing top predicted regulators in the CSF lymphoid cells across B-cell depleted vs. PRL-negative and PRL-high vs. PRL-low comparisons. e) Upset plot showing shared regulators across CSF and blood cell populations.

**Extended Figure 19:**
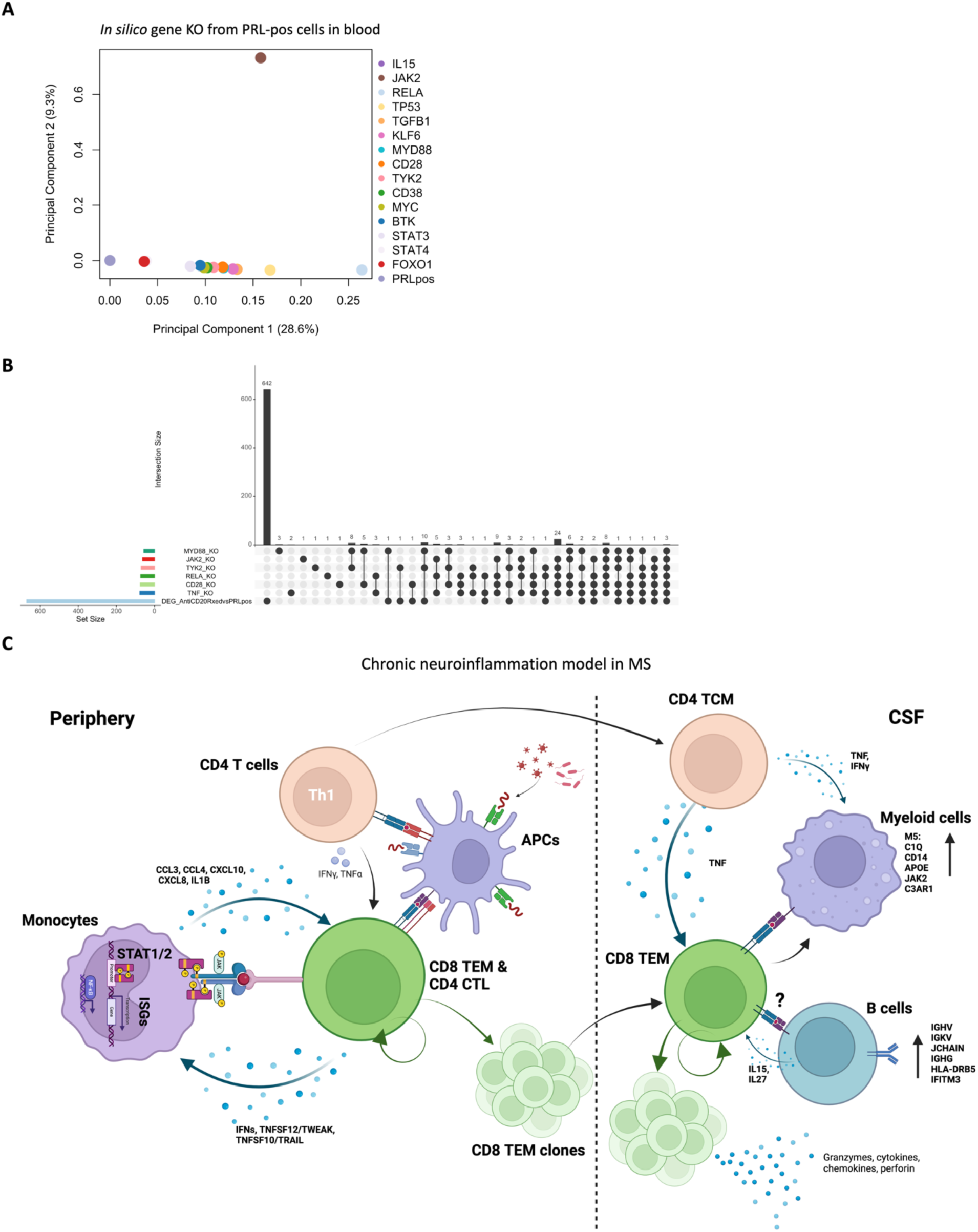
Curating genes likely linked to PRL pathology and predicted targets for modulation. a) PCA plot of the simulated distances from comparing the gene regulatory network (GRN) of the original PRL-positive state in the blood vs. the GRNs resulting from *in-silico* KO of the predicted regulators. b) Upset plot showing the overlap of DEG in *in vivo* B-cell depletion — using anti-CD20-treated vs. PRL-positive comparison in CSF — with the significantly perturbed genes resulting from *in silico* deletion of the regulators in PRL-positive (untreated and inactive) cells of CSF. Of interest are the genes that are not affected by CD20-depletion and are only predicted to be affected by *in silico* KO of *MYD88*, *TYK2*, *JAK2*, *RELA*, *CD28*, and *TNF* (Figure 4E). c) Proposed model for the maintenance of chronic neuroinflammation in MS.

## Notes

### Competing Interest Statement

The authors have declared no competing interest.

