## Supplementary Information for "Maintenance of chronic neuroinflammation in multiple sclerosis via interferon signaling and CD8 T cell-mediated cytotoxicity"

Table of contents

- Cluster annotations and pathway analyses
- Combined object and immune cells across CSF and blood compartments
- IL15 regulates immune responses in chronic active MS
- References
- Supplementary Figures

**Cluster annotations and pathway analyses**

**CSF cells**

L1 annotations of the CSF cells involved manual and Azimuth-based annotations. Subclustering was performed on the myeloid, T-cell and NK-related, and B-cell lineages (enclosed by dashed green, pink and blue lines) to obtain L2 cluster populations (Extended Figure 1A, C and Figure S5A-D).

<u>T-cell and NK-related lineages:</u> Among the T-cells, CD4 subsets were annotated as $CD4^+$ Naïve, $CD4^+$ Central Memory T (CD4 TCM), $CD4^+$ Effector Memory (CD4 TEM) and $CD4^+$ T-cells with Cytotoxic activity (CD4 CTL). Cluster defining genes for CD4 Naïve cells included *CCR7* (responsible for homing of T-cells to lymphoid organs), *FHIT*, and *NOG*[1]. CD4 TCM and CD4 TEM both expressed *INPP4B*, *AQP3*, *FLT3LG*, *MAL*, and *CD40LG*, with CD4 TEM additionally expressing *GZMK*, *IFNG-AS1*, *LYAR*, and *DPP4* as the cluster-defining genes. CD4 CTL expressed *GZMH*, *NKG7*, and *LYAR*[2]. $CD8^+$ cells were marked by *CD8A*, *CD8B* which serve as co-receptors for the TCR[3], and *KLRK1*, which encodes a cell-surface receptor expressed on activated $CD8^+$ T cells[4]. *GZMH*, *GZMK, NKG7*, *CTSW,* and *CST7* genes differentiated CD8 TEM from CD8 TCM, reflecting the cytotoxic effector profile of CD8 TEM cells[5,6]. *NELL2*, with its role in $CD8^+$ T-cell differentiation, was a marker of CD8 Naïve and CD8 TCM cells[7] but was also expressed by dnT cells. $CD8^+$ proliferating cells, albeit a small cluster of cells, were marked by *CDT1*, *CLSPN, GINS2*, *UHRF1*, and *DTL* genes, which are involved in cell cycle regulation and epigenetic modifications[8]. Similarly, a $CD4^+$ proliferating cell cluster comprised a small number of cells and was marked by *TYMS*, *RRM2*, *TK1*, *PCLAF,* and *UBEC2*, which are associated with cell proliferation and DNA synthesis and repair[8].

dnT cell markers included *INPP4B*, *IFNG-AS1*, *GZMK*, and *TOX*, suggestive of cytotoxic functions, consistent with prior reports[9]. Note, *IFNG-AS1* is a long noncoding RNA (lncRNA) that acts as an enhancer for IFN-$\gamma$ transcription and regulates its production at both transcriptional and translational levels[10–12] . gdT ($\gamma\delta$T-cells) were characterized by *TRDC*, *TRGV,* and *TRDV2* genes, which is expected since these are the TCR delta and gamma genes[13]. Mucosal-associated invariant T (MAIT) cells were identifiable using canonical markers, including *SLC4A10* (bicarbonate transporter with potential roles in maintaining pH and ion homeostasis for MAIT cells) and *TRAV1-2*, which is a TCR $\alpha$-chain variable chain strongly associated with MAIT cells[14,15].

Top markers for innate lymphoid cells (ILC) included *TRDC*, *SPINK2*, *KIT*, *XCL1*, and *XCL2* genes. *XCL1* and *XCL2* as chemotactic cytokines have been shown previously to be markers of ILC, including natural killer (NK) cells[16]. NK and NK_CD56bright cells differentially expressed the canonical markers including *KLRD1*, *KLRC1*, *KLRK1*, *GNLY*, *XCL1*, *XCL2*, *NKG7*, *TRDC*, and *CTSW*, discriminating these cells from the rest[17]. Regulatory T cells (Treg) exhibited the characteristic markers *FOXP3*, *TIGIT*, *IKZF2*, *RTKN2*[18,19]; interestingly, we had a non-coding RNA (ncRNA), *AL136456.1*, as a Treg discriminatory marker as well.

Myeloid cells: L1 CSF myeloid cells were annotated as CD14 Mono, Mono1, Mono2, cDC2, cDC1, ASDC, and pDC. "CD14 Mono" cluster exhibited canonical markers such as *LYZ*, *FCN1*, *S100A9*, *S100A8*, and *VCAN*, suggestive of responses to Toll-like receptor (TLR) signaling, regulation of inflammatory processes, and cell adhesion/migration. Conventional dendritic cells type 2 (cDC2) showed expression of *FCER1A* ($\alpha$-subunit of the high-affinity IgE receptor - Fc$\varepsilon$RI), *HLA-DQA1*, *CST3*, *CD1C*, and *HLA-DQB1* genes, representing of major histocompatibility complex (MHC) class II molecules and lipid antigen presentation[20,21]. Previously defined markers, *XCR1* and *IRF8*, involved in cross-presentation of antigens to $CD8^+$ T cells, development of cDC1 (conventional dendritic cells type 1), and antiviral responses, including regulation of IFN-stimulated genes (ISG), were found as discriminatory markers of cDC1 from cDC2 in our dataset[22,23] – although we had additional discriminatory markers (*LGALS2*, *DNASE1L3*, *IDO1* and *TACSTD2*) as well. *TCF4* as one of the master regulator transcription factors (TF) for plasmacytoid dendritic cells (pDC) was found to be expressed in our dataset. In addition, *RHEX*, *CLEC4C*, and *LILRA4* were also expressed by pDC[21]. Presence of SIGLEC6 and PPP1R14A differentiated $AXL^+SIGLEC6^+$ dendritic cells (ASDC) from pDC[21] in our dataset.

The cluster annotated as "Mono1" was marked by *C1QC*, *C1QA*, *C1QB*, *TYROBP*, *HLA-DRA*, *AIF1*, and *FCER1G* genes, while absent to decreased levels of expression of *LYZ*, *HLA-DQA1*, *HLA-DQB1*, and *MEF2C* in "Mono2" parsed the latter from the Mono1 cluster (Extended Figure 1C, Figure S5B). We performed unsupervised subclustering on all myeloid cell populations to obtain L2 clusters (Figure S6), with the overall interpretation of the subclusters and the pathway analyses provided below (see Methods — top 30 cluster defining genes were used):

- CD14M cluster showed enrichment of genes like *FCN1*, *VCAN*, *S100A8*, *S100A9*, *LGALS3*, *DYPD*, and *MT-ND2*. These genes represent terms like leukocyte aggregation, granulocyte chemotaxis, and positive regulation of inflammatory and defense responses (Figure S6B, C).
- M1 cluster showed association with genes responsible for regulation of $K^+$ ion transmembrane transport and monoatomic cationic homeostasis (*HAMP*, *SEL1L2*, *SLC12A5*, and *AL031056.1*).
- M2 was characterized by *TREM2*, *C3*, *A2M*, *SPP1*, *LIPA*, and *FOLR2* as the top marker genes, which represented terms such as regulation of receptor-mediated endocytosis (*C3, APOC1, APOC2*), synapse pruning (*C3*, *TREM2*), positive regulation of leukocyte chemotaxis (*CSF1R*, *CXCL12*, *CALR*), and regulation of lipid metabolic process (*APOC2*, *APOC1*, and *TREM2*). Notably, M3 expressed some of the same genes as M2, but the differentiating genes included *RHOB, TNFRSF1B*, *C5AR1*, *KLF2*, *JUNB*, *CEBPD*, and *ZFP36*. Pathway analyses for M3 showed response to lipopolysaccharide/LPS (*ZFP36*, *SLC11A1*, *CD14*, *TNFRSF1B*), phagocytosis and endocytosis (*MSR1*, *SLC11A1*, *CD14*, *CXCL16*) and inflammatory response (*CSF1R*, *SLC11A1*, *C5AR1, CD14*). The transcriptional profile of M3 bears some resemblance to previously described 'microglia-like' cells in the CSF[24] and to microglial phenotypes previously described in various neurodegenerative disorders[25–27].
- M4 cluster enriched for markers like *IL32*, *CD3E*, *IL7R*, *CCL5, SKAP1*, and *CD247*, responsible for antigen receptor-mediated signaling (*ITK*, *LCK*, *CD247*, *SKAP1*) and T-cell activation (*CD2*, *ITK*, *LCK*, *CD247*).

- Cluster M5 had *PLXDC2*, *MT-ND1*, *NEAT1*, *LRMDA*, *SLC8A1*, *ARHGAP15*, *ZBTB20*, *FOXP2*, *SSH2*, and *ELMO1* as the cluster-defining genes. The pathways enriched included sodium ion transport across plasma membrane (*SLC9A9*, *SLC8A1*), myeloid leukocyte differentiation (*GAB2*, *FOXP1*), and regulation of small GTPase-mediated signal transduction (*STARD13*, *ARHGAP15*, *ARHGAP26*), among others. Note, comparison of cluster M5 to the immune cells from a previously published MRI-informed snRNAseq study showed that the clusters M3 and M5 were transcriptionally most like the microglia inflamed in MS (MIMS) phenotypes (MIMS-iron and MIMS-foamy) found at the CAL edge (Figure 2C)[28]. In addition, M5 cluster transcriptionally bore resemblance to the microglia isolated from human Alzheimer's disease (AD) brains (Extended Figure 3C)[29].
- Cluster M6 likely represents "cycling myeloid cells," enriched in terms like microtubule cytoskeleton organization involved in mitosis (*STMN1*, *PLK1*, *NUSAP1*, *CDK1*, *BIRC5*, *AURKB*, *SPC25*), mitotic cell cycle phase transition (*CCNA2*, *CCNB2*, *UBEC2*, *PLK1*, *CDK1*, *PKMYT1*, *CDKN3*), mitotic spindle organization (*TPX2*, *STMN1*, *PLK1*, *BIRC5*, *AURKB*, *SPC25*) and cell cycle G2/M phase transition (*CCNA2*, *PLK1*, *CDK1*, *PKMYT1*, *AURKB*), among others. The cluster-defining genes included *STMN1*, *TYMS*, *PCLAF*, *UBEC2*, *CDK1*, *NUSAP1*, *TK1*, *RMM2*, *TOP2A*, and *CDKN3.*
- M7 was marked by *ISG15*, *IFIT1*, *IFIT3*, *MX1*, *CXCL10*, *IFI4L*, *STAT1, GBP1*, *OAS3*, and *CMPK2* genes, and enriched in pathways including defense to virus/symbiont (*IFITM1*, *STAT1*, *MX2*, *MX1*, *IFI6*, *ISG15*, *IFIT1*, *USP18, IFIT3, IFI44L, IFIT2, OASL, CXCL10, OAS2*, *OAS3*), response to IFN-$\beta$ (*IFITM1*, *PLSCR1*, *STAT1*, *XAF1*), and response to type-I IFN (*MX1*, *ISG15*, *USP18*).
- Cluster M8 was defined by terms including regulation of TLR-9 signaling and negative regulation of viral-induced cytoplasmic pattern recognition receptors (*PTPRS*, *LILRA4*), and differentiated from other clusters by *RHEX*, *BCL11A*, *UGCG*, *LILRA4*, *PLAC8*, *JCHAIN*, and *CLEC4C*. M9 was determined by presence of *CCL3*, *CCL4L2*, *IL1B*, *CCL4*, *EGR1*, *CCL3L1*, and *EGR2*. Pathway analyses showed enrichment for cellular response to interleukin-1/IL1 (*EGR1*, *CCL8*, *CCL3L1*, *IL1B*, *CCL4*, *CCL3*, *CCL2*), lymphocyte chemotaxis (*CH25H*, *CCL8*, *CCL3L1, CCL4, CCL3, CCL2*), regulation of ERK1 and ERK2 cascade/positive regulation of MAPK cascade (*PDGFRA*, *CCL8*, *CCL3L1*, *IL1B*, *CCL4, CCL3*, *CCL2, FGF10*), and response to IL1.
- M10 was defined by tryptophan metabolism (*IDO2*, *IDO1*) and BMP signaling pathway (*NOG*, *GDF7*).
- M11 was defined by *CCR7*, *LAMP3*, *BIRC3*, *CLLU1*, *LAD1*, *NOS1, NCCRP*, and *TFP12*, characterized by enrichment of processes like negative regulation of dendritic cell apoptosis, response to prostaglandin E, and nitric oxide (*CCR7*, *CCL19*).
- Cluster cDC2 enriched for regulation of leukocyte/T-cell mediated cytotoxicity (*CD1E*, *CD1C*), response to cytokines (*CIITA*, *FLT3*, *AFF3*)*,* positive regulation of $Ca^{2+}$ ion transmembrane transporter activity (*P2RY6*, *AP1B1*), C-type lectin receptors (*CLEC10A*), cytokine receptor activity (*FLT3*, *IL1R2, IL2RG*), and additional gene markers including *FCER1A*.
- Amongst the top genes discriminating cDC1 from cDC2 cluster included *XCR1*, *LGALS2*, and *IDO1*. Cluster cDC1 enriched for genes involved in regulation of mitotic nuclear division (*EPGN, RGCC*), tertiary granule and clathrin-coated endocytotic vesicle (*TCN1*, *EPGN*).

<u>B-lymphoid lineage cells:</u> Gene markers common to L1 annotations of B-naïve, B-intermediate and B-memory cells included *MS4A1*, *BANK1*, *CD79A*, and *RALGPS2*[30–32]. Markers specific to B-naïve cells included *TCL1A* and *IGHM*, with *BLK* being discriminatory for B-memory cells versus other B-cell types. *AFF3* was expressed to a lesser extent by B-memory versus other clusters. Plasmablasts were enriched in *JCHAIN*, *MZB1*, *IGHA1*, *TXNDC5*, and *TNFRSF17*

genes[32]. For detailed characterization, subclustering was performed with the following clusters and functions (Figure S7):

- Cluster b1 is enriched in B-cell-receptor signaling and B-cell activation (*BLK*, *CD79B*, *MS4A1*, *BANK1*), response to cytokines (*SELL*, *CXCR4*, *AFF3*), regulation of signal transduction (*GNG7*, *CD24*, *CD55*), and positive regulation of T-cell proliferation (*TNFRSF13C*, *CD24*). The cluster-defining genes included *CXCR4*, *VPREB3*, *AFF3*, *TNFRSF13C*, *NIBAN3*, *TSC22D3*, *LINC00926*, *SNHG7*, and *SELL*.
- Cluster b2 was defined by expression of *IGKV1D-8*, *CD3E*, *IL7R*, *IGHV4-59*, *IL32*, *IGKV3-15*, *CST3*, *FYB1*, *GIMAP7*, *CD3G*, *INPP4B*, *IGHV4-61*, *IGKV3-11*, *GIMAP4*, *LINC00861*, *CD3D*, *HIST1H1E*, *MS4AGA*, and *KIF5C*. Pathways enriched for this cluster included T-cell activation, antigen-receptor mediated signaling pathway, and positive regulation of IL4 and FcεR-signaling pathway. Transcriptionally, cluster b2 most resembles the cells labeled as Plasmablast in L1 annotation (Figure S9E). Note, cluster b2 is also marked by IL7R, which has been shown to be expressed in early stages of B-cell lineage development[33,34]. Interestingly, this cluster also showed expression of CD3-pertinent genes, which may be suggestive of a bi-phenotypic lymphocytic subset displaying both T- and B-cell functions[35], considering prior reports of the presence of CD20+ T cells in the CSF and blood of MS patients[36], and the aberrant presence of T-cell-associated antigens in B-cell lymphomas[35,37], or vice versa[38]. This cluster also showed expression of immunoglobulin genes (Figure S7B), suggestive of B-cell function. The presence of canonical T-cell transcripts in b2 cluster could also be a possible result of prolonged contact between T and b2 cells. A prior study also reported CD3-positive B cells being a consequence of ex vivo storage of PBMC samples[39]; however, it is pertinent to note that all the samples used (CSF and PBMC) were fresh in our study.
- Cluster b3 enriched for markers like *SOX5*, *ACP5*, *TNFRSF13B*, *ANXA4*, *CXCR3*, *FGR*, *ADGRG5*, *HMOX1*, and *RGCC* (Figure S7B). The terms associated with b3 cluster included positive regulation of cytokine production (*CD86*, *PYCARD*, *FGR*, *RGCC*, *SYK*, *HMOX1*, *POU2F2*), regulation of phagocytosis (*PYCARD*, FGR*, HCK*, *SYK*), Fcγ-receptor signaling pathway (*FGR*, *HCK*, *SYK*), and B-cell homeostasis (*TNFRSF13B*, *GAPT*)[40,41].
- Cluster b4 was characterized by mediation of the endoplasmic reticulum (ER)-associated degradation (ERAD) pathway (*XBP1*, *HSPA5*, *SELENOS*, *DERL3*, *MAN1A1*), response to ER stress (*XBP1*, *HSPA5*, *SELENOS, CREB3L2*, *PDIA4*), ubiquitin-dependent ERAD pathway, and negative regulation of apoptotic process (*MYDGF*, *XBP1*, *HSPA5*, *CD38*, *IGF1*, *TXNDC5*)[42–44]. Since ERAD pathway is critical for transitioning of large pre-B cells to small pre-B cells, it could be possible that these cells represent a lineage transition toward small pre-B cells, subsequently into immature B and/or mature B cells.
- Cluster b5 represented B-cell activation and differentiation (*PRKCB*, *PTPRJ*, *PAX5*, *HDAC9*), regulation of interleukin-2 (IL2) production (*HDAC9*, *CARD11*, *RUNX1*), regulation of B-cell receptor signaling (*PRKCB*, *FCRL3*), and negative regulation of $Na^+$ transmembrane transporter activity (*CAMK2D*, *PRKCE*)[45,46]. Significant cluster-defining markers included *PRKCB*, *PTPRJ*, *PAX5*, *HDAC9*, *CAMK2D*, *PRKCE*, *FCRL3*, *EIF4G3, SLC9A7*, and *SIPA1L3*[30,33].
- Cluster b6 — a very small cluster — enriched for *LAMP3*, *FSCN1*, *TPFPI2*, and *MGLL*. It showed terms like monocyte chemotaxis and response to tumor necrosis factor/TNF (*CCL22*, *TNFRSF11A*, *CCL19*) and tryptophan metabolic process (*IL4I1*, *IDO1*).

**PBMC**

L1 annotations of the PBMC were performed using a reference dataset[47]. Subclustering was performed on the myeloid, T-cell and NK-related, and B-cell lineages to obtain L2 cluster populations (Extended Figure 1B, D and Figure S5E-H). Most of the L1 cluster-defining genes for the PBMC were the same as L1 CSF cluster-defining genes, particularly in the T-cell and NK-related lineages and B-lymphoid lineages, unless otherwise stated.

T-cell and NK-related lineages: For PBMC in the periphery, cluster-defining genes for CD4 Naïve were the same as for CSF: *CCR7* and *FHIT*[1], but also *LEF1* and *TCF7* (encodes TCF1 – a key transcription factor in T-cell development and differentiation)[48,49]. For CD8 Naïve cells, the cluster-defining genes were similar with the addition of *CD8A* and *CD8B*. CD4 TCM and CD4 TEM were both defined by expression of the canonical marker *IL7R*[34], but CD4 TEM expressed *GZMK* while CD4 TCM *INPP4B* and *MAL*. CD4 CTL expressed similar markers as CSF but had additionally the expression of *CCL5*[2]. CD8 TEM was differentiated from CD8 TCM by expression of *GZMH*, *NKG7, GNLY*, *GZMA*, *CCL5/RANTES* in the effector memory phenotype, suggestive of a cytotoxic profile like that seen in the CSF[5,6]. dnT was characterized by expression of *IFNG-AS1* and *GZMK*. $\gamma\delta$T expressed TCR-delta and gamma genes, in addition to *DUSP2*. MAIT were identifiable by *KLRB1*, *SLC4A10*, and *DUSP2*[14,15,50]. NK and NK_CD56bright expressed the same canonical genes (*KLRD1*, *GZMH, GNLY*, *NKG7*) as described above for CSF, with *XCL1* and *XCL2* discriminating latter from the prior in the blood[49]. Treg were identifiable using essentially the same markers in the periphery as the CSF.

Myeloid lineage cells: L1 annotations of myeloid lineage cells in the blood included CD14 Mono, CD16 Mono, cDC1, cDC2, ASDC, and pDC. Note, we have been characterizing pDC as part of myeloid lineage cells despite there being evidence of pDC being derived from common lymphoid progenitors in addition to myeloid-based lineage, based on the clustering[51–54]. cDC2 and cDC1 were identifiable by the canonical expression of genes like *HLA-DQA1*, *HLA-DQB1*, *HLA-DRB5*, and *CPVL*[20,21,55], with *FCER1A, S100A9*, and *VCAN* expression in cDC2 parsing it from cDC1. cDC1 cells were characterized by the expression of CLEC9A, *IDO1*, *DNASE1L3* and *XCR1*. CD14 Mono was defined by *S100A8, S100A9, VCAN,* and *FCN1* expression which is identical to CSF, while CD16 Mono cells were characterized by the expression of *FCGR3A (CD16/Fc$\gamma$RIII*; also expressed by NK), *MS4A7*, *SMIM25*, *SERPINA1*, and *TCF7L2*[21]. ASDC in the periphery expressed the same markers as in the CSF (Figure S5B, F). pDC were identifiable using *TCF4*, *PLD4*, *CCDC50*, *JCHAIN*, *LILRA4*, *IL3RA* (*CD123*), *CLEC4A*, and *SERPINF1* expression[52–54].

We subclustered the above-mentioned myeloid cells for better understanding of cell phenotypes (Figure S8A-C). The following are the subclusters defined:

- CD14M_1 was characterized by negative regulation of peptidase/endopeptidase activity (*SERPINB10*, *SERPINB2*, *SERPINI2*, *NRG1*) and lipid oxidation (*ALOX15B*, *PLA2G7*) while CD14M_2 enriched for vitamin transport, water-soluble vitamin metabolic process (*SLC2A3*, *FOLR3*, *VNN2*), and organic anion transport (*ABCC6*, *SLC2A3*).
- CD14M_2 was discriminated from CD14M_1 by expression of genes involved in detoxification processes, lipid hydrolysis, and citrullination for NETosis in neutrophils (*MGST1*, *CES1*, *PADI4*)[56–58], clearance of apoptotic cells (*STAB1*)[59], and phospholipid metabolism (*PLA2G7*), among others.

- CD14M_3 showed gene expression significant for antigen presentation of peptide antigen via MHC class II (*HLA-DMB*, *HLA-DOA*, *HLA-DQA2*, *HLA-DQA1*) and positive regulation of leukocyte chemotaxis (*CSF1R*, *CALR*). The cluster defining genes included *HLA-DQA1*, *CSF1R*, *MARCKS*, *CALR*, and *LIPA*[21,60].
- CD14M_4 cluster enriched for defense responses against symbiont/virus or negative regulation of viral processes, IL-27 signaling, antiviral innate immune response, positive regulation of type-I IFN production, IFN-$\alpha/\beta$ signaling, ISG15 antiviral mechanism, and IFN-$\gamma$ signaling (*STAT1*, STAT2, *MX1*, *MX2, IFI6*, *EIF2AK2*, *ISG15, IFIT1*, *IFI44L*, *IFIT3*, *IFIT2*, *OASL*, *OAS2*, *OAS3*, *IRF7*, *RSAD2*, *HERC, OASL*)[61,62].
- CD14M_5 was defined by inflammatory response (*CXCL8*, *CCL3L1*, *TNFAIP6*, *IRAK2*, *RIPK2*, *IL1B*, *CCL4*, *CCL3*, *NLRP3*, *CXCL2*), cellular response to IL-1 (*EGR1*, *CXCL8*, *CCL3L1*, *IRAK2*, *IL1B*, *CCL4*, CCL3), cellular response to LPS, granulocyte migration, chemokine activity (*CXCL8, CCL3L1, CCL4*, *CCL3, CXCL2*), and NF$\kappa$B signaling (*CXCL8*, *CCL4L2*, *IL1B*, *CCL4, TNFAIP3*, *PTGS2*, *CXCL2*, *ICAM1*).
- CD14M_6 cluster was enriched in activation of GTPase activity (*TBC1D5*, *TBC1D22A*, *ARHGAP24*), positive regulation of protein tyrosine kinase activity (*FBXW7*, *NEDD9*), regulation of glycolytic process (*PKRAG2*), positive regulation of myeloid leukocyte differentiation and biosynthetic processes (*NEDD9*, *RUNX1*, *ZBTB20*). This is suggestive of cytoskeletal dynamics, cell differentiation and proliferation, and immune activation[63].
- CD14M_7 was defined by chemokine-mediated signaling (*CCL5*, *PPBP*, *PF4V1, PF4*), myeloid cell and megakaryocyte development (*GP9*, *MPIG6B, GP1BB*), and negative regulation of $Ca^{2+}$ sequestering (*CLU*, *GP1BB*).
- CD14M_8 enriched for a multitude of lncRNAs (Figure S8B), including *VCAN-AS1, CPB2-AS1*, and *DPYD-AS1*, suggesting that this cluster may have regulatory roles in gene expression and additional processes.
- CD16M_1 cluster was characterized by positive regulation of actin filament bundle assembly, cellular response to fluid shear stress, cell-cell junction maintenance (*EVL*, *RHOC*, *MTSS1*, *KLF2*, *CSF1R)*, negative regulation of cell population proliferation, and apoptosis (*CDKN1C*, *IFITM1*, *HMOX1*, *MTSS1*, *TCF7L2*).
- CD16M_2 showed enrichment of similar terms, including positive regulation of intracellular transduction (*CX3CR1*, *TCF7L2*, *PTP4A3*, *HMOX1*, *LTB*, *RHOC*).
- cDC represented similar MHC class-II protein class assembly and MHC-II restricted antigen presentation terms (*HLA-DOA*, *HLA-DQA2*, *HLA-DQA1, FCER1A)*, progenitor cell differentiation (*FLT3*, *PLD4*), and antigen (including lipid) presentation (*CD1C*, *CD1E*), as described above[21].
- pDC showed the same markers and functions as above.
- The "Mono" cluster showed antigen receptor-mediated signaling and T-cell activation (*ITK*, *TRBC2*, *CD247*, *CD3E*, *CD3D*, *SKAP1*), which could be a result of the myeloid cell cluster being in contact with adaptive immune cells or a clustering artifact.

B-lymphoid lineage cells: L1 annotations for B-lymphoid cells in the blood had the same markers as in CSF, including *MS4A1*, *BANK1*, *CD79A*, *RALGPS2*, and *BLK*[30–32]. The following are the subclusters and pathway analyses performed (Figure S9A-D):

- Cluster b1 enriched in cellular response to cytokines (*TCL1A*, *IL4R*, *CCR7*, *YBX3*) and negative regulation of cytokine production (*APLP2*, *CD200*). This cluster may represent $IgM^{low}$ transitional B-cells[33].
- Cluster b2 was defined by B-cell receptor signaling pathway (*IGHG1*, *IGHG2*, *IGHA1*) and regulation of B cell proliferation and B-cell activation (*TNFRSF13B*, *GPR183*, *CD27*).

- Cluster b3 showed expression of genes involvement in negative regulation of inflammatory response (*FGR*, *HCK, GRN*, *TNFRSF1B*) and positive regulation of T-helper 1 type immune response (*SLC11A1*, *TBX21*).
- Cluster b4 is defined by positive regulation of cell-substrate adhesion (*CDK6*, *JUP*, *LIMS2*), regulation of B-cell proliferation, and B-cell-receptor signaling pathway (*MZB1*, *CD38*, *IGLC1*).
- Cluster b5 enriched in regulation of $Na^+$ ion transmembrane transporter activity (*CAMK2D*, *PRKCE*, *UTRN*), $F_c\gamma$- and $F_c$-receptor signaling pathway in phagocytosis (*VAV3*, *PRKCE*), and actin cytoskeleton reorganization (*SIPA1L1*, *AUTS2*) and phosphorylation (*CAMK2D*, *PRKCE*, *SIK3, CDK14*).
- Cluster b6 represented terms including $\alpha\beta$ T-cell activation (*NKG7*, *CD247*, *CD3E, CD3D*), granulocyte chemotaxis (*ANXA1*, *CCL5*), and genes like *IL32*, *TYROBP*, and *GNLY*. This peripheral B-cell cluster was transcriptionally most similar to the b2 cluster in the CSF (Figure S9D), therefore, more likely to resemble plasmablast (see above for CSF b2 cluster). Other cluster markers are shown in Figure S9B.

Note that for both the CSF cells and PBMC, we projected the L2 annotations onto L1 space, supporting the sub-clustering scheme (Figure S5C-D, G-H).

**Combined object and immune cells across CSF and blood compartments**

Integrating the CSF and PBMC samples, we created a combined object (Figure S8B), with level 2 (L2) cell annotations mapped onto it (Figure S8C). In this combined object, the myeloid cells from CSF clustered separately from the myeloid cells of PBMC origin (Figure S8C, D). This is consistent with the diversity of CSF macrophages and the unique niche they occupy[64]. Most of the overlap between CSF- and PBMC-derived myeloid cells occurred among dendritic cells across the two compartments.

The adaptive immune cells from both CSF and blood, on the other hand, clustered together. However, in the non-perturbed samples (untreated and radiologically inactive PRL-positive vs. PRL-negative) and across the compartments, CD8-TEM and CD4-TCM cells upregulated genes associated with integrin signaling and migration (*CXCR3*, *CXCR4*, *CXCR6*, *CCL5*, *ITGA4*) and cytotoxicity (*GZMK*, *GZMA*) in CSF relative to blood (Figure S8E, F). This aligns with a recent report describing upregulation of genes associated with migration in $CD8^+$ and $CD4^+$ T cells in the CSF relative to PBMC[65].

The NK cells in the CSF were enriched in pathways pertinent to eukaryotic translation and lymphotoxin receptor signaling relative to blood (Figure S8F). Notably, GZMK was one of the most enriched genes in CD8-TEM cells from CSF relative to blood (Figure S8E). Since we find clonal expansion of CD8-TEM cells in PRL-positive vs. PRL-negative cases in both blood and CSF, this suggests that clonally expanded CD8-TEM cells may acquire an even more cytotoxic signature upon trafficking from the blood to the CSF, consistent with their relevance to mediation of chronic neuroinflammation in PRL-positive cases.

**IL15 regulates immune responses in chronic active MS**

To determine how the immune environment might be modulating the transcriptomic changes across cells from PRL-positive versus PRL-negative cases in both blood and CSF, we performed NicheNet analyses, which interrogate predicted ligand-receptor interactions. We found *IL15* and *TNFSF12* (TWEAK) to be the predicted regulatory ligands of the target genes for myeloid (Figure S9A) and CD4 T-cells (Figure S9B) in blood. This is consistent with prior reports implicating IL15-induced pro-inflammatory responses in MS and experimental autoimmune encephalomyelitis (EAE)[66,67]. Previously, TWEAK+ cells were found to be frequent at the edges of chronic active lesion edges and in subpial cortical lesions, suggesting a potential role of TNFSF12-mediated signaling in such lesions[68]. We also found *TNFSF10* (TRAIL) to be modulating the transcriptional changes in blood myeloid cells in the PRL-positive state.

Interestingly, *CD40*, expressed by B cells, was also predicted to be a critical regulator of the transcriptional changes in CD4 T-cells in blood (Figure S9B), whereas *CD40LG* was a predicted regulatory ligand for myeloid cells. CD40-CD40LG costimulatory signaling is critical in regulating the initiation of adaptive and innate immune cellular processes, with recent approaches involving the therapeutic targeting of *CD40LG* with a monoclonal antibody in MS[69,70]. For NK cells, *LTB*, *TYROBP*, *CD40LG,* and *APP* were the among the top regulatory ligands driving the differentially expressed genes across PRL-positive versus PRL-negative cells (Figure S9C). In the case of *LTB*, its interaction with *LTβR* on NK-cell induces innate immune responses[71]. For CD8 T-cells, *ITGAL*, *GRN*, and *TIMP1* were the main predicted regulatory ligands, with the latter regulating *MYC*, a transcription factor involved in CD8 T-cell fate disposition toward an effector phenotype (Figure S9D)[72]. For B-cells, *CD2* was the molecule predicted to regulate the expression of TCF4, which is critical for germinal center B-cell development (Figure S9E)[73].

In terms of the prioritized differential ligand-receptor interactions in blood, we found HLA-I molecules on NK cells to be interacting with various receptors, including *LILRB2*, *LILRB1*, and *SIGLEC1*, on myeloid cells; *KLRC1* and *KLRD1* on NK cells; and *CD8A* and *KLRK1* on CD8 T-cells (Figure S9F). These ligand-receptor interactions have an immunoregulatory potential, with interactions involving *CD8A*, *KLRC1*, and *KLRK1* receptors promoting T-cell activation and cytotoxicity, and *LILRB2* and *LILRB1* acting as inhibitory receptors for MHC-I class molecules, thereby dampening myeloid-mediated immune responses[74]. Similarly, galectins (LGALS1 and LGALS9) on myeloid cells interacted with PTPRC on NK cells, and they have been shown to modulate immune responses[75].

In the CSF, the top predicted molecules regulating the target genes in PRL-positive cases included *TNF* from CD4 T-cells and *TGFB1* from NK cells (Figure S9G). In the $CD8^+$ T-cells, *TNF* from CD4 T-cells emerged as the predicted molecule driving the differentially expressed genes across PRL-positive vs. PRL-negative cells (Figure S10A). The central target gene modulated by the predicted regulators in both myeloid and $CD8^+$ T-cells of the CSF was *IRF1*, which is also strongly induced by IFN-$\gamma$-mediated signaling[76]. *B2M* and *TYROBP* from $CD8^+$ T-cells and cDC, respectively, were the main predicted ligands for NK cells, while *TGFB1* and *HMGB1* were the predicted ligands for CD4 T-cells (Figure S10B, C). The prioritized ligand-receptor interactions in CSF were significant for myeloid *SPP1* as the ligand for *ITGA4* on myeloid, and *ITGB5* and *CD44* receptors on CD4 T-cells, respectively (Figure S10E). Prior studies have suggested the role of *SPP1* (osteopontin) in regulating $T_H1$ responses in an EAE model, with its ablation resulting in less IFN-$\gamma$ production by myelin-reactive T cells[77]. We also identified APOE as a prioritized ligand interacting with TREM2 receptors on myeloid cells in the CSF of PRL-positive cases. APOE-TREM2 interaction has been implicated in switching of the microglia to a neurodegenerative phenotype[78]. Concordantly, our proteomic data showed abundance of TREM2 in the CSF of patients with a high burden of PRL (PRL $\geq$ 4) vs. low burden (PRL 1-3;

p = 0.06) and no-PRL (p = 0.1). In addition, we also found enrichment of IL15 in the CSF of cases with high PRL burden relative to low PRL (p = 0.004) and no PRL (p = 0.0008) (Figure S10F).

**Fig. S1**

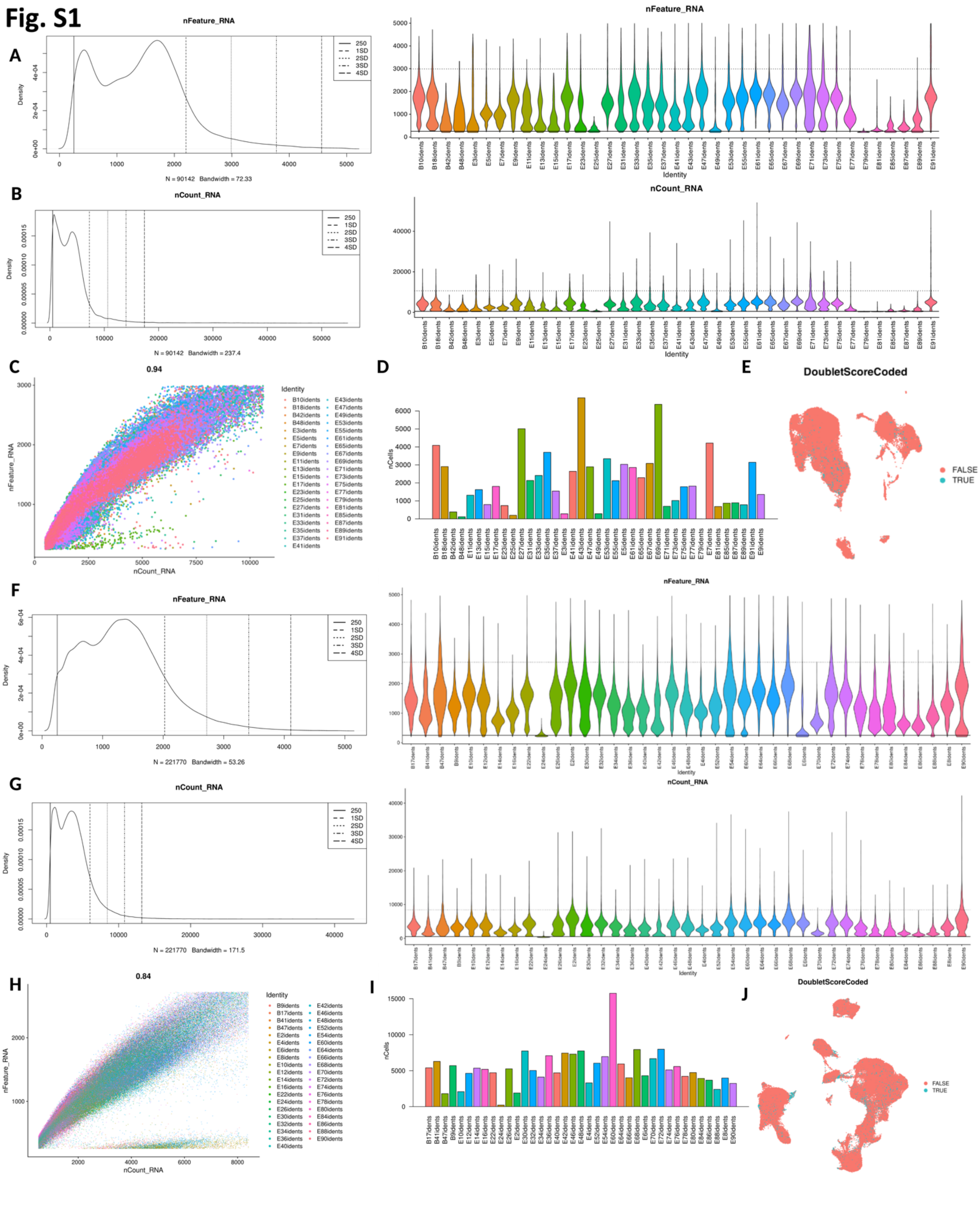

**Figure S1: Quality control (QC) metrics for CSF and PBMC objects**

a) Pre-filtering density curve for nFeature_RNA (left panel) — unique features (genes) — identified and violin plot (right panel) showing nFeature_RNA per CSF sample.
b) Pre-filtering density curve for nCount_RNA (left panel) — total number of genes detected — and violin plot (right panel) demonstrating counts per CSF sample.
c) Post-filtering scatter plot showing high correlation between nCount_RNA and nFeature_RNA for the CSF object.
d) Post-filtering cell counts per CSF sample. E79 was excluded from further analyses.
e) UMAP illustrating doublets that were excluded from analyses.
f) Pre-filtering density curve for nFeature_RNA (left panel) and violin plot (right panel) showing nFeature_RNA per PBMC sample.
g) Pre-filtering density curve for nCount_RNA (left panel) and violin plot (right panel) demonstrating counts per PBMC sample.
h) Post-filtering scatter plot between nCount_RNA and nFeature_RNA for the PBMC object.
i) Total cell counts per PBMC sample following filtering.
j) Doublets identified shown in the UMAP for PBMC object. These were removed from further analyses.

#### Fig. S2

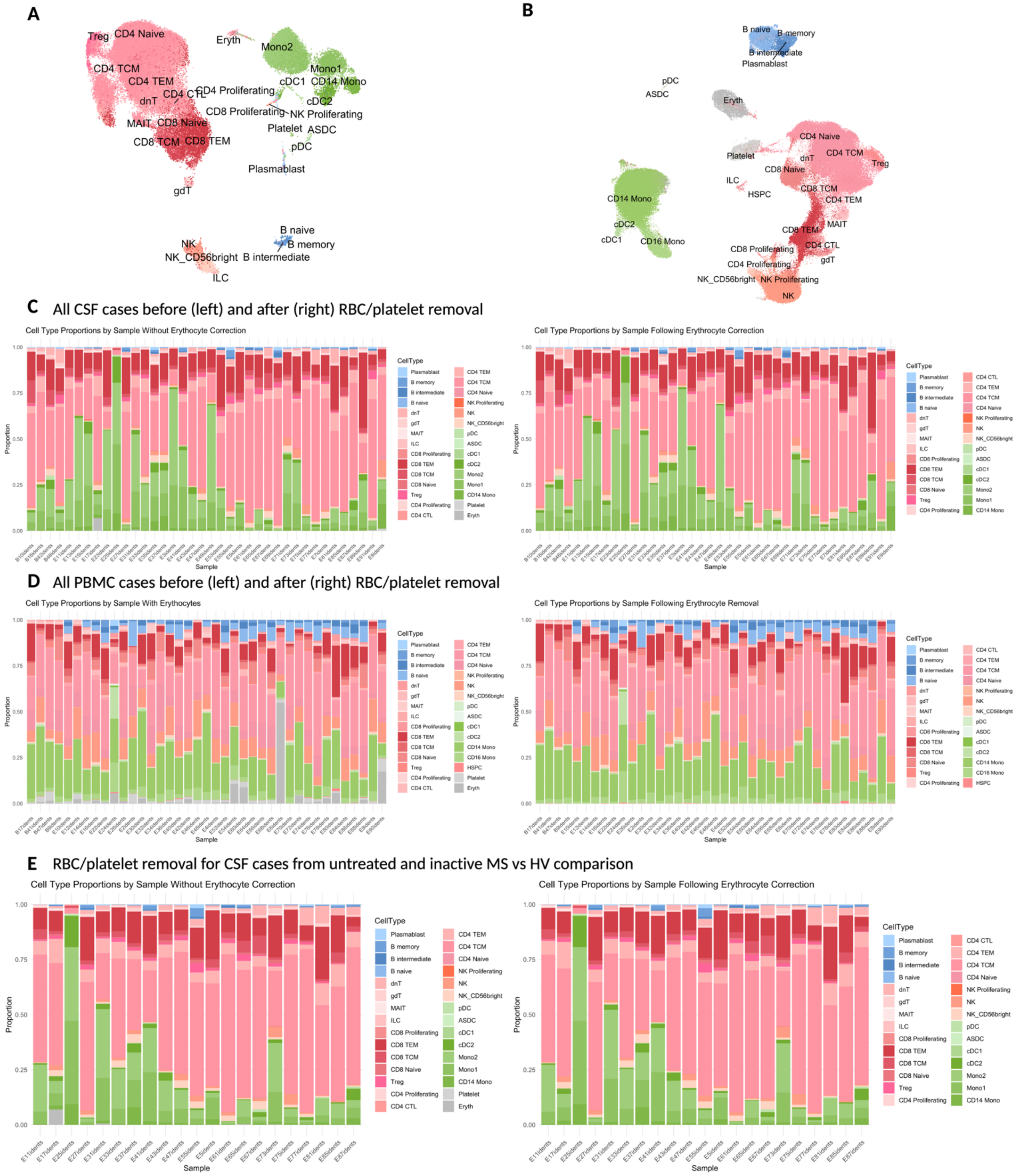
A
Treg
CD4 Naive
CD4 TCM
CD4 TEM
CD4 CTL
CD4 Proliferating
dnT
CD8 Proliferating
MAIT
CD8 Naive
CD8 TCM
CD8 TEM
gdT
Eryth
Mono2
Mono1
cDC1
CD14 Mono
cDC2
NK Proliferating
Platelet
ASDC
pDC
Plasmablast
NK
NK_CD56bright
B naive
B memory
B intermediate
ILC
B
B naive
B memory
B intermediate
Plasmablast
pDC
ASDC
Eryth
CD4 Naive
Platelet
dnT
CD4 TCM
Treg
CD8 Naive
ILC
HSPC
CD8 TCM
CD4 TEM
CD14 Mono
MAIT
CD8 TEM
cDC2
cDC1
CD16 Mono
CD4 CTL
CD8 Proliferating
gdT
CD4 Proliferating
NK_CD56bright
NK Proliferating
NK
C All CSF cases before (left) and after (right) RBC/platelet removal
Cell Type Proportions by Sample Without Erythocyte Correction
Cell Type Proportions by Sample Following Erythrocyte Correction
Proportion
Sample
CellType
D All PBMC cases before (left) and after (right) RBC/platelet removal
Cell Type Proportions by Sample With Erythocytes
Cell Type Proportions by Sample Following Erythrocyte Removal
E RBC/platelet removal for CSF cases from untreated and inactive MS vs HV comparison
Cell Type Proportions by Sample Without Erythocyte Correction
Cell Type Proportions by Sample Following Erythrocyte Correction

### Fig. S2 continued

**F** RBC/platelet removal for PBMC cases from untreated and inactive MS vs HV comparison

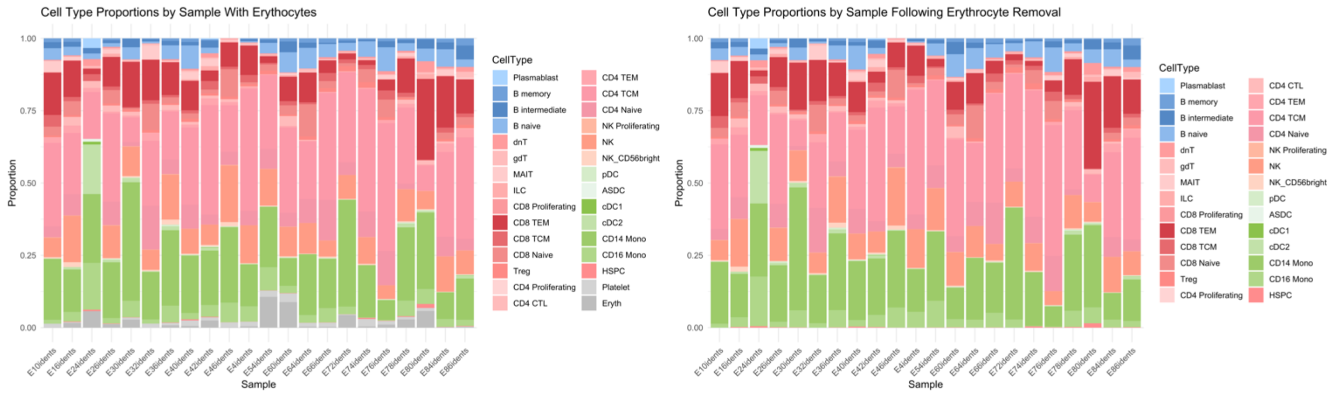


**G** RBC/platelet removal for CSF cases from untreated and inactive PRL-positive vs PRL-negative comparison

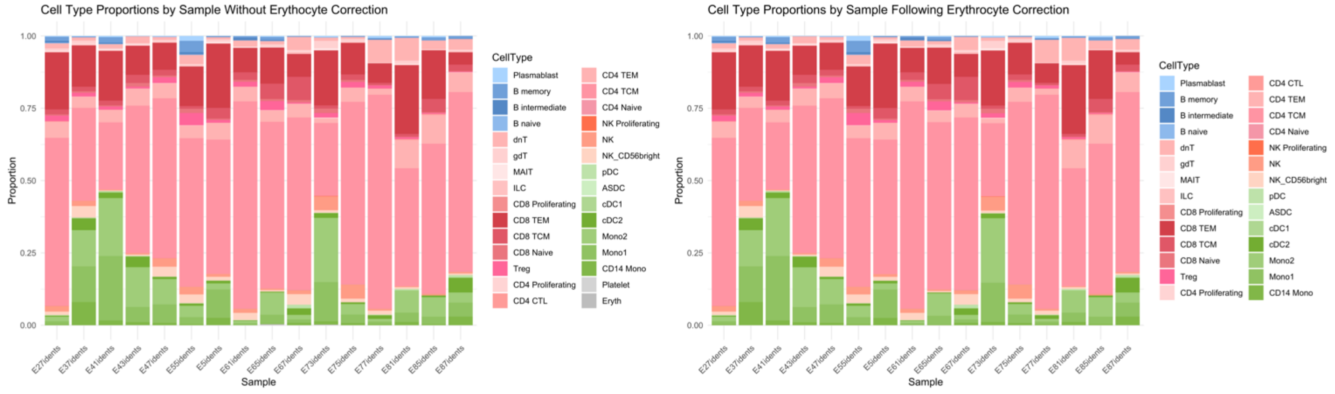


**H** RBC/platelet removal for PBMC cases from untreated and inactive PRL-positive vs PRL-negative comparison

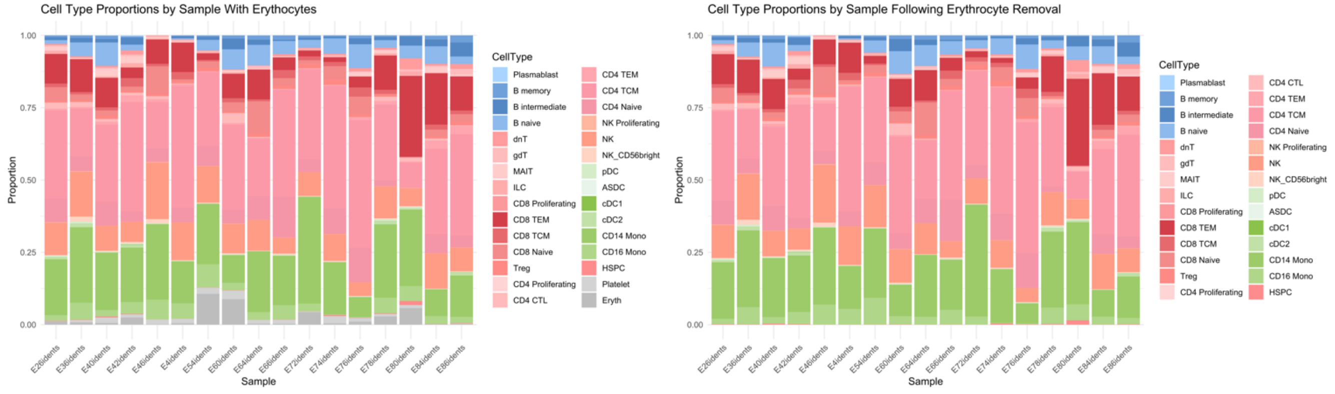

**Figure S2: Erythrocyte and platelet removal (see Methods) does not impact cell proportions in the CSF and PBMC**

a) Overall CSF L1 UMAP representing cell populations, including erythrocytes and platelets.
b) Overall PBMC L1 UMAP including erythrocytes and platelets.
c) Stacked bar graph summarizing CSF L1 cell proportions prior to (left panel) and following (right panel) erythrocyte and platelet removal.
d) Stacked bar graph summarizing PBMC L1 cell proportions prior to (left panel) and following (right panel) erythrocyte and platelet removal.
e) CSF L1 cell proportions for the untreated and inactive MS vs. HV samples subsampled from the main CSF object, showing no to minimal change in proportions before and following erythrocyte and platelet removal.
f) PBMC L1 cell proportions for the untreated and inactive MS vs. HV samples before and following erythrocyte and platelet removal, showing no to minimal change in proportions before and following erythrocyte and platelet corrections.
g) Stacked bar graph showing CSF L1 cell proportions without and with erythrocyte and platelet removal for the PRL-positive vs. PRL-negative samples.
h) Stacked bar graph showing PBMC L1 cell proportions without and with erythrocyte and platelet removal for the PRL-positive vs. PRL-negative samples.

### Fig. S3

CSF

**A**

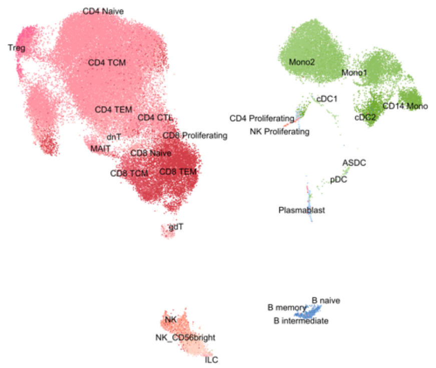


**B**

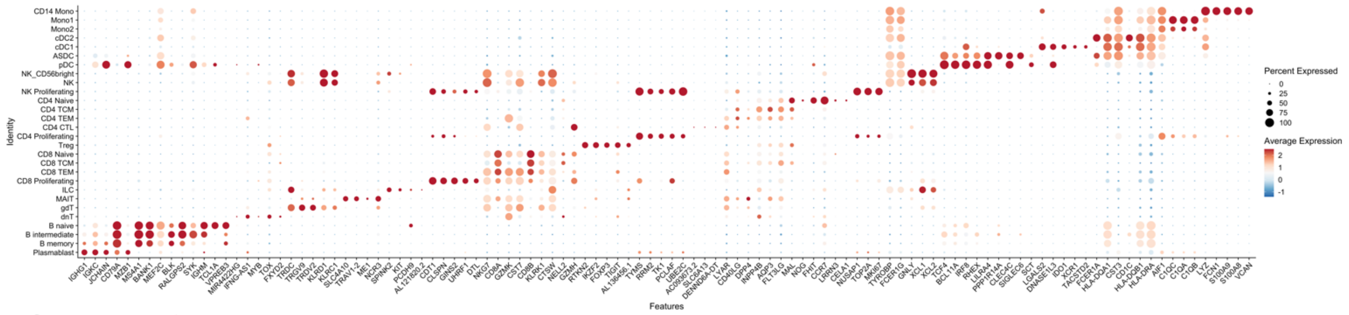


**C**

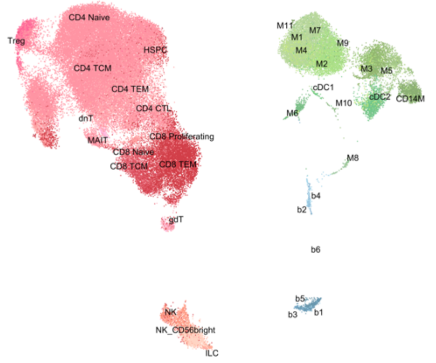


**D**

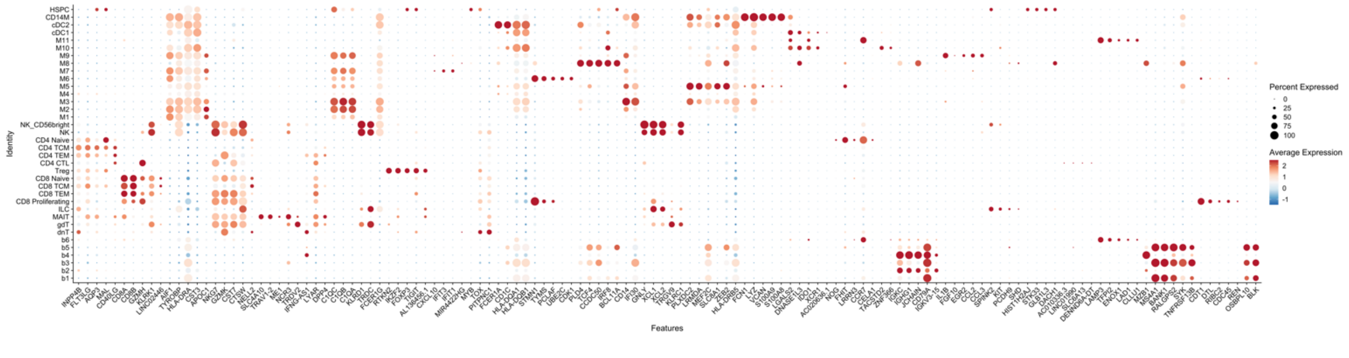

#### Fig. S3 continued

PBMC

E

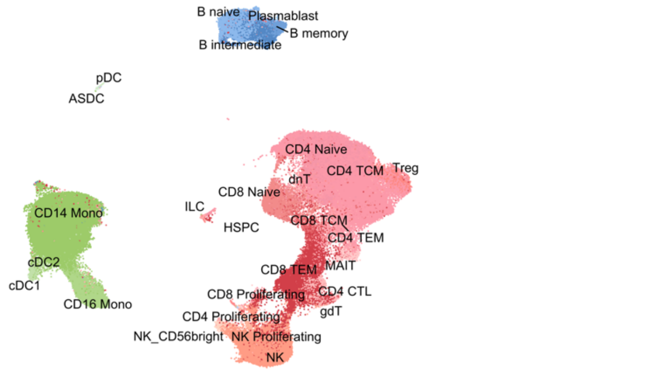


F

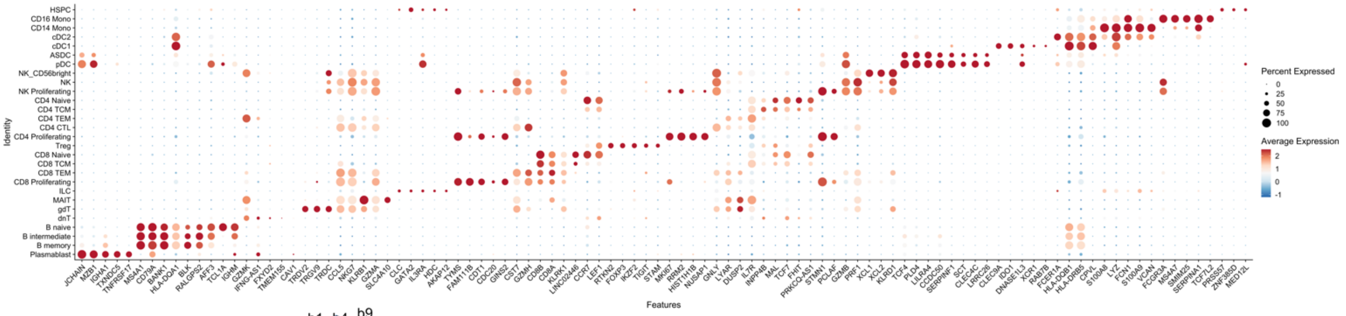


G

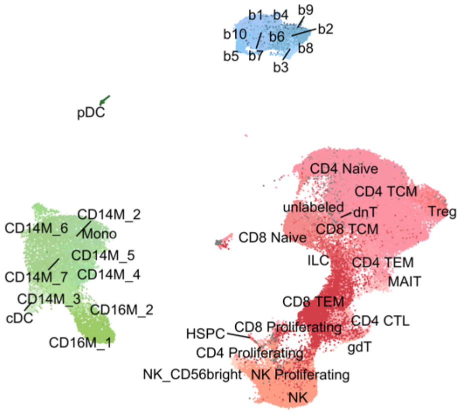


H

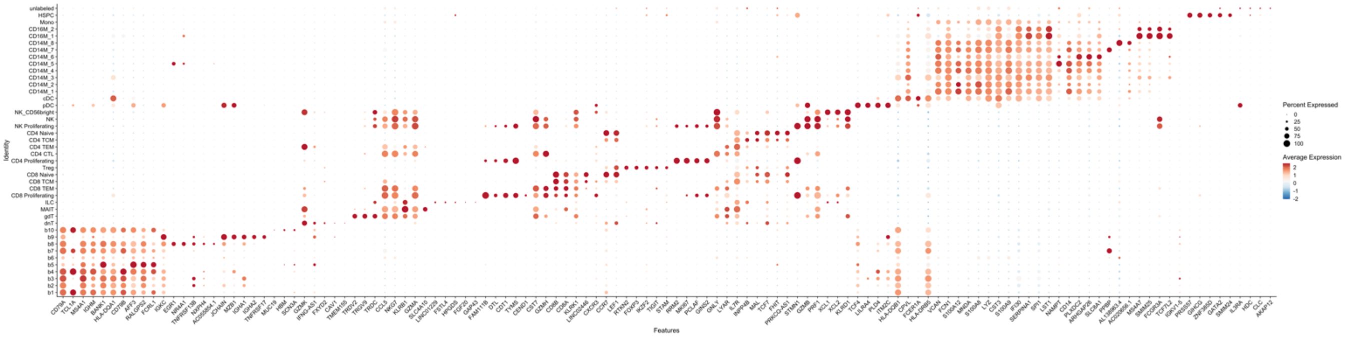

**Figure S3: Top in-depth gene markers used for inferring cell populations in the CSF and PBMC**

a) CSF L1 UMAP with the cell populations shown.
b) Dotplot representing top gene markers for the various L1 cell populations in the CSF.
c) CSF UMAP with L2 cell annotations mapped onto L1 clusters.
d) Dotplot summarizing the top gene markers for L2 annotated cell populations in the CSF.
e) Scatterplot showing the L1 annotations for the PBMC.
f) Dotplot representing the top gene markers for L1 cell annotated populations of the PBMC.
g) PBMC UMAP with L2 cell annotations mapped onto L1 clusters.
h) Dotplot summarizing the top gene markers for L2 annotated cell populations of the PBMC.

**Fig. S4**

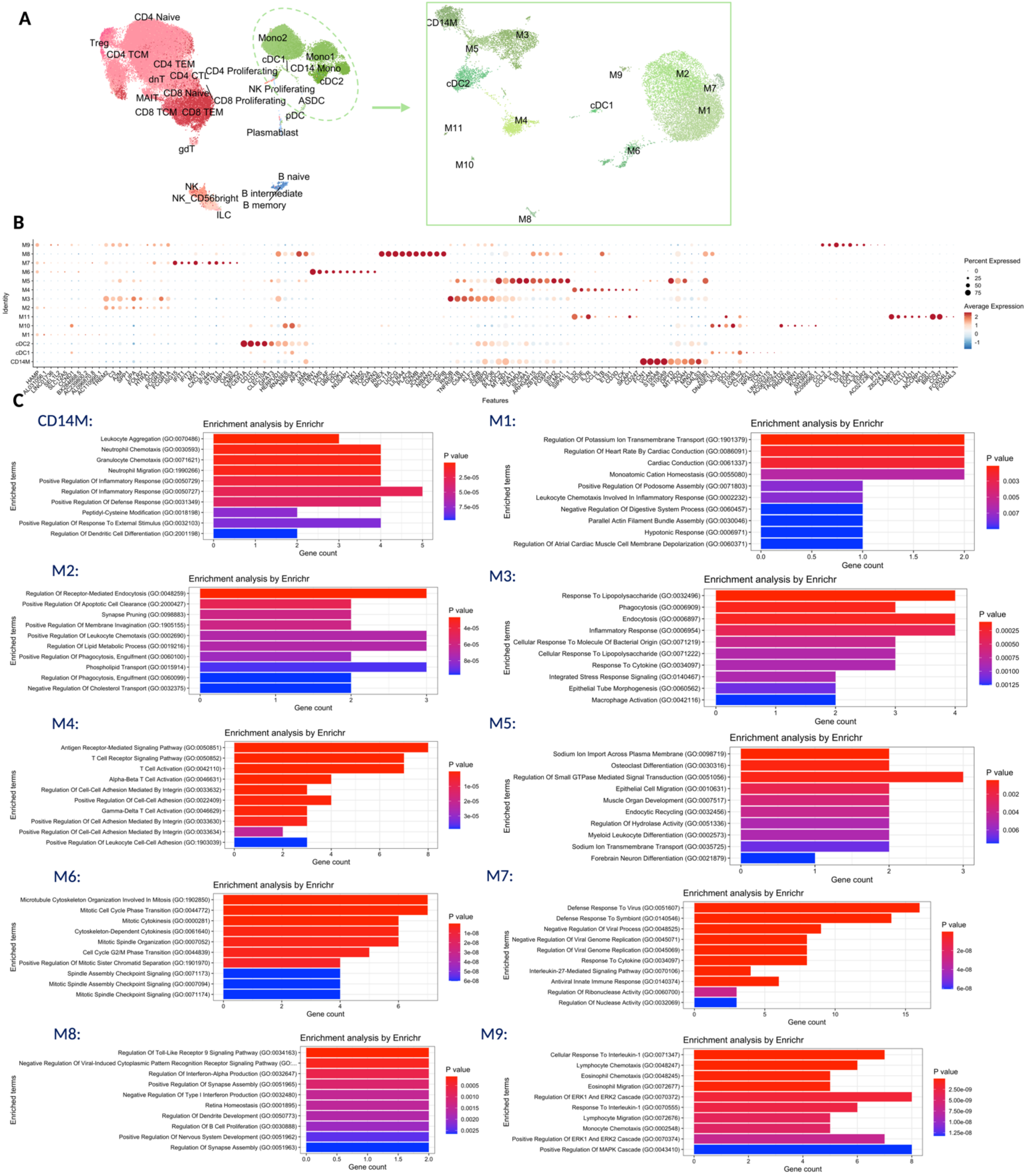
A
CD4 Naive
Treg
CD4 TCM
CD4 TEM
dnT
CD4 CTL
CD4 Proliferating
MAIT
CD8 Naive
CD8 Proliferating
CD8 TCM
CD8 TEM
gdT
Mono2
cDC1
Mono1
CD14 Mono
cDC2
NK Proliferating
ASDC
pDC
Plasmablast
NK
NK_CD56bright
ILC
B naive
B intermediate
B memory
CD14M
M3
M5
cDC2
M9
M2
M7
M1
cDC1
M4
M11
M6
M10
M8
B
Identity
Features
Percent Expressed
Average Expression
C
CD14M:
Enrichment analysis by Enrichr
Leukocyte Aggregation (GO:0070486)
Neutrophil Chemotaxis (GO:0030593)
Granulocyte Chemotaxis (GO:0071621)
Neutrophil Migration (GO:1990266)
Positive Regulation Of Inflammatory Response (GO:0050729)
Regulation Of Inflammatory Response (GO:0050727)
Positive Regulation Of Defense Response (GO:0031349)
Peptidyl-Cysteine Modification (GO:0018198)
Positive Regulation Of Response To External Stimulus (GO:0032103)
Regulation Of Dendritic Cell Differentiation (GO:2001198)
Enriched terms
Gene count
P value
M1:
Enrichment analysis by Enrichr
Regulation Of Potassium Ion Transmembrane Transport (GO:1901379)
Regulation Of Heart Rate By Cardiac Conduction (GO:0086091)
Cardiac Conduction (GO:0061337)
Monoatomic Cation Homeostasis (GO:0055080)
Positive Regulation Of Podosome Assembly (GO:0071803)
Leukocyte Chemotaxis Involved In Inflammatory Response (GO:0002232)
Negative Regulation Of Digestive System Process (GO:0060457)
Parallel Actin Filament Bundle Assembly (GO:0030046)
Hypotonic Response (GO:0006971)
Regulation Of Atrial Cardiac Muscle Cell Membrane Depolarization (GO:0060371)
Enriched terms
Gene count
P value
M2:
Enrichment analysis by Enrichr
Regulation Of Receptor-Mediated Endocytosis (GO:0048259)
Positive Regulation Of Apoptotic Cell Clearance (GO:2000427)
Synapse Pruning (GO:0098883)
Positive Regulation Of Membrane Invagination (GO:1905155)
Positive Regulation Of Leukocyte Chemotaxis (GO:0002690)
Regulation Of Lipid Metabolic Process (GO:0019216)
Positive Regulation Of Phagocytosis, Engulfment (GO:0060100)
Phospholipid Transport (GO:0015914)
Regulation Of Phagocytosis, Engulfment (GO:0060099)
Negative Regulation Of Cholesterol Transport (GO:0032375)
Enriched terms
Gene count
P value
M3:
Enrichment analysis by Enrichr
Response To Lipopolysaccharide (GO:0032496)
Phagocytosis (GO:0006909)
Endocytosis (GO:0006897)
Inflammatory Response (GO:0006954)
Cellular Response To Molecule Of Bacterial Origin (GO:0071219)
Cellular Response To Lipopolysaccharide (GO:0071222)
Response To Cytokine (GO:0034097)
Integrated Stress Response Signaling (GO:0140467)
Epithelial Tube Morphogenesis (GO:0060562)
Macrophage Activation (GO:0042116)
Enriched terms
Gene count
P value
M4:
Enrichment analysis by Enrichr
Antigen Receptor-Mediated Signaling Pathway (GO:0050851)
T Cell Receptor Signaling Pathway (GO:0050852)
T Cell Activation (GO:0042110)
Alpha-Beta T Cell Activation (GO:0046631)
Regulation Of Cell-Cell Adhesion Mediated By Integrin (GO:0033632)
Positive Regulation Of Cell-Cell Adhesion (GO:0022409)
Gamma-Delta T Cell Activation (GO:0046629)
Positive Regulation Of Cell Adhesion Mediated By Integrin (GO:0033630)
Positive Regulation Of Cell-Cell Adhesion Mediated By Integrin (GO:0033634)
Positive Regulation Of Leukocyte Cell-Cell Adhesion (GO:1903039)
Enriched terms
Gene count
P value
M5:
Enrichment analysis by Enrichr
Sodium Ion Import Across Plasma Membrane (GO:0098719)
Osteoclast Differentiation (GO:0030316)
Regulation Of Small GTPase Mediated Signal Transduction (GO:0051056)
Epithelial Cell Migration (GO:0010631)
Muscle Organ Development (GO:0007517)
Endocytic Recycling (GO:0032456)
Regulation Of Hydrolase Activity (GO:0051336)
Myeloid Leukocyte Differentiation (GO:0002573)
Sodium Ion Transmembrane Transport (GO:0035725)
Forebrain Neuron Differentiation (GO:0021879)
Enriched terms
Gene count
P value
M6:
Enrichment analysis by Enrichr
Microtubule Cytoskeleton Organization Involved In Mitosis (GO:1902850)
Mitotic Cell Cycle Phase Transition (GO:0044772)
Mitotic Cytokinesis (GO:0000281)
Cytoskeleton-Dependent Cytokinesis (GO:0061640)
Mitotic Spindle Organization (GO:0007052)
Cell Cycle G2/M Phase Transition (GO:0044839)
Positive Regulation Of Mitotic Sister Chromatid Separation (GO:1901970)
Spindle Assembly Checkpoint Signaling (GO:0071173)
Mitotic Spindle Assembly Checkpoint Signaling (GO:0007094)
Mitotic Spindle Checkpoint Signaling (GO:0071174)
Enriched terms
Gene count
P value
M7:
Enrichment analysis by Enrichr
Defense Response To Virus (GO:0051607)
Defense Response To Symbiont (GO:0140546)
Negative Regulation Of Viral Process (GO:0048525)
Negative Regulation Of Viral Genome Replication (GO:0045071)
Regulation Of Viral Genome Replication (GO:0045069)
Response To Cytokine (GO:0034097)
Interleukin-27-Mediated Signaling Pathway (GO:0070106)
Antiviral Innate Immune Response (GO:0140374)
Regulation Of Ribonuclease Activity (GO:0060700)
Regulation Of Nuclease Activity (GO:0032069)
Enriched terms
Gene count
P value
M8:
Enrichment analysis by Enrichr
Regulation Of Toll-Like Receptor 9 Signaling Pathway (GO:0034163)
Negative Regulation Of Viral-Induced Cytoplasmic Pattern Recognition Receptor Signaling Pathway (GO:...
Regulation Of Interferon-Alpha Production (GO:0032647)
Positive Regulation Of Synapse Assembly (GO:0051965)
Negative Regulation Of Type I Interferon Production (GO:0032480)
Retina Homeostasis (GO:0001895)
Regulation Of Dendrite Development (GO:0050773)
Regulation Of B Cell Proliferation (GO:0030888)
Positive Regulation Of Nervous System Development (GO:0051962)
Regulation Of Synapse Assembly (GO:0051963)
Enriched terms
Gene count
P value
M9:
Enrichment analysis by Enrichr
Cellular Response To Interleukin-1 (GO:0071347)
Lymphocyte Chemotaxis (GO:0048247)
Eosinophil Chemotaxis (GO:0048245)
Eosinophil Migration (GO:0072677)
Regulation Of ERK1 And ERK2 Cascade (GO:0070372)
Response To Interleukin-1 (GO:0070555)
Lymphocyte Migration (GO:0072676)
Monocyte Chemotaxis (GO:0002548)
Positive Regulation Of ERK1 And ERK2 Cascade (GO:0070374)
Positive Regulation Of MAPK Cascade (GO:0043410)
Enriched terms
Gene count
P value

**Figure S4: CSF myeloid subclusters demonstrate heterogeneity in function**

a) CSF L1 UMAP showing the different populations, with the myeloid cells (encircled by a green dashed line) subclustered into L2 populations as shown in the right panel.
b) Dotplot representing the gene markers of the myeloid subclusters. Average gene expression and the percentage of cells expressing the genes are shown.
c) GO analyses of the top gene markers for the subclusters (see Methods to find how the top markers were found using FindMarker() function). GO terms, including majorly Biological Processes (BP) were used for annotating the marker gene lists. When required, additional subontologies including Molecular Function (MF), Cellular Component (CC), KEGG, Wiki, BioCarta, and Panther were used. CD14M subcluster was associated with leukocyte aggregation, neutrophil and granulocyte chemotaxis and positive regulation of inflammatory response. M1 enriched in regulation of potassium ion ($K^+$) transport, while M2 enriched in receptor-mediated endocytosis, synapse pruning, positive regulation of leukocyte chemotaxis and regulation of lipid metabolic process. M3 subcluster was associated with response to lipopolysaccharide (LPS), phagocytosis, endocytosis and inflammatory response. Gene markers identifying M4 subcluster enriched in antigen receptor-mediated signaling, while M5 showed association with sodium ($Na^+$) import, and regulation of small GTPase-mediated signaling. M6 was primarily associated with cell-cycle phases and mitotic spindle organization, while M7 was enriched for defense response to virus and regulation of viral genome replication. M8 enriched in Toll-like receptor (TLR) signaling, and M9 demonstrated enrichment for response to IL-1, ERK1/ERK2 cascade regulation, and chemotaxis of lymphocytes.

### Fig. S5

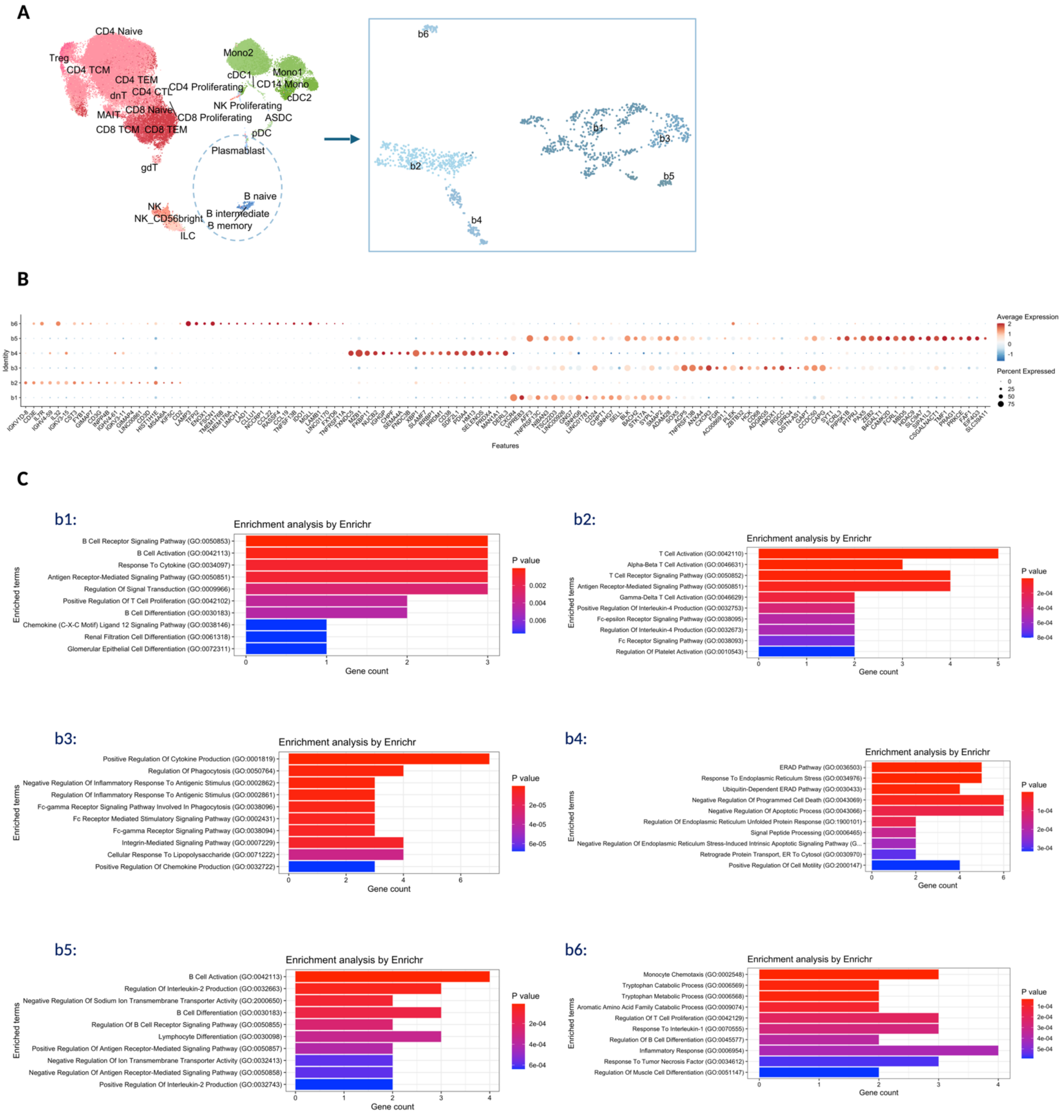
A
CD4 Naive
Treg
CD4 TCM
CD4 TEM
dnT
CD4 CTL
CD4 Proliferating
CD8 Naive
MAIT
CD8 TCM
CD8 TEM
CD8 Proliferating
NK Proliferating
gdT
Mono2
cDC1
Mono1
CD14 Mono
cDC2
ASDC
pDC
Plasmablast
B naive
B intermediate
B memory
NK
NK_CD56bright
ILC
b1
b2
b3
b4
b5
b6
B
Identity
Features
Average Expression
Percent Expressed
C
b1:
Enrichment analysis by Enrichr
B Cell Receptor Signaling Pathway (GO:0050853)
B Cell Activation (GO:0042113)
Response To Cytokine (GO:0034097)
Antigen Receptor-Mediated Signaling Pathway (GO:0050851)
Regulation Of Signal Transduction (GO:0009966)
Positive Regulation Of T Cell Proliferation (GO:0042102)
B Cell Differentiation (GO:0030183)
Chemokine (C-X-C Motif) Ligand 12 Signaling Pathway (GO:0038146)
Renal Filtration Cell Differentiation (GO:0061318)
Glomerular Epithelial Cell Differentiation (GO:0072311)
Enriched terms
Gene count
P value
b2:
Enrichment analysis by Enrichr
T Cell Activation (GO:0042110)
Alpha-Beta T Cell Activation (GO:0046631)
T Cell Receptor Signaling Pathway (GO:0050852)
Antigen Receptor-Mediated Signaling Pathway (GO:0050851)
Gamma-Delta T Cell Activation (GO:0046629)
Positive Regulation Of Interleukin-4 Production (GO:0032753)
Fc-epsilon Receptor Signaling Pathway (GO:0038095)
Regulation Of Interleukin-4 Production (GO:0032673)
Fc Receptor Signaling Pathway (GO:0038093)
Regulation Of Platelet Activation (GO:0010543)
Enriched terms
Gene count
P value
b3:
Enrichment analysis by Enrichr
Positive Regulation Of Cytokine Production (GO:0001819)
Regulation Of Phagocytosis (GO:0050764)
Negative Regulation Of Inflammatory Response To Antigenic Stimulus (GO:0002862)
Regulation Of Inflammatory Response To Antigenic Stimulus (GO:0002861)
Fc-gamma Receptor Signaling Pathway Involved In Phagocytosis (GO:0038096)
Fc Receptor Mediated Stimulatory Signaling Pathway (GO:0002431)
Fc-gamma Receptor Signaling Pathway (GO:0038094)
Integrin-Mediated Signaling Pathway (GO:0007229)
Cellular Response To Lipopolysaccharide (GO:0071222)
Positive Regulation Of Chemokine Production (GO:0032722)
Enriched terms
Gene count
P value
b4:
Enrichment analysis by Enrichr
ERAD Pathway (GO:0036503)
Response To Endoplasmic Reticulum Stress (GO:0034976)
Ubiquitin-Dependent ERAD Pathway (GO:0030433)
Negative Regulation Of Programmed Cell Death (GO:0043069)
Negative Regulation Of Apoptotic Process (GO:0043066)
Regulation Of Endoplasmic Reticulum Unfolded Protein Response (GO:1900101)
Signal Peptide Processing (GO:0006465)
Negative Regulation Of Endoplasmic Reticulum Stress-Induced Intrinsic Apoptotic Signaling Pathway (G...
Retrograde Protein Transport, ER To Cytosol (GO:0030970)
Positive Regulation Of Cell Motility (GO:2000147)
Enriched terms
Gene count
P value
b5:
Enrichment analysis by Enrichr
B Cell Activation (GO:0042113)
Regulation Of Interleukin-2 Production (GO:0032663)
Negative Regulation Of Sodium Ion Transmembrane Transporter Activity (GO:2000650)
B Cell Differentiation (GO:0030183)
Regulation Of B Cell Receptor Signaling Pathway (GO:0050855)
Lymphocyte Differentiation (GO:0030098)
Positive Regulation Of Antigen Receptor-Mediated Signaling Pathway (GO:0050857)
Negative Regulation Of Ion Transmembrane Transporter Activity (GO:0032413)
Negative Regulation Of Antigen Receptor-Mediated Signaling Pathway (GO:0050858)
Positive Regulation Of Interleukin-2 Production (GO:0032743)
Enriched terms
Gene count
P value
b6:
Enrichment analysis by Enrichr
Monocyte Chemotaxis (GO:0002548)
Tryptophan Catabolic Process (GO:0006569)
Tryptophan Metabolic Process (GO:0006568)
Aromatic Amino Acid Family Catabolic Process (GO:0009074)
Regulation Of T Cell Proliferation (GO:0042129)
Response To Interleukin-1 (GO:0070555)
Regulation Of B Cell Differentiation (GO:0045577)
Inflammatory Response (GO:0006954)
Response To Tumor Necrosis Factor (GO:0034612)
Regulation Of Muscle Cell Differentiation (GO:0051147)
Enriched terms
Gene count
P value

**Figure S5: CSF B-lineage subclusters show transcriptional differences and varying functions**

a) CSF L1 UMAP and the B cell-lineage populations (encircled by a blue dashed line) subclustered into L2 populations as shown in the right panel.
b) Dotplot showing the major gene markers for the subclusters.
c) GO analyses of the subcluster defining genes. Predominantly, BP subontology was used to determine biological relevance. Subcluster b1 showed enrichment of B-cell receptor (BCR) signaling and B-cell activation, while b2 enriched for antigen-receptor mediated signaling and T-cell activation. Subcluster b3 enriched for cytokine production and regulation of phagocytosis. ERAD (Endoplasmic-Reticulum-associated protein degradation) pathway and response to ER stress were the terms associated with b4 cells, and B-cell activation and differentiation for b5 subcluster. Monocyte chemotaxis and tryptophan metabolism were the terms annotated for b6 population.

### Fig. S6

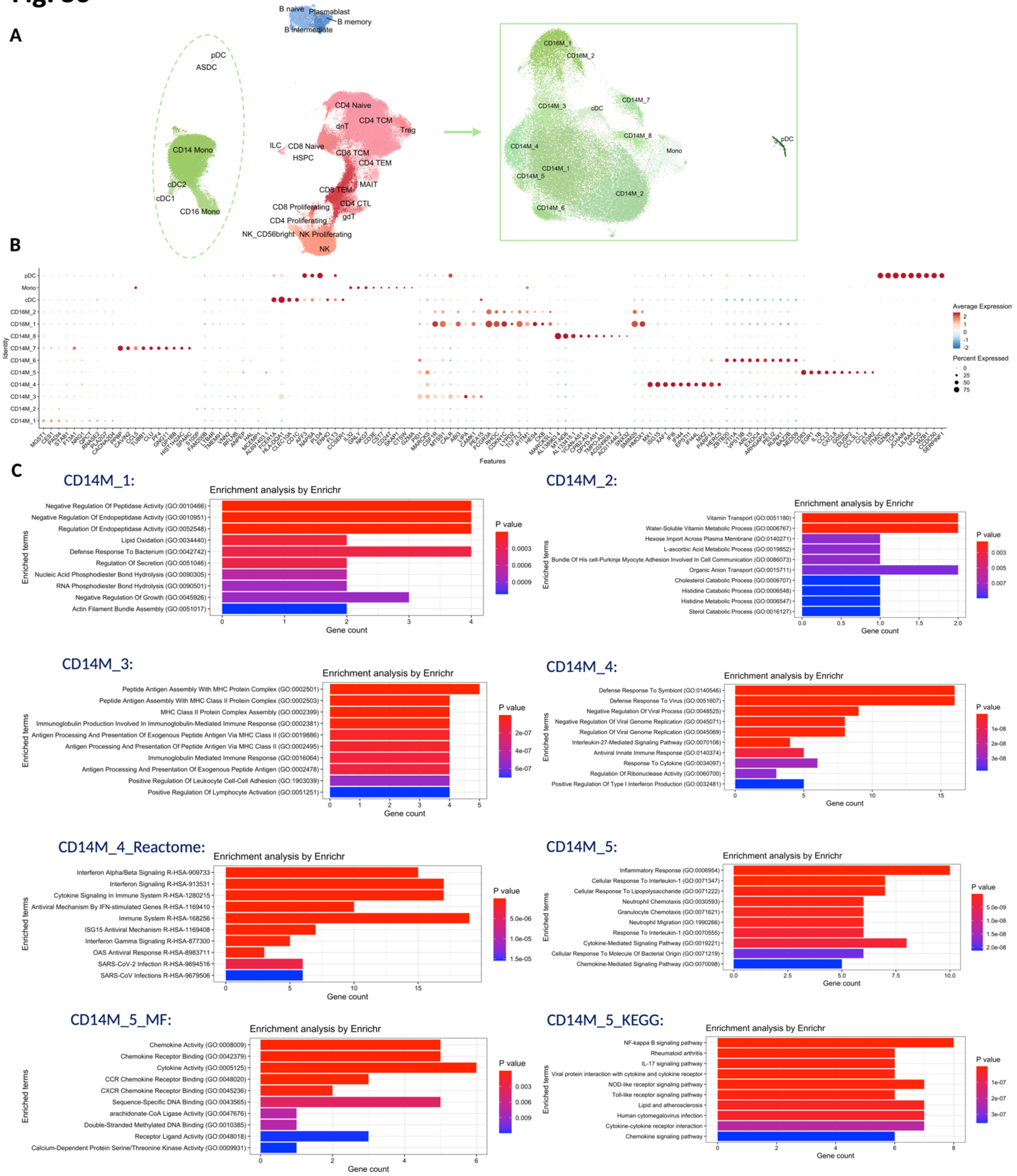

A
B naive
Plasmablast
B memory
B intermediate
pDC
ASDC
CD14 Mono
cDC2
cDC1
CD16 Mono
CD4 Naive
CD4 TCM
dnT
Treg
ILC
CD8 Naive
HSPC
CD8 TCM
CD4 TEM
CD8 TEM
MAIT
CD4 CTL
CD8 Proliferating
gdT
CD4 Proliferating
NK_CD56bright
NK Proliferating
NK
CD16M_1
CD16M_2
CD14M_7
CD14M_3
cDC
CD14M_8
CD14M_4
Mono
pDC
CD14M_1
CD14M_5
CD14M_2
CD14M_6
B
Identity
Features
Average Expression
Percent Expressed
C
CD14M_1:
Enrichment analysis by Enrichr
Negative Regulation Of Peptidase Activity (GO:0010466)
Negative Regulation Of Endopeptidase Activity (GO:0010951)
Regulation Of Endopeptidase Activity (GO:0052548)
Lipid Oxidation (GO:0034440)
Defense Response To Bacterium (GO:0042742)
Regulation Of Secretion (GO:0051046)
Nucleic Acid Phosphodiester Bond Hydrolysis (GO:0090305)
RNA Phosphodiester Bond Hydrolysis (GO:0090501)
Negative Regulation Of Growth (GO:0045926)
Actin Filament Bundle Assembly (GO:0051017)
Enriched terms
Gene count
P value
CD14M_2:
Enrichment analysis by Enrichr
Vitamin Transport (GO:0051180)
Water-Soluble Vitamin Metabolic Process (GO:0006767)
Hexose Import Across Plasma Membrane (GO:0140271)
L-ascorbic Acid Metabolic Process (GO:0019852)
Bundle Of His cell-Purkinje Myocyte Adhesion Involved In Cell Communication (GO:0086073)
Organic Anion Transport (GO:0015711)
Cholesterol Catabolic Process (GO:0006707)
Histidine Catabolic Process (GO:0006548)
Histidine Metabolic Process (GO:0006547)
Sterol Catabolic Process (GO:0016127)
Enriched terms
Gene count
P value
CD14M_3:
Enrichment analysis by Enrichr
Peptide Antigen Assembly With MHC Protein Complex (GO:0002501)
Peptide Antigen Assembly With MHC Class II Protein Complex (GO:0002503)
MHC Class II Protein Complex Assembly (GO:0002399)
Immunoglobulin Production Involved In Immunoglobulin-Mediated Immune Response (GO:0002381)
Antigen Processing And Presentation Of Exogenous Peptide Antigen Via MHC Class II (GO:0019886)
Antigen Processing And Presentation Of Peptide Antigen Via MHC Class II (GO:0002495)
Immunoglobulin Mediated Immune Response (GO:0016064)
Antigen Processing And Presentation Of Exogenous Peptide Antigen (GO:0002478)
Positive Regulation Of Leukocyte Cell-Cell Adhesion (GO:1903039)
Positive Regulation Of Lymphocyte Activation (GO:0051251)
Enriched terms
Gene count
P value
CD14M_4:
Enrichment analysis by Enrichr
Defense Response To Symbiont (GO:0140546)
Defense Response To Virus (GO:0051607)
Negative Regulation Of Viral Process (GO:0048525)
Negative Regulation Of Viral Genome Replication (GO:0045071)
Regulation Of Viral Genome Replication (GO:0045069)
Interleukin-27-Mediated Signaling Pathway (GO:0070106)
Antiviral Innate Immune Response (GO:0140374)
Response To Cytokine (GO:0034097)
Regulation Of Ribonuclease Activity (GO:0060700)
Positive Regulation Of Type I Interferon Production (GO:0032481)
Enriched terms
Gene count
P value
CD14M_4_Reactome:
Enrichment analysis by Enrichr
Interferon Alpha/Beta Signaling R-HSA-909733
Interferon Signaling R-HSA-913531
Cytokine Signaling In Immune System R-HSA-1280215
Antiviral Mechanism By IFN-stimulated Genes R-HSA-1169410
Immune System R-HSA-168256
ISG15 Antiviral Mechanism R-HSA-1169408
Interferon Gamma Signaling R-HSA-877300
OAS Antiviral Response R-HSA-8983711
SARS-CoV-2 Infection R-HSA-9694516
SARS-CoV Infections R-HSA-9679506
Enriched terms
Gene count
P value
CD14M_5:
Enrichment analysis by Enrichr
Inflammatory Response (GO:0006954)
Cellular Response To Interleukin-1 (GO:0071347)
Cellular Response To Lipopolysaccharide (GO:0071222)
Neutrophil Chemotaxis (GO:0030593)
Granulocyte Chemotaxis (GO:0071621)
Neutrophil Migration (GO:1990266)
Response To Interleukin-1 (GO:0070555)
Cytokine-Mediated Signaling Pathway (GO:0019221)
Cellular Response To Molecule Of Bacterial Origin (GO:0071219)
Chemokine-Mediated Signaling Pathway (GO:0070098)
Enriched terms
Gene count
P value
CD14M_5_MF:
Enrichment analysis by Enrichr
Chemokine Activity (GO:0008009)
Chemokine Receptor Binding (GO:0042379)
Cytokine Activity (GO:0005125)
CCR Chemokine Receptor Binding (GO:0048020)
CXCR Chemokine Receptor Binding (GO:0045236)
Sequence-Specific DNA Binding (GO:0043565)
arachidonate-CoA Ligase Activity (GO:0047676)
Double-Stranded Methylated DNA Binding (GO:0010385)
Receptor Ligand Activity (GO:0048018)
Calcium-Dependent Protein Serine/Threonine Kinase Activity (GO:0009931)
Enriched terms
Gene count
P value
CD14M_5_KEGG:
Enrichment analysis by Enrichr
NF-kappa B signaling pathway
Rheumatoid arthritis
IL-17 signaling pathway
Viral protein interaction with cytokine and cytokine receptor
NOD-like receptor signaling pathway
Toll-like receptor signaling pathway
Lipid and atherosclerosis
Human cytomegalovirus infection
Cytokine-cytokine receptor interaction
Chemokine signaling pathway
Enriched terms
Gene count
P value

#### Fig. S6 continued

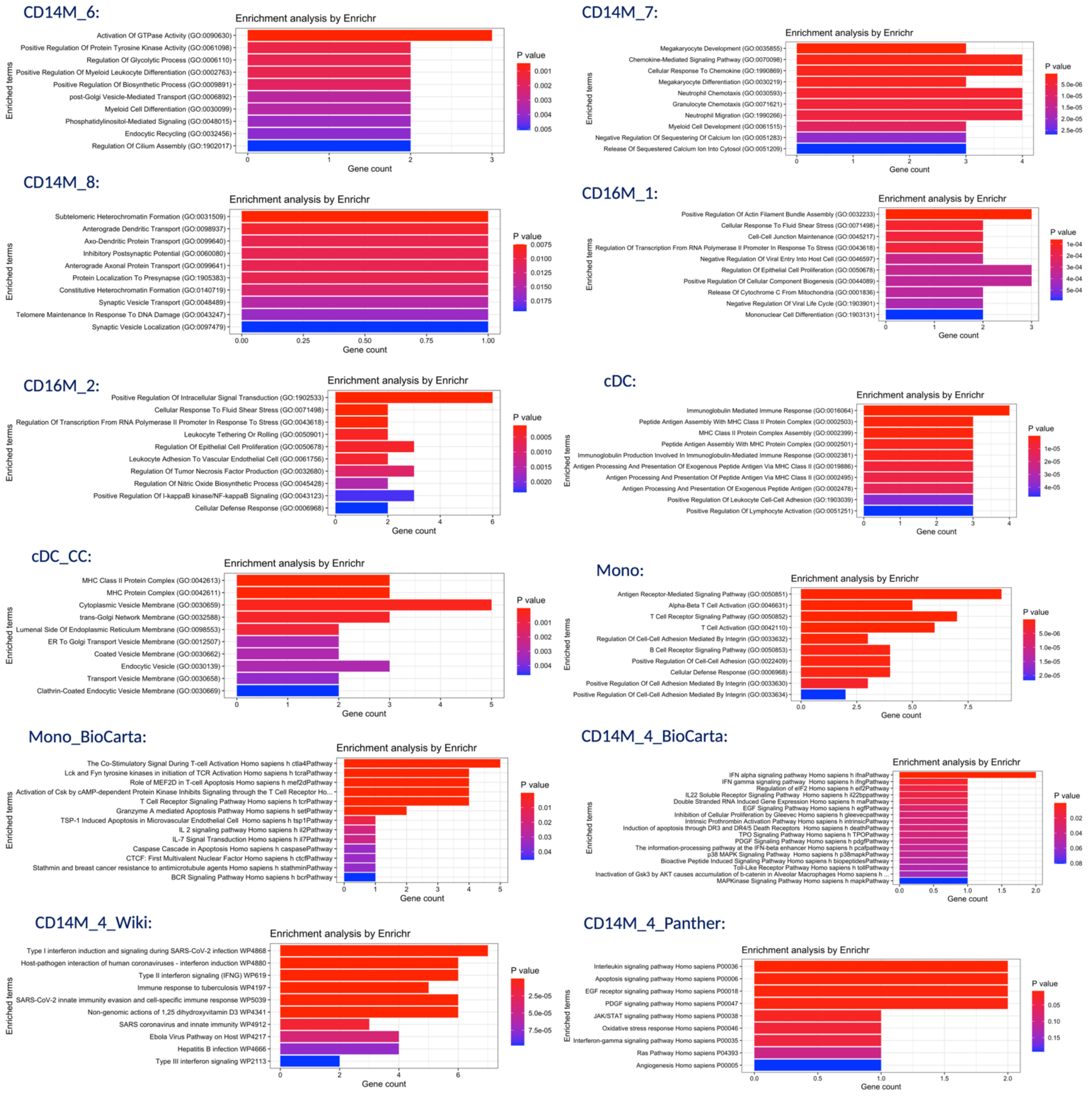

**Figure S6: Peripheral myeloid cells subclusters and their ontologies**

a) PBMC L1 UMAP showing the myeloid cells and the subclusters in the right panel. The subclustering was performed on CD14 monocytes, CD16 monocytes, dendritic cells (cDC1, cDC2, ASDC, and pDCs).
b) Dotplot illustrating the main cell markers for the peripheral myeloid subclusters.
c) GO terms associated with CD14M_1 included negative regulation of peptidase activity and lipid oxidation. CD14M_2 monocytes enriched for vitamin-pertinent transport and metabolic processes. CD14M_3 monocytes had associations with peptide antigen assembly with MHC protein complex and MHC-II protein complex assembly. Notably, CD14M_4 monocytes enriched for terms including defense response to virus, antiviral immune response, positive regulation of type-I interferon production and type-II interferon signaling. Reactome subontology for CD14M_4 monocytes illustrate its role in interferon $\alpha/\beta$ (IFN-$\alpha/\beta$) signaling, cytokine signaling, ISG15-antiviral mechanisms, and IFN-$\gamma$ signaling. CD14M_5 monocytes enrich for inflammatory responses relevant to IL1 and LPS, and various chemotactic responses involving CXCR and CCR chemokine receptor binding. This subcluster is also enriched in NF-$\kappa$B, IL-17, NOD-like receptor and TLR signaling pathways. Note, additional subontologies mentioned above helped identify these annotations. CD14M_6 subcluster is enriched for activation of GTPase activity, positive regulation of protein tyrosine kinase and positive regulation of myeloid leukocyte differentiation. CD16M subclusters associated with cellular response to fluid shear stress and intracellular signal transduction. cDC, as expected, enrich in peptide antigen assembly with MHC-II protein complex, trans-Golgi network and ER-related processes.

#### Fig. S7

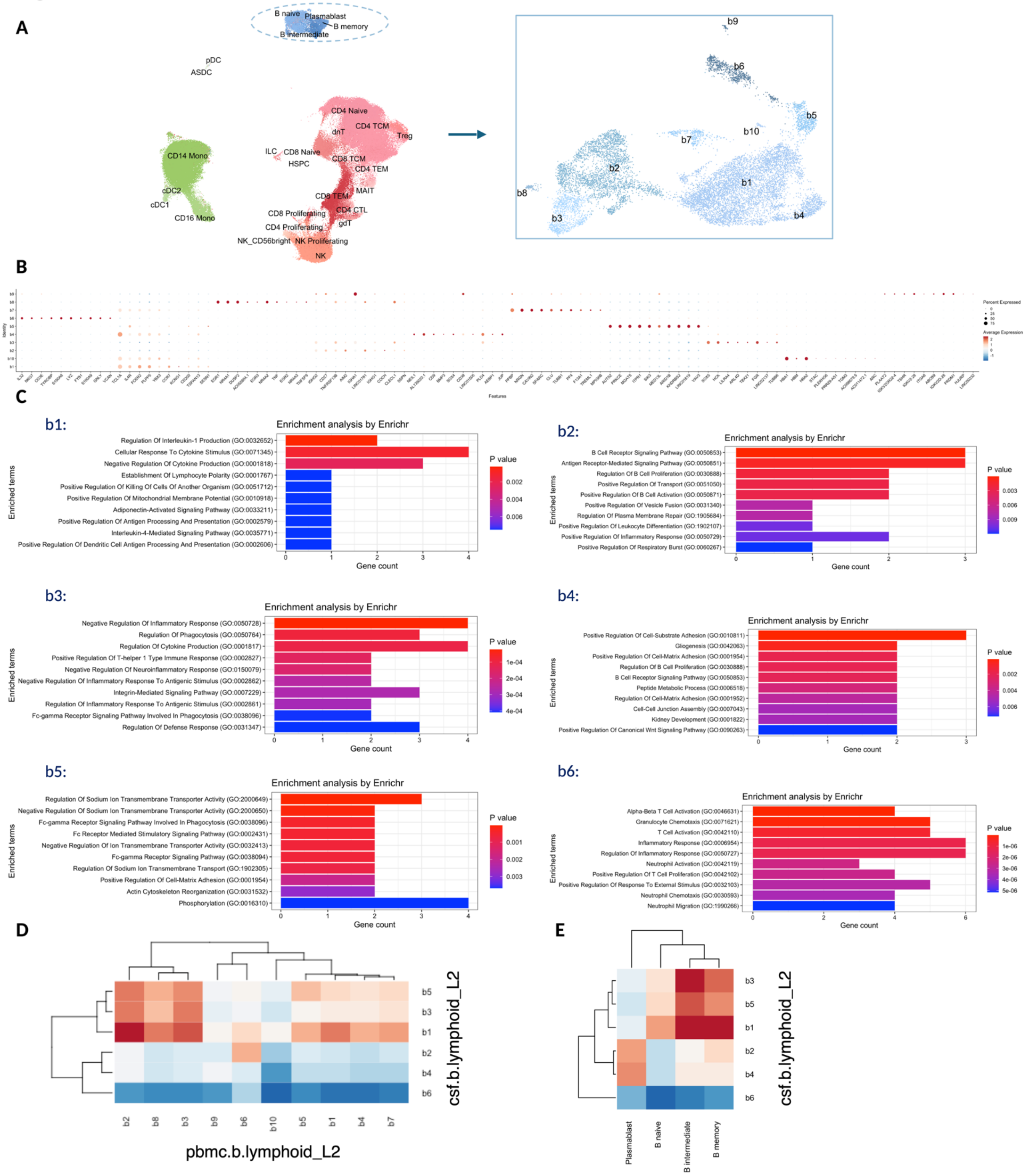
A
B naive
Plasmablast
B memory
B intermediate
pDC
ASDC
CD4 Naive
CD4 TCM
dnT
Treg
CD14 Mono
ILC
CD8 Naive
HSPC
CD8 TCM
CD4 TEM
MAIT
CD8 TEM
cDC2
cDC1
CD16 Mono
CD8 Proliferating
CD4 CTL
CD4 Proliferating
gdT
NK_CD56bright
NK Proliferating
NK
b9
b6
b5
b10
b7
b2
b1
b8
b3
b4
B
C
b1:
Enrichment analysis by Enrichr
Regulation Of Interleukin-1 Production (GO:0032652)
Cellular Response To Cytokine Stimulus (GO:0071345)
Negative Regulation Of Cytokine Production (GO:0001818)
Establishment Of Lymphocyte Polarity (GO:0001767)
Positive Regulation Of Killing Of Cells Of Another Organism (GO:0051712)
Positive Regulation Of Mitochondrial Membrane Potential (GO:0010918)
Adiponectin-Activated Signaling Pathway (GO:0033211)
Positive Regulation Of Antigen Processing And Presentation (GO:0002579)
Interleukin-4-Mediated Signaling Pathway (GO:0035771)
Positive Regulation Of Dendritic Cell Antigen Processing And Presentation (GO:0002606)
Enriched terms
Gene count
P value
b2:
Enrichment analysis by Enrichr
B Cell Receptor Signaling Pathway (GO:0050853)
Antigen Receptor-Mediated Signaling Pathway (GO:0050851)
Regulation Of B Cell Proliferation (GO:0030888)
Positive Regulation Of Transport (GO:0051050)
Positive Regulation Of B Cell Activation (GO:0050871)
Positive Regulation Of Vesicle Fusion (GO:0031340)
Regulation Of Plasma Membrane Repair (GO:1905684)
Positive Regulation Of Leukocyte Differentiation (GO:1902107)
Positive Regulation Of Inflammatory Response (GO:0050729)
Positive Regulation Of Respiratory Burst (GO:0060267)
Enriched terms
Gene count
P value
b3:
Enrichment analysis by Enrichr
Negative Regulation Of Inflammatory Response (GO:0050728)
Regulation Of Phagocytosis (GO:0050764)
Regulation Of Cytokine Production (GO:0001817)
Positive Regulation Of T-helper 1 Type Immune Response (GO:0002827)
Negative Regulation Of Neuroinflammatory Response (GO:0150079)
Negative Regulation Of Inflammatory Response To Antigenic Stimulus (GO:0002862)
Integrin-Mediated Signaling Pathway (GO:0007229)
Regulation Of Inflammatory Response To Antigenic Stimulus (GO:0002861)
Fc-gamma Receptor Signaling Pathway Involved In Phagocytosis (GO:0038096)
Regulation Of Defense Response (GO:0031347)
Enriched terms
Gene count
P value
b4:
Enrichment analysis by Enrichr
Positive Regulation Of Cell-Substrate Adhesion (GO:0010811)
Gliogenesis (GO:0042063)
Positive Regulation Of Cell-Matrix Adhesion (GO:0001954)
Regulation Of B Cell Proliferation (GO:0030888)
B Cell Receptor Signaling Pathway (GO:0050853)
Peptide Metabolic Process (GO:0006518)
Regulation Of Cell-Matrix Adhesion (GO:0001952)
Cell-Cell Junction Assembly (GO:0007043)
Kidney Development (GO:0001822)
Positive Regulation Of Canonical Wnt Signaling Pathway (GO:0090263)
Enriched terms
Gene count
P value
b5:
Enrichment analysis by Enrichr
Regulation Of Sodium Ion Transmembrane Transporter Activity (GO:2000649)
Negative Regulation Of Sodium Ion Transmembrane Transporter Activity (GO:2000650)
Fc-gamma Receptor Signaling Pathway Involved In Phagocytosis (GO:0038096)
Fc Receptor Mediated Stimulatory Signaling Pathway (GO:0002431)
Negative Regulation Of Ion Transmembrane Transporter Activity (GO:0032413)
Fc-gamma Receptor Signaling Pathway (GO:0038094)
Regulation Of Sodium Ion Transmembrane Transport (GO:1902305)
Positive Regulation Of Cell-Matrix Adhesion (GO:0001954)
Actin Cytoskeleton Reorganization (GO:0031532)
Phosphorylation (GO:0016310)
Enriched terms
Gene count
P value
b6:
Enrichment analysis by Enrichr
Alpha-Beta T Cell Activation (GO:0046631)
Granulocyte Chemotaxis (GO:0071621)
T Cell Activation (GO:0042110)
Inflammatory Response (GO:0006954)
Regulation Of Inflammatory Response (GO:0050727)
Neutrophil Activation (GO:0042119)
Positive Regulation Of T Cell Proliferation (GO:0042102)
Positive Regulation Of Response To External Stimulus (GO:0032103)
Neutrophil Chemotaxis (GO:0030593)
Neutrophil Migration (GO:1990266)
Enriched terms
Gene count
P value
D
b5
b3
b1
b2
b4
b6
b2 b8 b3 b9 b6 b10 b5 b1 b4 b7
csf.b.lymphoid_L2
pbmc.b.lymphoid_L2
E
b3
b5
b1
b2
b4
b6
Plasmablast
B naive
B intermediate
B memory
csf.b.lymphoid_L2

**Figure S7: Peripheral B-cell subclusters and congruence with CSF B-lineage cells**

a) PBMC L1 UMAP and representative B-cell populations with the subclusters shown in the right panel
b) Dotplot representing main markers for the subcluster populations of peripheral B-lineage cells
c) GO analyses of the individual peripheral B-lineage subclusters. The B-cell subcluster b1 enriched in regulation of IL-1 production and response to cytokine stimulus, b2 in B-cell receptor signaling, antigen-receptor mediated signaling and B-cell proliferation, b3 in negative regulation of inflammatory response and regulation of phagocytosis, b4 in positive regulation of cell-substrate adhesion, b5 in regulation of $Na^+$ ion transmembrane transporter activity and $F_C\gamma$-receptor mediated pathway involved in phagocytosis, and b6 in $\alpha/\beta$ T cell activation, granulocyte chemotaxis and T cell activation.
d) Correlation heatmap showing transcriptional similarity between blood and CSF B-lineage clusters based on averaged transcriptional expression. Peripheral b6 B cell subcluster resembles the b2 subcluster in the CSF, which aligns with the matching ontologies of their respective gene markers.
e) Correlation heatmap demonstrating transcriptional similarity between L2 CSF B-lymphoid clusters and main L1 CSF B-cell annotations.

### Fig. S8

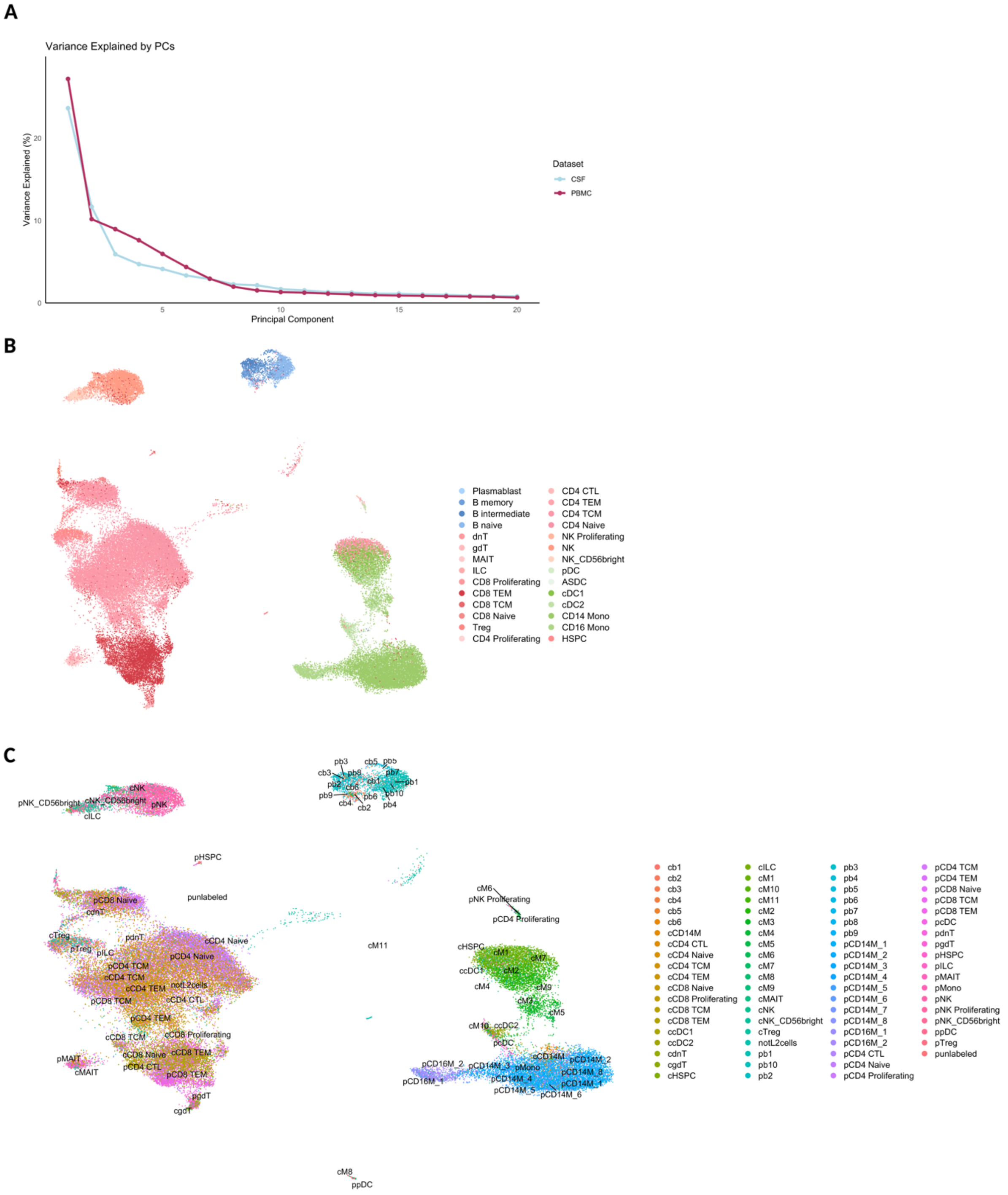
A
Variance Explained by PCs
Variance Explained (%)
Principal Component
Dataset
CSF
PBMC
B
Plasmablast
B memory
B intermediate
B naive
dnT
gdT
MAIT
ILC
CD8 Proliferating
CD8 TEM
CD8 TCM
CD8 Naive
Treg
CD4 Proliferating
CD4 CTL
CD4 TEM
CD4 TCM
CD4 Naive
NK Proliferating
NK
NK_CD56bright
pDC
ASDC
cDC1
cDC2
CD14 Mono
CD16 Mono
HSPC
C
cb1
cb2
cb3
cb4
cb5
cb6
cCD14M
cCD4 CTL
cCD4 Naive
cCD4 TCM
cCD4 TEM
cCD8 Naive
cCD8 Proliferating
cCD8 TCM
cCD8 TEM
ccDC1
ccDC2
cdnT
cgdT
cHSPC
cILC
cM1
cM10
cM11
cM2
cM3
cM4
cM5
cM6
cM7
cM8
cM9
cMAIT
cNK
cNK_CD56bright
cTreg
notL2cells
pb1
pb10
pb2
pb3
pb4
pb5
pb6
pb7
pb8
pb9
pCD14M_1
pCD14M_2
pCD14M_3
pCD14M_4
pCD14M_5
pCD14M_6
pCD14M_7
pCD14M_8
pCD16M_1
pCD16M_2
pCD4 CTL
pCD4 Naive
pCD4 Proliferating
pCD4 TCM
pCD4 TEM
pCD8 Naive
pCD8 TCM
pCD8 TEM
pcDC
pdnT
pgdT
pHSPC
pILC
pMAIT
pMono
pNK
pNK Proliferating
pNK_CD56bright
ppDC
pTreg
punlabeled

**Fig. S8 continued**

D

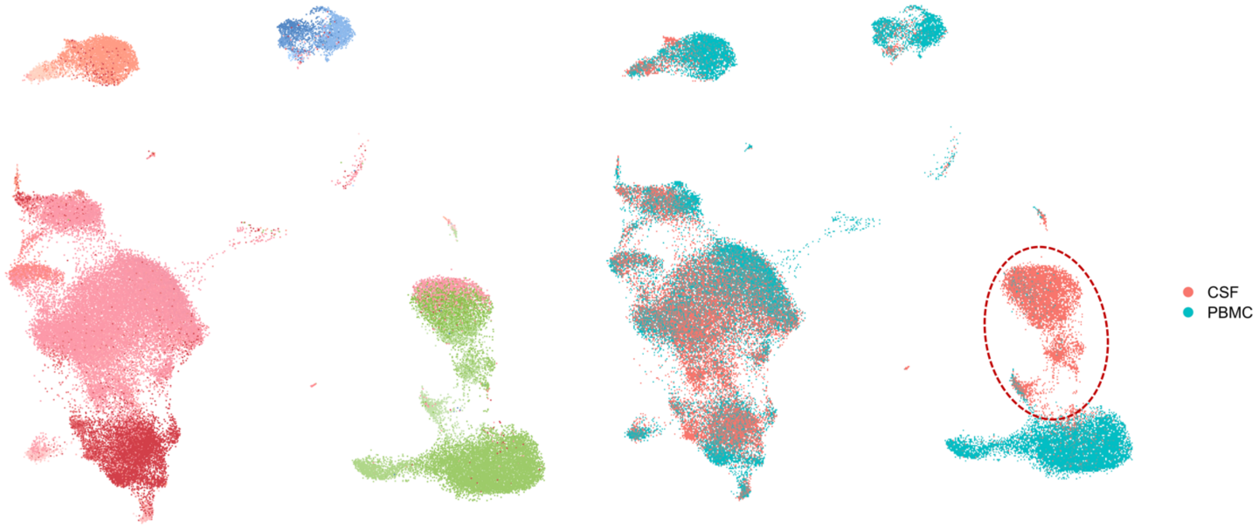


E

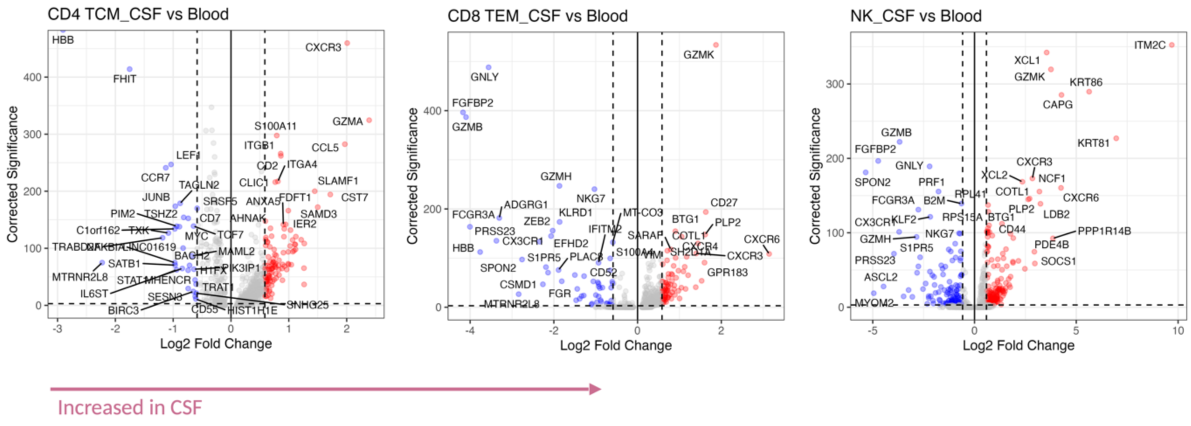


F

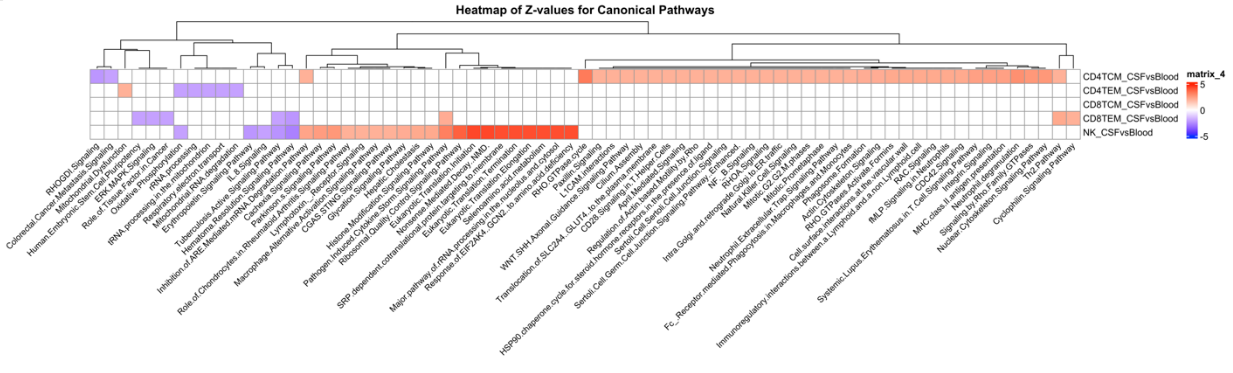

**Figure S8: CD8 and CD4 T-cells upregulate genes associated with trafficking in CSF relative to blood**

a) Elbow plots for the variance explained by the PCs for CSF and PBMC objects.
b) Combined UMAP (CSF cells and PBMC). Note that each sample was randomly down-sampled to 1000 cells before combination (if the sample had less than 1000 cells, the original number of cells were kept).
c) Scatter plot illustrating L2 annotations from CSF cells and PBMC mapped back onto the combined UMAP object. The prefix 'p' represents cells from blood, while the prefix 'c' shows cells from the CSF.
d) Scatter plot showing the immune-cell populations, and the separate clustering of CSF- and blood-derived myeloid cells.
e) Volcano plots contrasting DEG across CSF relative to blood for the CD4-TCM, CD8-TEM, and NK cells from untreated and inactive PRL-positive and PRL-negative cases.
f) Heatmap showing the Z-scores for pathway enrichment across the CSF versus blood comparison for T-lymphoid and NK cells.

**Fig. S9**

A

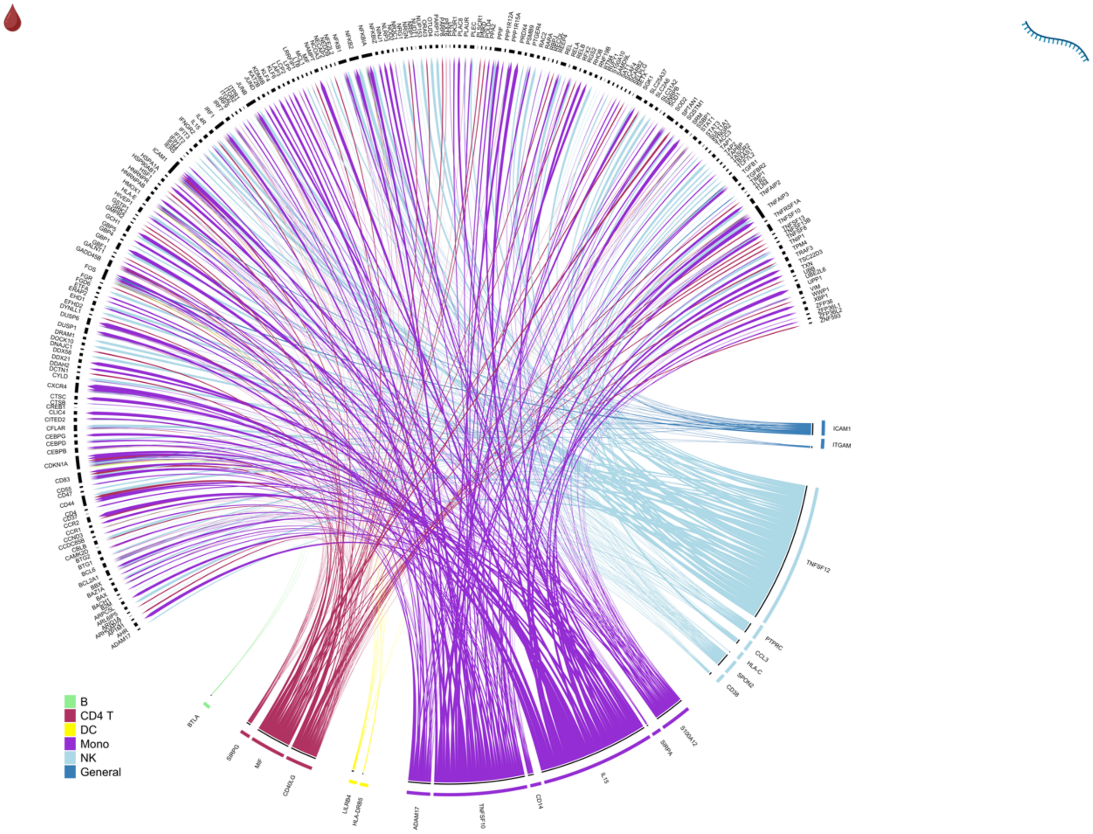

B
CD4 T
DC
Mono
NK
General


B

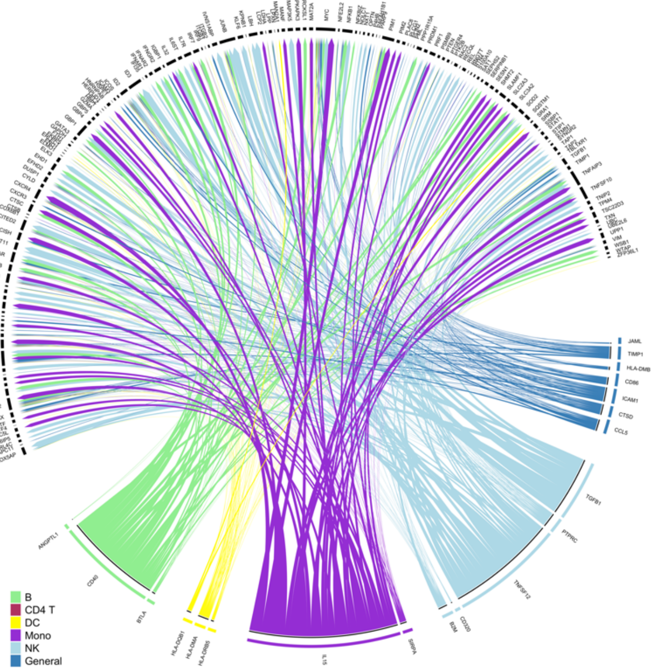

B
CD4 T
DC
Mono
NK
General


C

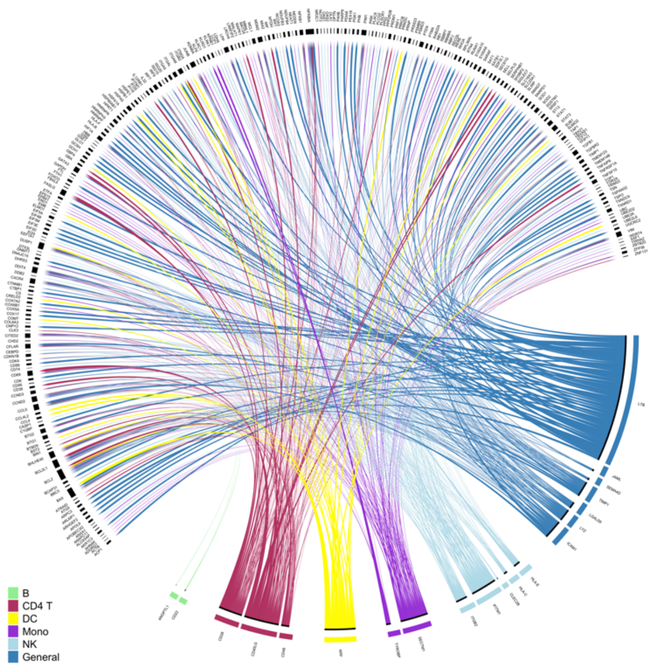

B
CD4 T
DC
Mono
NK
General

#### Fig. S9 continued

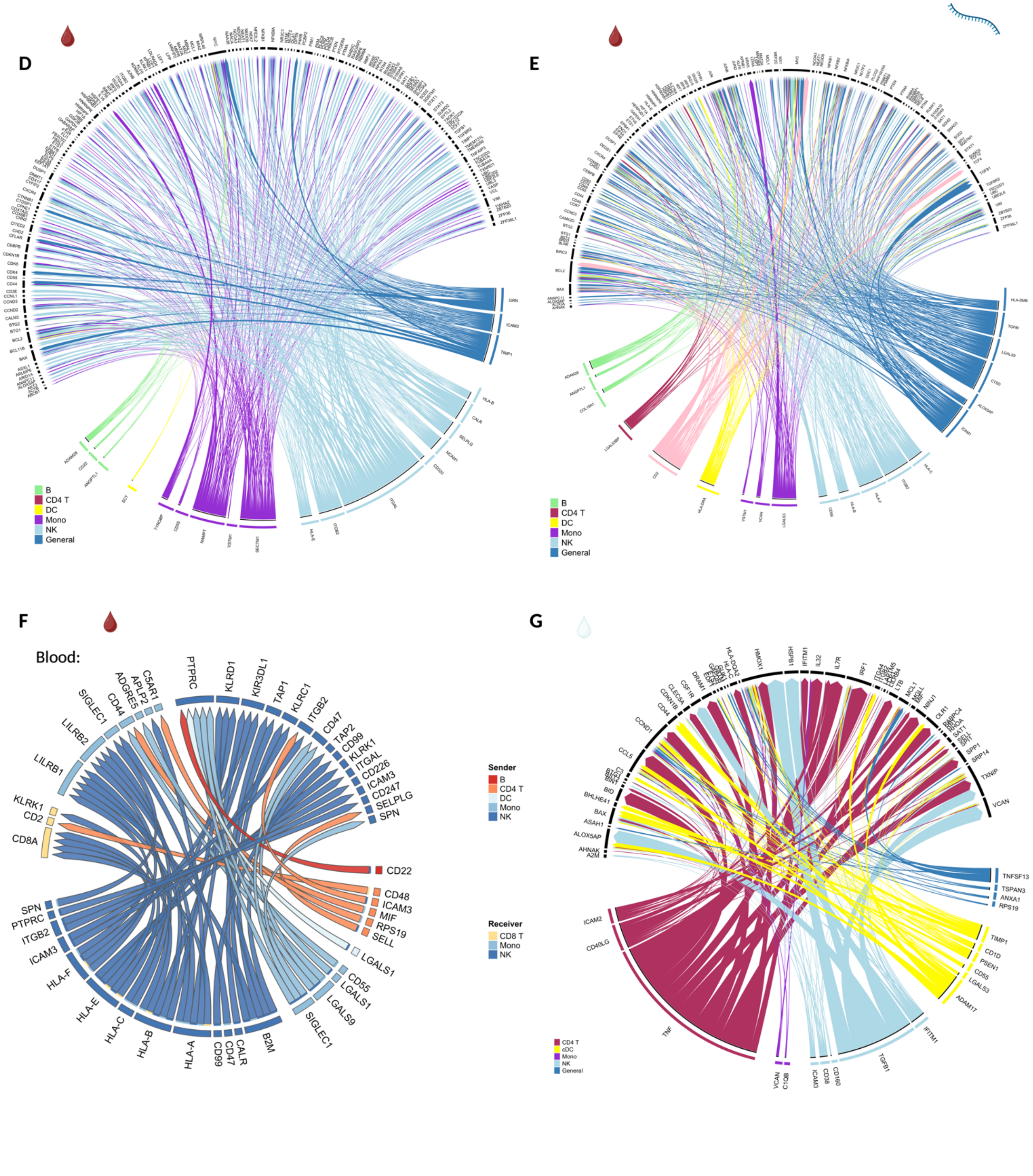

**Figure S9: Interaction analyses between prioritized ligands (senders) and differentially expressed target genes across PRL-positive versus PRL-negative (receivers) reveal a myriad of ligands in regulating PRL-relevant pathology in the blood**

Circos plots summarizing the interactions between prioritized ligands and target genes of myeloid (a), CD4-T (b), NK (c), CD8-T (d), and B (e) cells for PRL-positive versus PRL-negative in the blood. Note the different prioritized ligands for each receiving cell type. These include *HLA-E*, *HLA-A*, *IFITM1*, *LTB*, *CD40LG,* and *CD28* for NK cells; *ITGAL*, *TYROBP,* and *TIMP1* for CD8 T-cells; *IL15*, *TNFSF12*, *TGFB1*, *CD40*, and *CCL5* for CD4 T-cells, and *CD2, ICAM1*, *CTSD*, and *LGALS3* for B-cells*.*

Prioritized LR-network for peripheral immune cells (f) representing differentially abundant interactions with senders being primarily NK cells and receivers including CD8-T, Mono and NK cells. Note, the MHC-I related genes act as ligands in the peripheral network with receptors including *CD8A*, *KLRK1*, *KLRKC*, *LILRB1*, and *SIGLEC1* to name a few.

**Fig. S10**

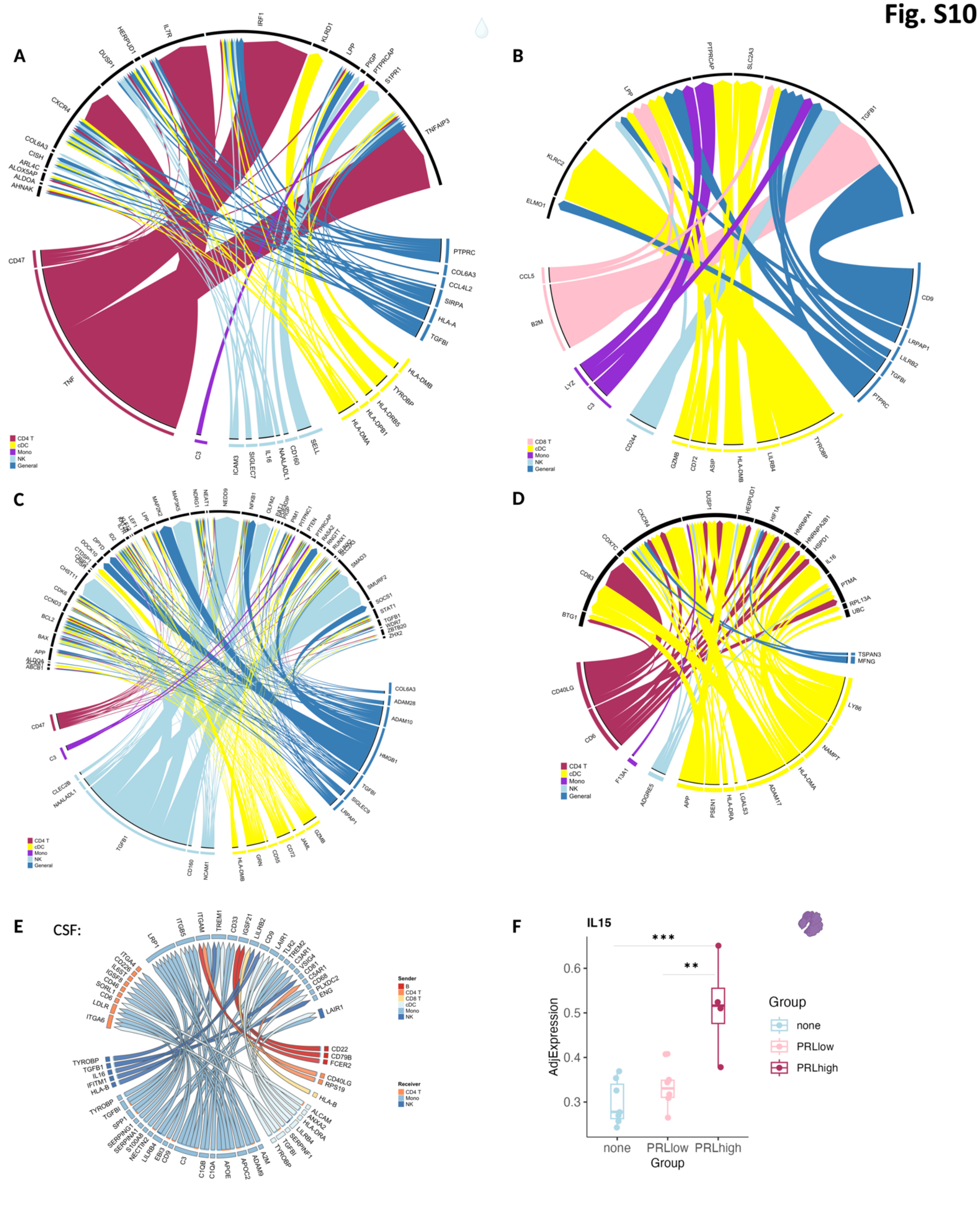

**Figure S10: Interaction analyses between prioritized ligands (senders) and differentially expressed genes across PRL-positive versus PRL-negative as the target genes in the CSF cells reveal the regulatory roles of TNF, TGFB1, and CD47 among many molecules. Regulatory potential of IL15.**

Circos plots summarize the interaction analyses between the prioritized ligands and target genes for myeloid cells (Figure S9G), CD8-T (a), NK (b), CD4-T (c), and B-cells (d) for PRL-positive vs. PRL-negative in the CSF. These include *TNF*, *TGFB1*, and *CD40LG* for myeloid cells (labeled as Mono for these analyses – see **Methods**). For CD8 T-cells, the prioritized ligands regulating the PRL-positive state included *TNF*, *SELL*, and *C3*, and for CD4 T-cells, the prioritized ligands included *TGFB1* and *HMGB1*. *TYROBP, B2M, CD244, C3,* and *CD9* were the prioritized ligands for NK-cells while for B-cells, *ADAM17*, *NAMPT*, and *APP* were the key prioritized ligands.

Prioritized LR-network for the CSF (e) demonstrating differentially abundant interactions contributing to PRL pathology. There is relative predominance of myeloid (Mono) cells as the senders and receivers in the prioritized networks for CSF. Significant interactions involving myeloid activation include *C3*, *C1QA, C1QB, APOE*, *APOC2*, *SPP1*, *TGFBI*, and *TYROBP* as ligands for receptors mostly present on myeloid cells. Notable also are the CD4 T-cells acting as receivers, while B, NK and CD8 T-cells act as the senders. The annotations used for the NicheNet analyses are the simplest level annotations to improve the significance and strength of intercellular communication level analyses.

IL15 abundance in the CSF across patients from different PRL-categories (f).
