## Supplementary File 5 for "Maintenance of chronic neuroinflammation in multiple sclerosis via interferon signaling and CD8 T cell-mediated cytotoxicity"

**M11 greenyellow module**

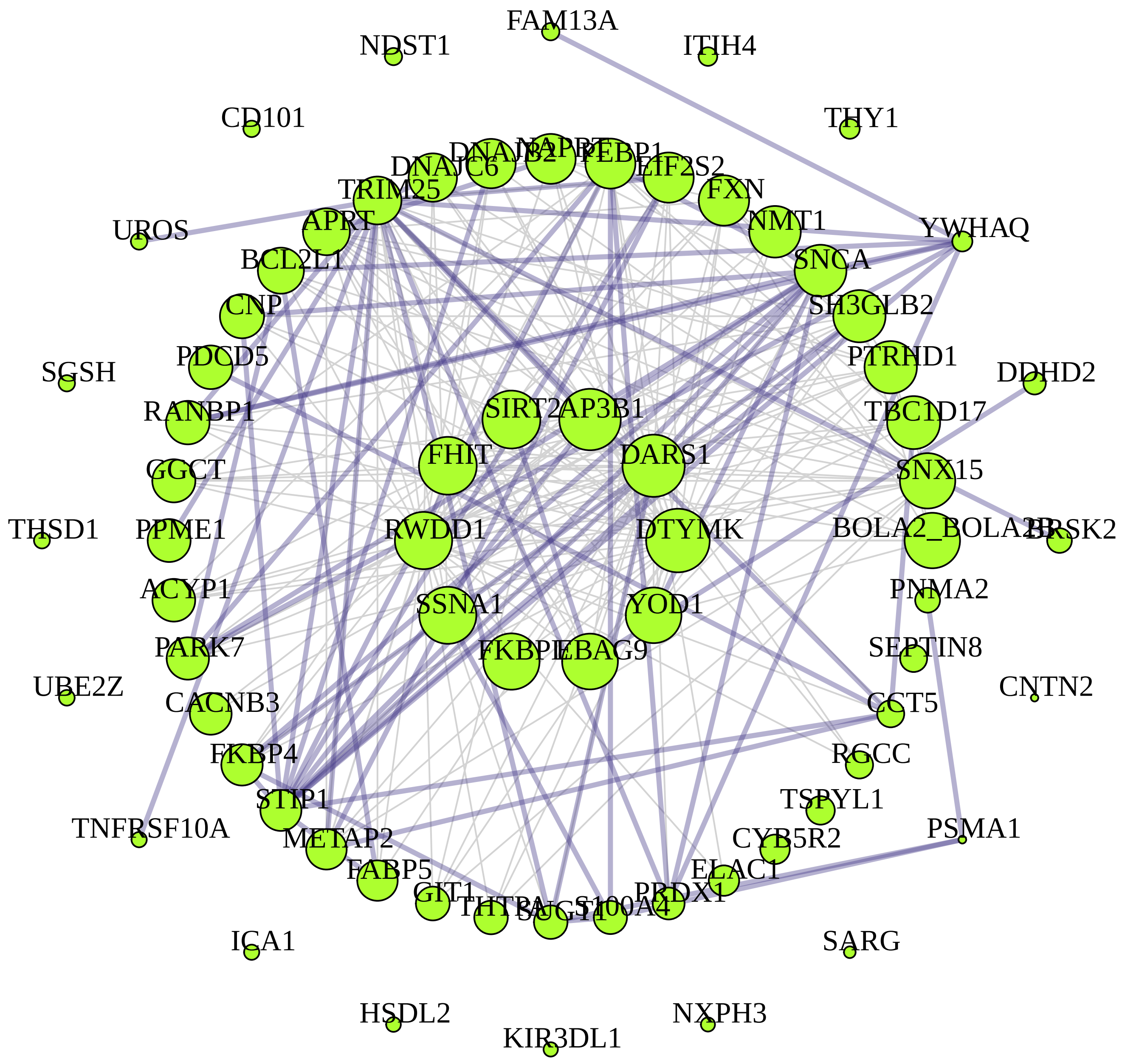


M11 greenyellow module hubs connected by top 374 TOM edges: $HUB^{degree}$

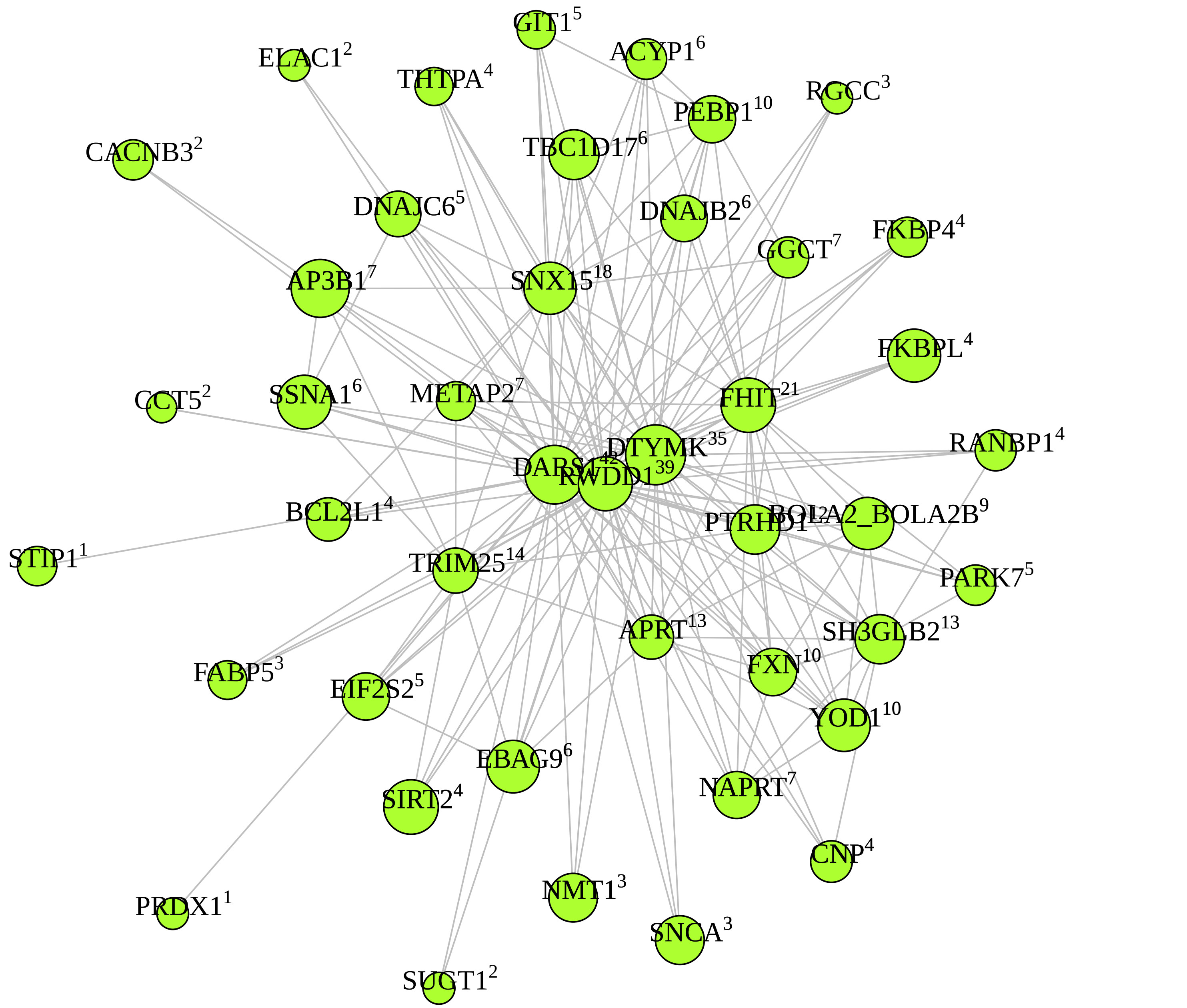

**M1 turquoise module**

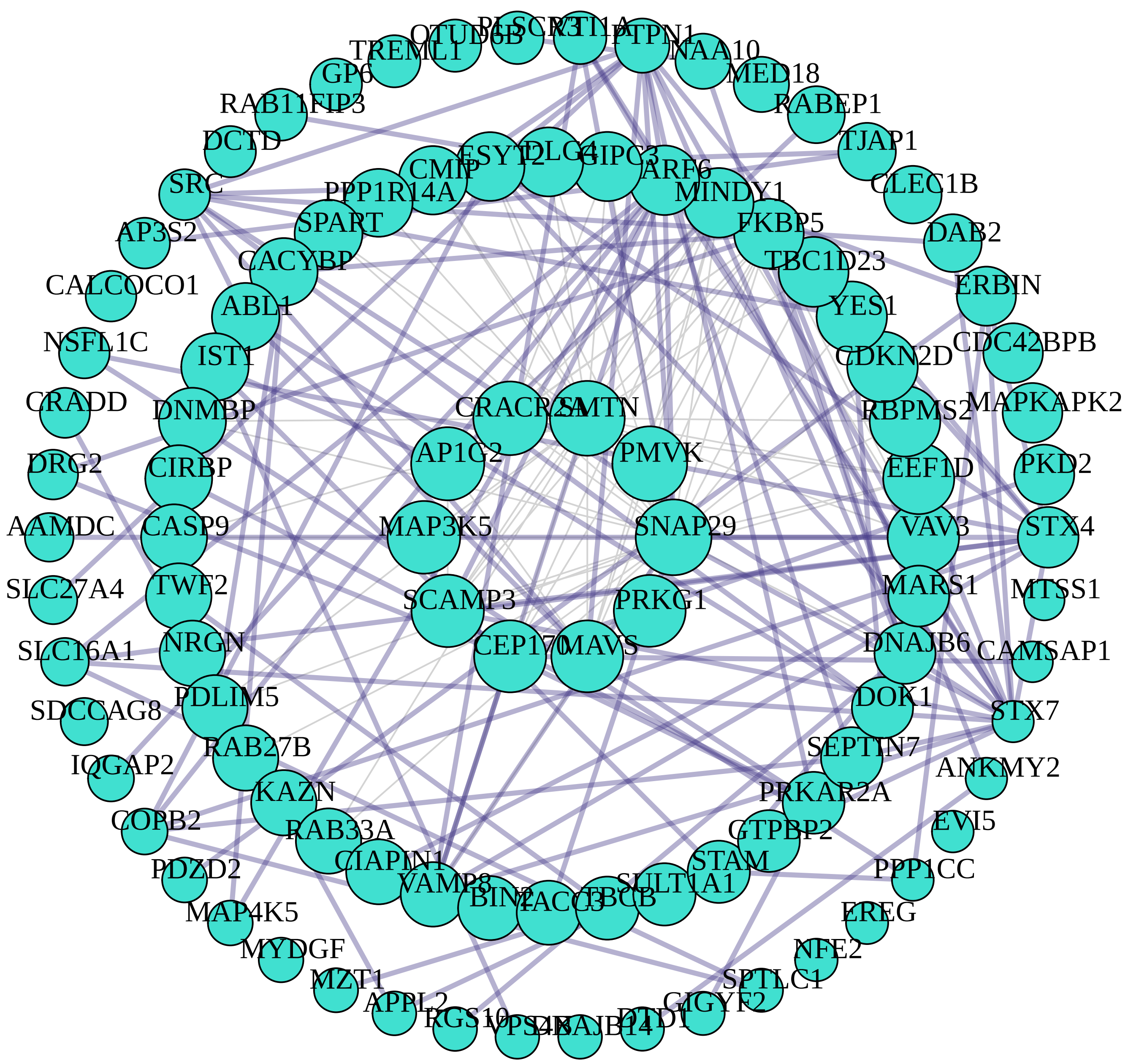


M1 turquoise module hubs connected by top 150 TOM edges: HUB$^{degree}$

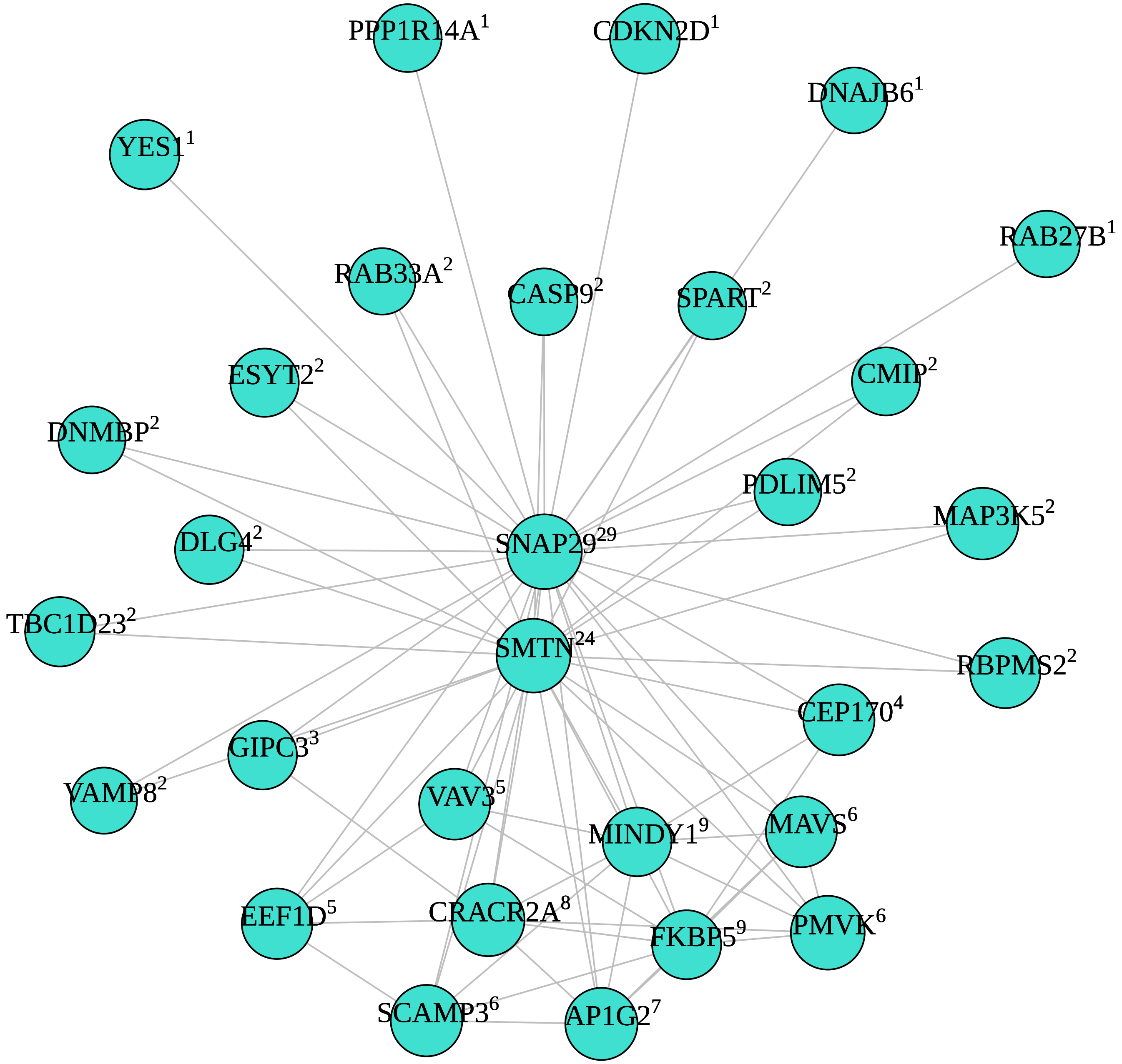

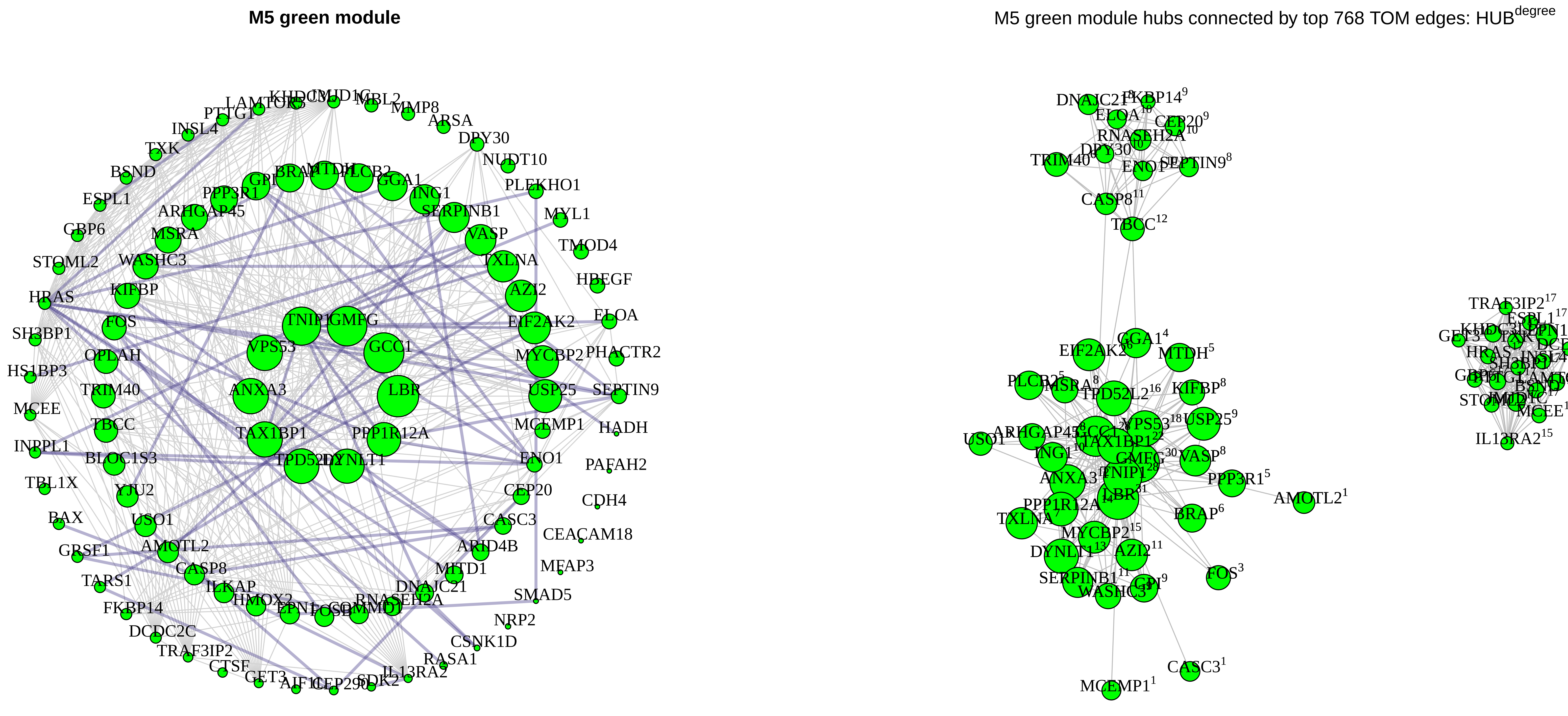

M5 green module
M5 green module hubs connected by top 768 TOM edges: HUB^degree
TNIP1
GMFG
GCC1
VPS53
ANXA3
LBR
TAX1BP1
PPP1R12A
DYNLT1
TXLNA
AZI2
EIF2AK2
MYCBP2
USP25
VASP
SERPINB1
ING1
GGA1
PLCB2
MTDH
BRAP
PPP3R1
ARHGAP45
MSRA
WASHC3
KIFBP
FOS
OPLAH
TRIM40
TBCC
BLOC1S3
YJU2
USO1
AMOTL2
CASP8
ILKAP
HMOX2
EPN1
COMMD1
RNASEH2A
DNAJC21
MITD1
ARID4B
CASC3
CEP20
ENO1
MCEMP1
HRAS
SH3BP1
HS1BP3
MCEE
INPPL1
TBL1X
BAX
GRSF1
TARS1
FKBP14
DCDC2C
TRAF3IP2
CTSF
GET3
AIF1L
SDK2
IL13RA2
RASA1
CSNK1D
NRP2
SMAD5
MFAP3
CEACAM18
CDH4
PAFAH2
HADH
SEPTIN9
PHACTR2
ELOA
HBEGF
TMOD4
MYL1
PLEKHO1
NUDT10
DPY30
ARSA
MMP8
MBL2
INSL4
TXK
BSND
ESPL1
GBP6
STOML2

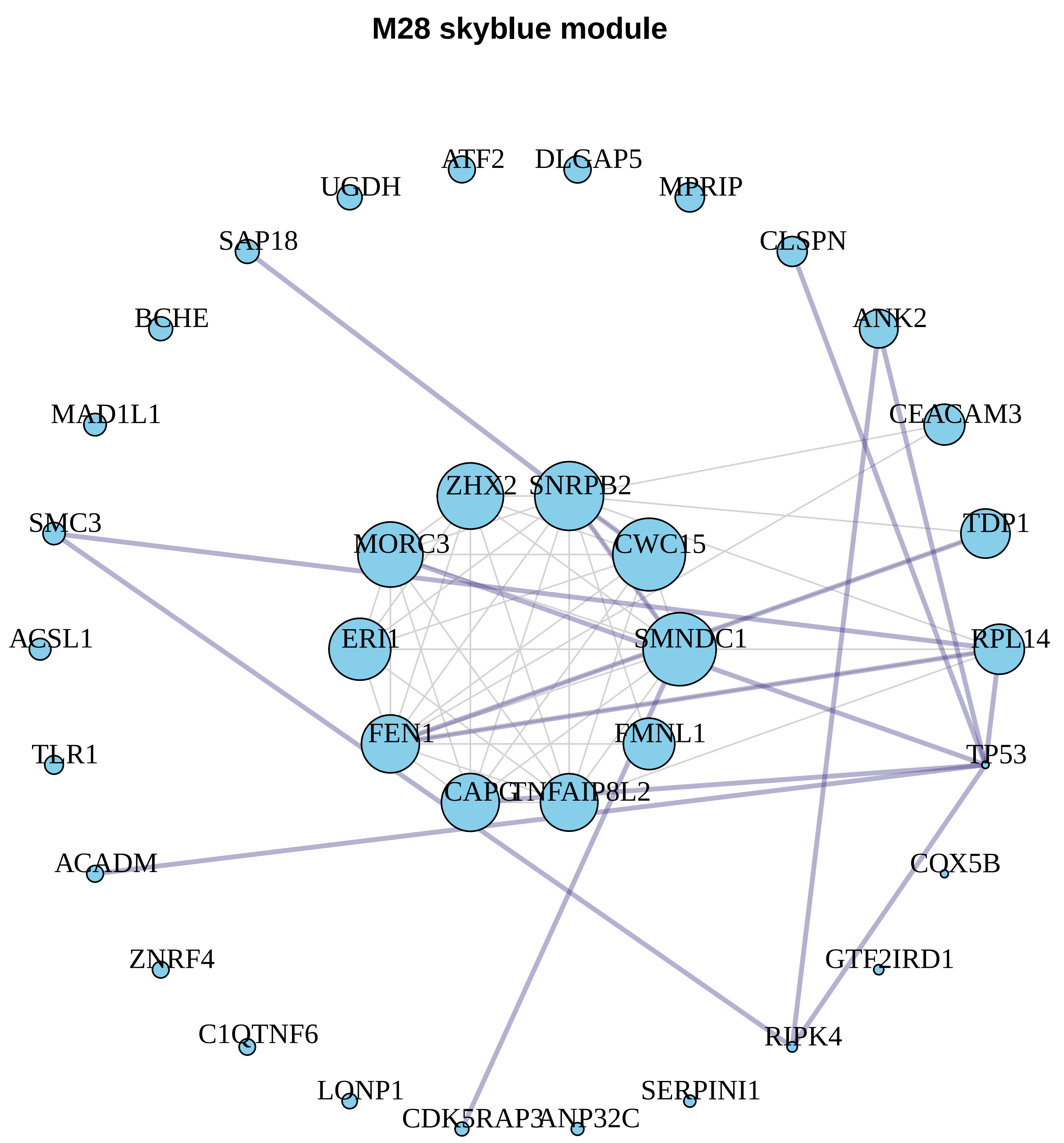
M28 skyblue module
ATF2
DLGAP5
UGDH
MPRIP
SAP18
CLSPN
BCHE
ANK2
MAD1L1
CEACAM3
ZHX2
SNRPB2
SMC3
TDP1
MORC3
CWC15
ACSL1
ERI1
SMNDC1
RPL14
FEN1
FMNL1
TLR1
TP53
CAPG
TNFAIP8L2
ACADM
COX5B
ZNRF4
GTF2IRD1
C1QTNF6
RIPK4
LONP1
SERPINI1
CDK5RAP3
ANP32C

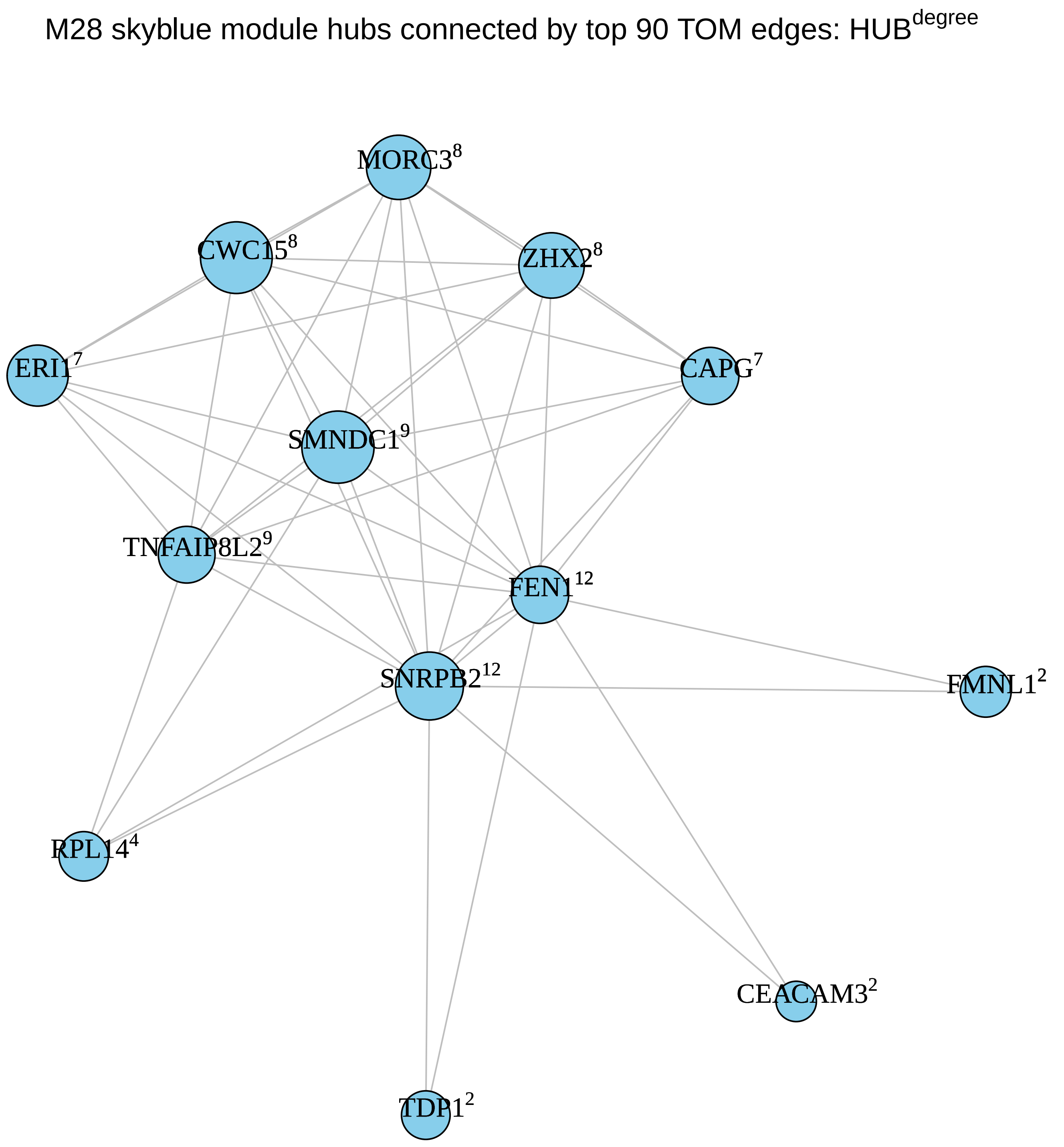
M28 skyblue module hubs connected by top 90 TOM edges: HUB$^{degree}$
MORC3$^{8}$
CWC15$^{8}$
ZHX2$^{8}$
ERI1$^{7}$
CAPG$^{7}$
SMNDC1$^{9}$
TNFAIP8L2$^{9}$
FEN1$^{12}$
SNRPB2$^{12}$
FMNL1$^{2}$
RPL14$^{4}$
CEACAM3$^{2}$
TDP1$^{2}$

**M4 yellow module**

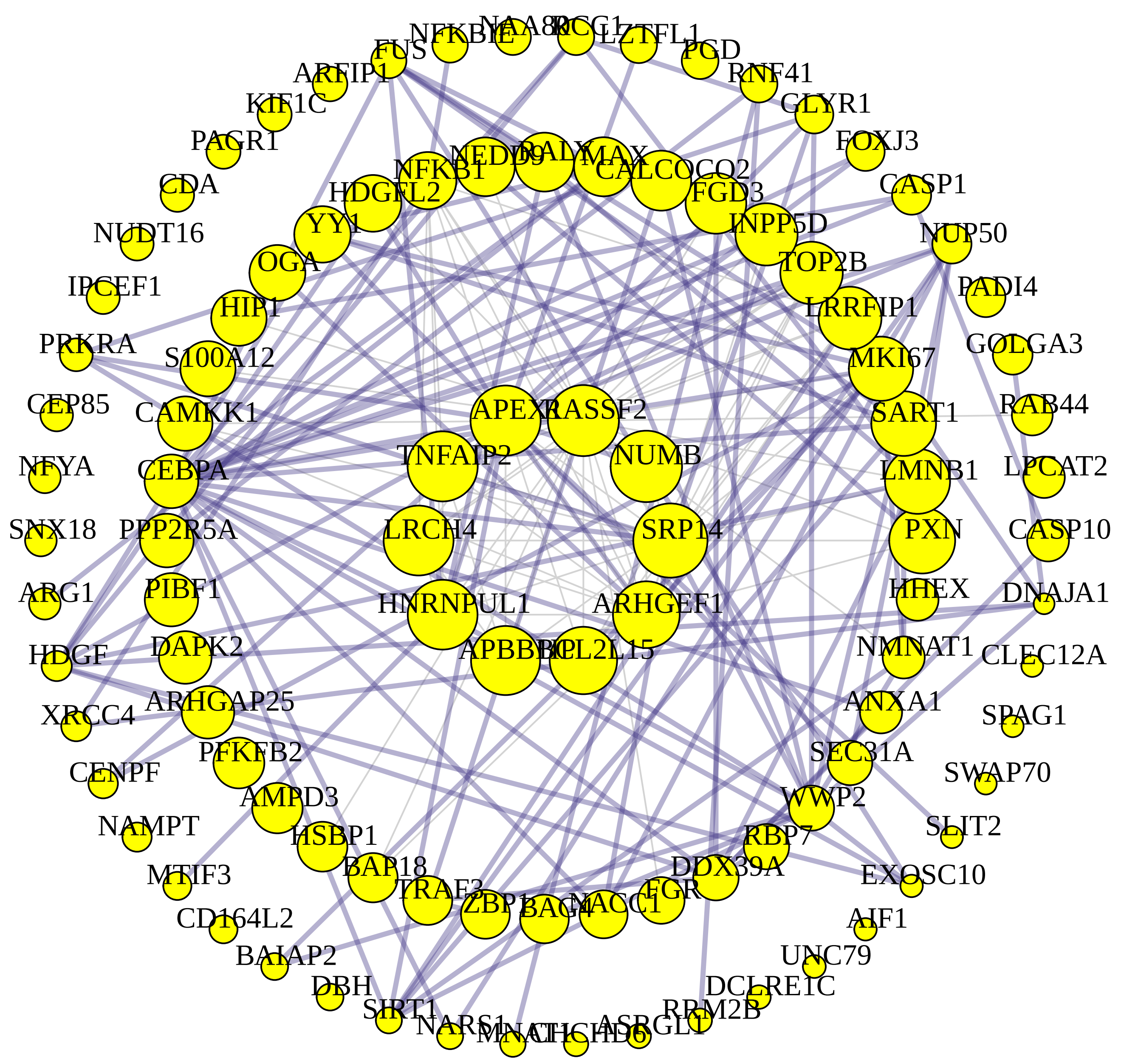


M4 yellow module hubs connected by top 150 TOM edges: HUB$^{degree}$

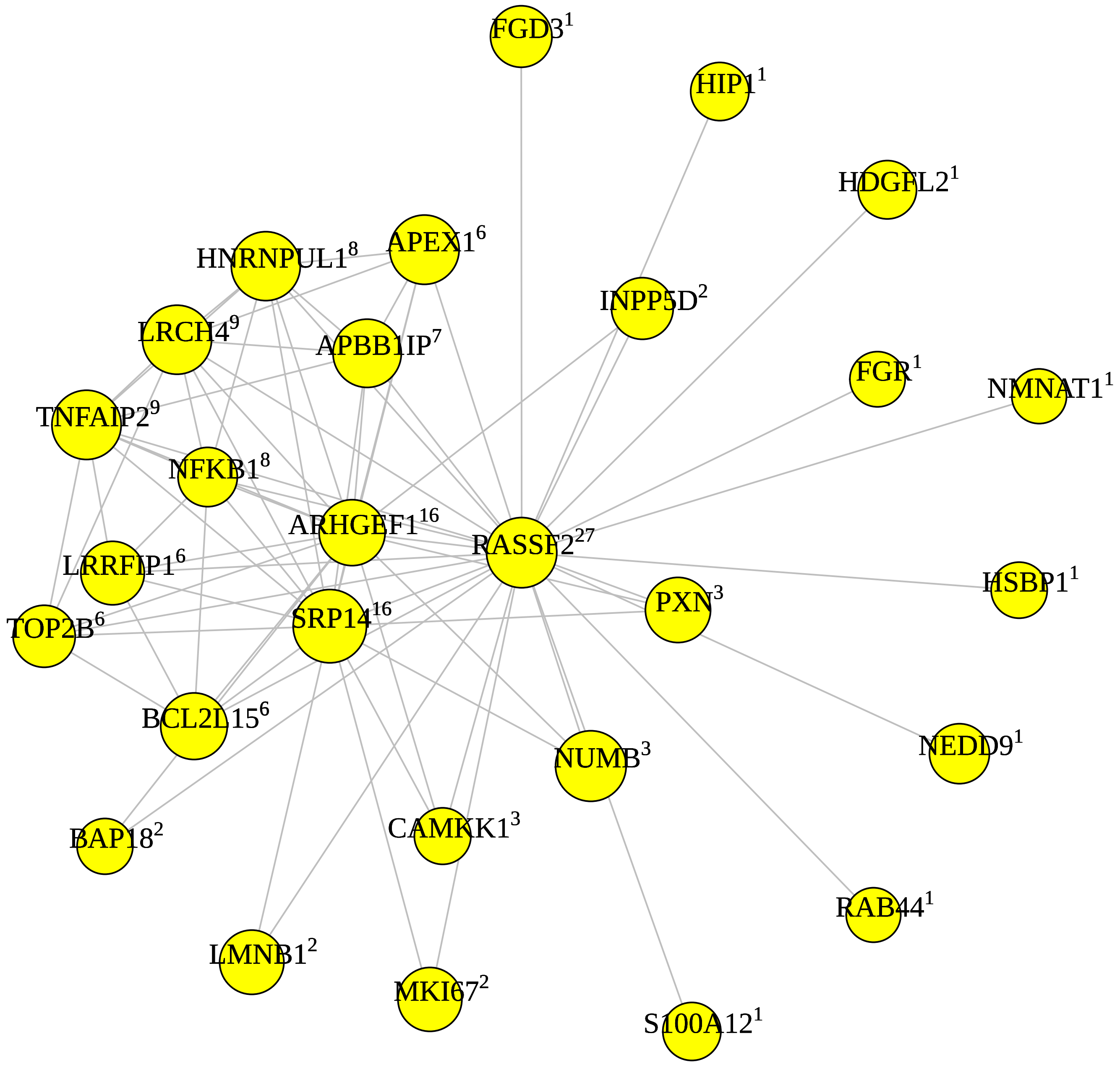

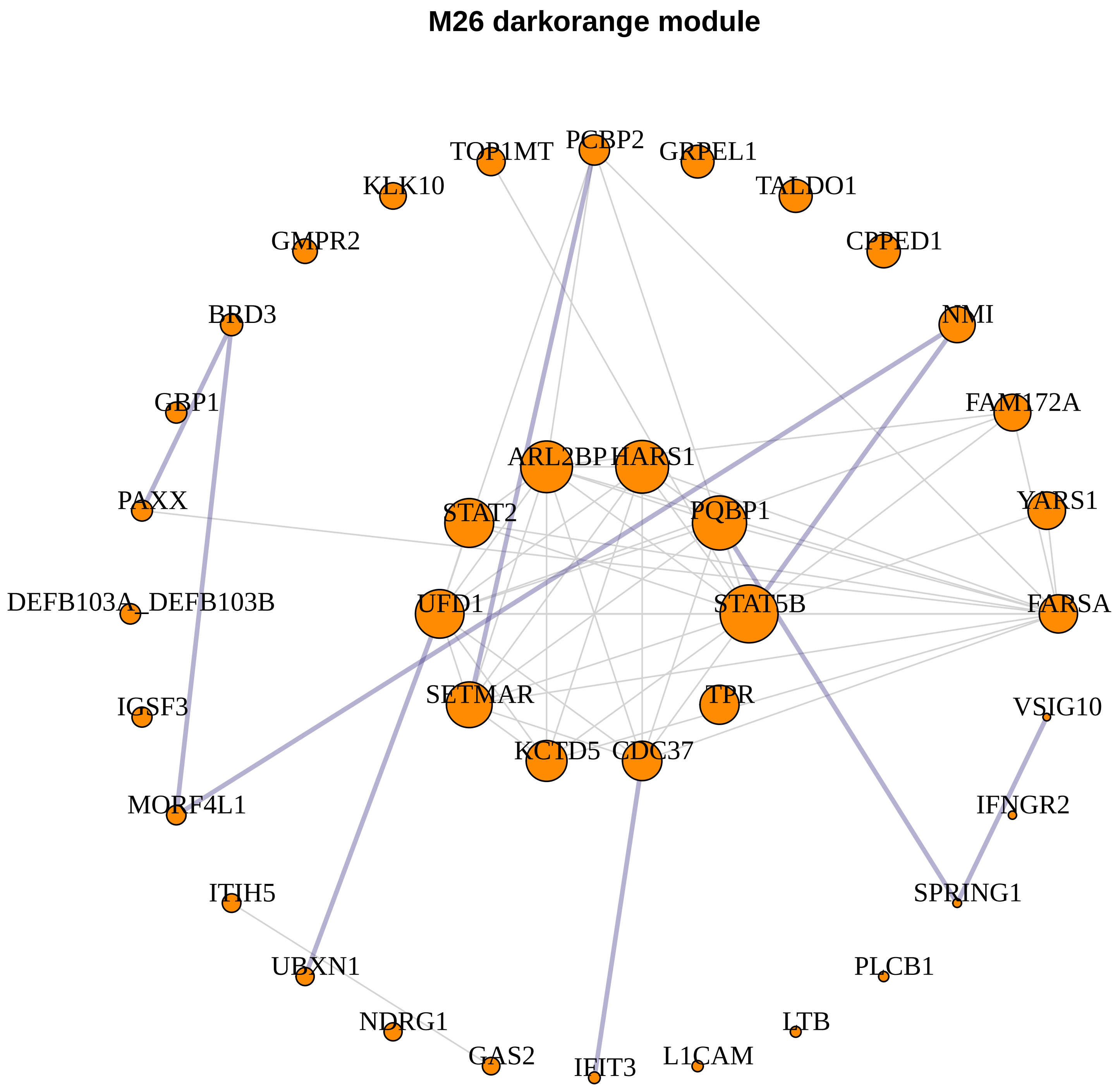
M26 darkorange module
TOP1MT
PCBP2
GRPEL1
KLK10
TALDO1
GMPR2
CPPED1
BRD3
NMI
GBP1
FAM172A
ARL2BP
HARS1
PAXX
STAT2
PQBP1
YARS1
DEFB103A_DEFB103B
UFD1
STAT5B
FARSA
SETMAR
TPR
IGSF3
VSIG10
KCTD5
CDC37
MORF4L1
IFNGR2
ITIH5
SPRING1
UBXN1
PLCB1
NDRG1
LTB
GAS2
IFIT3
L1CAM

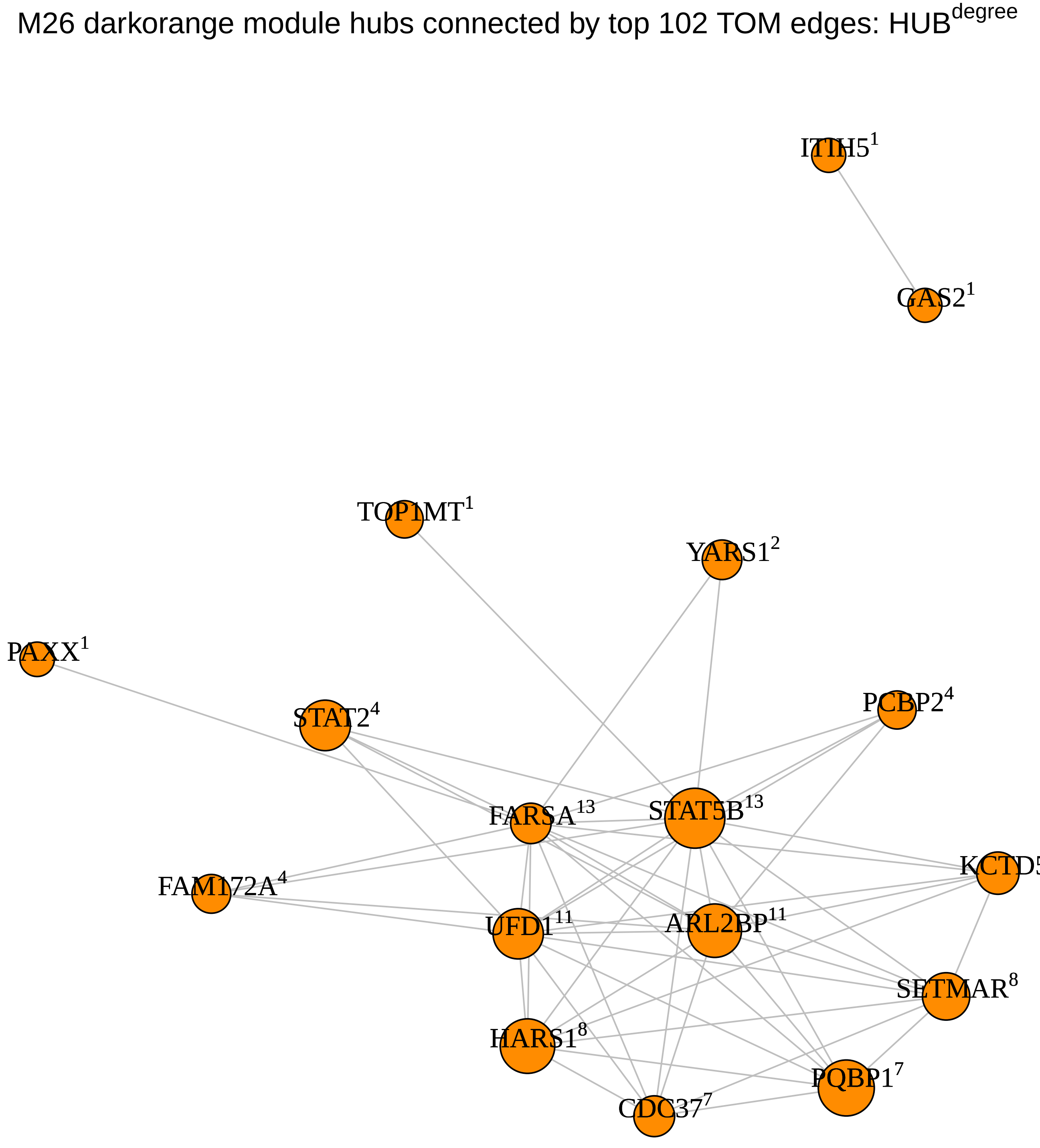
M26 darkorange module hubs connected by top 102 TOM edges: HUB$^{degree}$
ITIH5$^{1}$
GAS2$^{1}$
TOP1MT$^{1}$
YARS1$^{2}$
PAXX$^{1}$
STAT2$^{4}$
PCBP2$^{4}$
FARSA$^{13}$
STAT5B$^{13}$
FAM172A$^{4}$
KCTD5$^{6}$
UFD1$^{11}$
ARL2BP$^{11}$
SETMAR$^{8}$
HARS1$^{8}$
PQBP1$^{7}$
CDC37$^{7}$

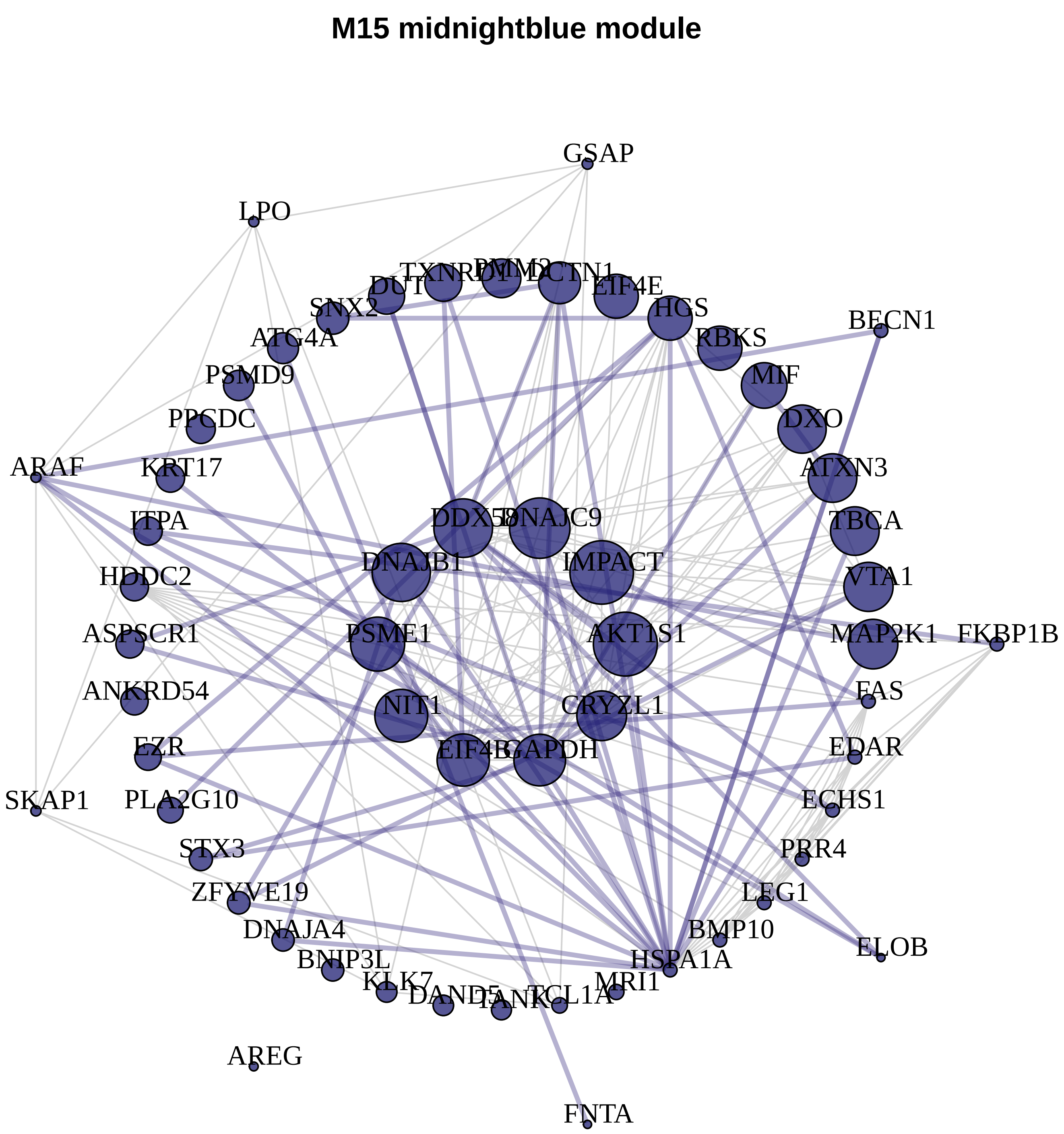
M15 midnightblue module
GSAP
LPO
TXNRD1
PMM2
DUT
DCTN1
EIF4E
SNX2
HGS
BECN1
ATG4A
RBKS
PSMD9
MIF
PPCDC
DXO
ARAF
KRT17
ATXN3
ITPA
DDX58
DNAJC9
TBCA
DNAJB1
IMPACT
HDDC2
VTA1
ASPSCR1
PSME1
AKT1S1
MAP2K1
FKBP1B
ANKRD54
FAS
NIT1
CRYZL1
EZR
EIF4B
GAPDH
EDAR
SKAP1
PLA2G10
ECHS1
STX3
PRR4
ZFYVE19
LEG1
DNAJA4
BMP10
BNIP3L
HSPA1A
ELOB
KLK7
DAND5
TANK
TCL1A
MRI1
AREG
FNTA

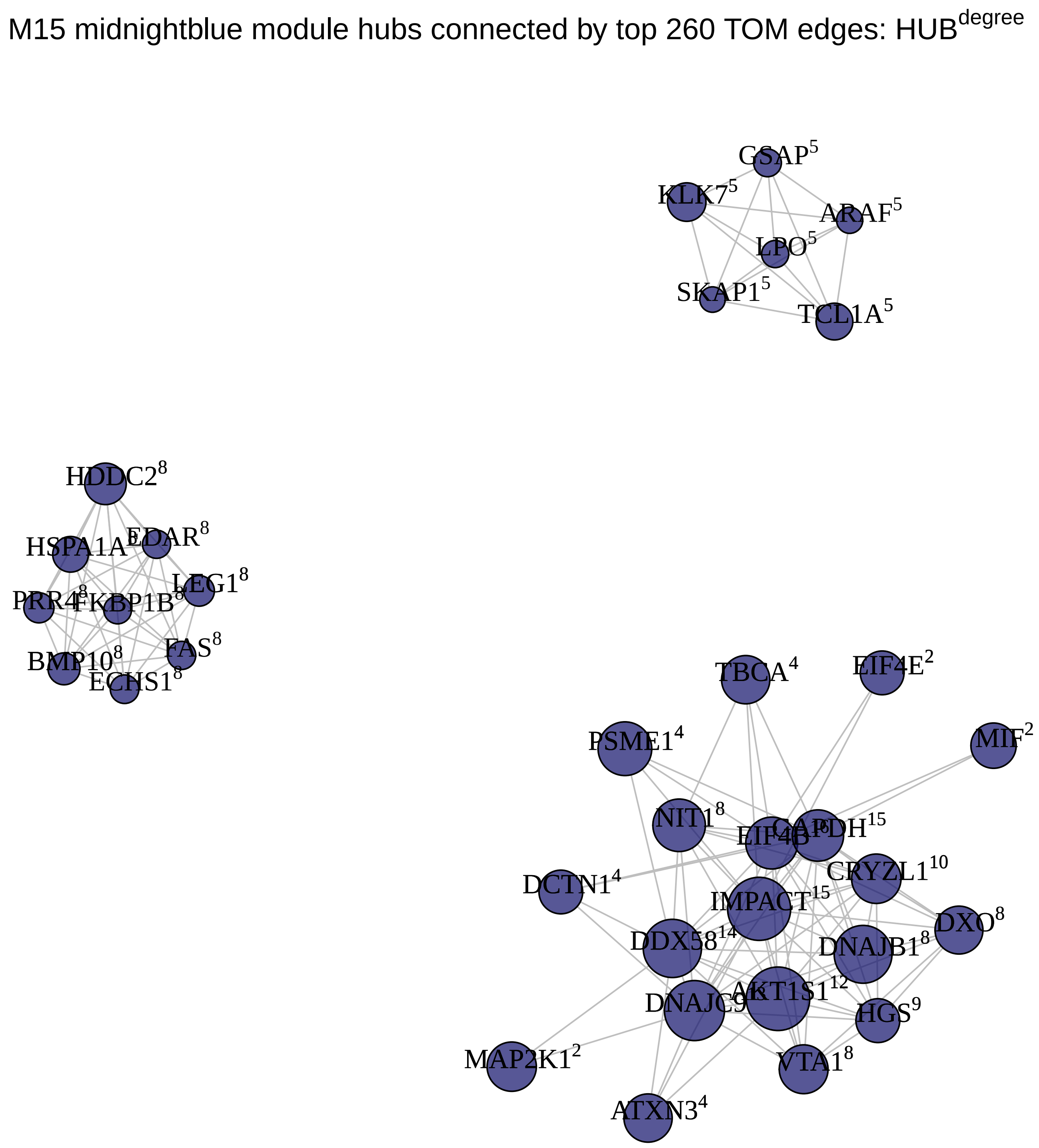
M15 midnightblue module hubs connected by top 260 TOM edges: HUB$^{degree}$
GSAP$^{5}$
KLK7$^{5}$
ARAF$^{5}$
LPO$^{5}$
SKAP1$^{5}$
TCL1A$^{5}$
HDDC2$^{8}$
HSPA1A$^{8}$
EDAR$^{8}$
LEG1$^{8}$
PRR4$^{8}$
FKBP1B$^{8}$
FAS$^{8}$
BMP10$^{8}$
ECHS1$^{8}$
TBCA$^{4}$
EIF4E$^{2}$
MIF$^{2}$
PSME1$^{4}$
NIT1$^{8}$
EIF4B
GAPDH$^{15}$
CRYZL1$^{10}$
DCTN1$^{4}$
IMPACT$^{15}$
DXO$^{8}$
DDX58$^{14}$
DNAJB1$^{8}$
AKT1S1$^{12}$
DNAJC9
HGS$^{9}$
MAP2K1$^{2}$
VTA1$^{8}$
ATXN3$^{4}$

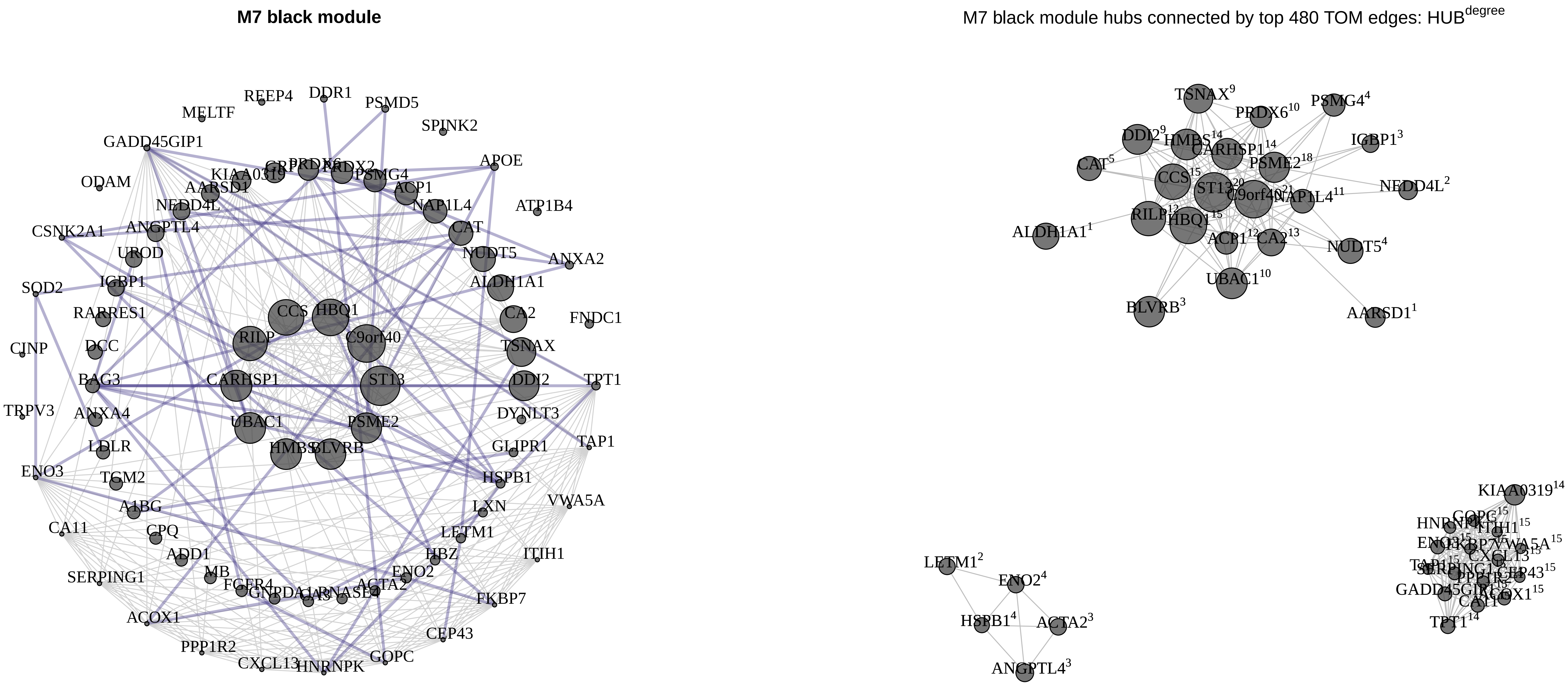
M7 black module
M7 black module hubs connected by top 480 TOM edges: HUB^degree
REEP4
DDR1
PSMD5
MELTF
SPINK2
GADD45GIP1
APOE
ODAM
KIAA0319
AARSD1
PSMG4
ACP1
NEDD4L
NAP1L4
ATP1B4
CSNK2A1
ANGPTL4
CAT
UROD
NUDT5
ANXA2
SOD2
IGBP1
ALDH1A1
RARRES1
CCS
HBQ1
CA2
FNDC1
CINP
DCC
RILP
C9orf40
TSNAX
BAG3
CARHSP1
ST13
DDI2
TPT1
TRPV3
ANXA4
UBAC1
PSME2
DYNLT3
LDLR
HMBS
BLVRB
GLIPR1
TAP1
ENO3
TGM2
HSPB1
A1BG
LXN
VWA5A
CA11
CPQ
LETM1
ADD1
HBZ
ITIH1
SERPING1
MB
ENO2
FGFR4
ACTA2
FKBP7
ACOX1
CEP43
PPP1R2
CXCL13
HNRNPK
GOPC
TSNAX9
PRDX610
PSMG44
DDI29
IGBP13
CAT5
PSME218
NEDD4L2
CCS15
NAP1L411
ALDH1A11
ACP112
CA213
NUDT54
UBAC110
BLVRB3
AARSD11
KIAA031914
LETM12
ENO24
HSPB14
ACTA23
ANGPTL43
VWA5A15
CEP4315
TPT114

**M24 darkgrey module**

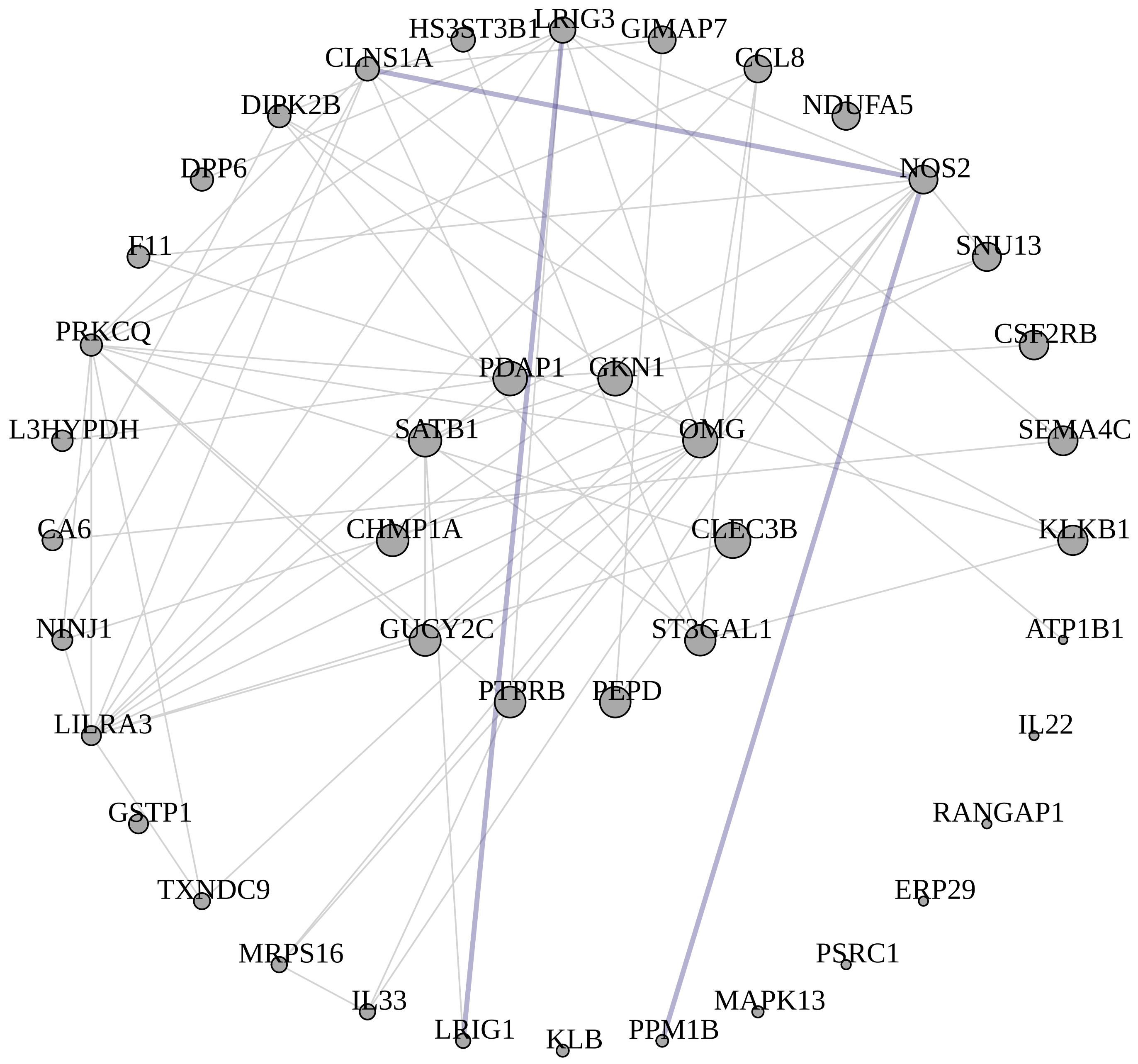


M24 darkgrey module hubs connected by top 126 TOM edges: $HUB^{degree}$

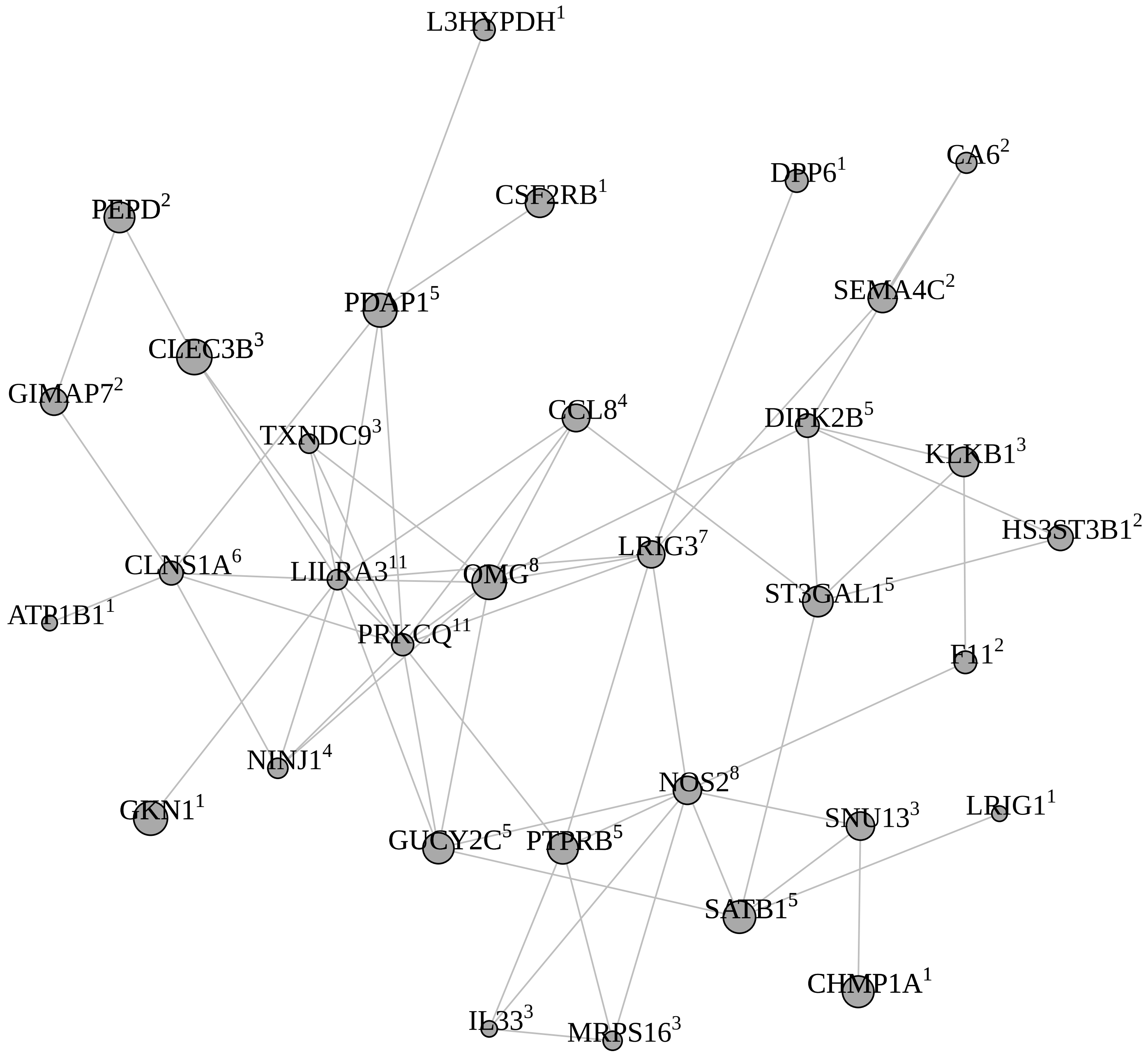

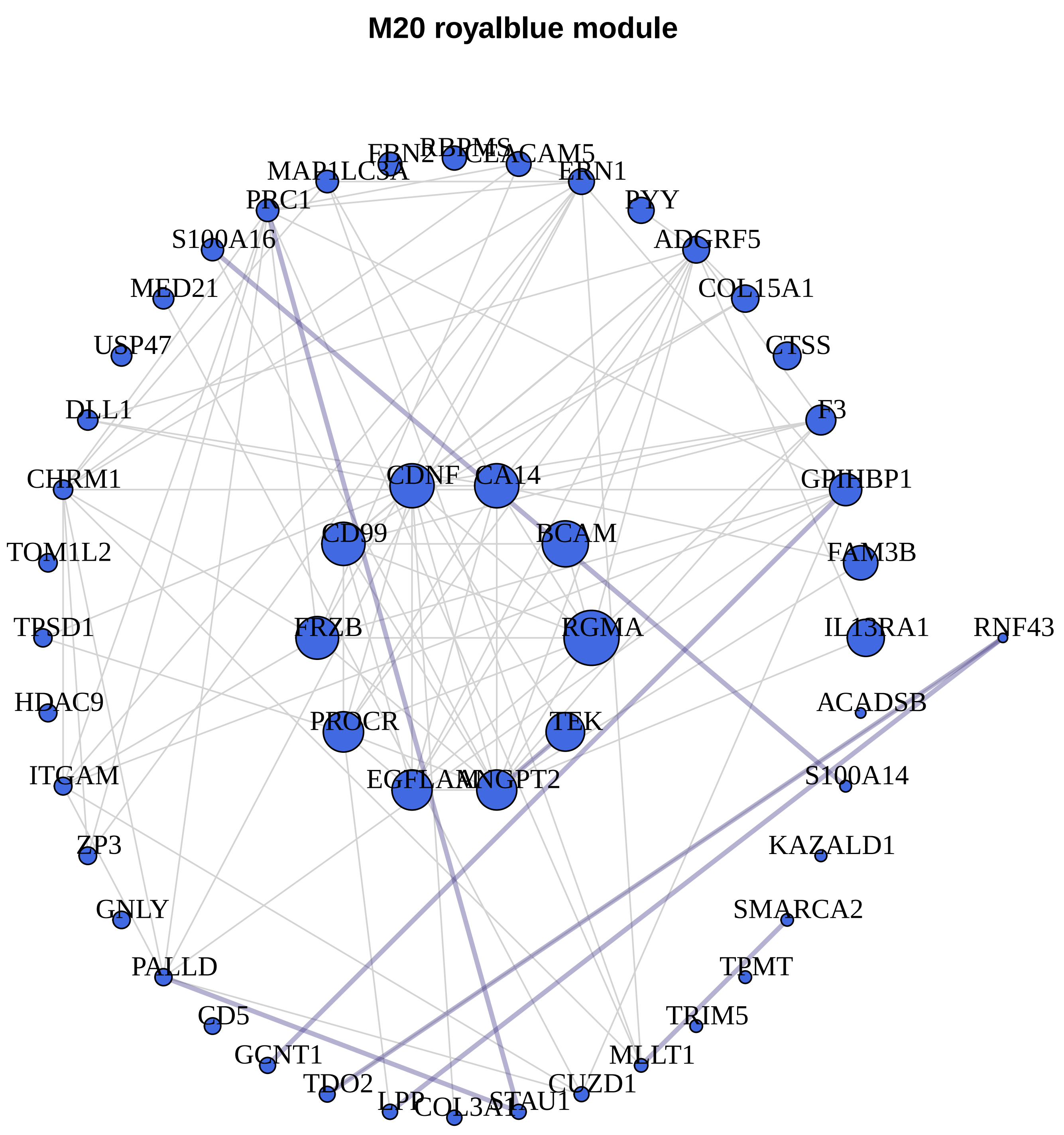
M20 royalblue module
ERN2
RBPMS
CEACAM5
MAP1LC3A
ERN1
PRC1
PYY
S100A16
ADGRF5
MED21
COL15A1
USP47
CTSS
DLL1
F3
CHRM1
CDNF
CA14
GPIHBP1
TOM1L2
CD99
BCAM
FAM3B
TPSD1
FRZB
RGMA
IL13RA1
RNF43
HDAC9
ACADSB
PROCR
TEK
ITGAM
EGFLAM
ANGPT2
S100A14
ZP3
KAZALD1
GNLY
SMARCA2
PALLD
TPMT
CD5
TRIM5
GCNT1
MLLT1
TDO2
CUZD1
LPP
COL3A1
STAU1
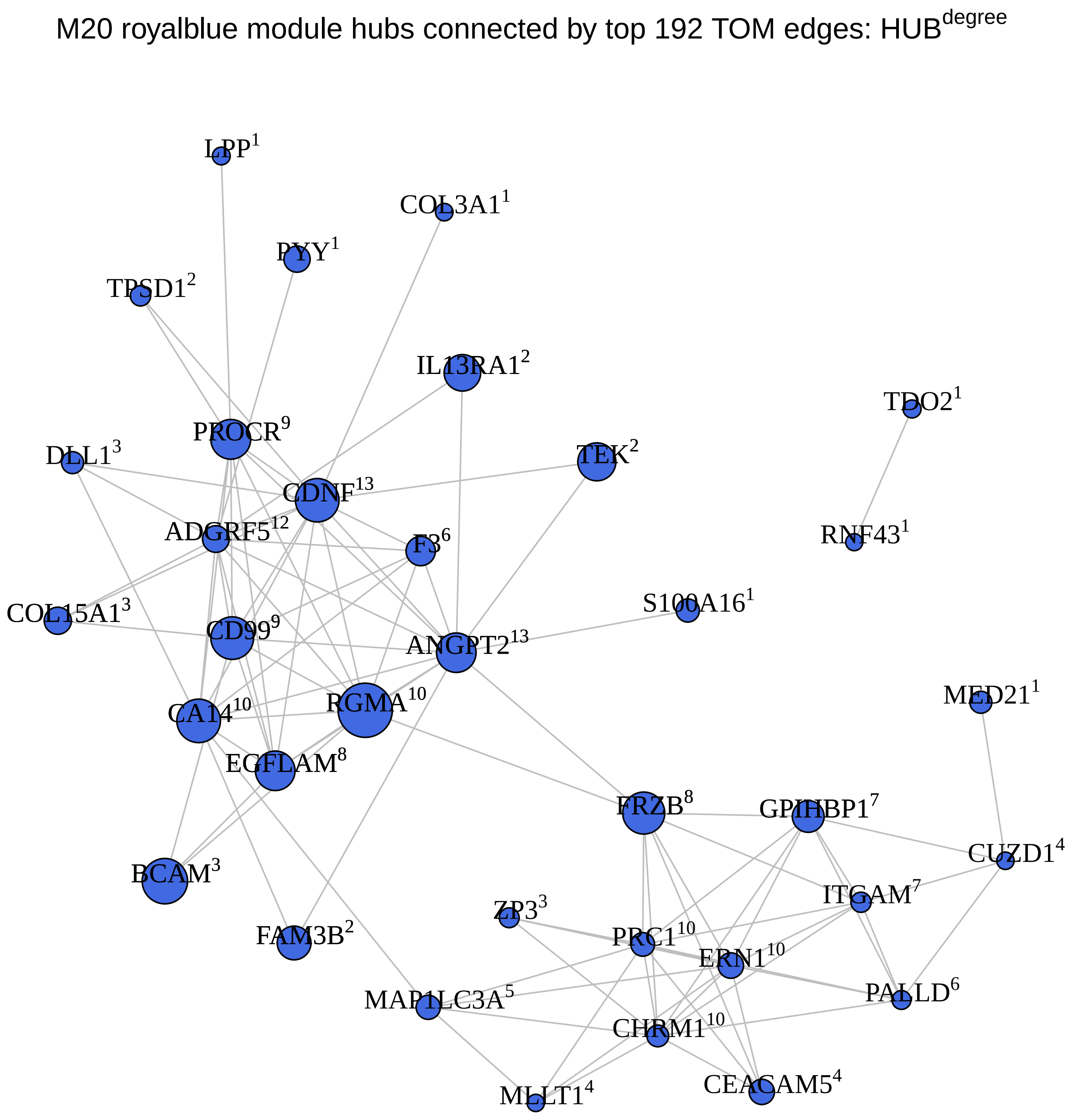
M20 royalblue module hubs connected by top 192 TOM edges: HUB[degree]
LPP[1]
COL3A1[1]
PYY[1]
TPSD1[2]
IL13RA1[2]
TDO2[1]
PROCR[9]
DLL1[3]
TEK[2]
CDNF[13]
ADGRF5[12]
F3[6]
RNF43[1]
S100A16[1]
COL15A1[3]
CD99[9]
ANGPT2[13]
MED21[1]
CA14[10]
RGMA[10]
EGFLAM[8]
FRZB[8]
GPIHBP1[7]
CUZD1[4]
BCAM[3]
ITGAM[7]
ZP3[3]
FAM3B[2]
PRC1[10]
ERN1[10]
PALLD[6]
MAP1LC3A[5]
CHRM1[10]
CEACAM5[4]
MLLT1[4]

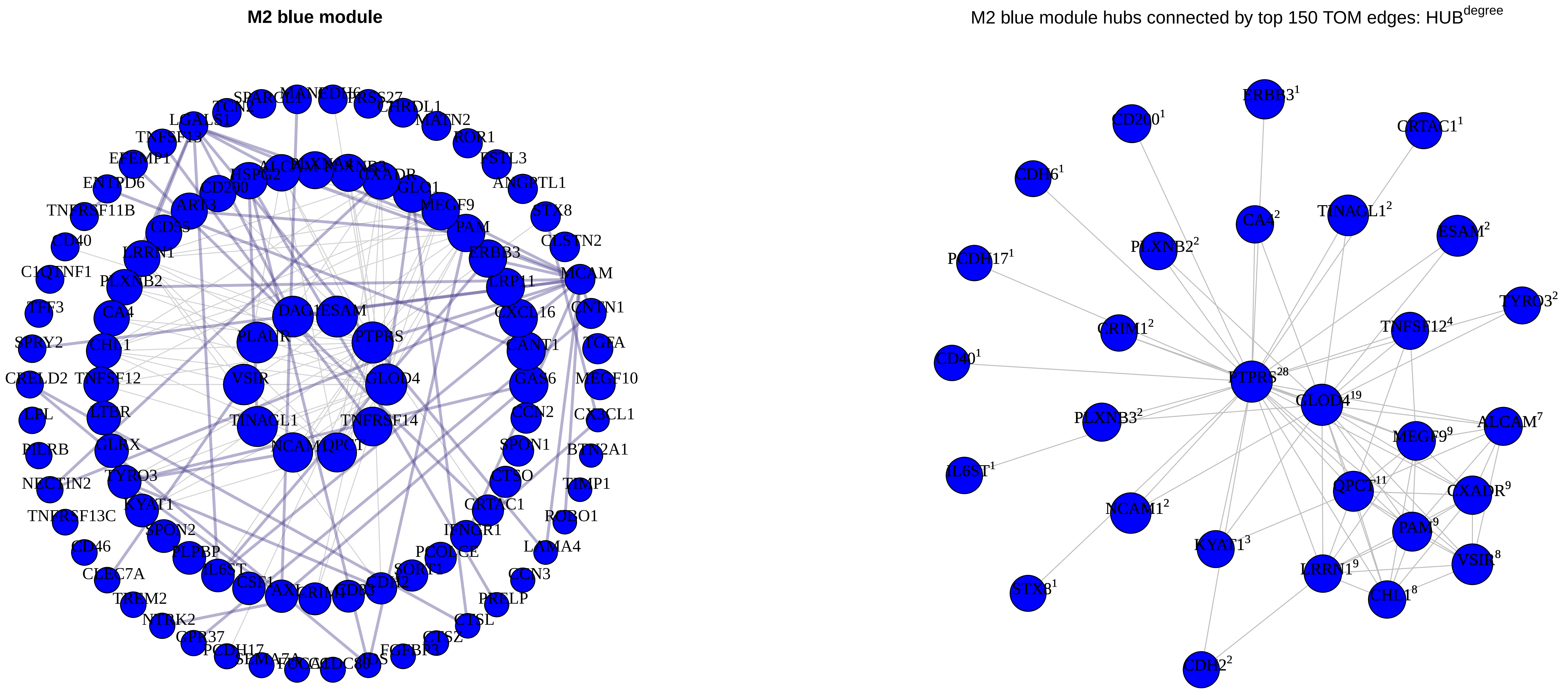
M2 blue module
M2 blue module hubs connected by top 150 TOM edges: HUB^degree
ERBB3^1
CD200^1
CRTAC1^1
CDH6^1
CA4^2
TINAGL1^2
ESAM^2
PLXNB2^2
PCDH17^1
TYRO3^2
CRIM1^2
TNFSF12^4
CD40^1
PTPRS^28
GLOD4^19
PLXNB3^2
ALCAM^7
MEGF9^9
IL6ST^1
QPCT^11
CXADR^9
NCAM1^2
PAM^9
KYAT1^3
VSIR^8
LRRN1^9
CHL1^8
STX8^1
CDH2^2

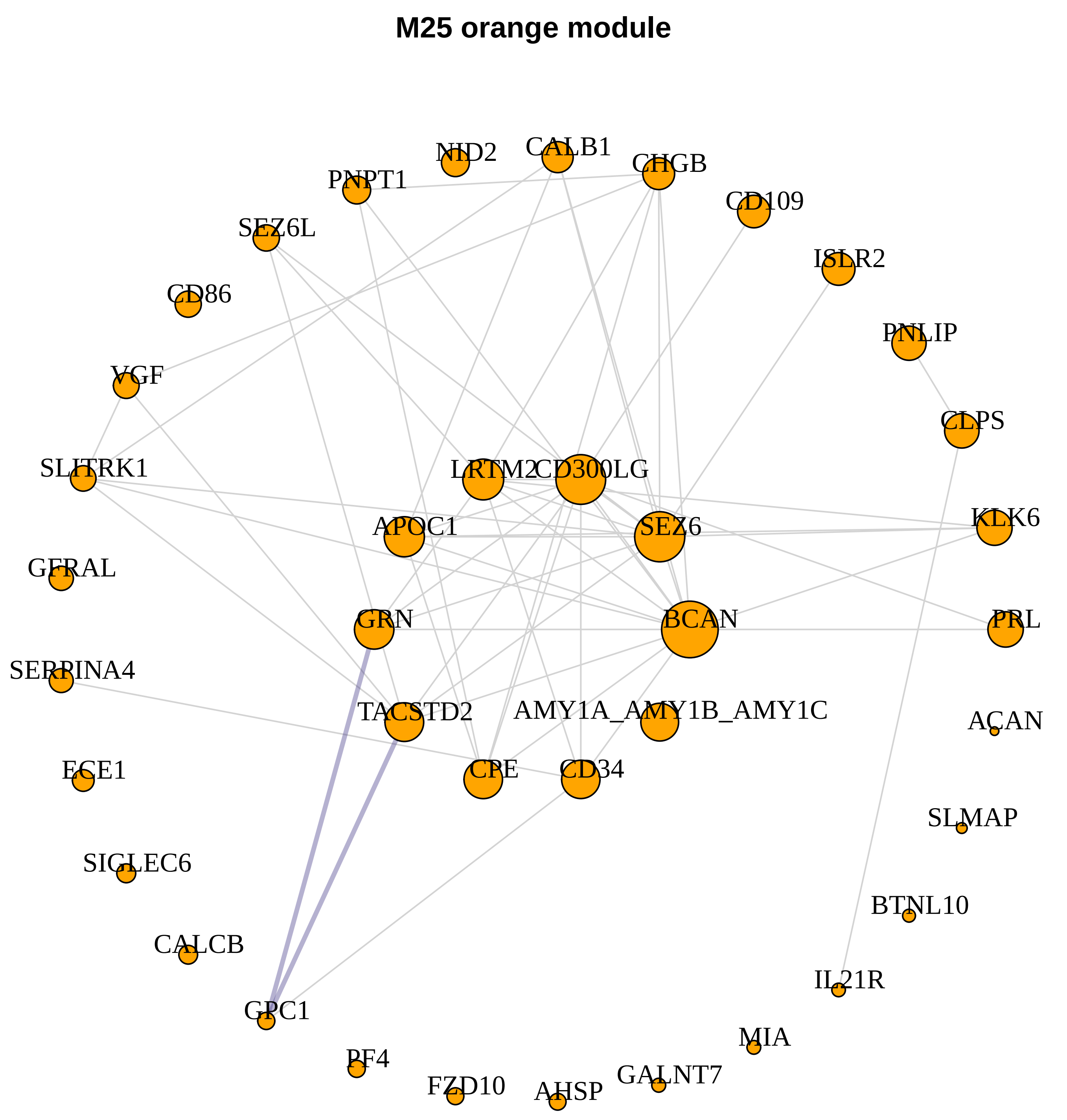
M25 orange module
NID2
CALB1
CHGB
PNPT1
CD109
SEZ6L
ISLR2
CD86
PNLIP
VGF
CLPS
SLITRK1
LRTM2
CD300LG
KLK6
APOC1
SEZ6
GFRAL
GRN
BCAN
PRL
SERPINA4
TACSTD2
AMY1A_AMY1B_AMY1C
ACAN
ECE1
CPE
CD34
SLMAP
SIGLEC6
BTNL10
CALCB
IL21R
GPC1
MIA
PF4
FZD10
AHSP
GALNT7

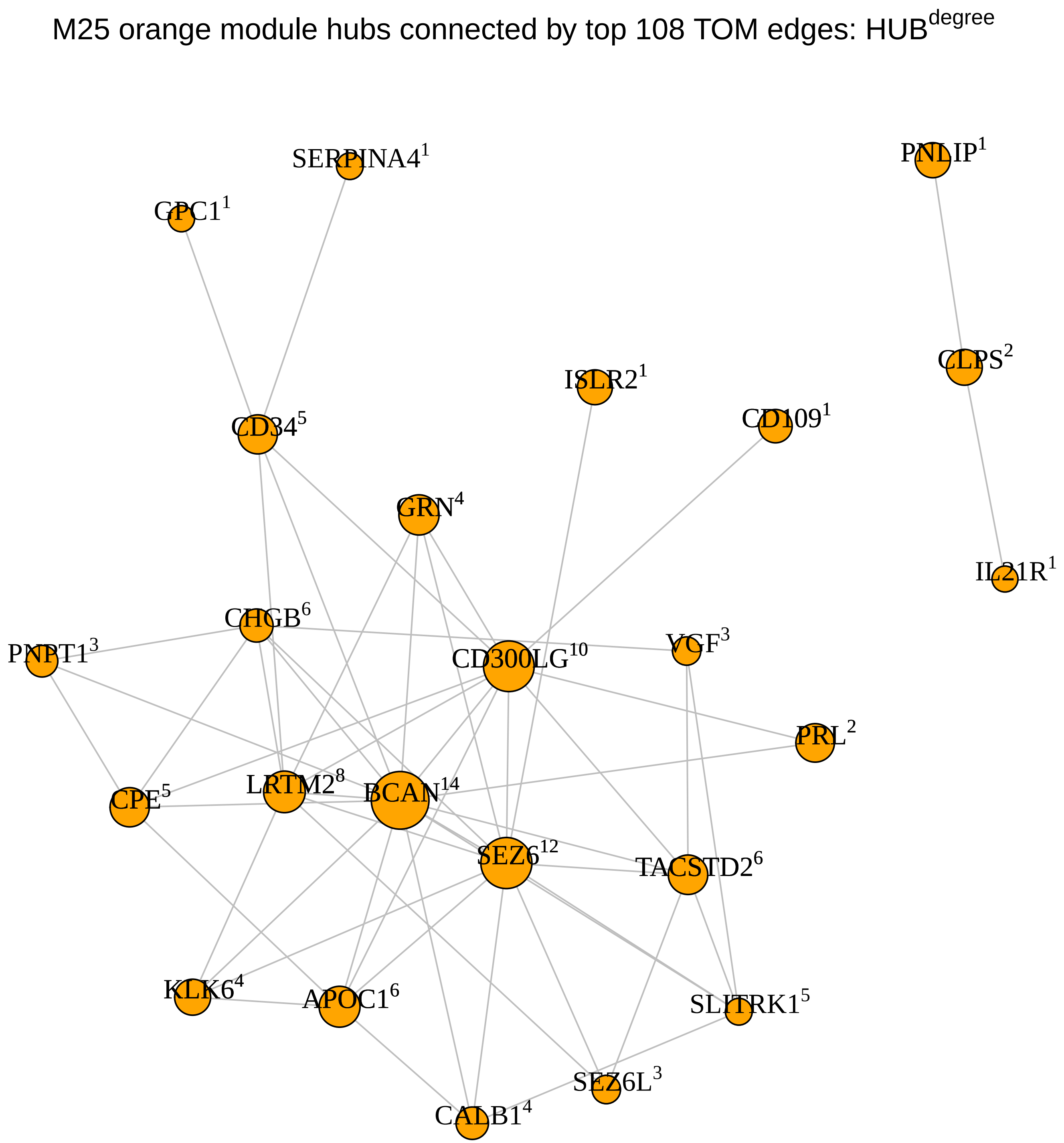
M25 orange module hubs connected by top 108 TOM edges: HUB^degree
SERPINA4 1
PNLIP 1
GPC1 1
CLPS 2
ISLR2 1
CD34 5
CD109 1
GRN 4
IL21R 1
CHGB 6
PNPT1 3
CD300LG 10
VGF 3
PRL 2
CPE 5
LRTM2 8
BCAN 14
SEZ6 12
TACSTD2 6
KLK6 4
APOC1 6
SLITRK1 5
SEZ6L 3
CALB1 4

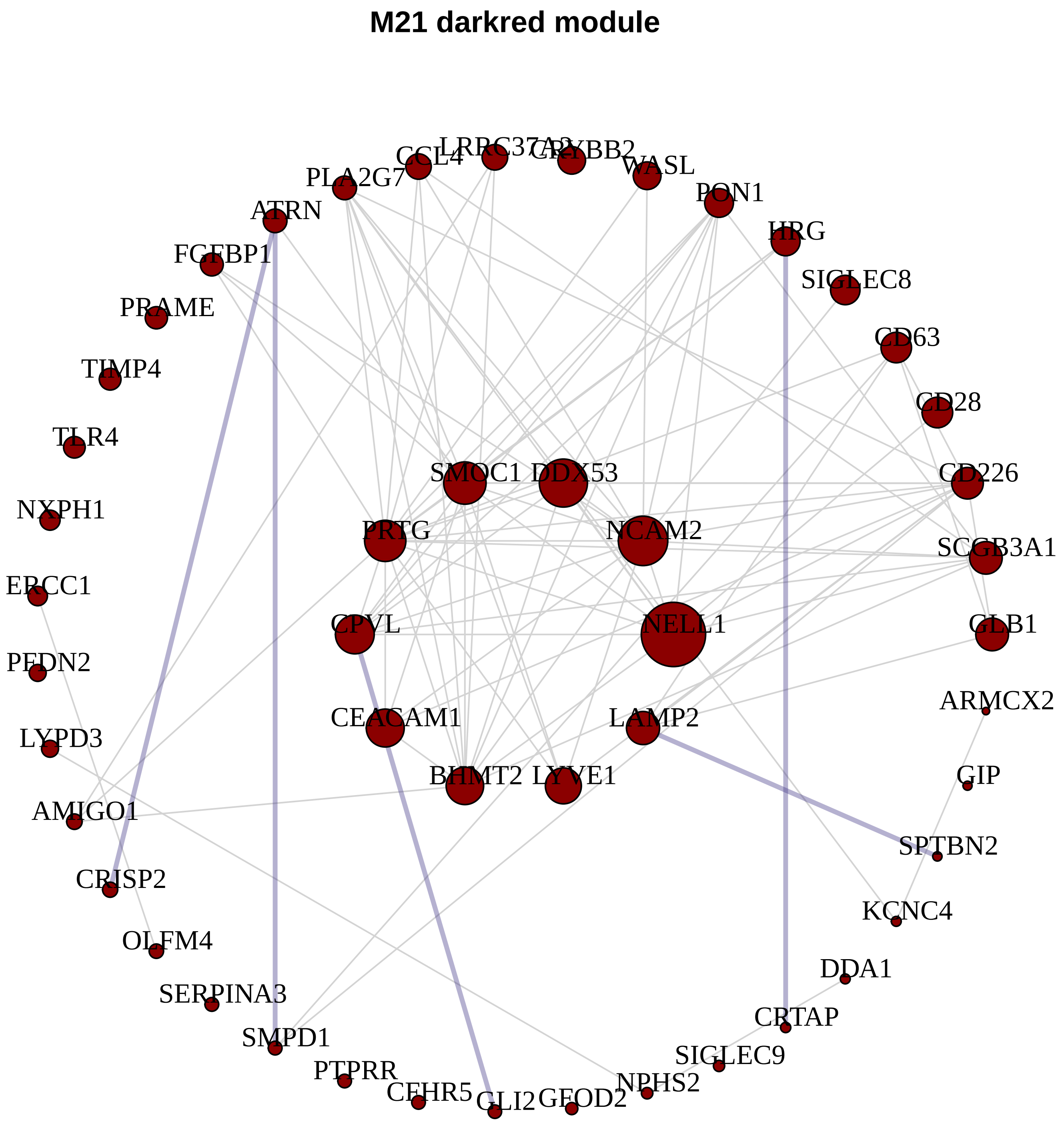
M21 darkred module
PLA2G7
CCL4
LRRC37A2
CRYBB2
WASL
PON1
ATRN
HRG
FCGBP1
SIGLEC8
PRAME
CD63
TIMP4
CD28
TLR4
SMOC1
DDX53
CD226
NXPH1
PRTG
NCAM2
SCGB3A1
ERCC1
CPVL
NELL1
ARMCX2
LYPD3
CEACAM1
LAMP2
GIP
AMIGO1
BHMT2
LYVE1
SPTBN2
CRISP2
KCNC4
OLFM4
DDA1
SERPINA3
CRTAP
SMPD1
SIGLEC9
PTPRR
CFHR5
GLI2
GFOD2
NPHS2

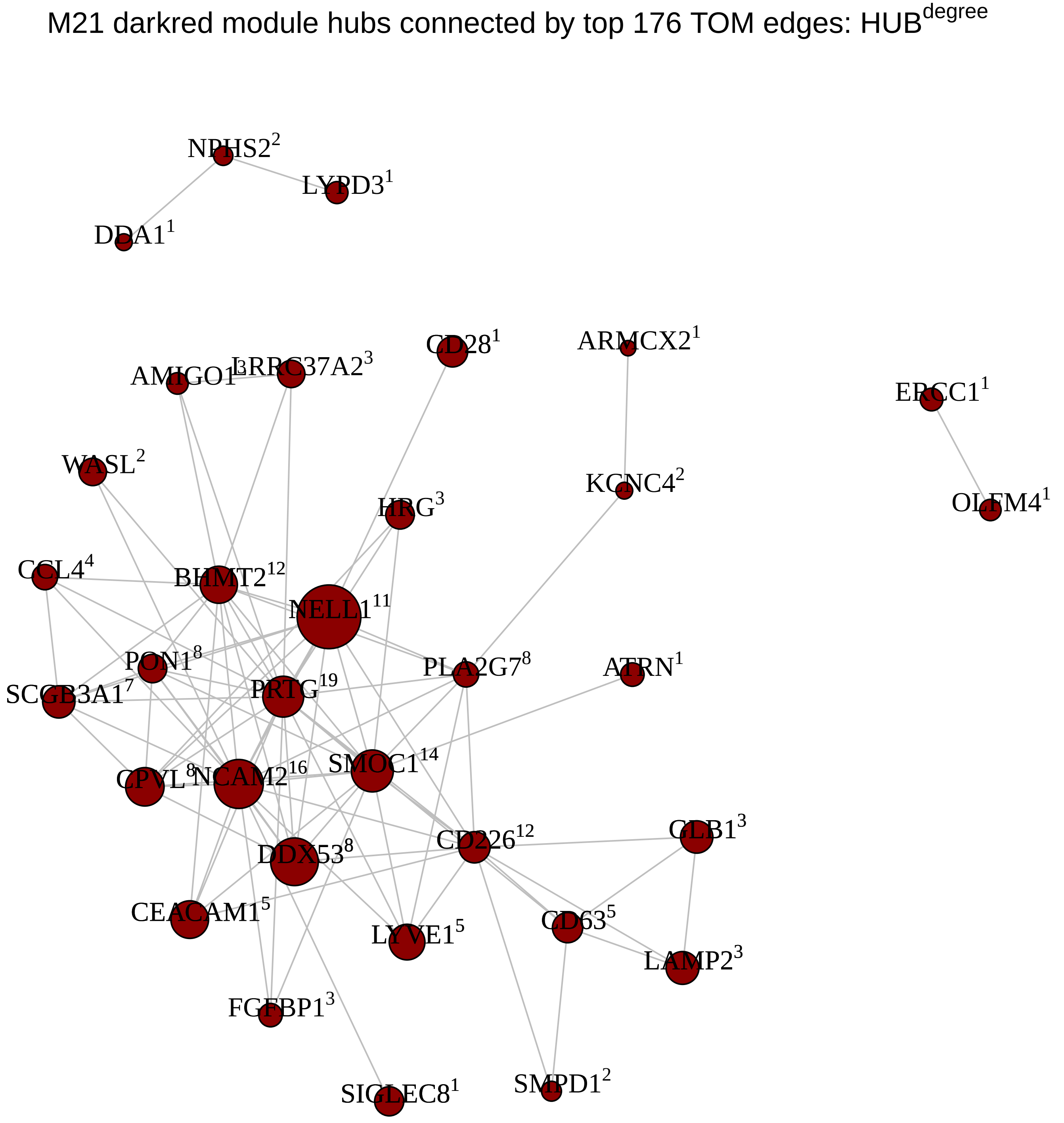
M21 darkred module hubs connected by top 176 TOM edges: HUB^degree
NPHS2^2
LYPD3^1
DDA1^1
CD28^1
ARMCX2^1
LRRC37A2^3
ERCC1^1
WASL^2
KCNC4^2
HRG^3
OLFM4^1
CCL4^4
BHMT2^12
NELL1^11
PON1^8
PLA2G7^8
ATRN^1
SCGB3A1^7
PRTG^19
SMOC1^14
NCAM2^16
DDX53^8
CD226^12
CEACAM1^5
LYVE1^5
CD63^5
LAMP2^3
FCGBP1^3
SIGLEC8^1
SMPD1^2

**M8 pink module**

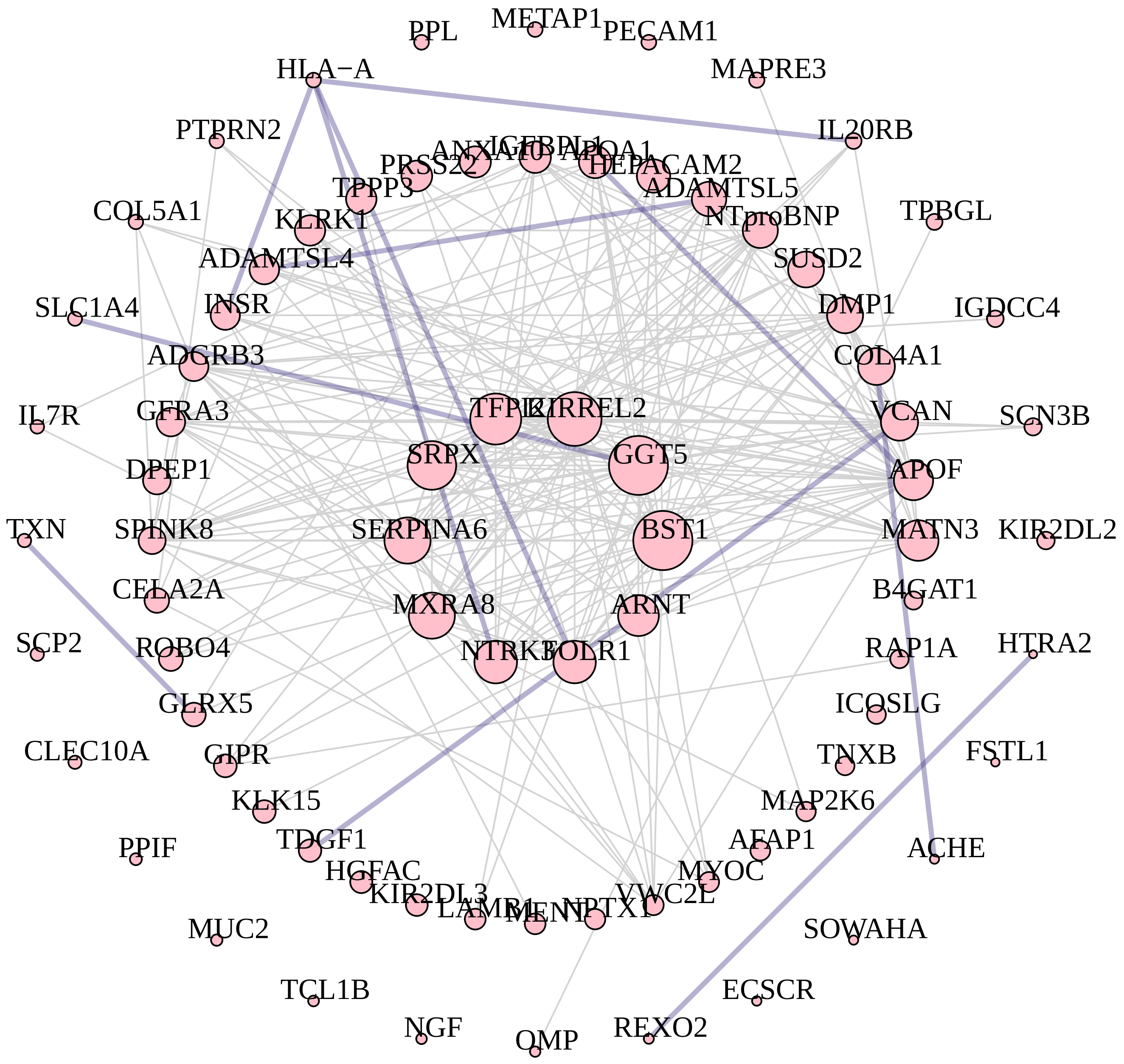


M8 pink module hubs connected by top 468 TOM edges: HUB$^{degree}$

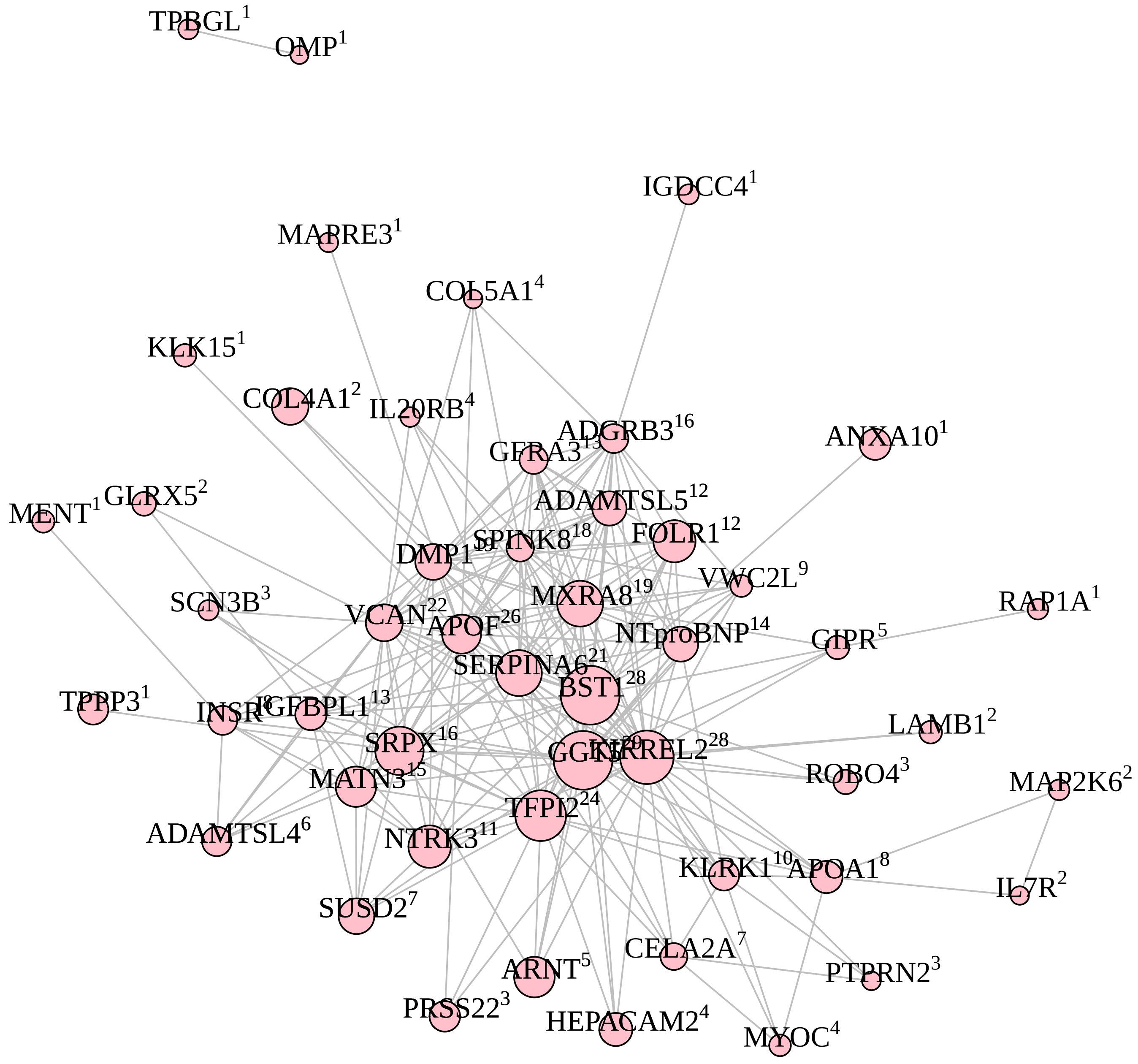

**M19 lightyellow module**

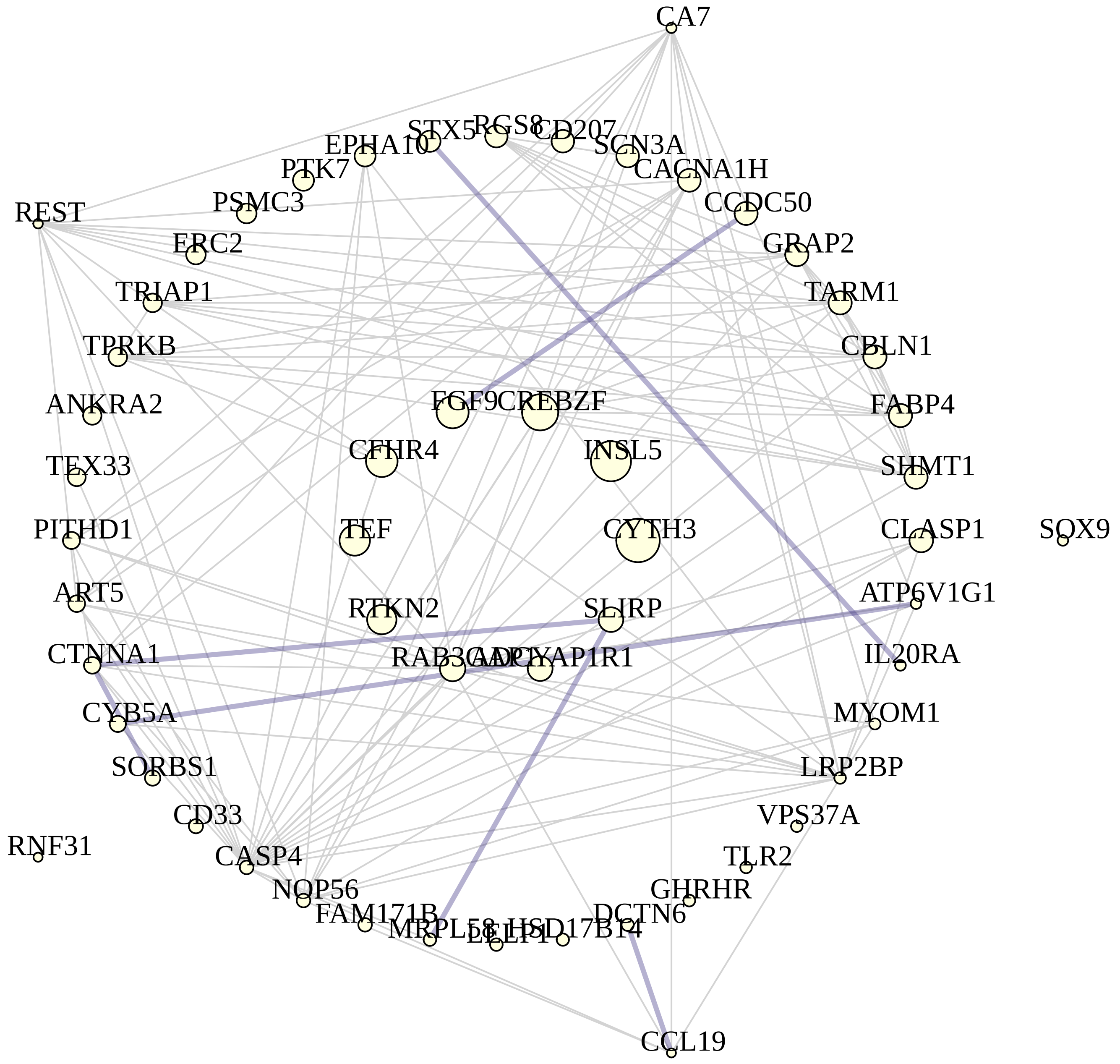


M19 lightyellow module hubs connected by top 224 TOM edges: $HUB^{degree}$

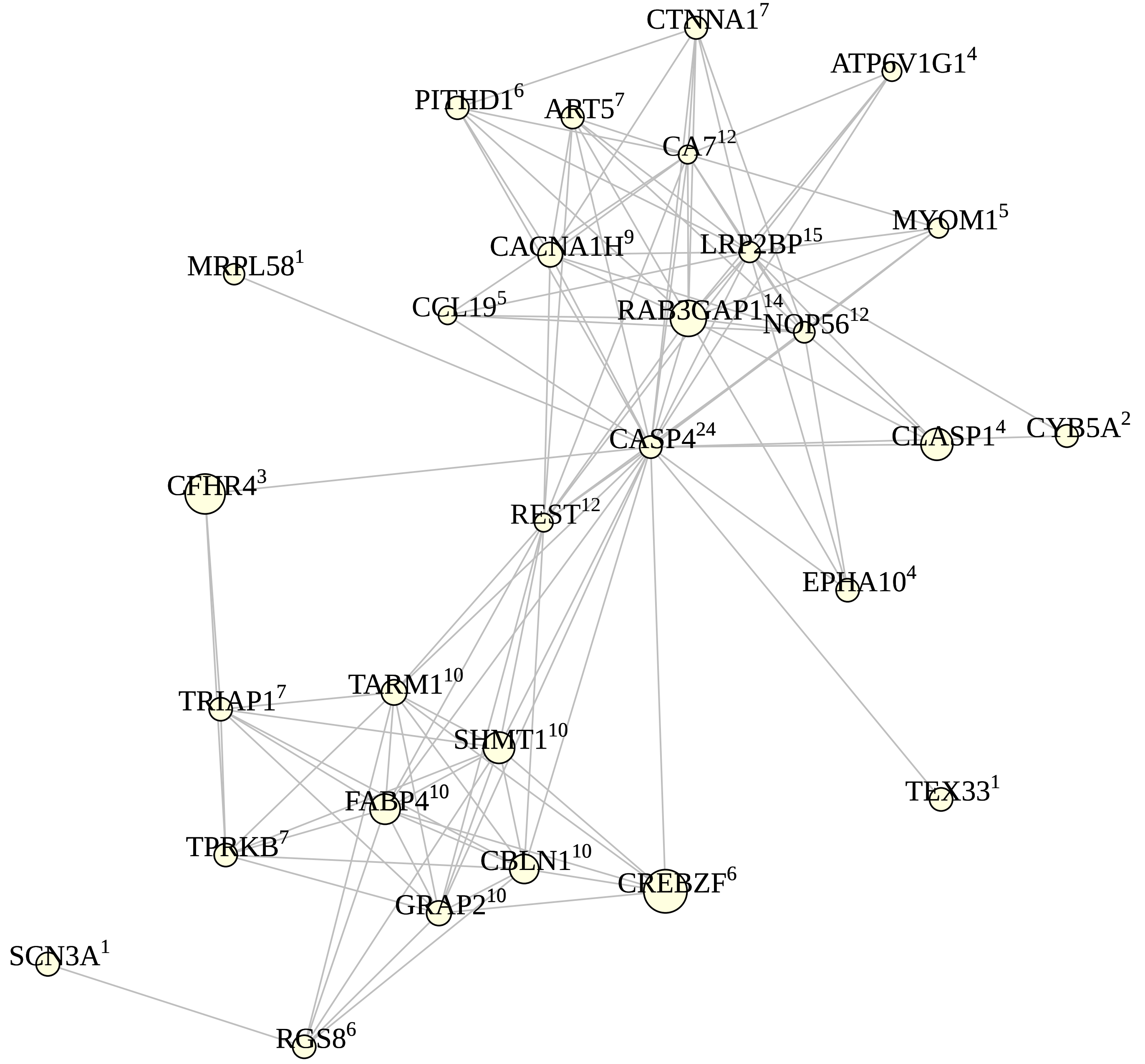

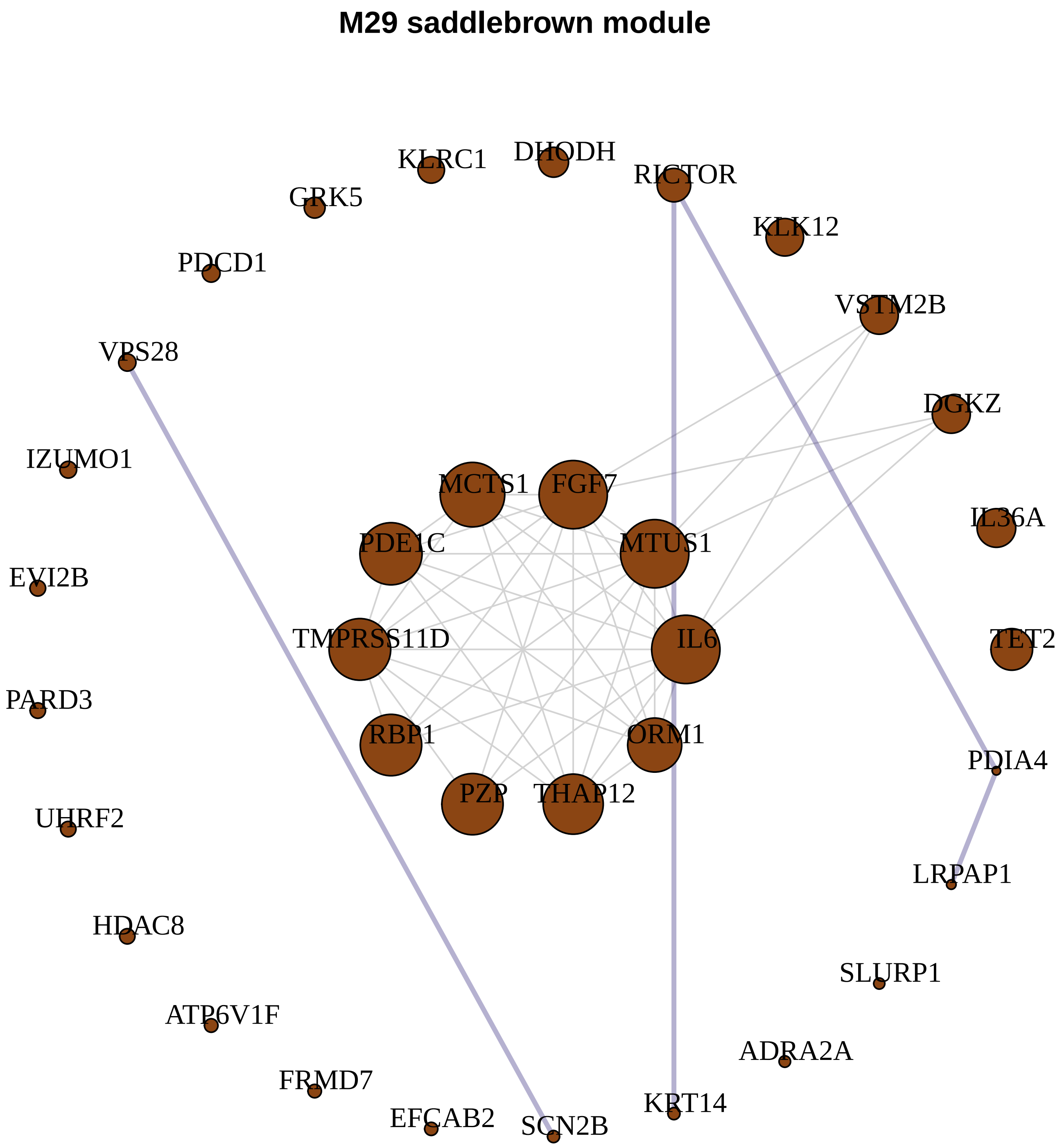
M29 saddlebrown module
KLRC1
DHODH
RICTOR
GRK5
KLK12
PDCD1
VSTM2B
VPS28
DGKZ
IZUMO1
MCTS1
FGF7
IL36A
PDE1C
MTUS1
EVI2B
TMPRSS11D
IL6
TET2
PARD3
RBP1
ORM1
PDIA4
PZP
THAP12
UHRF2
LRPAP1
HDAC8
SLURP1
ATP6V1F
ADRA2A
FRMD7
KRT14
EFCAB2
SCN2B

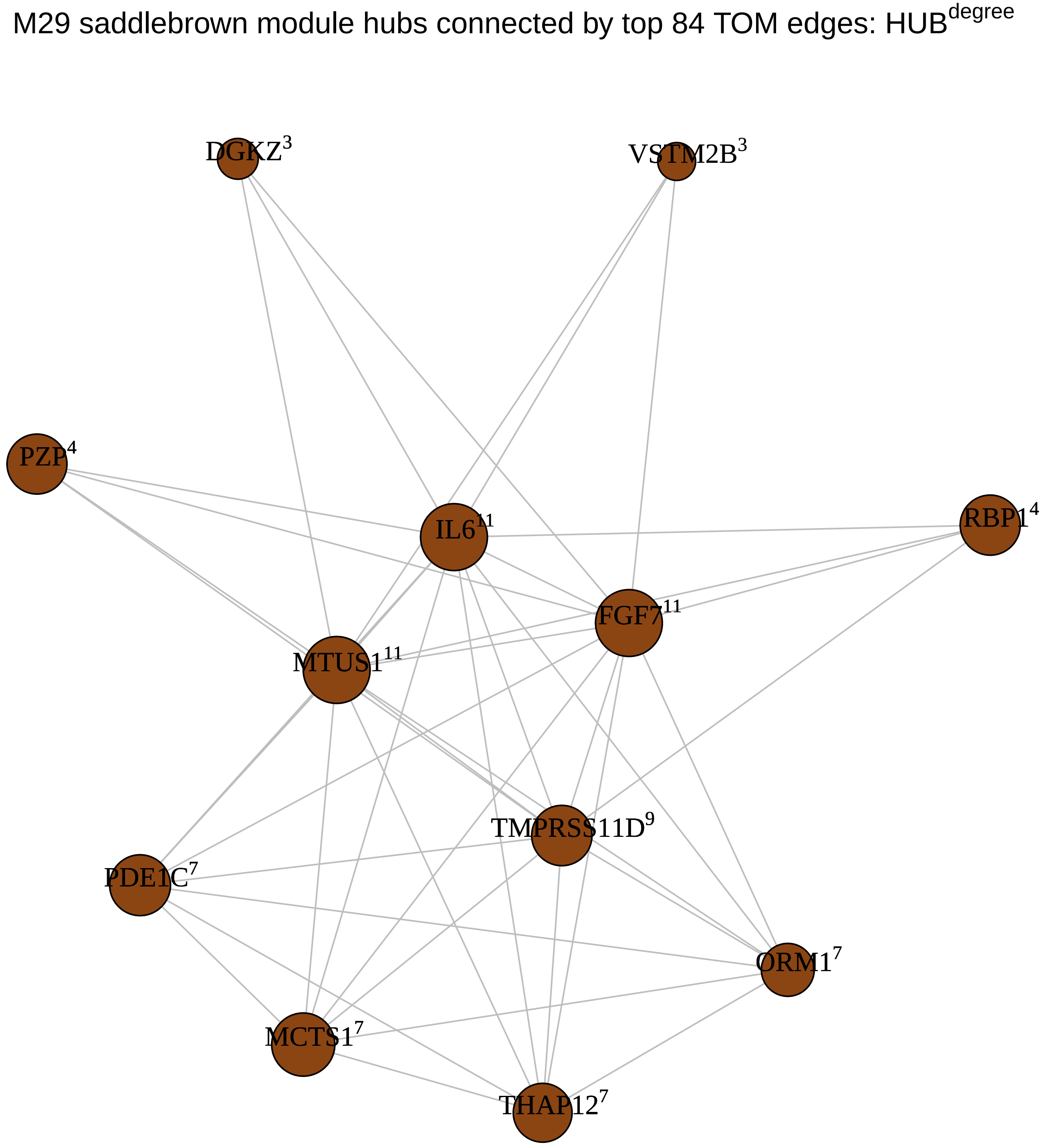
M29 saddlebrown module hubs connected by top 84 TOM edges: HUB^degree
DGKZ^3
VSTM2B^3
PZP^4
IL6^11
RBP1^4
FGF7^11
MTUS1^11
TMPRSS11D^9
PDE1C^7
ORM1^7
MCTS1^7
THAP12^7

**M17 grey60 module**

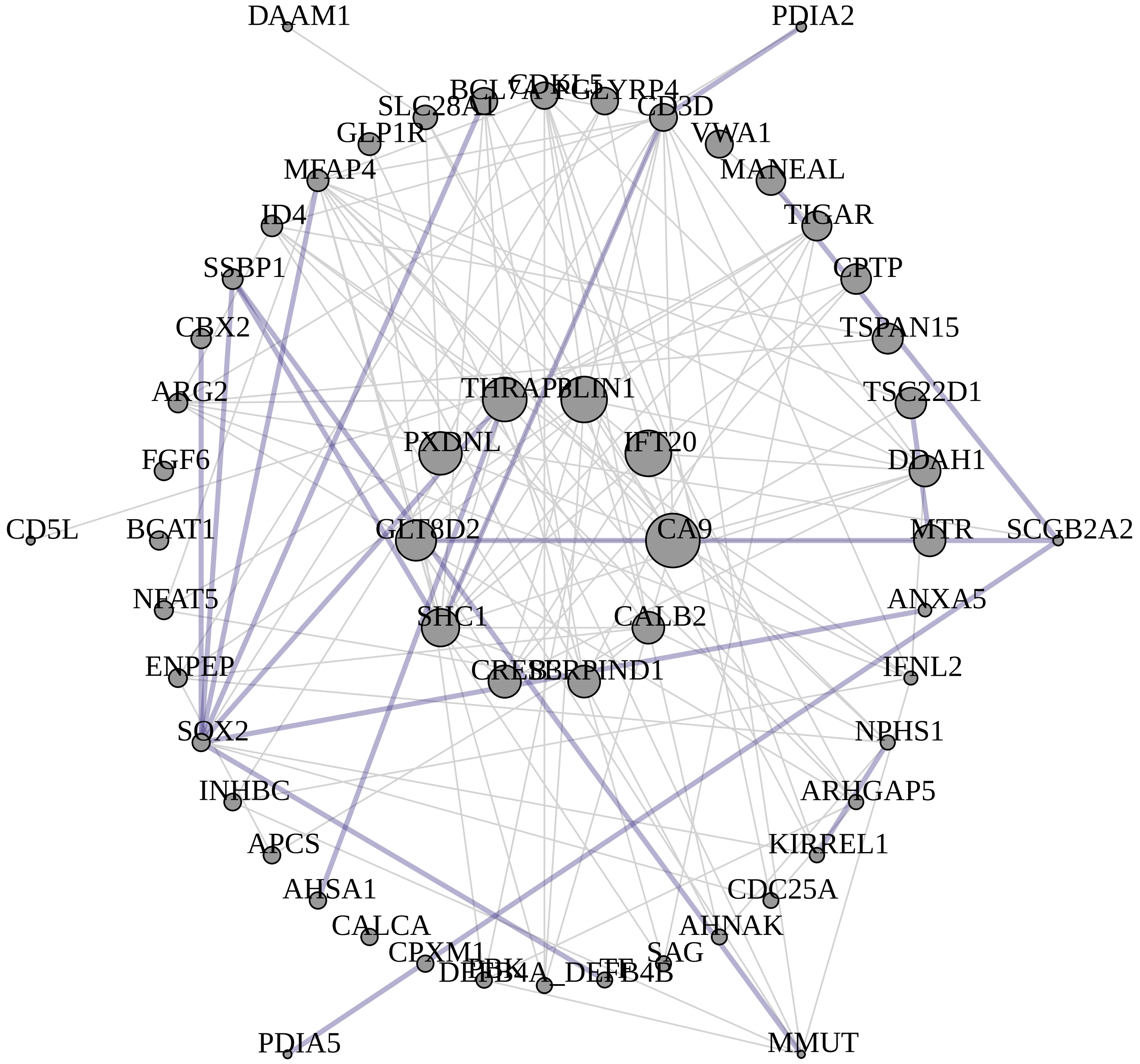


M17 grey60 module hubs connected by top 234 TOM edges: HUB$^{degree}$

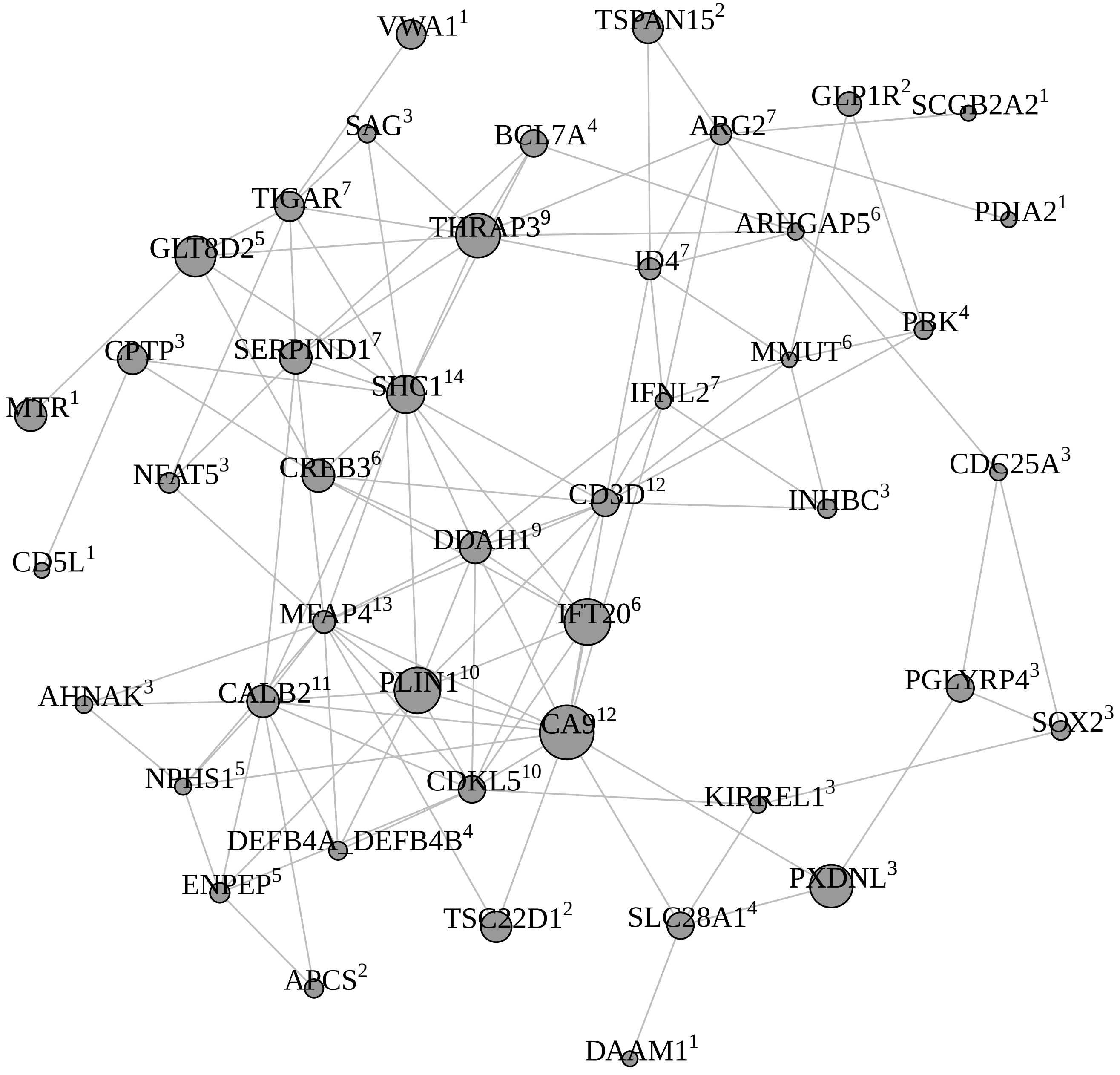

**M9 magenta module**

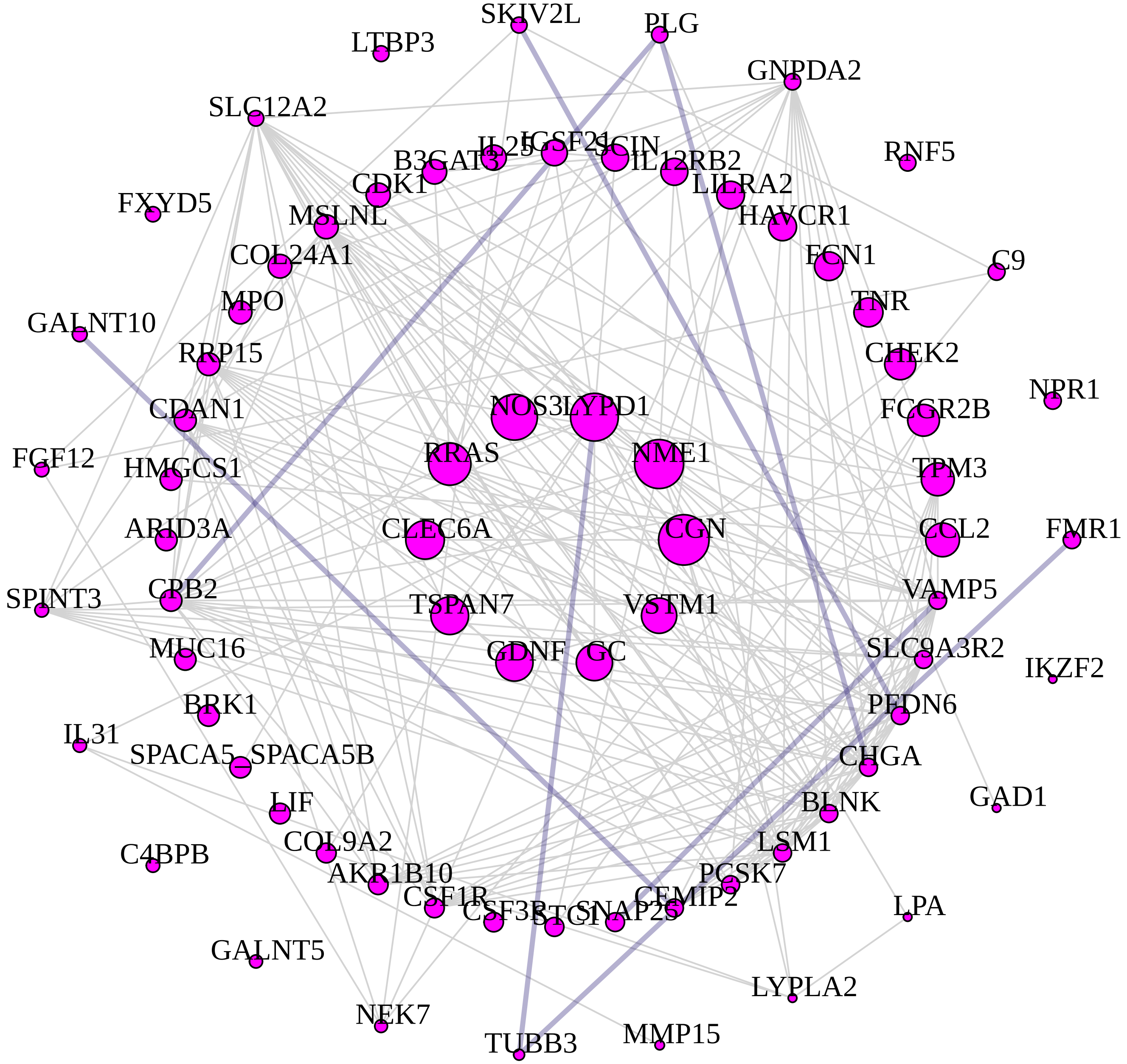


M9 magenta module hubs connected by top 408 TOM edges: HUB$^{degree}$

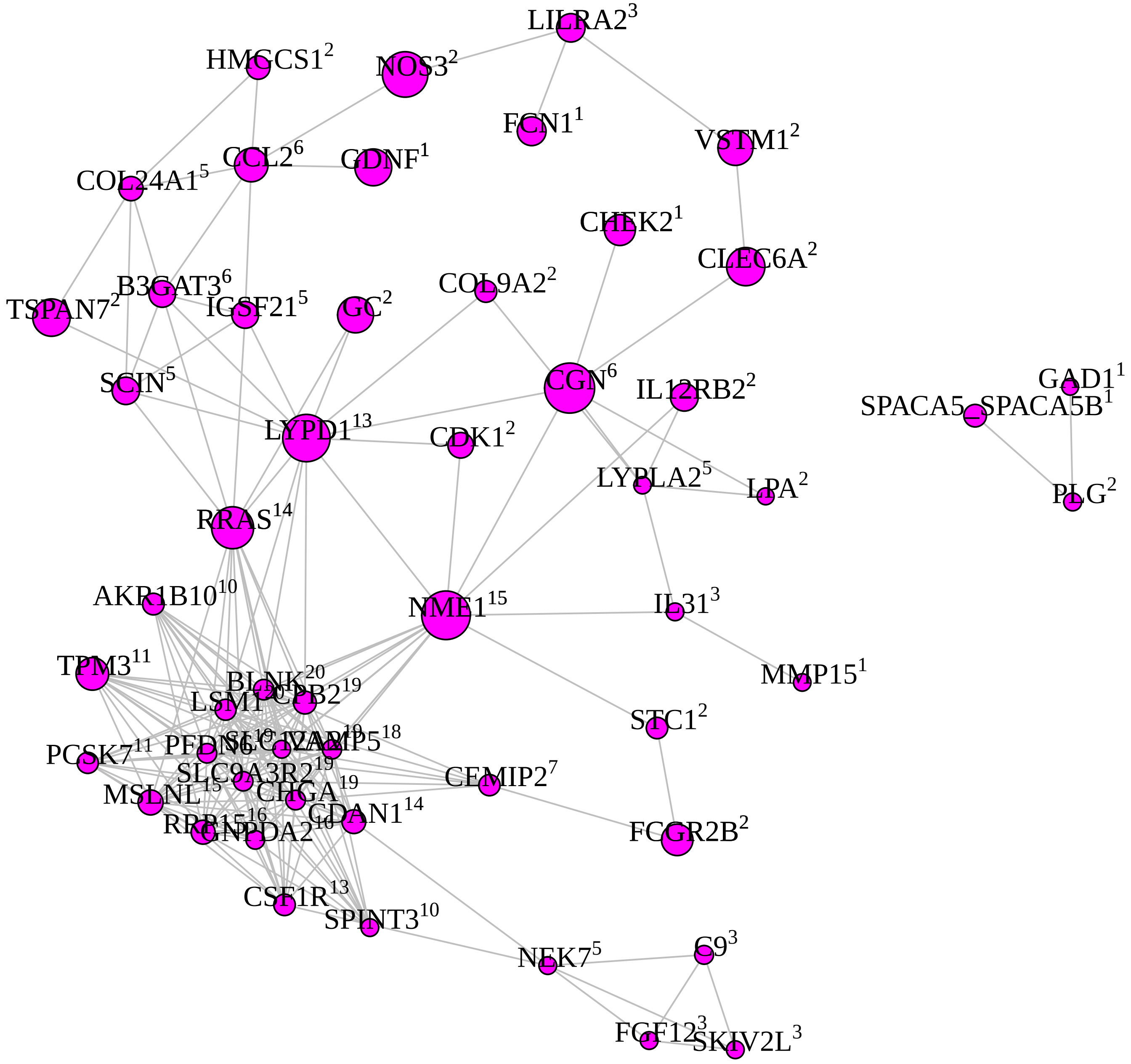

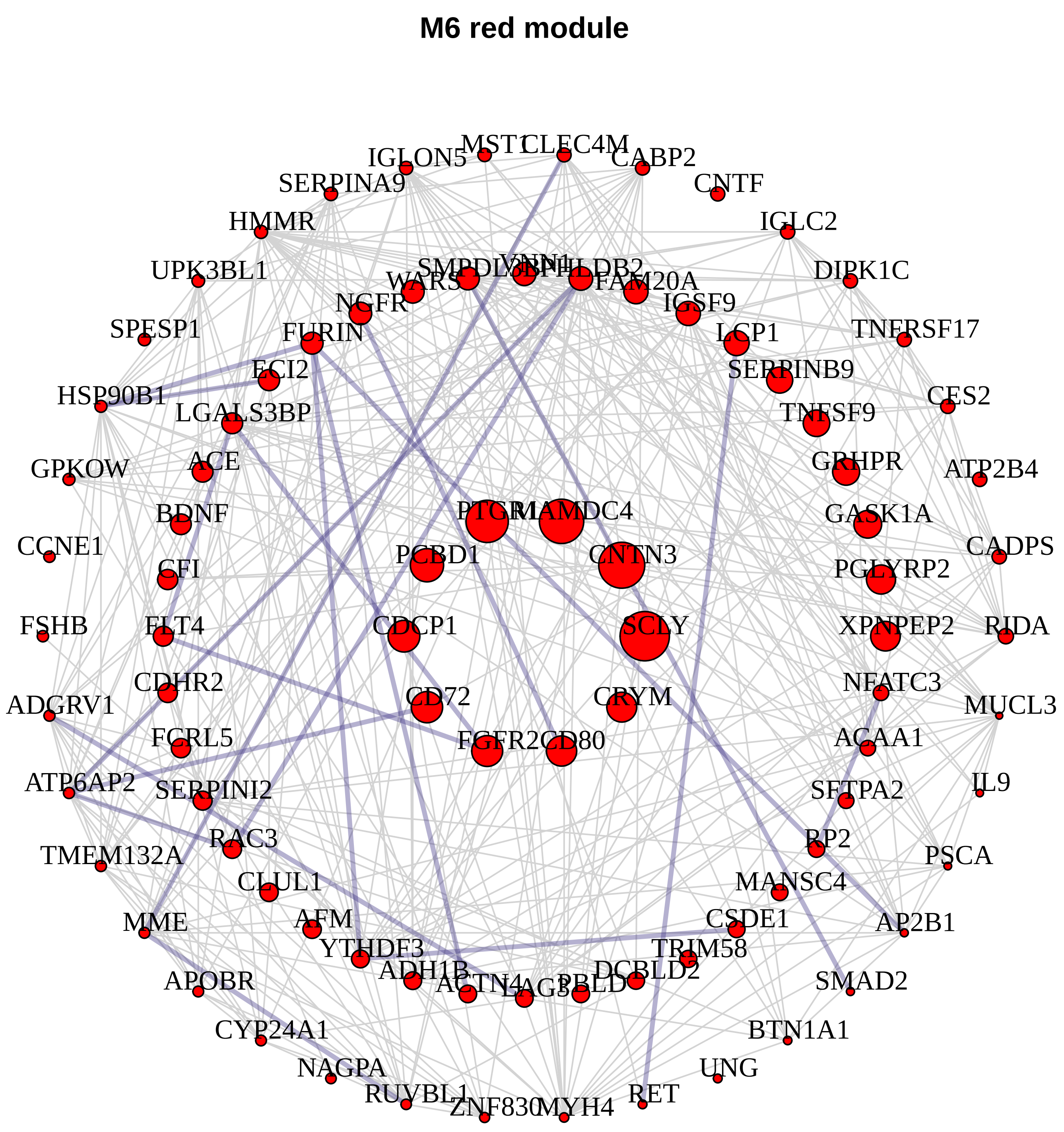

M6 red module
IGLON5
MST1
CLEC4M
CABP2
SERPINA9
CNTF
HMMR
IGLC2
UPK3BL1
DIPK1C
WARS
SMPDL3B
VNN1
PHLDB2
FAM20A
NGFR
IGSF9
SPESP1
FURIN
TNFRSF17
LCP1
ECI2
SERPINB9
HSP90B1
CES2
LGALS3BP
TNFSF9
GPKOW
ACE
GRHPR
ATP2B4
BDNF
PTGR1
MAMDC4
GASK1A
CCNE1
PCBD1
CNTN3
CADPS
CFI
PGLYRP2
FSHB
FLT4
CDCP1
SCLY
XPNPEP2
RIDA
CDHR2
NFATC3
ADGRV1
CD72
CRYM
MUCL3
FCRL5
FGFR2
CD80
ACAA1
ATP6AP2
SERPINI2
SFTPA2
IL9
RAC3
RP2
TMEM132A
PSCA
CLUL1
MANSC4
MME
AFM
CSDE1
AP2B1
YTHDF3
TRIM58
ADH1B
ACTN4
LAG3
PBLD
DCBLD2
APOBR
SMAD2
CYP24A1
BTN1A1
NAGPA
UNG
RUVBL1
RET
ZNF830
MYH4


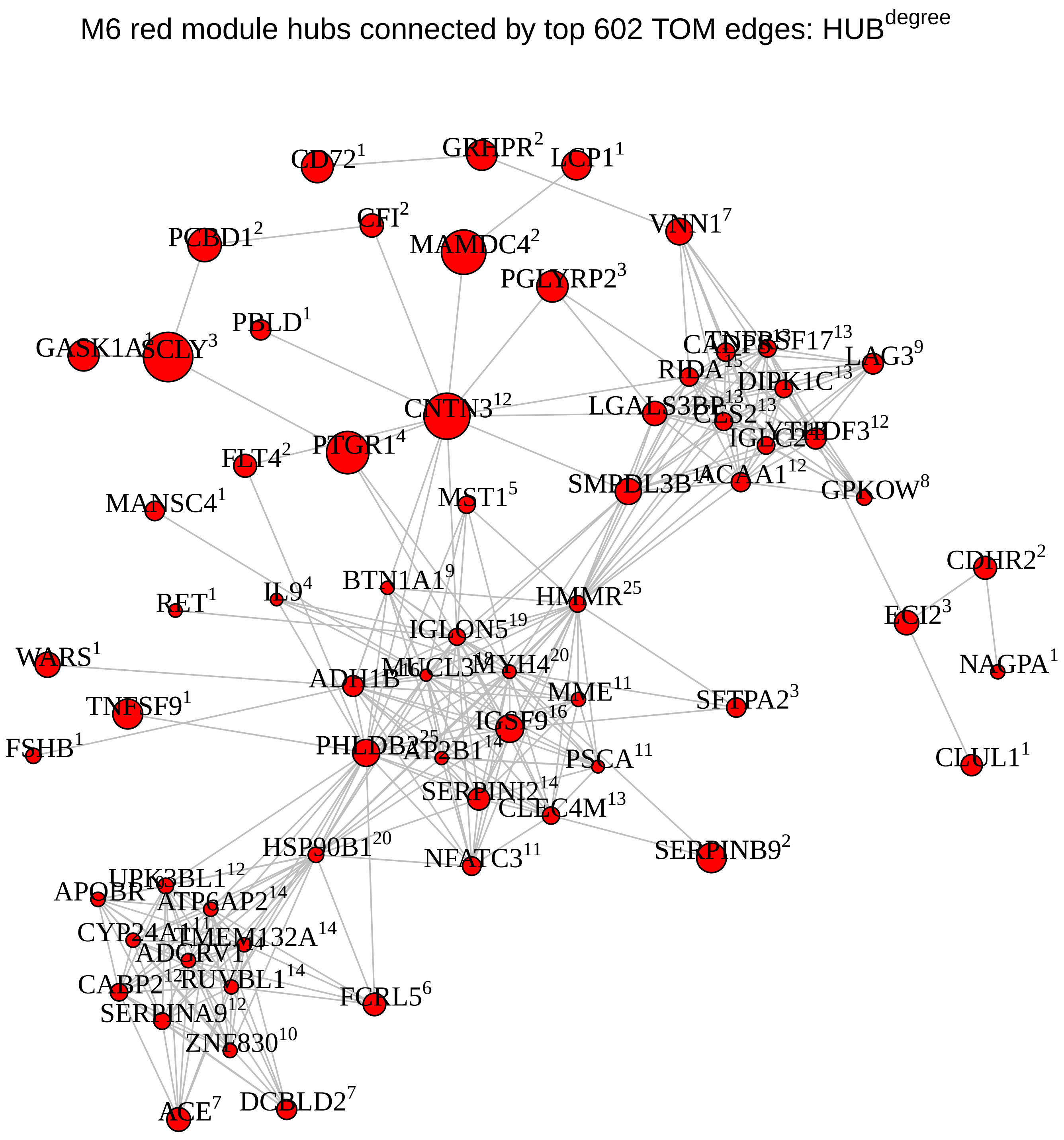

M6 red module hubs connected by top 602 TOM edges: HUB^degree

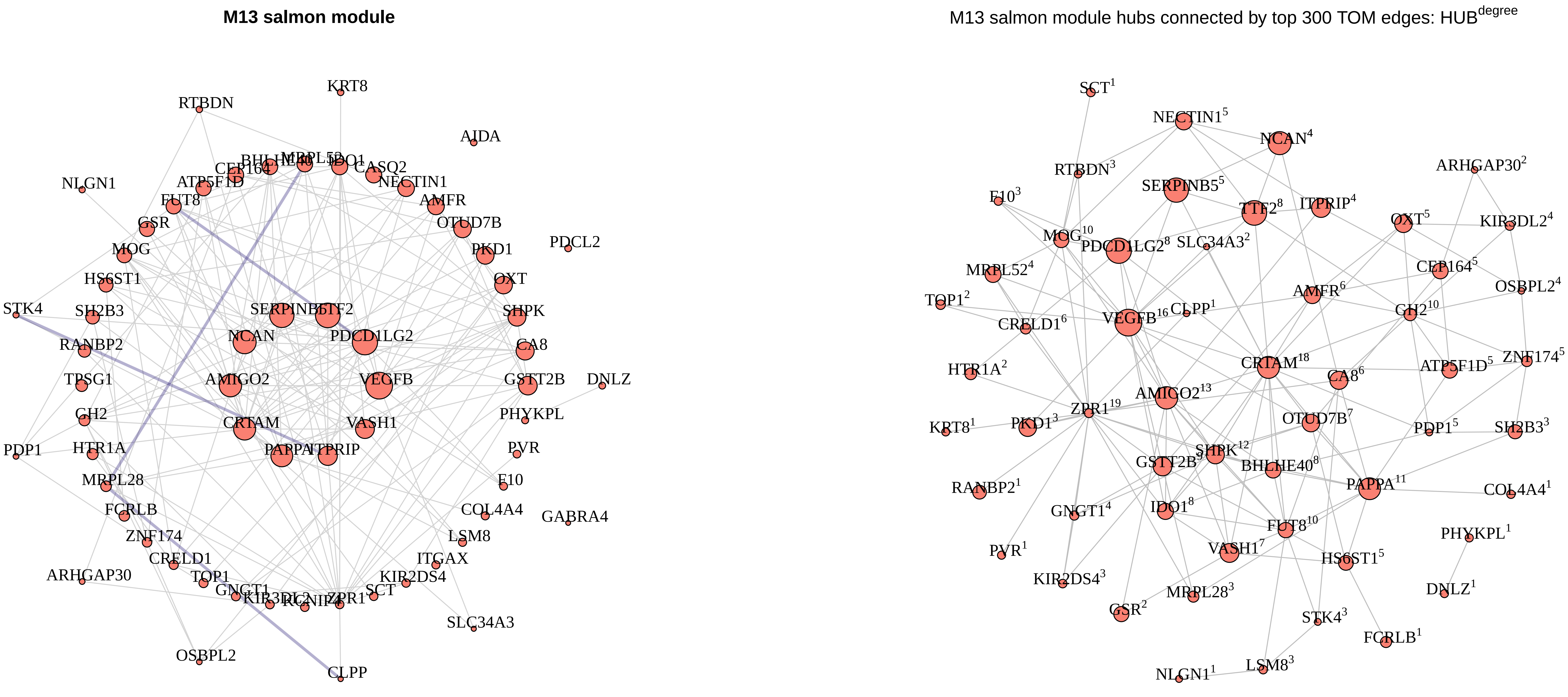

M13 salmon module
KRT8
RTBDN
AIDA
CASQ2
CEP164
NLGN1
ATP5F1D
NECTIN1
FUT8
AMFR
GSR
OTUD7B
MOG
PKD1
PDCL2
HS6ST1
OXT
STK4
SH2B3
SERPINB5
TTF2
SHPK
NCAN
PDCD1LG2
RANBP2
CA8
TPSG1
AMIGO2
VEGFB
GSTT2B
DNLZ
GH2
CRTAM
VASH1
PHYKPL
PDP1
HTR1A
PAPPA
ITPRIP
PVR
MRPL28
F10
FCRLB
COL4A4
GABRA4
ZNF174
LSM8
CRELD1
ITGAX
ARHGAP30
TOP1
KIR2DS4
GNGT1
ZPR1
SCT
SLC34A3
OSBPL2
CLPP
M13 salmon module hubs connected by top 300 TOM edges: HUB$^{degree}$
SCT[1]
NECTIN1[5]
NCAN[4]
RTBDN[3]
ARHGAP30[2]
F10[3]
SERPINB5[5]
TTF2[8]
ITPRIP[4]
OXT[5]
KIR3DL2[4]
MOG[10]
PDCD1LG2[8]
SLC34A3[2]
MRPL52[4]
CEP164[5]
AMFR[6]
OSBPL2[4]
TOP1[2]
VEGFB[16]
CLPP[1]
GH2[10]
CRELD1[6]
CRTAM[18]
ZNF174[5]
HTR1A[2]
CA8[6]
ATP5F1D[5]
AMIGO2[13]
ZPR1[19]
OTUD7B[7]
KRT8[1]
PKD1[3]
PDP1[5]
SH2B3[3]
SHPK[12]
GSTT2B[5]
BHLHE40[8]
RANBP2[1]
PAPPA[11]
COL4A4[1]
GNGT1[4]
IDO1[8]
FUT8[10]
PHYKPL[1]
PVR[1]
VASH1[7]
HS6ST1[5]
KIR2DS4[3]
MRPL28[3]
DNLZ[1]
GSR[2]
STK4[3]
FCRLB[1]
NLGN1[1]
LSM8[3]

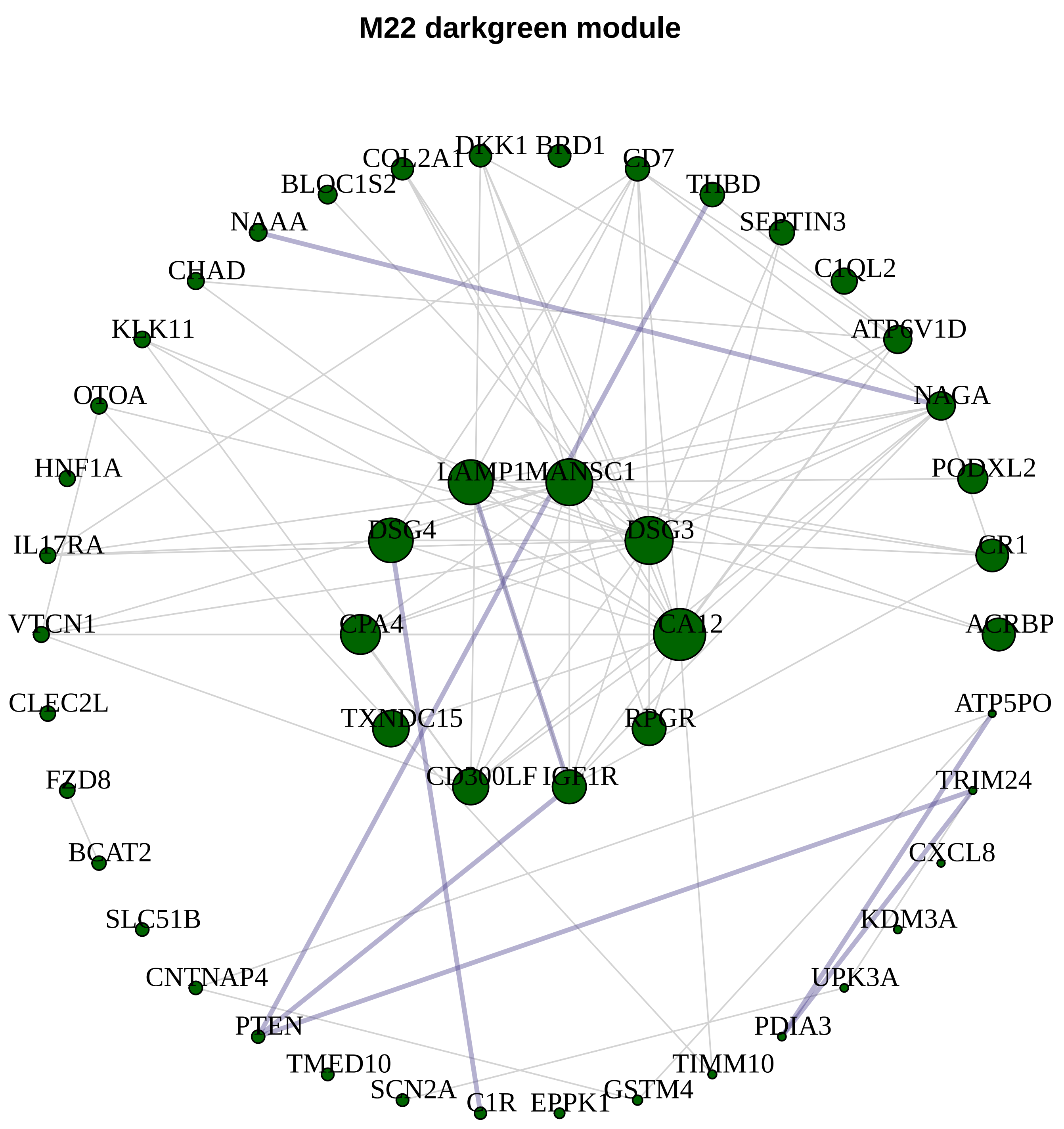
M22 darkgreen module
COL2A1
DKK1
BRD1
CD7
THBD
BLOC1S2
NAAA
SEPTIN3
C1QL2
CHAD
ATP6V1D
KLK11
NAGA
OTOA
HNF1A
LAMP1
MANSC1
PODXL2
IL17RA
DSG4
DSG3
CR1
VTCN1
CPA4
CA12
ACRBP
CLEC2L
TXNDC15
RPGR
ATP5PO
FZD8
CD300LF
IGF1R
TRIM24
BCAT2
CXCL8
SLC51B
KDM3A
CNTNAP4
UPK3A
PTEN
PDIA3
TMED10
TIMM10
SCN2A
C1R
EPPK1
GSTM4

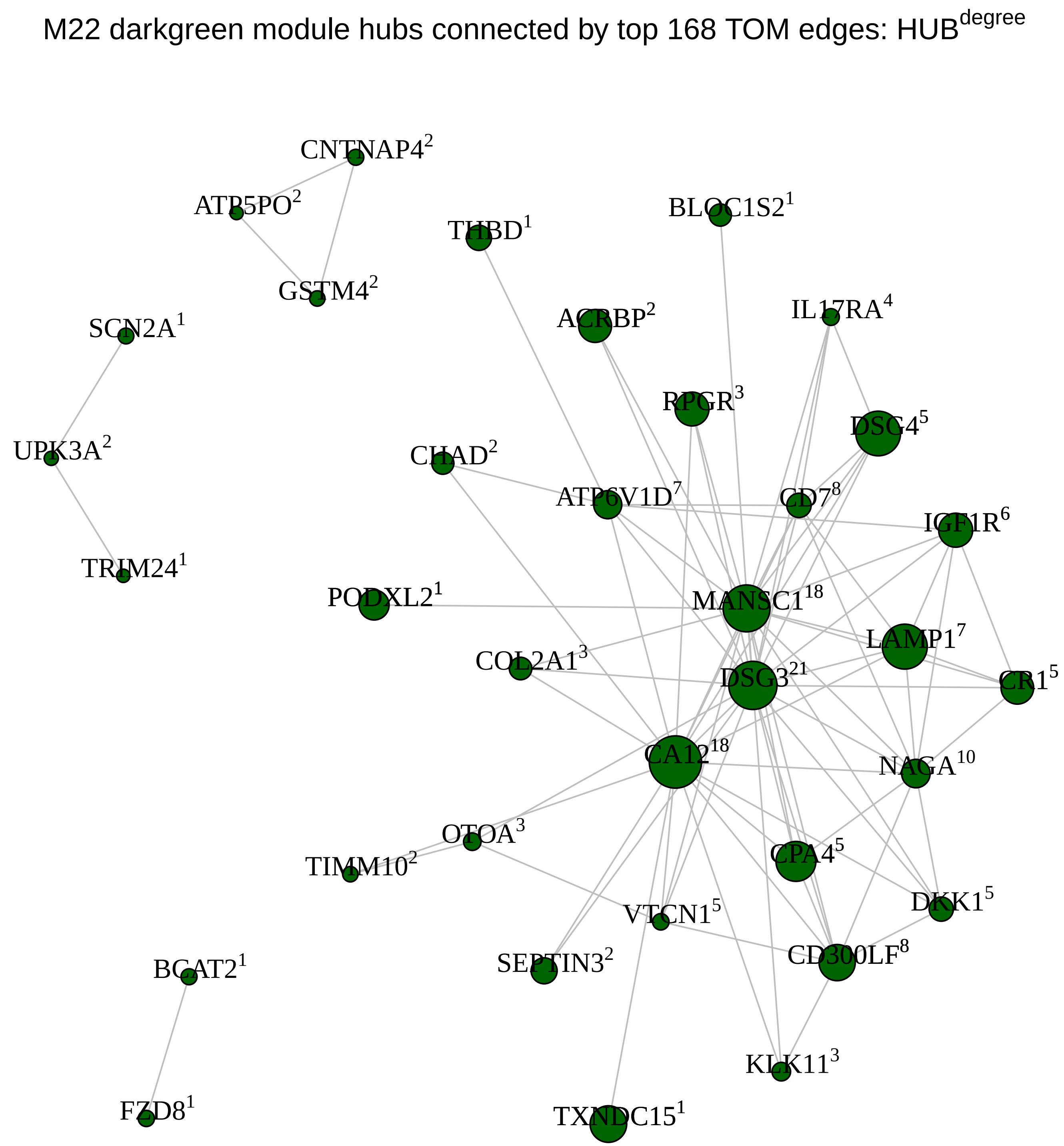
M22 darkgreen module hubs connected by top 168 TOM edges: HUB$^{degree}$
CNTNAP4$^{2}$
ATP5PO$^{2}$
THBD$^{1}$
BLOC1S2$^{1}$
GSTM4$^{2}$
SCN2A$^{1}$
ACRBP$^{2}$
IL17RA$^{4}$
RPGR$^{3}$
UPK3A$^{2}$
DSG4$^{5}$
CHAD$^{2}$
ATP6V1D$^{7}$
CD7$^{8}$
IGF1R$^{6}$
TRIM24$^{1}$
PODXL2$^{1}$
MANSC1$^{18}$
LAMP1$^{7}$
COL2A1$^{3}$
DSG3$^{21}$
CR1$^{5}$
CA12$^{18}$
NAGA$^{10}$
OTOA$^{3}$
TIMM10$^{2}$
CPA4$^{5}$
DKK1$^{5}$
VTCN1$^{5}$
BCAT2$^{1}$
SEPTIN3$^{2}$
CD300LF$^{8}$
KLK11$^{3}$
FZD8$^{1}$
TXNDC15$^{1}$

**M14 cyan module**

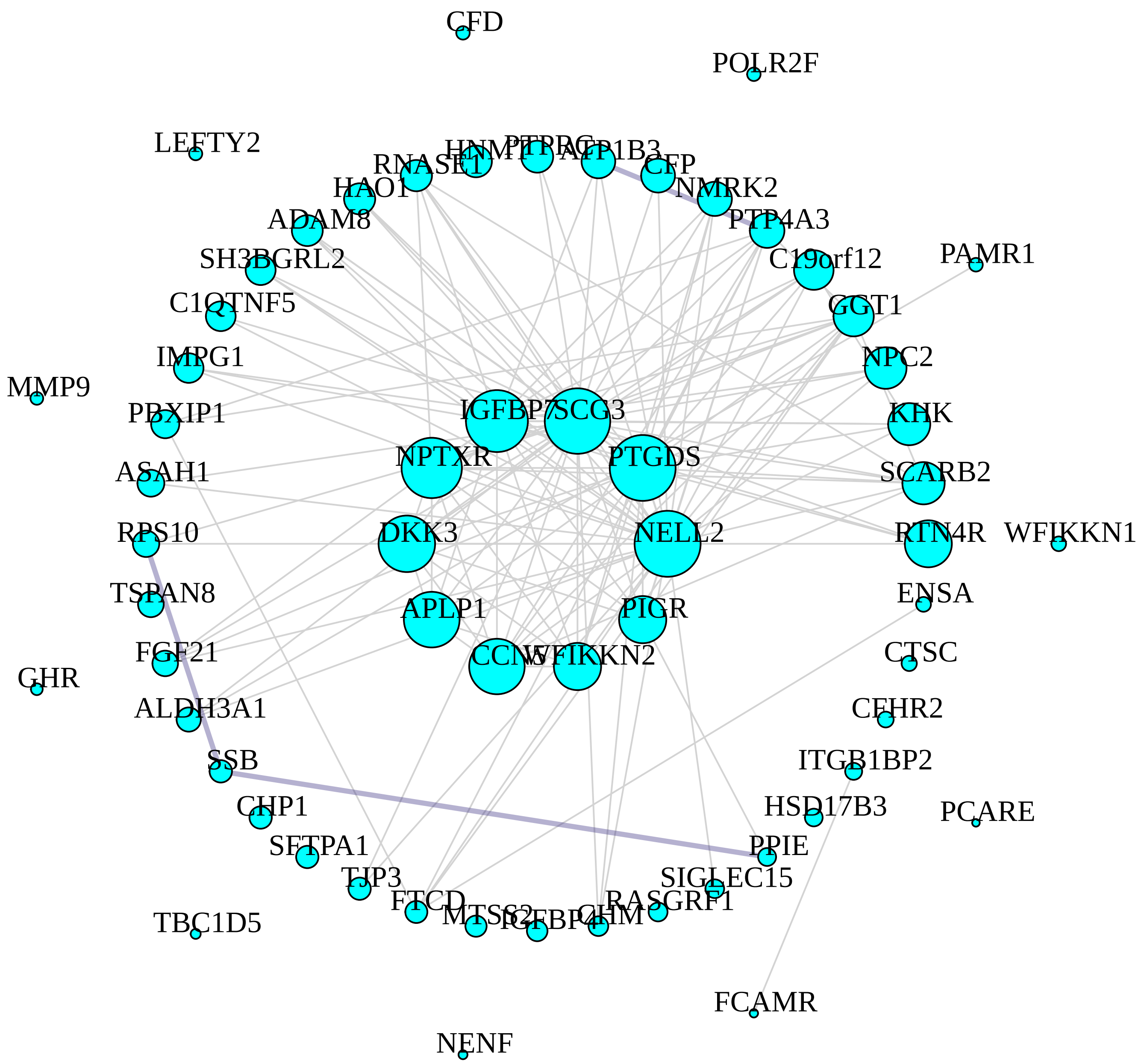


M14 cyan module hubs connected by top 280 TOM edges: HUB$^{degree}$

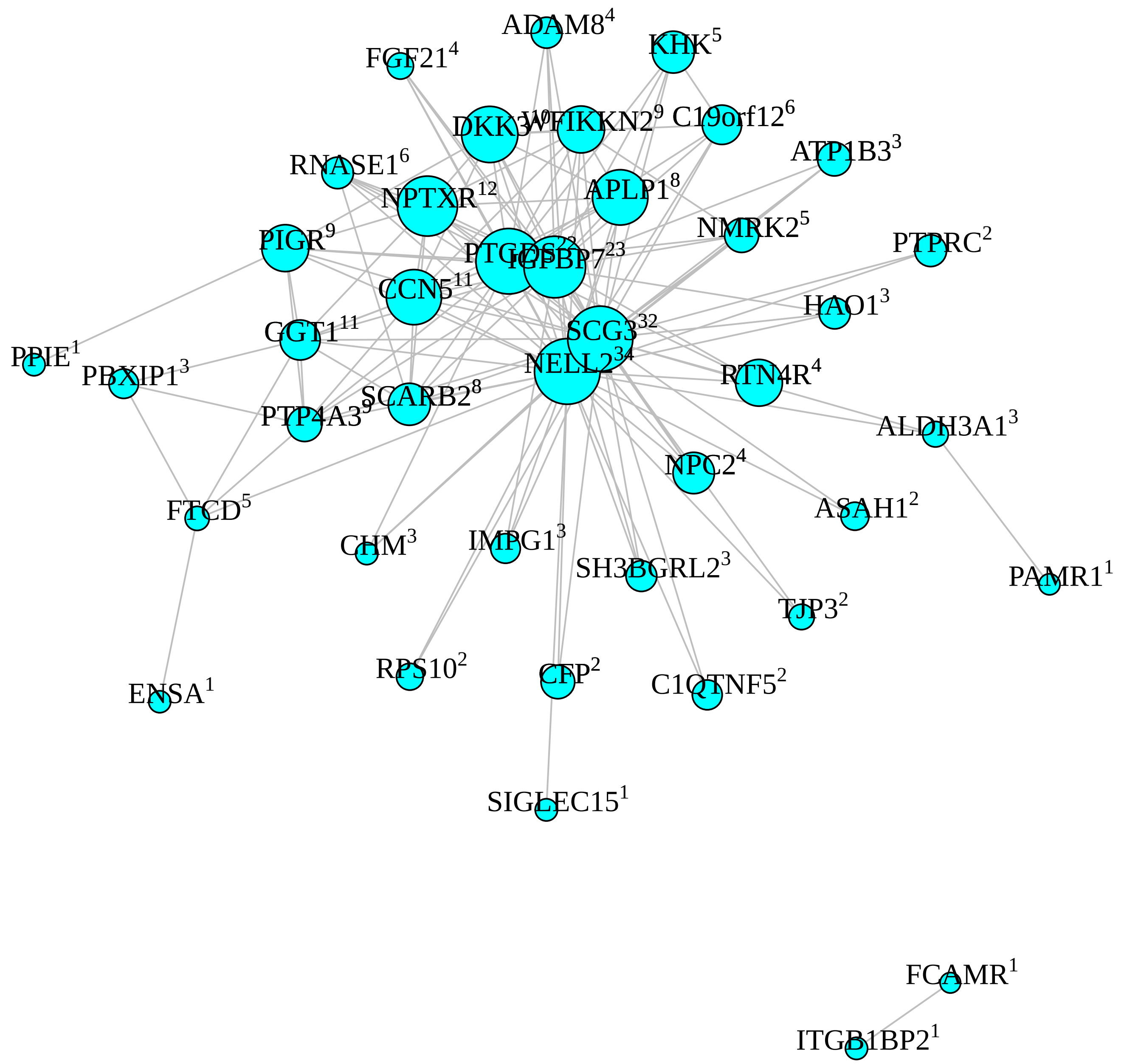

**M10 purple module**

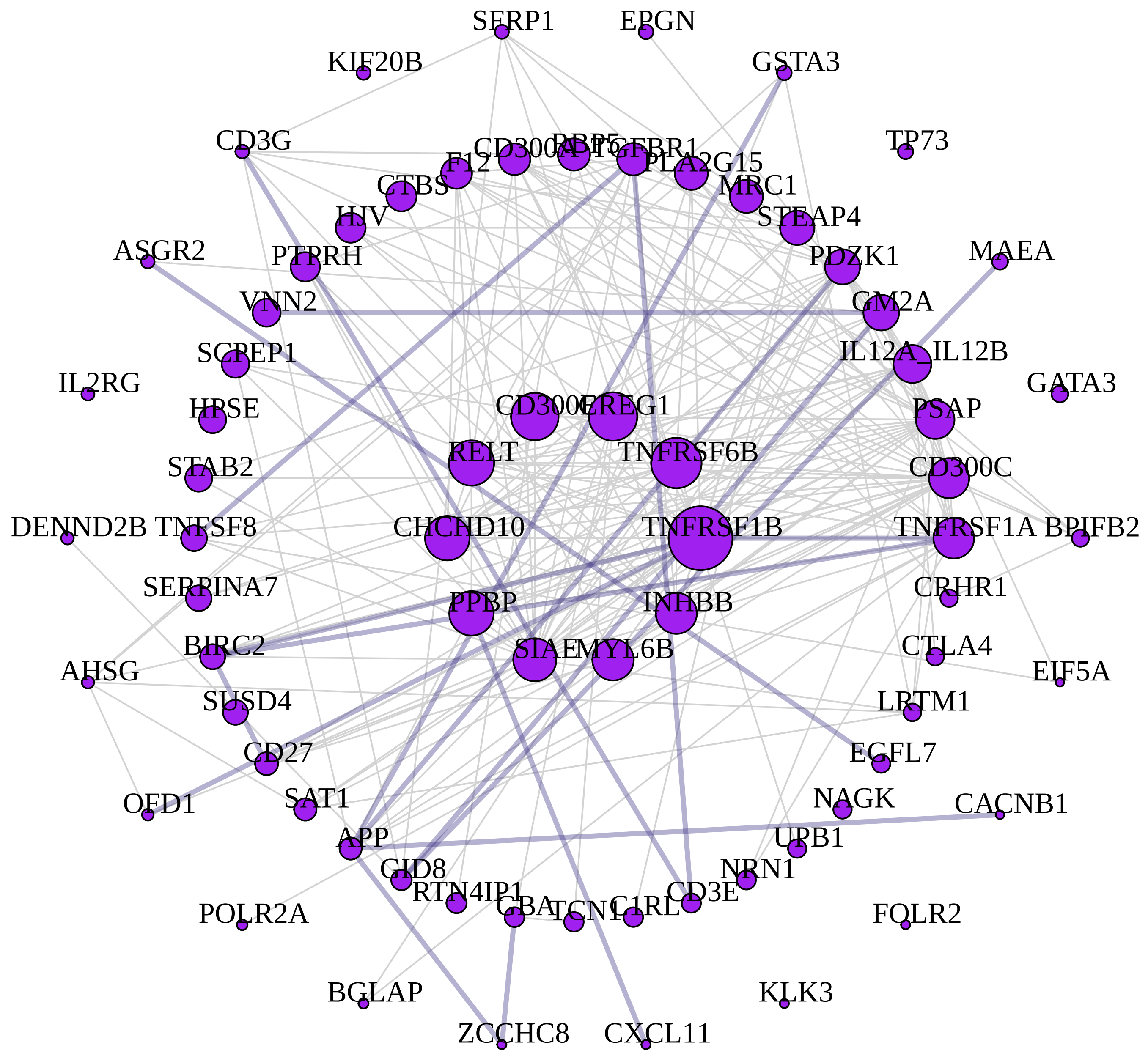


M10 purple module hubs connected by top 396 TOM edges: HUB$^{degree}$

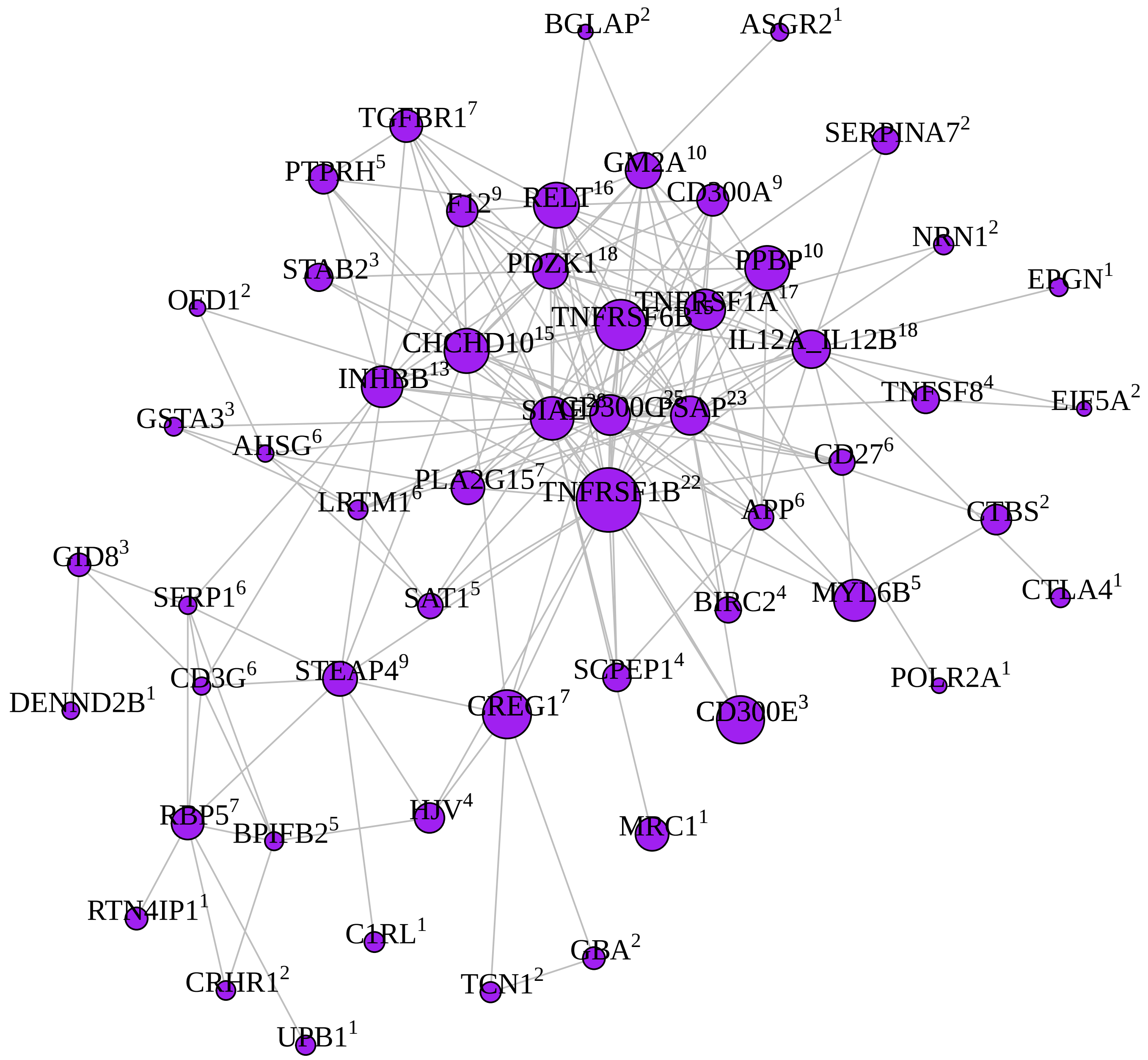

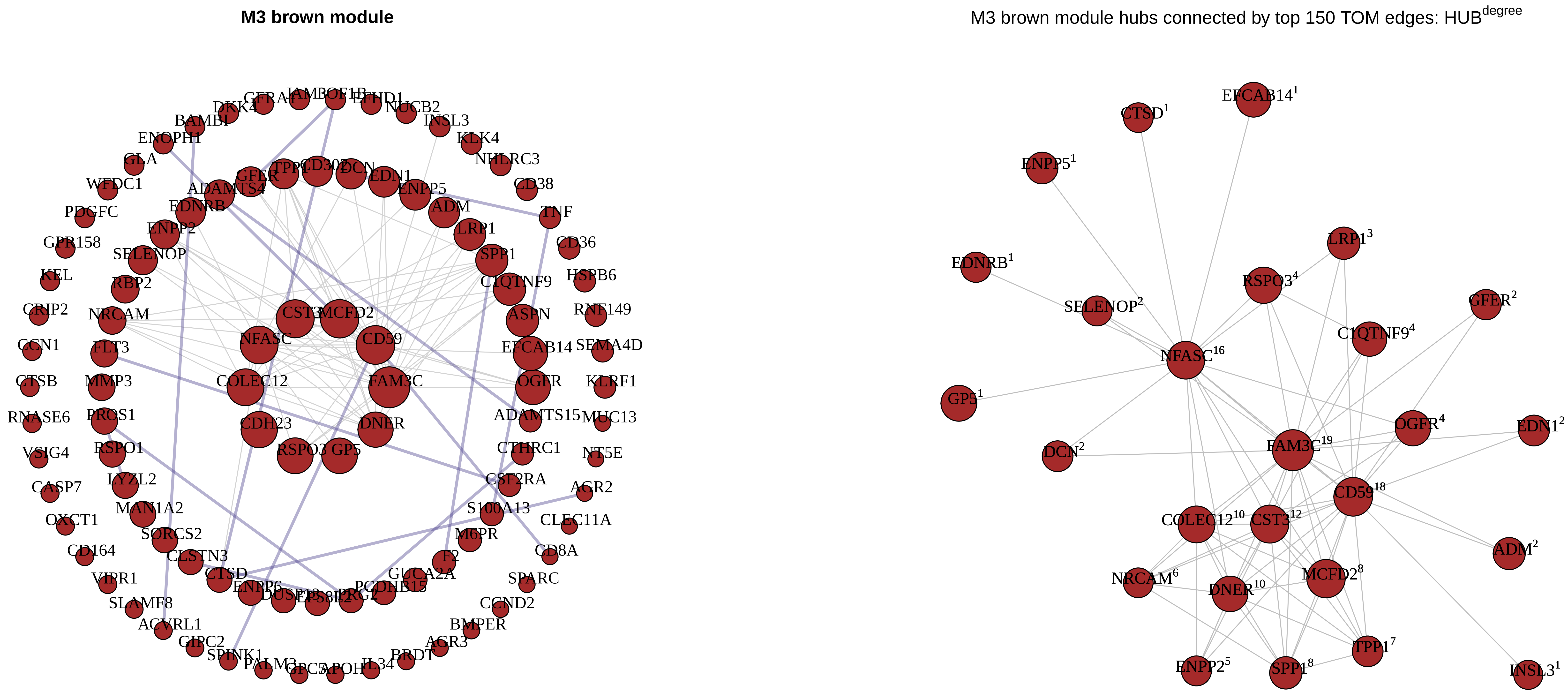
M3 brown module
DKK4
BAMBI
ENOPH1
GLA
WFDC1
PDGFC
GPR158
KEL
CRIP2
CCN1
CTSB
RNASE6
VSIG4
CASP7
OXCT1
CD164
VIPR1
SLAMF8
ACVRL1
GIPC2
SPINK1
PALM3
GPC5
APOH
IL34
BRDT
AGR3
BMPER
CCND2
SPARC
CD8A
CLEC11A
AGR2
NT5E
MUC13
KLRF1
SEMA4D
RNF149
HSPB6
CD36
TNF
CD38
NHLRC3
KLK4
INSL3
NUCB2
EFHD1
GFER
TPP1
CD302
DCN
EDN1
ENPP5
ADM
LRP1
SPP1
C1QTNF9
ASPN
EFCAB14
OGFR
ADAMTS15
CTHRC1
CSF2RA
S100A13
M6PR
F2
GUCA2A
PCDHB15
PRG2
ENPP6
CTSD
CLSTN3
SORCS2
MAN1A2
LYZL2
RSPO1
PROS1
MMP3
FLT3
NRCAM
RBP2
SELENOP
ENPP2
EDNRB
ADAMTS4
CST3
MCFD2
NFASC
CD59
COLEC12
FAM3C
CDH23
DNER
RSPO3
GP5
M3 brown module hubs connected by top 150 TOM edges: HUB$^{degree}$
EFCAB14$^{1}$
CTSD$^{1}$
ENPP5$^{1}$
EDNRB$^{1}$
LRP1$^{3}$
RSPO3$^{4}$
GFER$^{2}$
SELENOP$^{2}$
C1QTNF9$^{4}$
NFASC$^{16}$
GP5$^{1}$
OGFR$^{4}$
EDN1$^{2}$
DCN$^{2}$
FAM3C$^{19}$
CD59$^{18}$
COLEC12$^{10}$
CST3$^{12}$
ADM$^{2}$
NRCAM$^{6}$
DNER$^{10}$
MCFD2$^{8}$
TPP1$^{7}$
ENPP2$^{5}$
SPP1$^{8}$
INSL3$^{1}$

**M12 tan module**

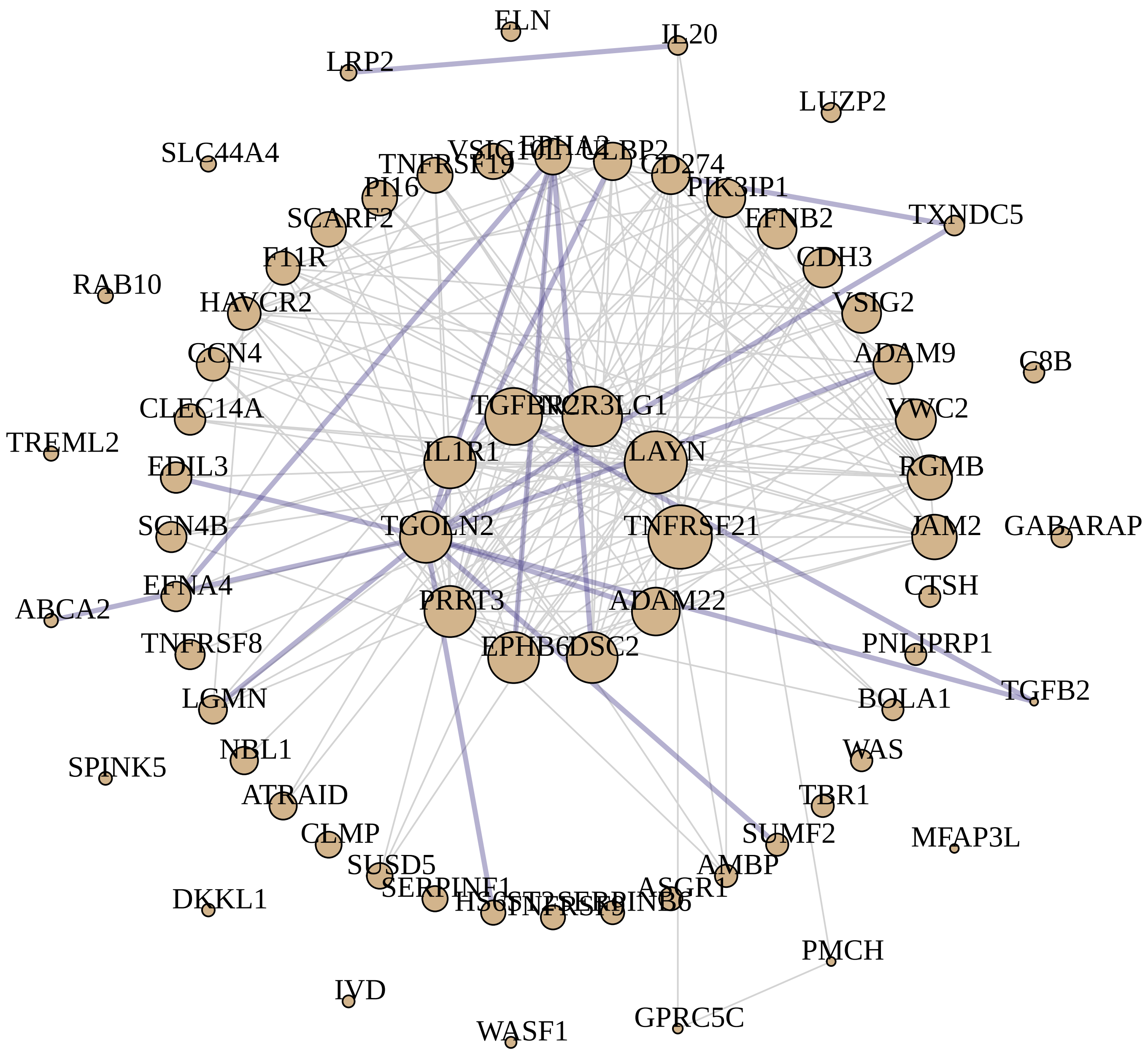


M12 tan module hubs connected by top 362 TOM edges: HUB$^{degree}$

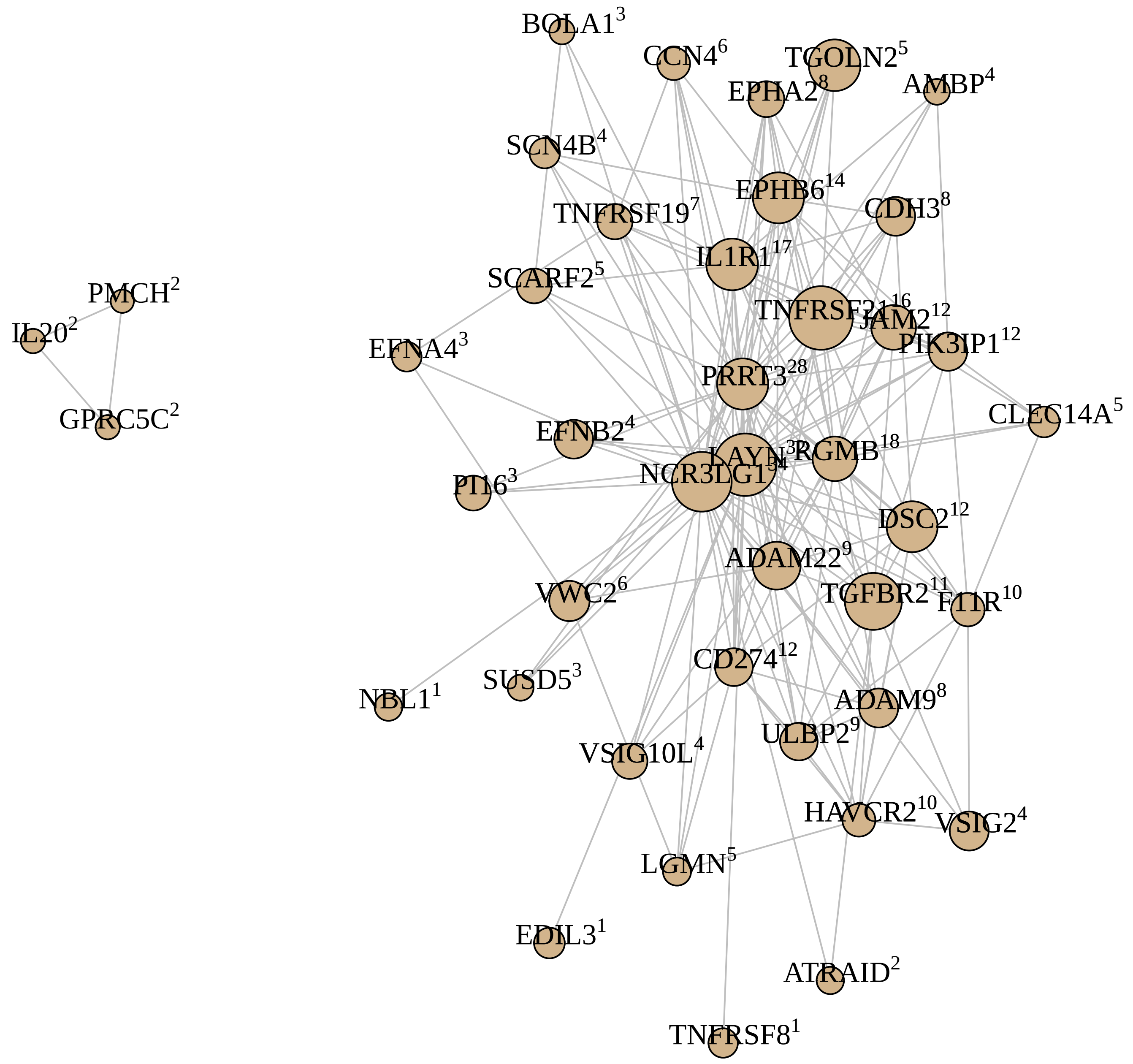

**M27 white module**

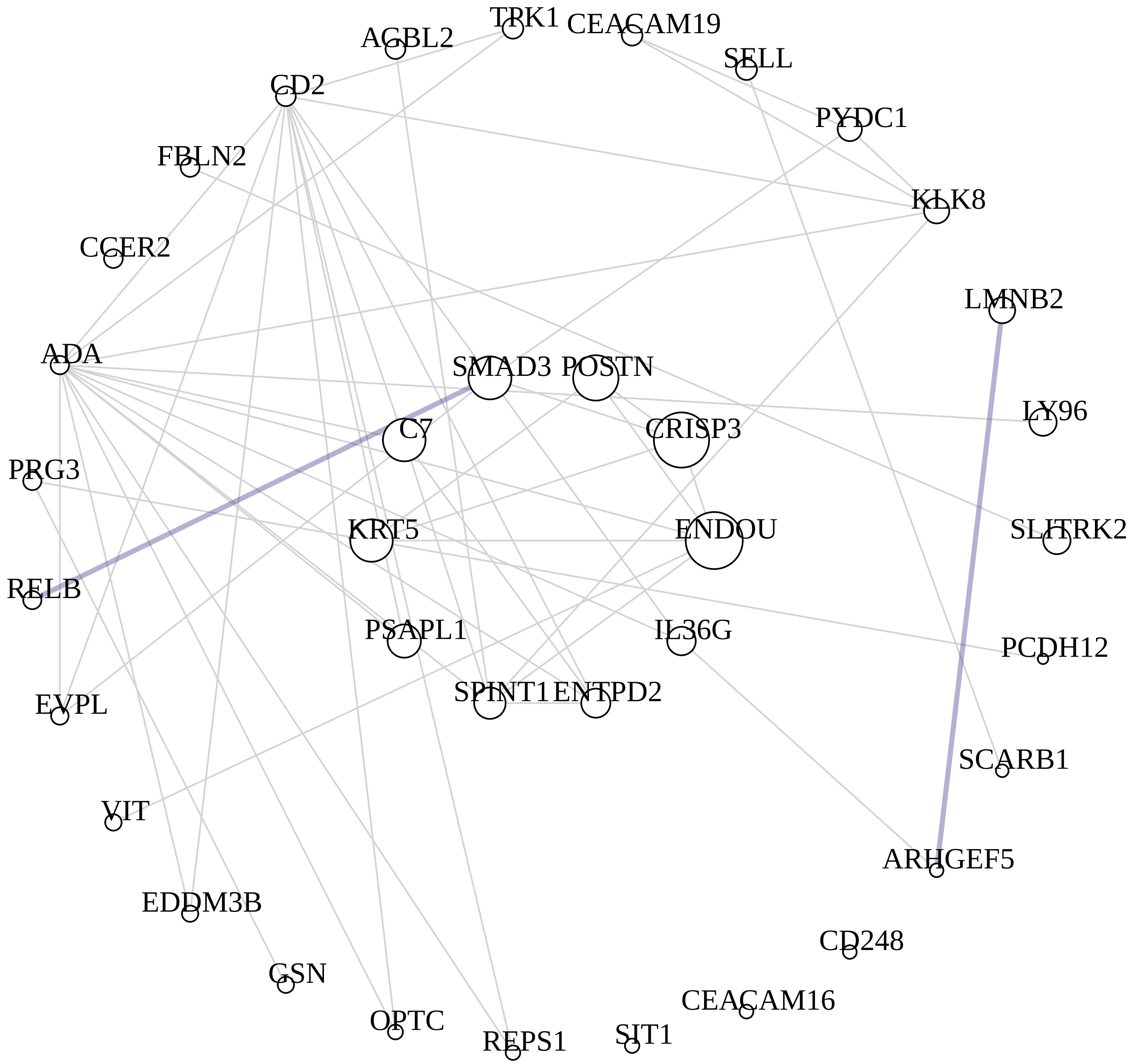


M27 white module hubs connected by top 96 TOM edges: $HUB^{degree}$

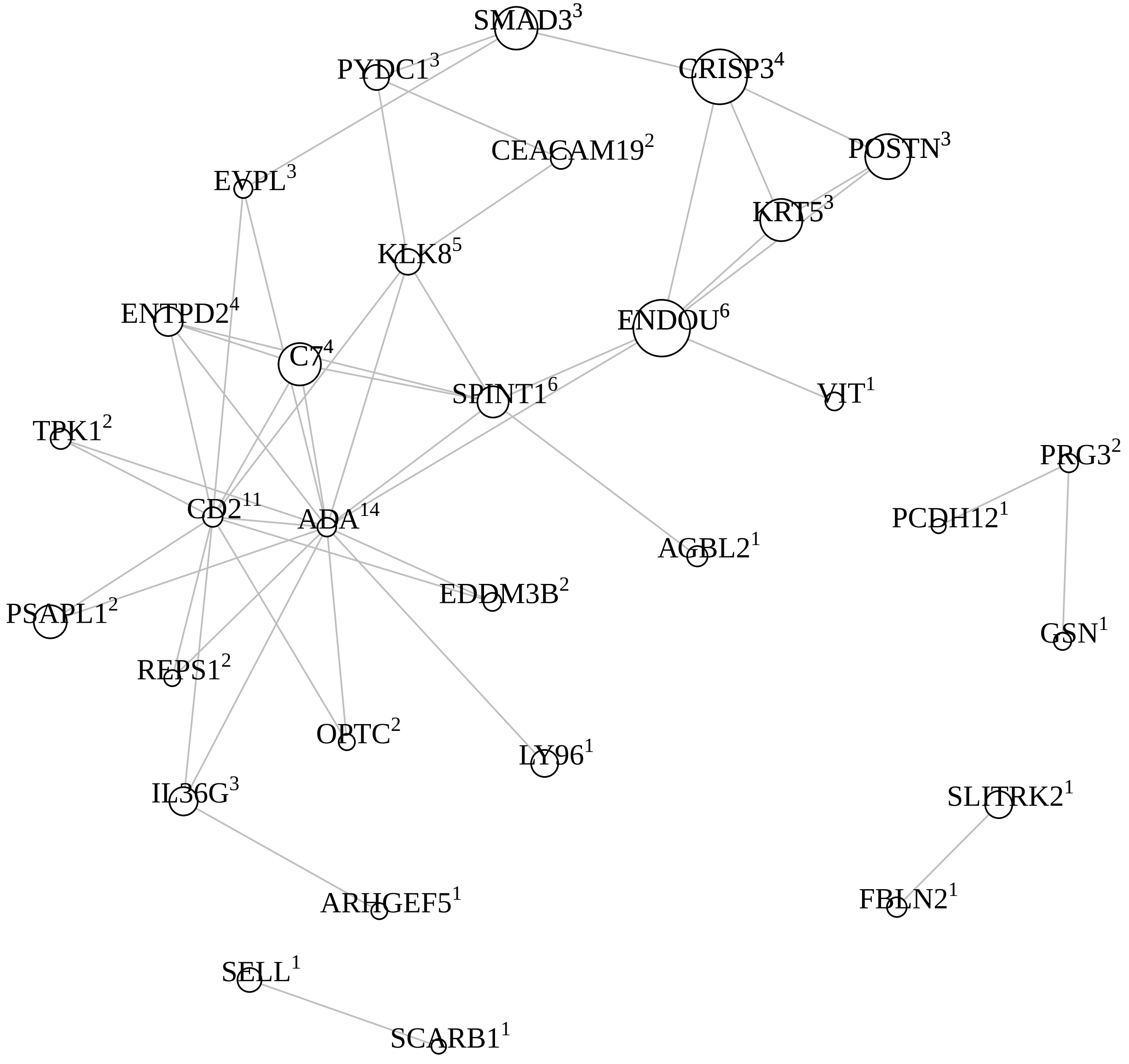

**M23 darkturquoise module**

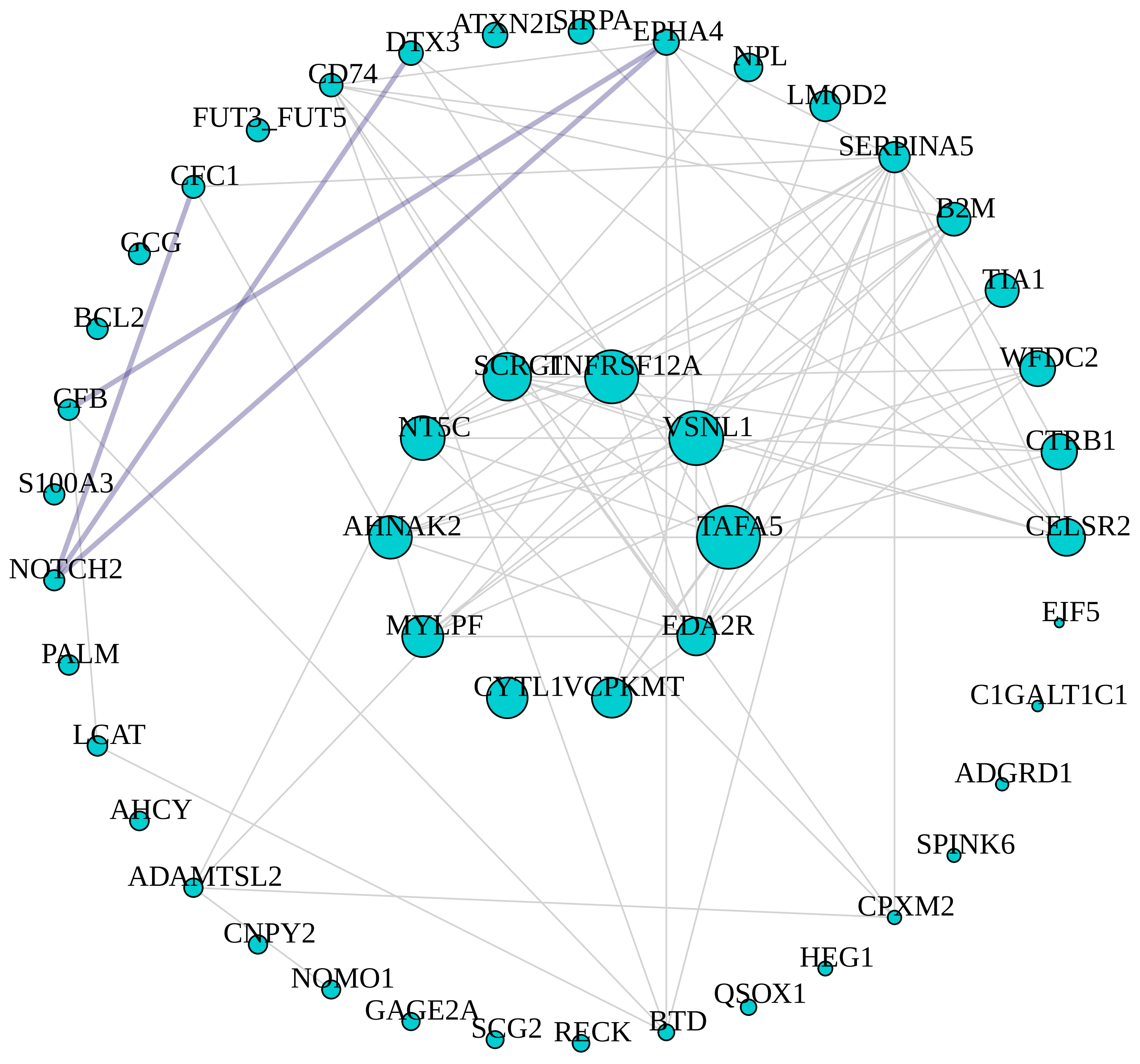


M23 darkturquoise module hubs connected by top 160 TOM edges: HUB$^{degree}$

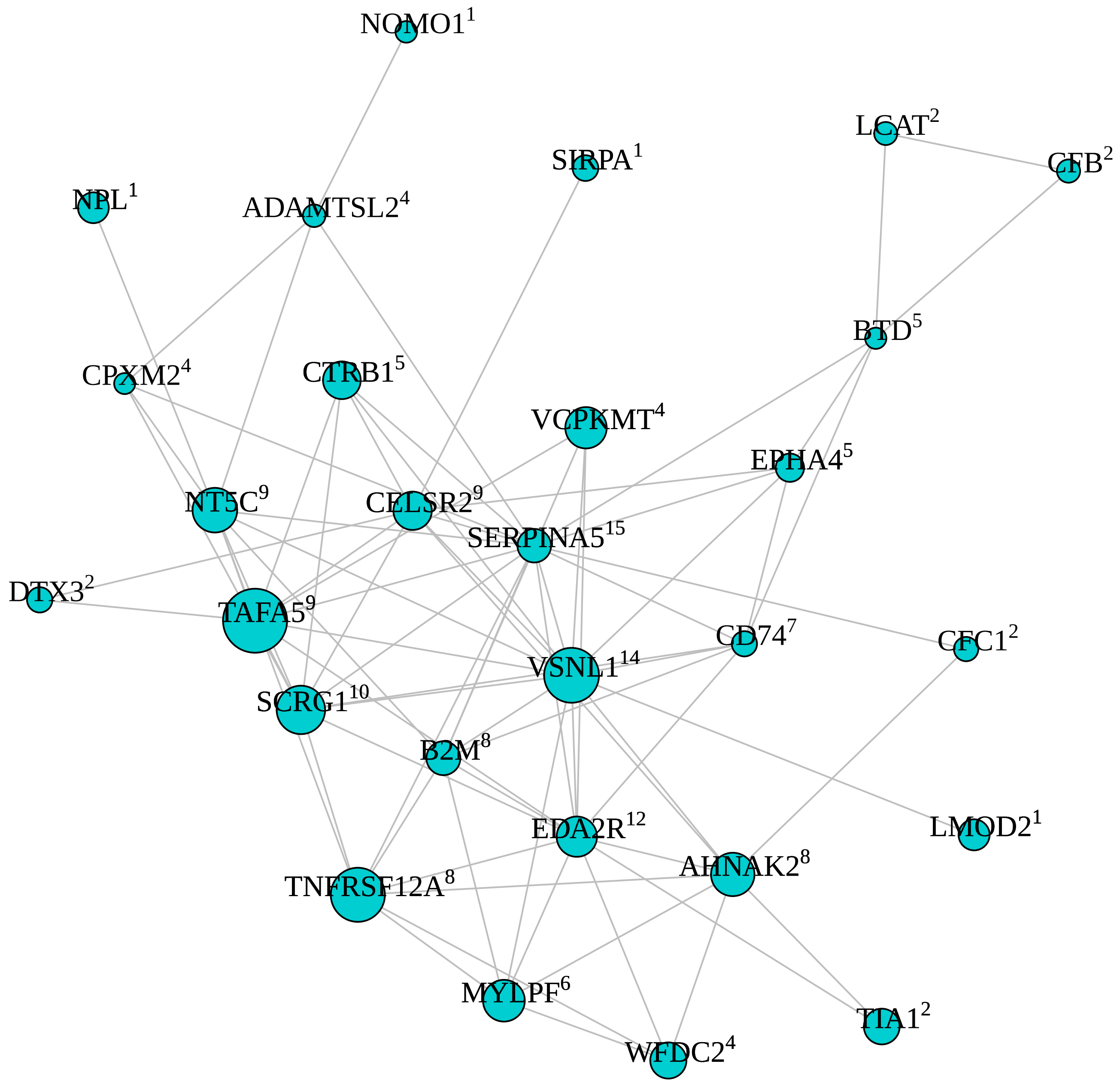

**M16 lightcyan module**

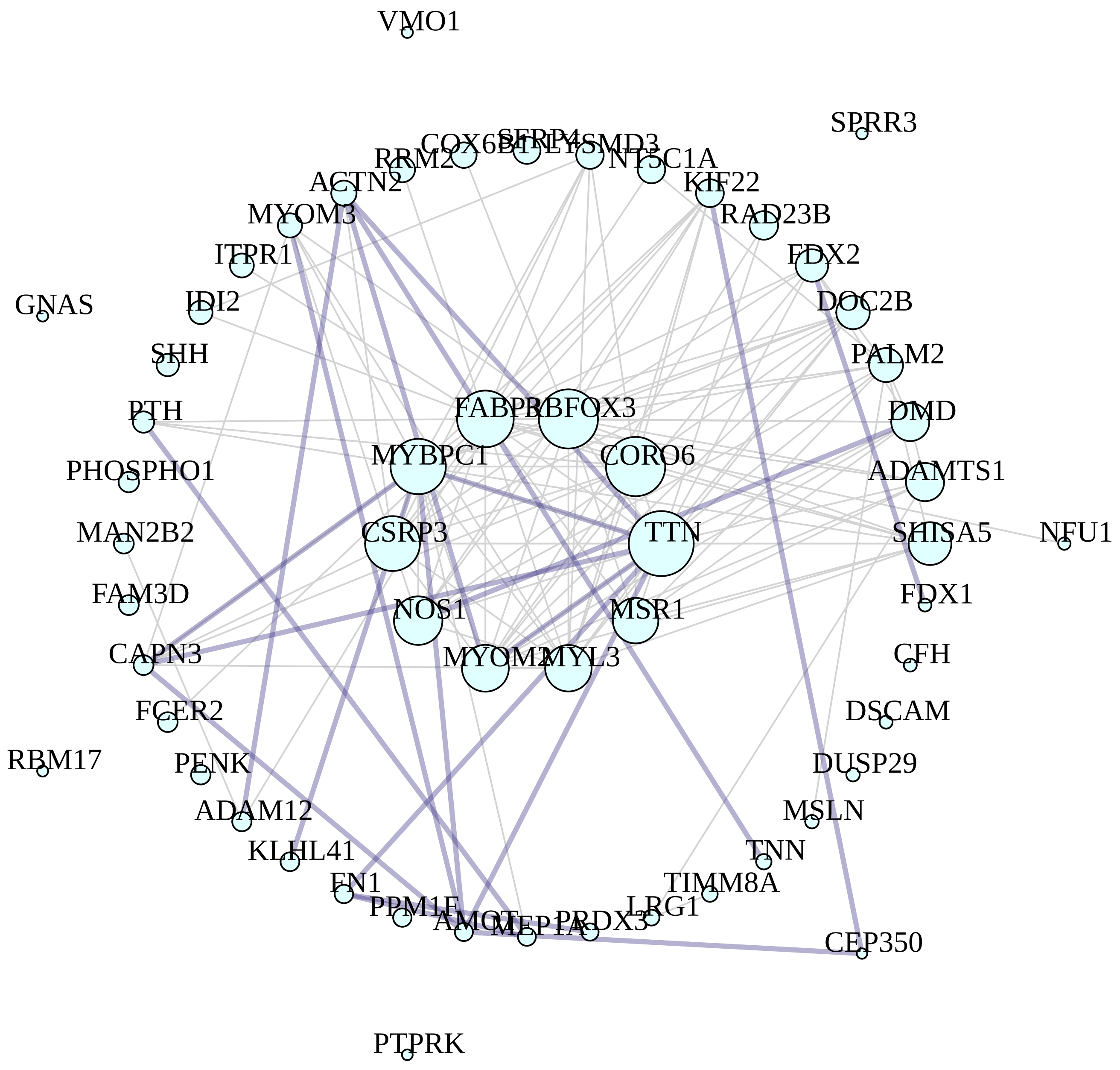


M16 lightcyan module hubs connected by top 242 TOM edges: HUB$^{degree}$

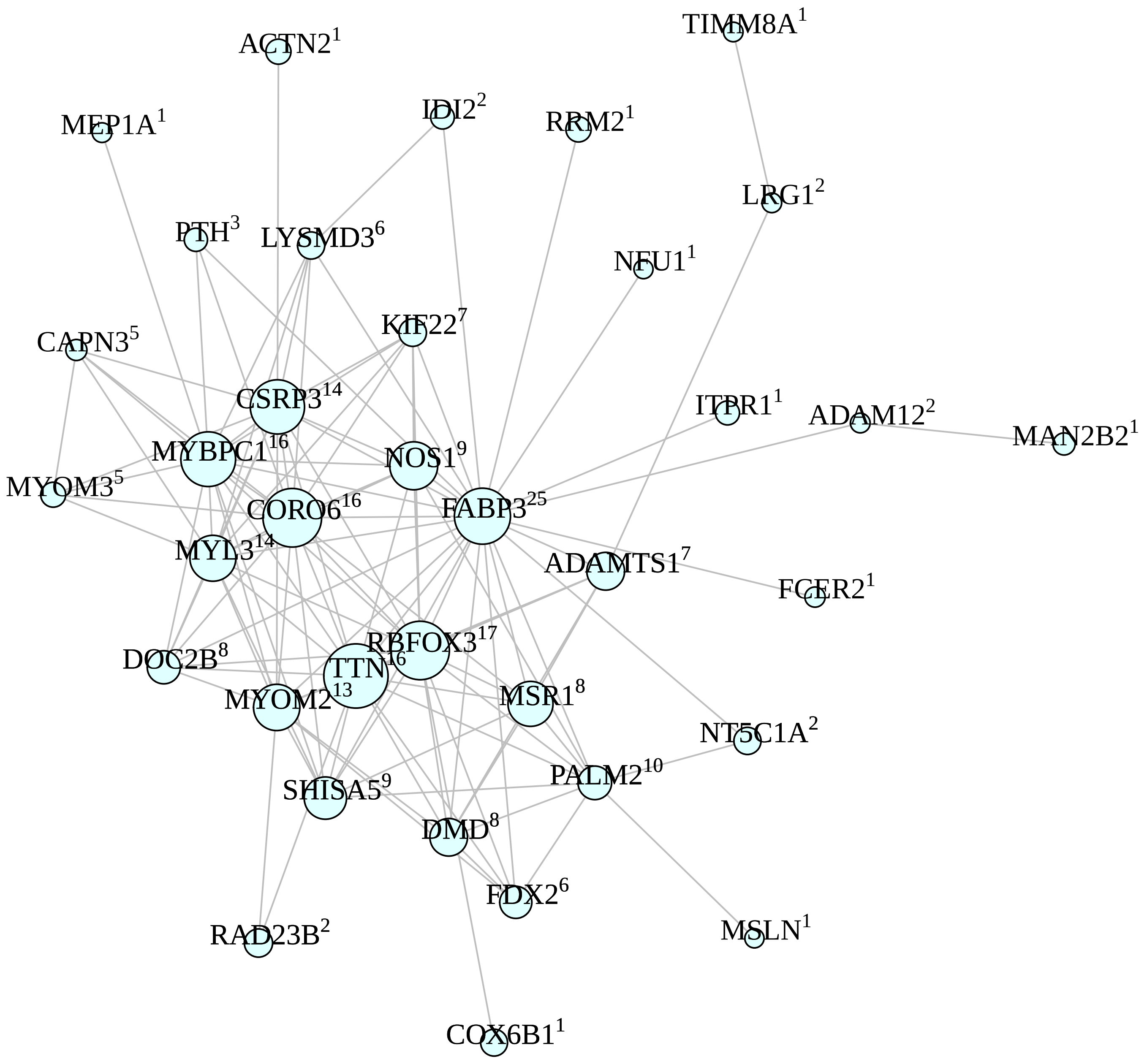

**M18 lightgreen module**

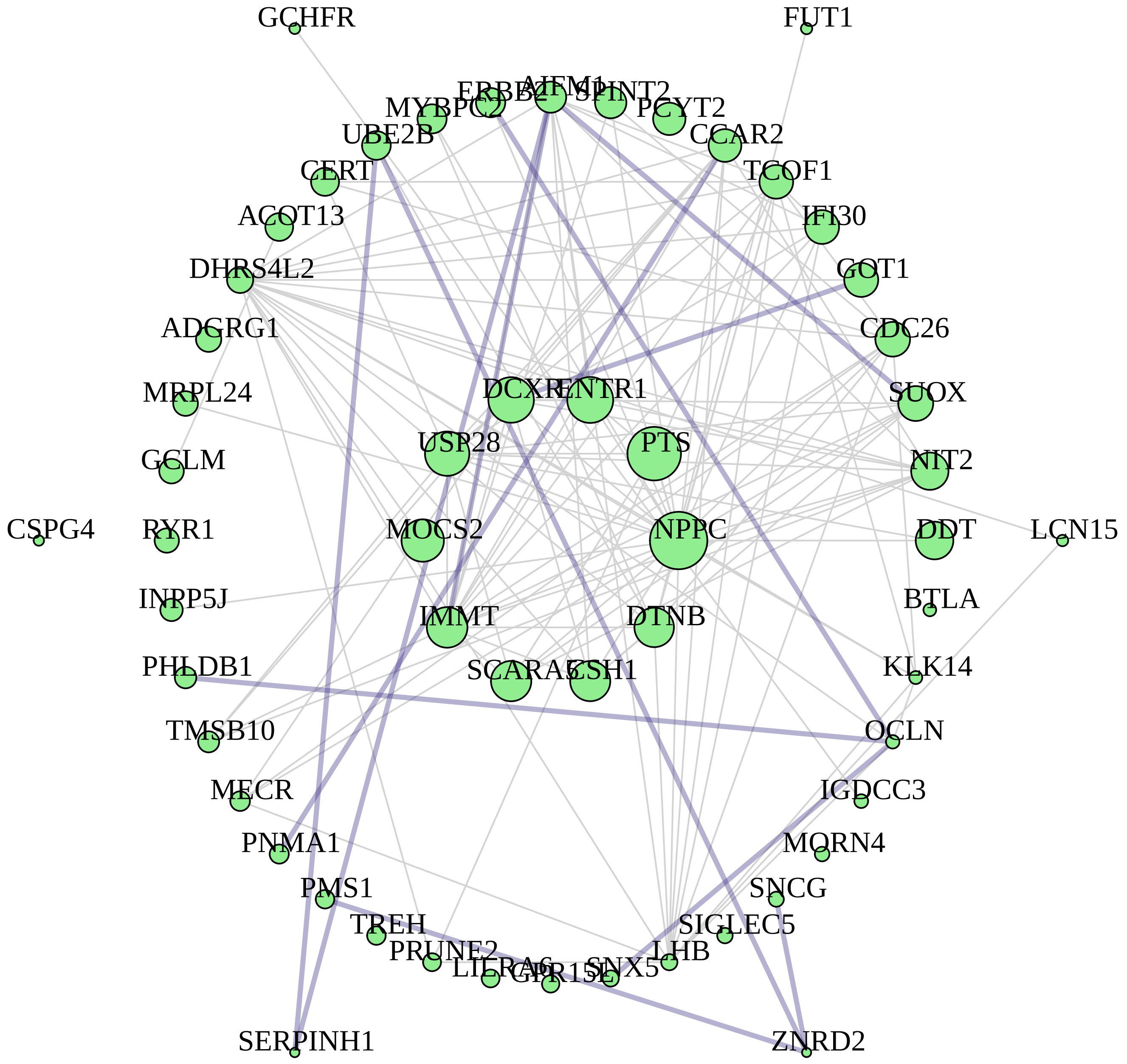


M18 lightgreen module hubs connected by top 234 TOM edges: HUB$^{degree}$

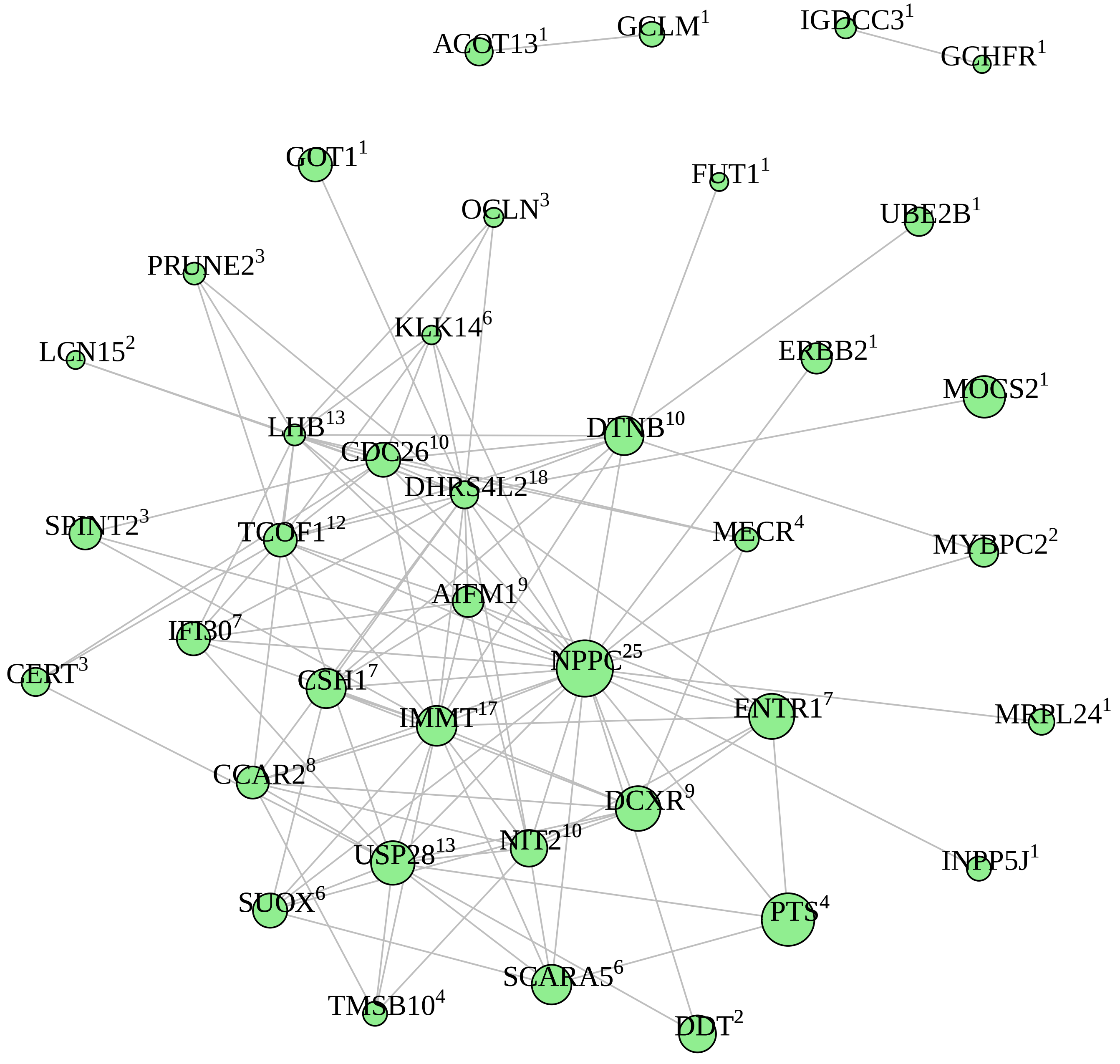
