## Supplementary File 6 for "Maintenance of chronic neuroinflammation in multiple sclerosis via interferon signaling and CD8 T cell-mediated cytotoxicity"

**M11 greenyellow module**

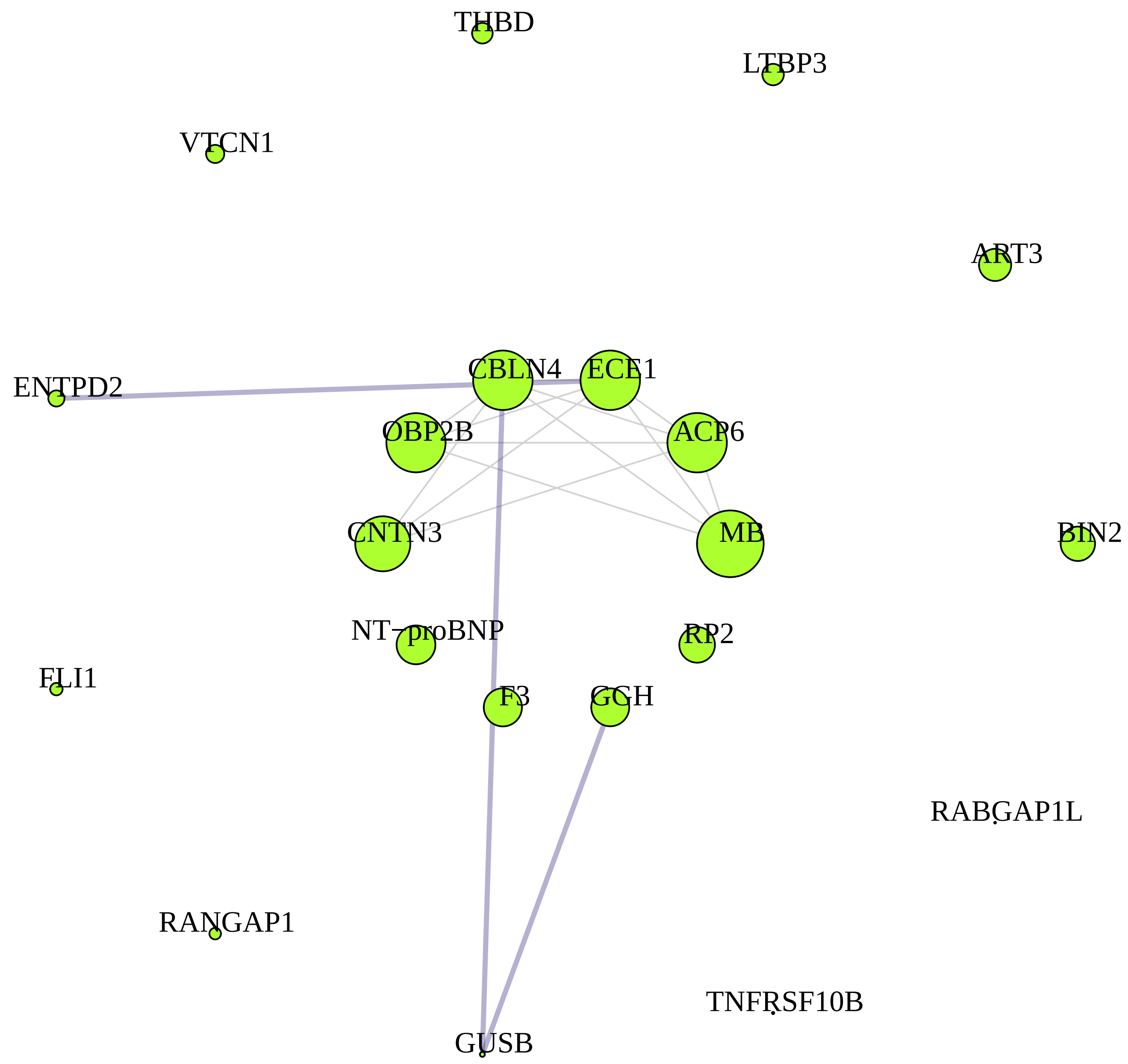


M11 greenyellow module hubs connected by top 26 TOM edges: $HUB^{degree}$

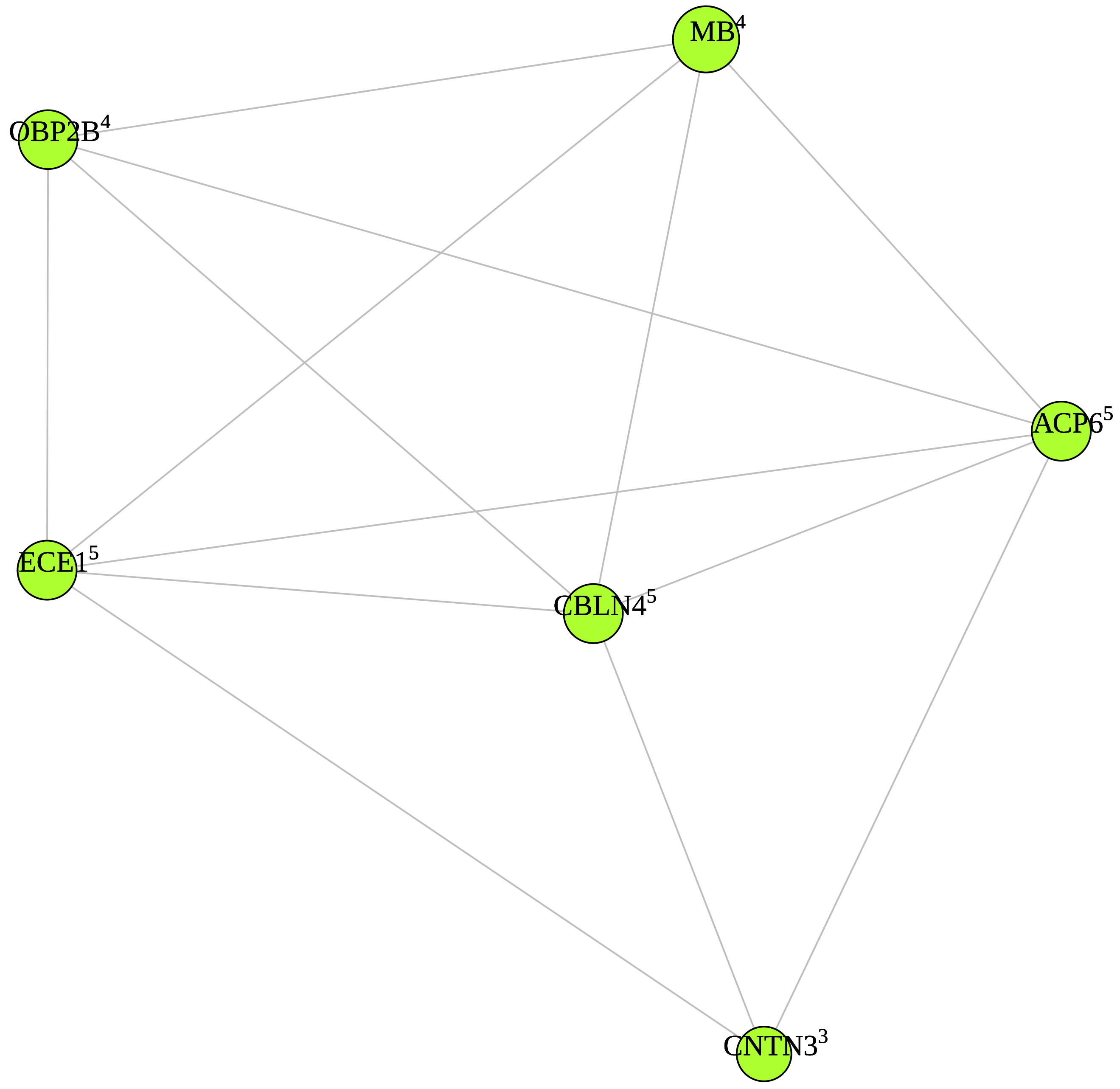

**M4 yellow module** M4 yellow module hubs connected by top 420 TOM edges: HUB$^{degree}$

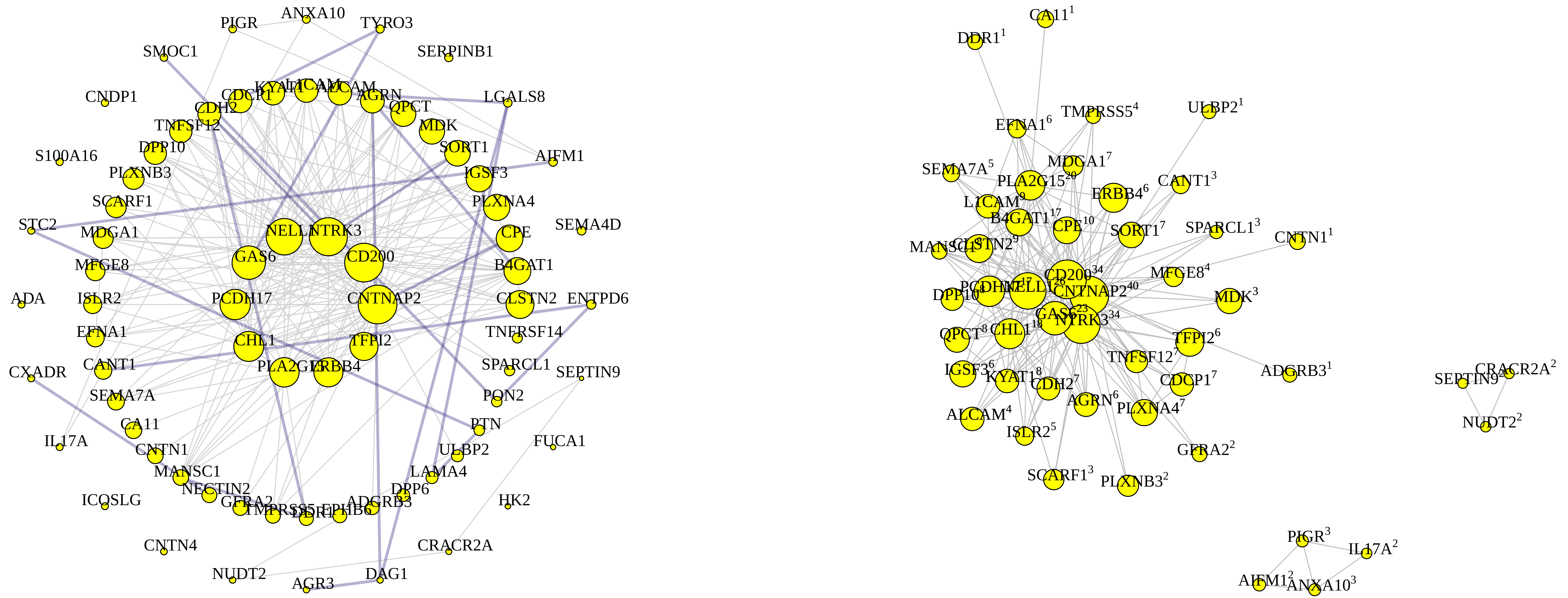

**M7 black module**

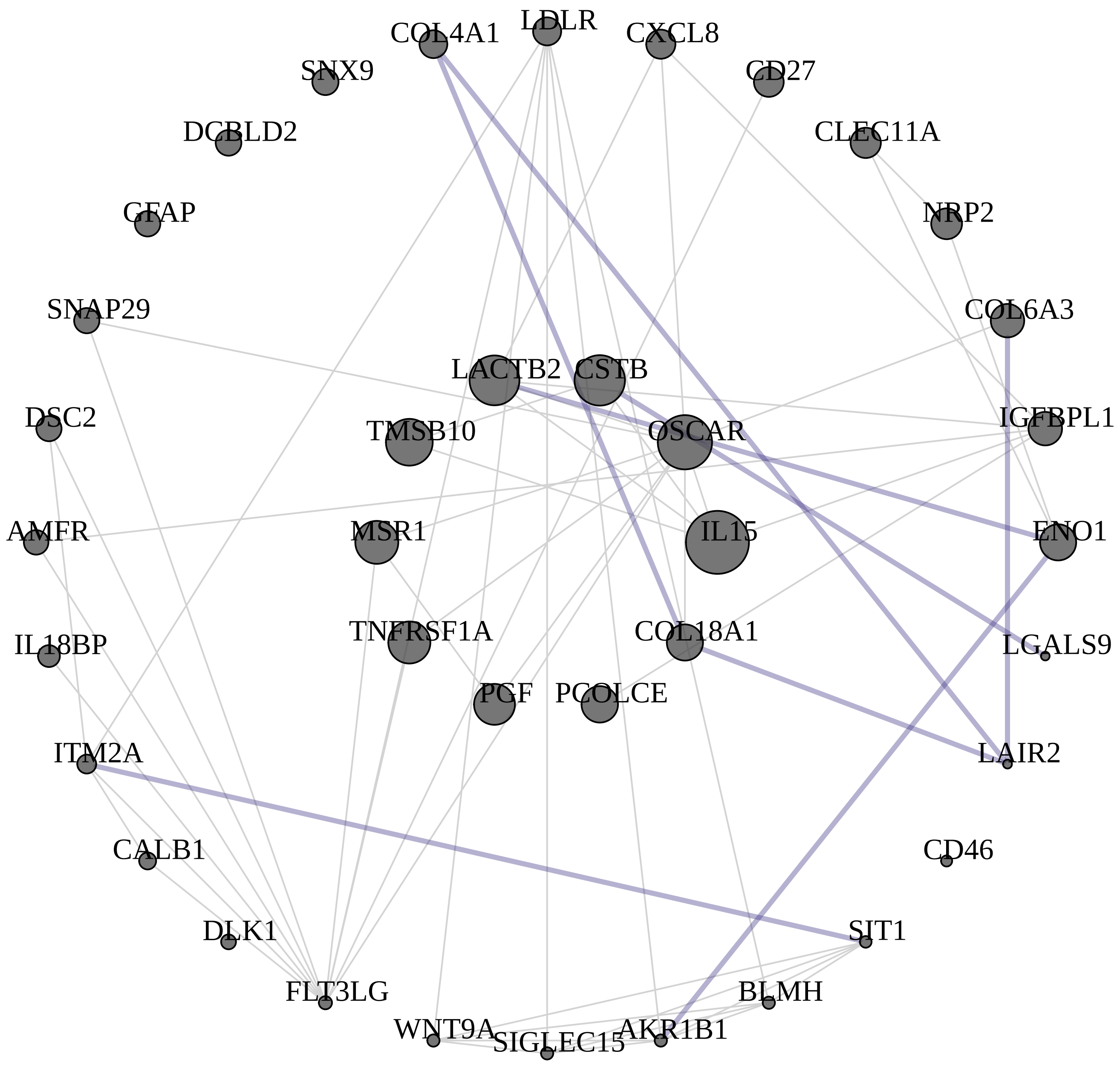


M7 black module hubs connected by top 102 TOM edges: HUB$^{degree}$

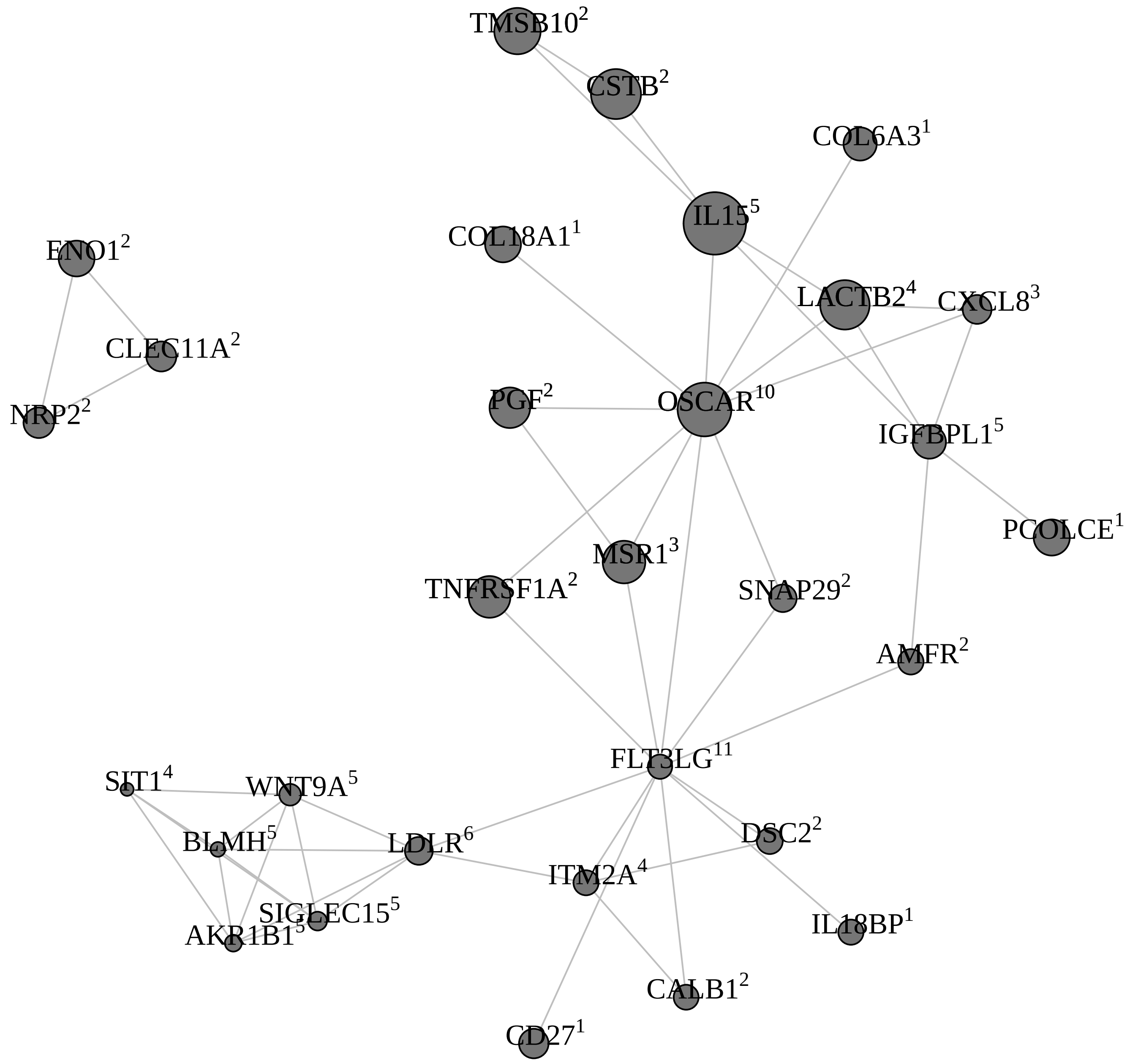

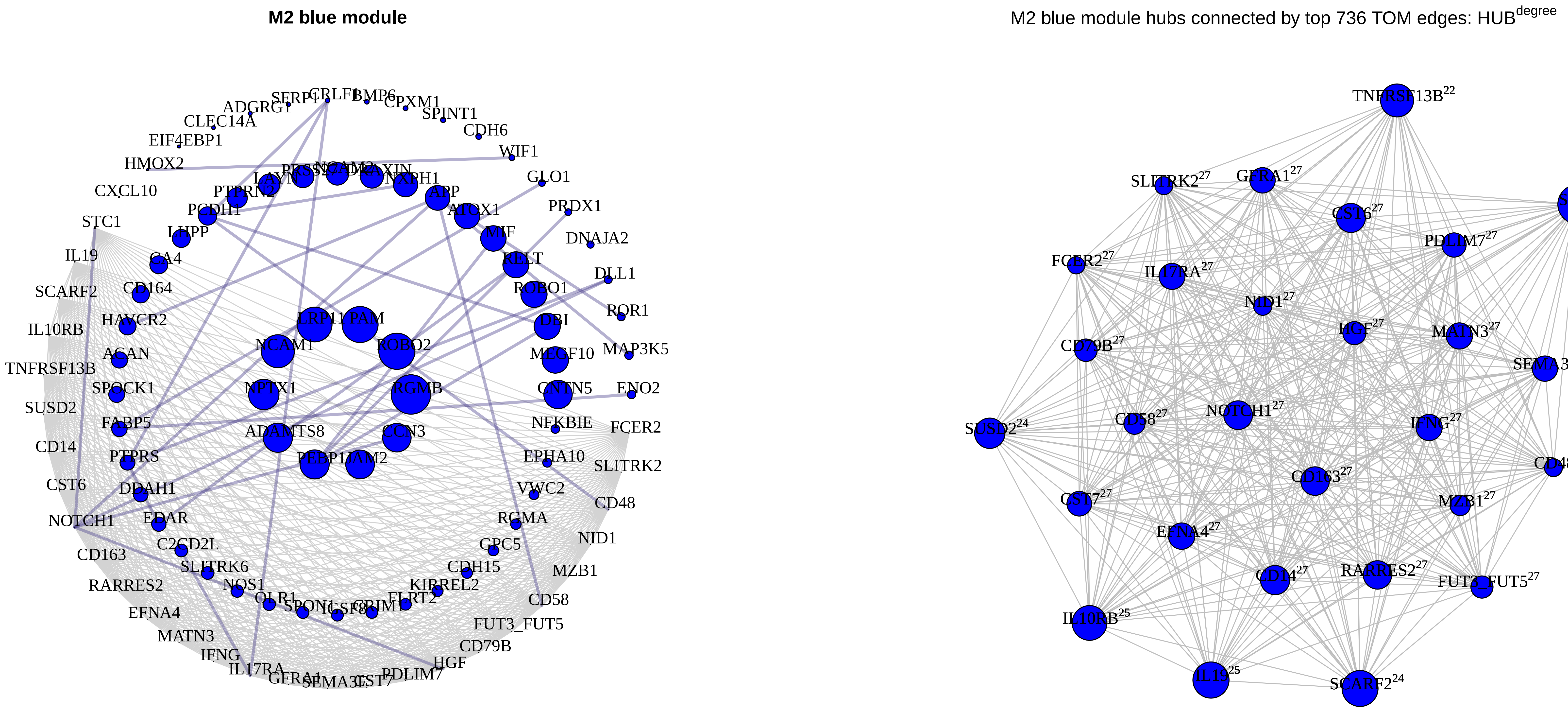
M2 blue module
SERPF1
CRLF1
BMP6
CPXM1
ADGRG1
SPINT1
CLEC14A
CDH6
EIF4EBP1
WIF1
HMOX2
GLO1
CXCL10
PTPRN2
NXPH1
APP
PRDX1
STC1
PCDH1
ATOX1
LHPP
MIF
DNAJA2
IL19
CA4
RELT
DLL1
SCARF2
CD164
ROBO1
IL10RB
HAVCR2
LRP11
PAM
DBI
ROR1
ACAN
NCAM1
ROBO2
MEGF10
MAP3K5
TNFRSF13B
SPOCK1
NPTX1
RGMB
CNTN5
ENO2
SUSD2
FABP5
NFKBIE
FCER2
CD14
ADAMTS8
CCN3
PTPRS
PEBP1
JAM2
EPHA10
SLITRK2
CST6
DDAH1
VWC2
CD48
NOTCH1
EDAR
RGMA
NID1
CD163
C2CD2L
GPC5
SLITRK6
CDH15
MZB1
RARRES2
NOS1
KIRREL2
OLR1
FLRT2
CD58
EFNA4
FUT3_FUT5
MATN3
CD79B
IFNG
HGF
IL17RA
PDLIM7
M2 blue module hubs connected by top 736 TOM edges: HUB^degree
TNFRSF13B22
SLITRK227
GFRA127
STC122
CST627
PDLIM727
FCER227
IL17RA27
NID127
HGF27
MATN327
CD79B27
SEMA3F27
SUSD224
CD5827
NOTCH127
IFNG27
CD4827
CD16327
CST727
MZB127
EFNA427
CD1427
RARRES227
FUT3_FUT527
IL10RB25
IL1925
SCARF224

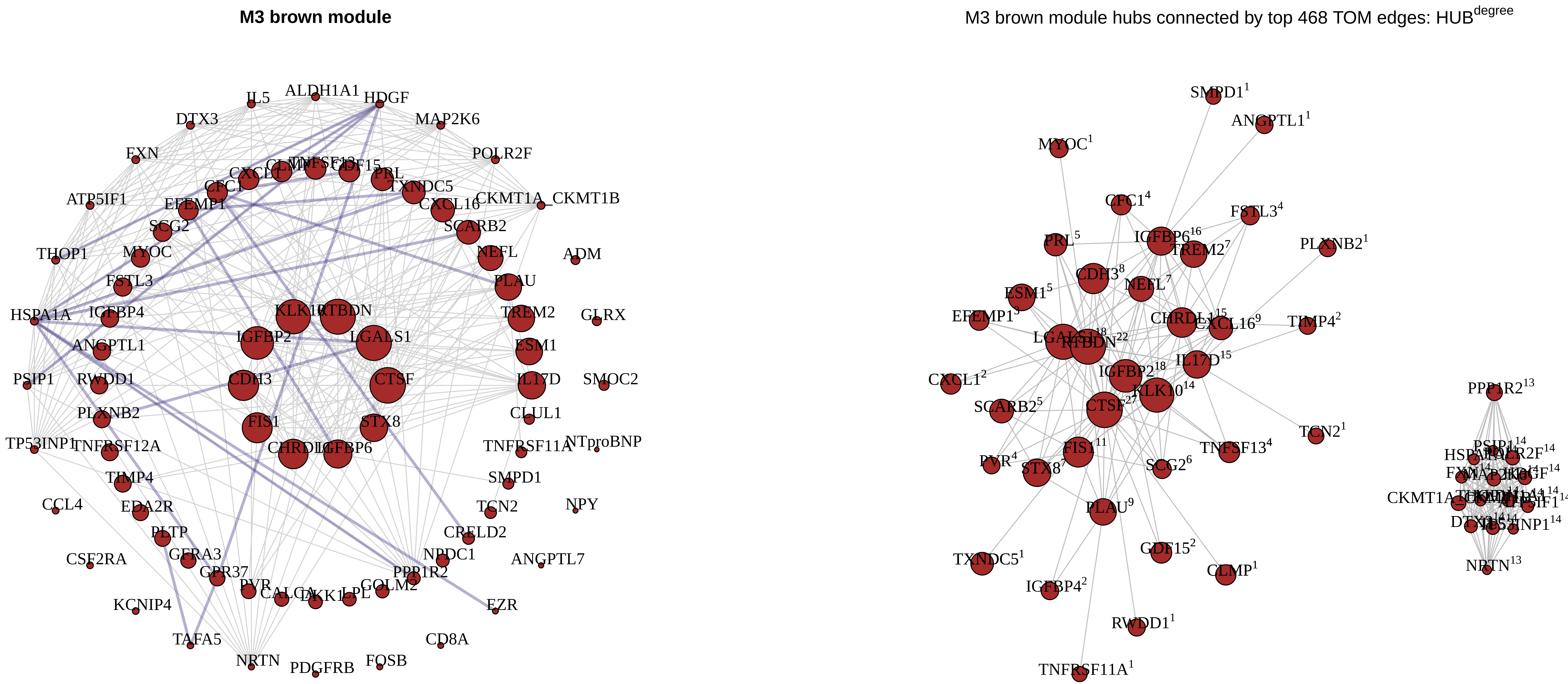
M3 brown module
IL5
ALDH1A1
HDGF
DTX3
MAP2K6
FXN
POLR2F
ATP5IF1
CKMT1A_CKMT1B
THOP1
ADM
HSPA1A
GLRX
PSIP1
SMOC2
TP53INP1
NTproBNP
CCL4
NPY
CSF2RA
ANGPTL7
KCNIP4
EZR
TAFA5
CD8A
NRTN
PDGFRB
FOSB
KLK10
RTBDN
IGFBP2
LGALS1
CDH3
CTSF
FIS1
STX8
CHRDL1
IGFBP6
PRL
TXNDC5
CXCL16
SCARB2
NEFL
PLAU
TREM2
ESM1
IL17D
CLUL1
TNFRSF11A
SMPD1
TCN2
CRELD2
NPDC1
PPP1R2
GOLM2
LPL
DKK1
CALCA
PVR
GPR37
GFRA3
PLTP
EDA2R
TIMP4
TNFRSF12A
PLXNB2
RWDD1
ANGPTL1
IGFBP4
FSTL3
MYOC
SCG2
EFEMP1
CFC1
CXCL1
M3 brown module hubs connected by top 468 TOM edges: HUB degree
SMPD1 1
ANGPTL1 1
MYOC 1
CFC1 4
FSTL3 4
PLXNB2 1
PRL 5
IGFBP6 16
TREM2 7
CDH3 8
NEFL 7
ESM1 5
EFEMP1 3
CHRDL1 15
CXCL16 9
TIMP4 2
RTBDN 22
IL17D 15
CXCL1 2
IGFBP2 18
KLK10 14
SCARB2 5
CTSF 27
TCN2 1
FIS1 11
TNFSF13 4
PVR 4
STX8 7
SCG2 6
PLAU 9
TXNDC5 1
GDF15 2
CLMP 1
IGFBP4 2
RWDD1 1
TNFRSF11A 1
PPP1R2 13
NRTN 13
CKMT1A

**M8 pink module**

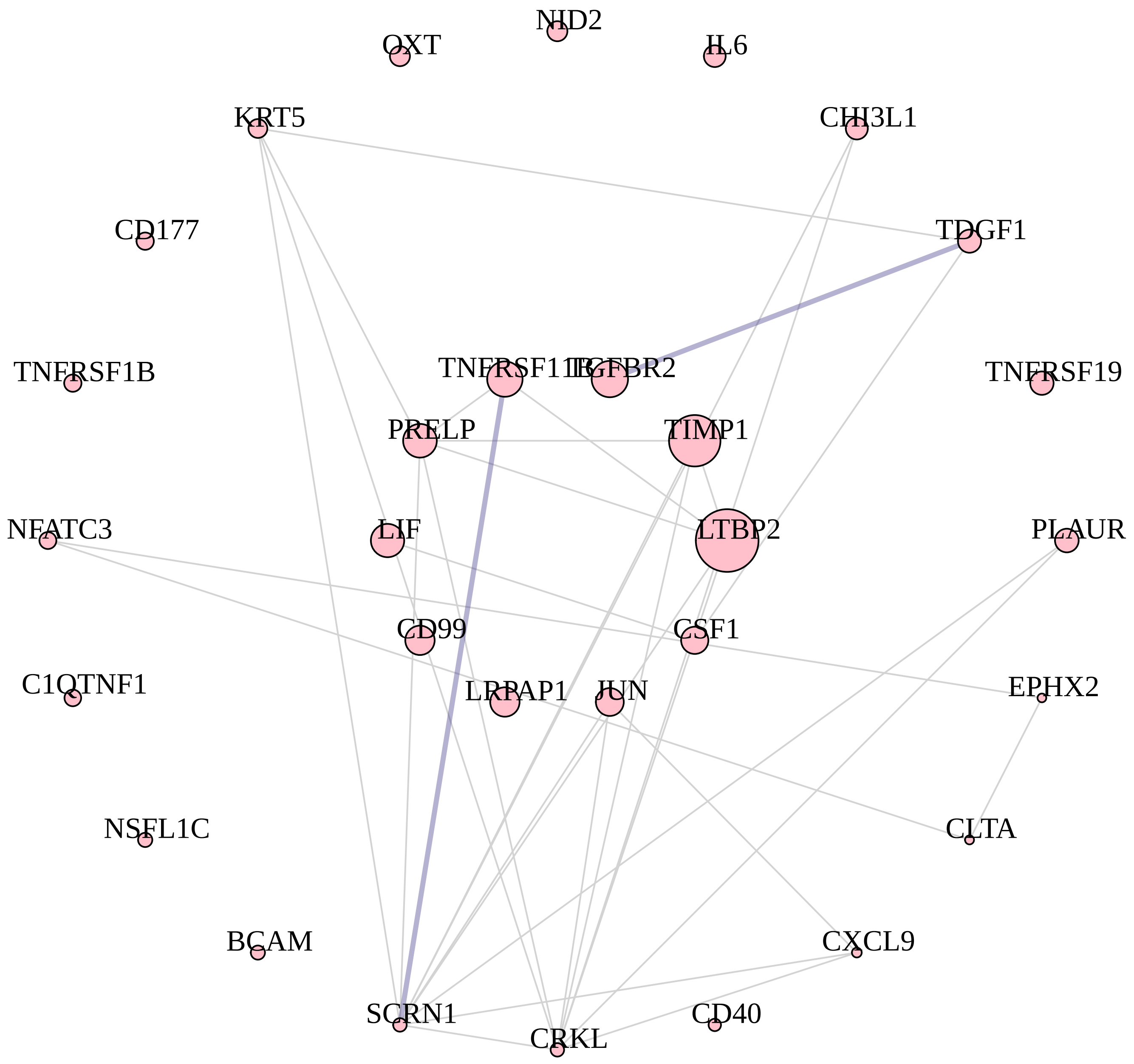


M8 pink module hubs connected by top 60 TOM edges: HUB$^{degree}$

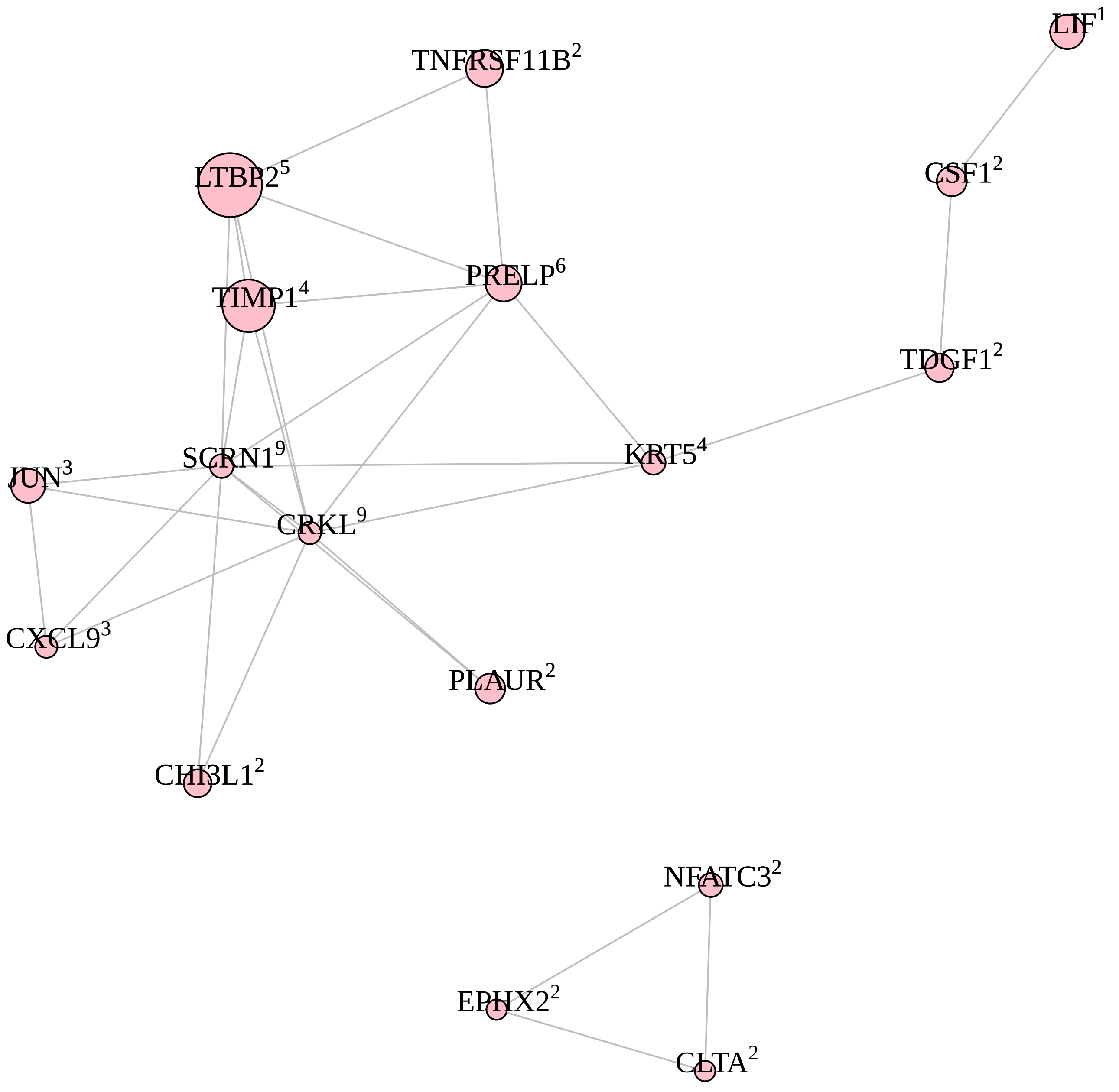

**M5 green module**

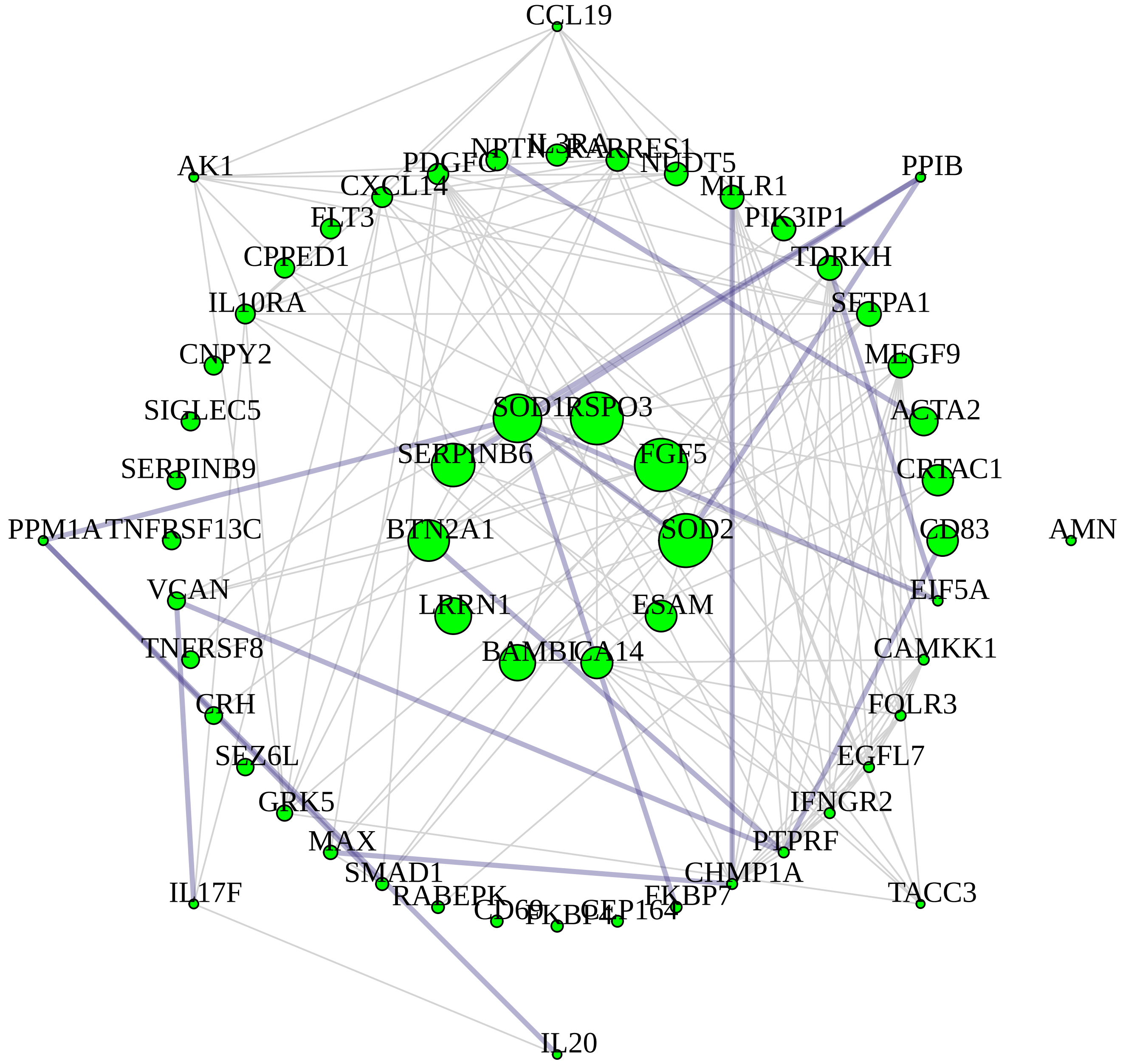


M5 green module hubs connected by top 252 TOM edges: HUB$^{degree}$

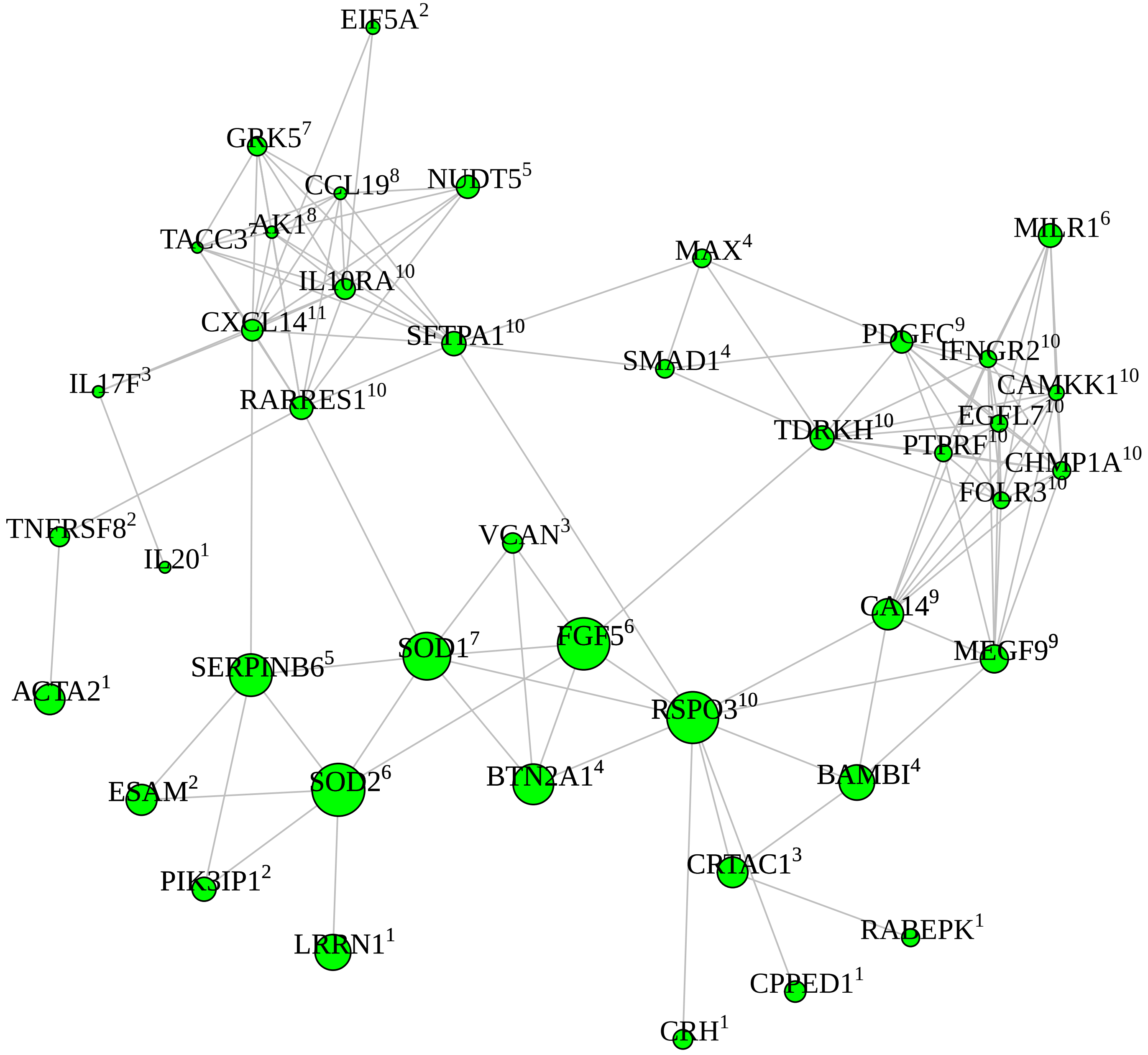

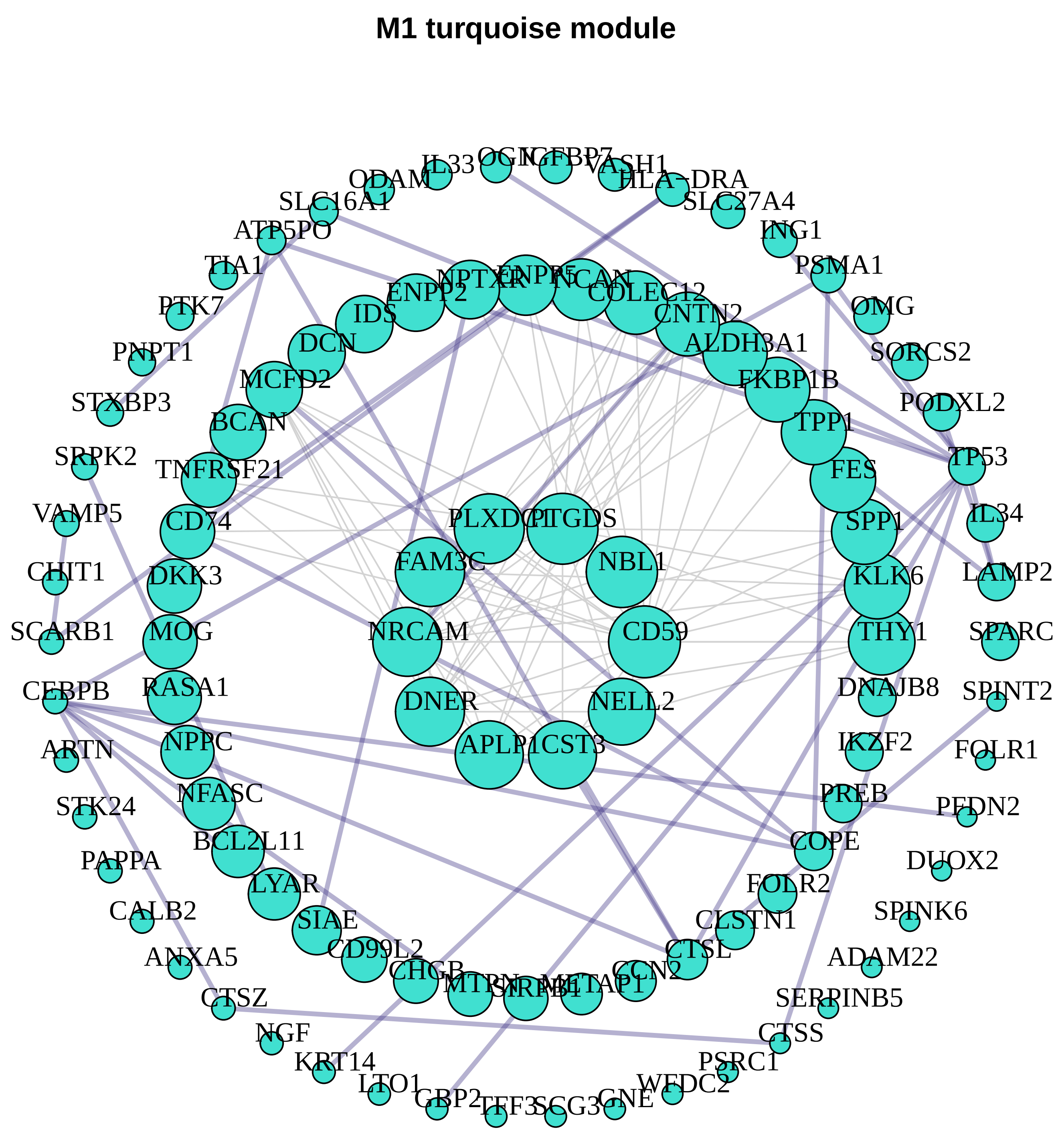
M1 turquoise module

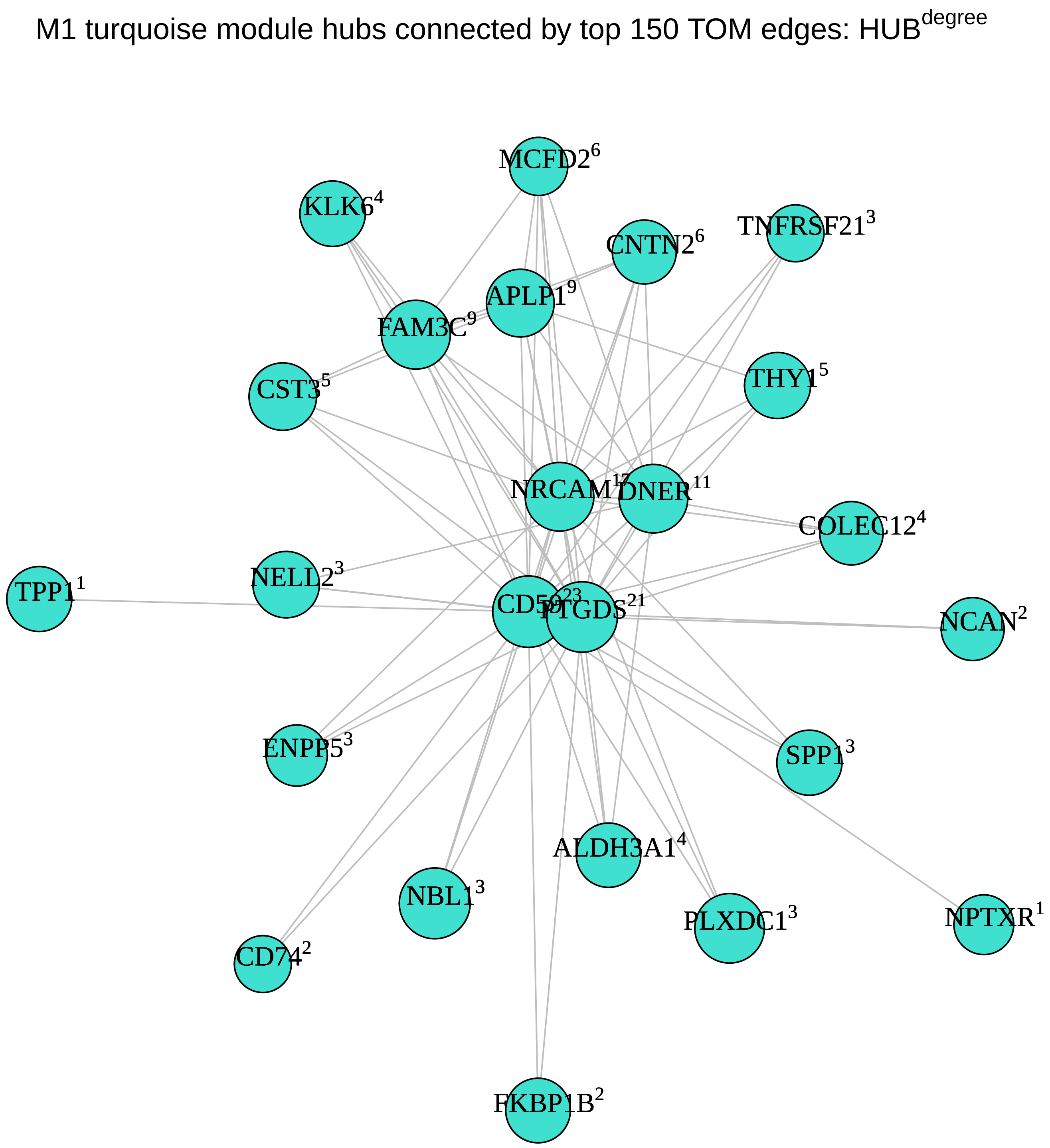
M1 turquoise module hubs connected by top 150 TOM edges: HUB$^{degree}$
MCFD2$^{6}$
KLK6$^{4}$
CNTN2$^{6}$
TNFRSF21$^{3}$
APLP1$^{9}$
FAM3C$^{9}$
CST3$^{5}$
THY1$^{5}$
NRCAM$^{15}$
DNER$^{11}$
COLEC12$^{4}$
TPP1$^{1}$
NELL2$^{3}$
CD59$^{23}$
PTGDS$^{21}$
NCAN$^{2}$
ENPP5$^{3}$
SPP1$^{3}$
ALDH3A1$^{4}$
NBL1$^{3}$
PLXDC1$^{3}$
NPTXR$^{1}$
CD74$^{2}$
FKBP1B$^{2}$

**M6 red module**

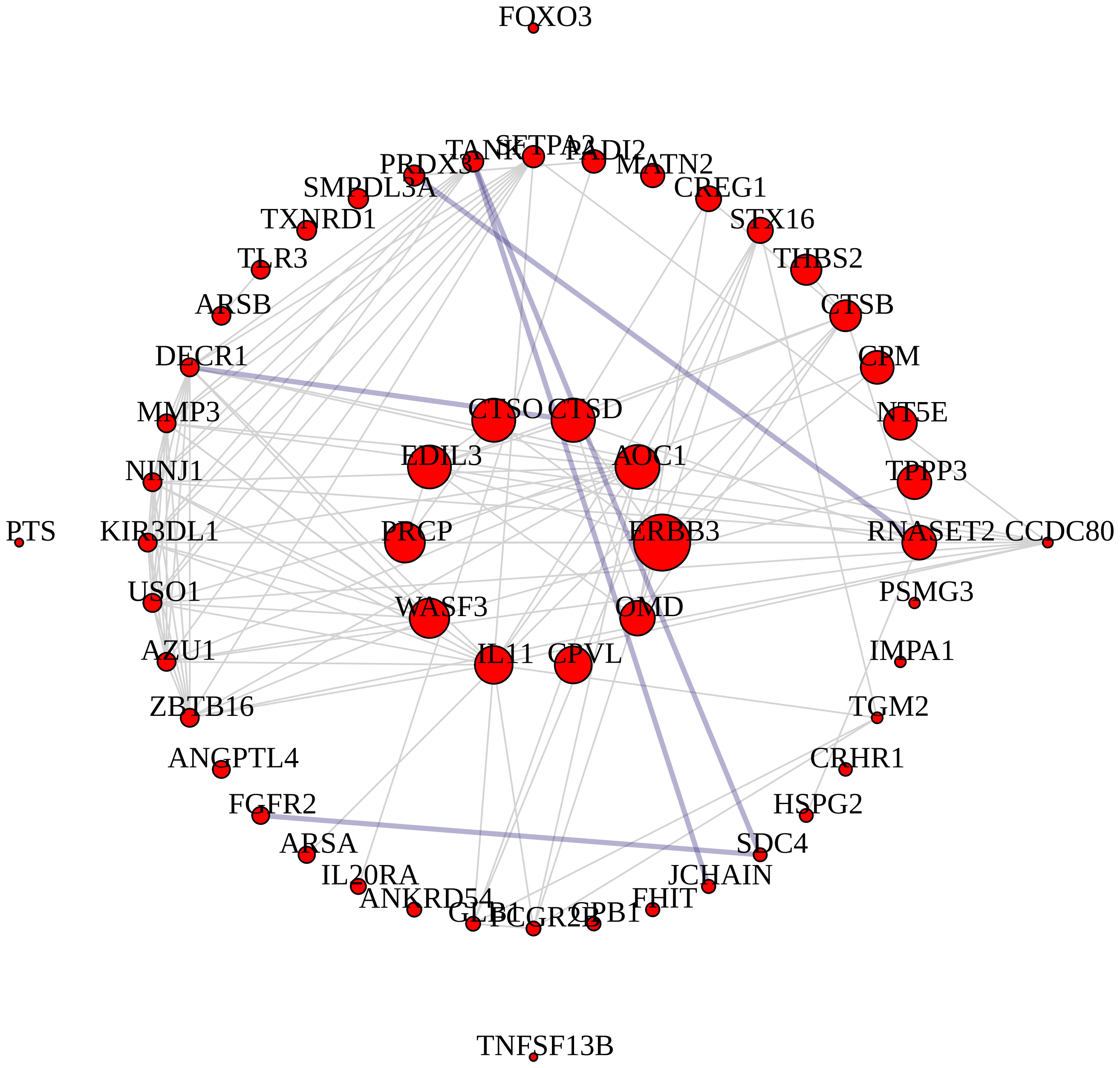


M6 red module hubs connected by top 218 TOM edges: HUB$^{degree}$

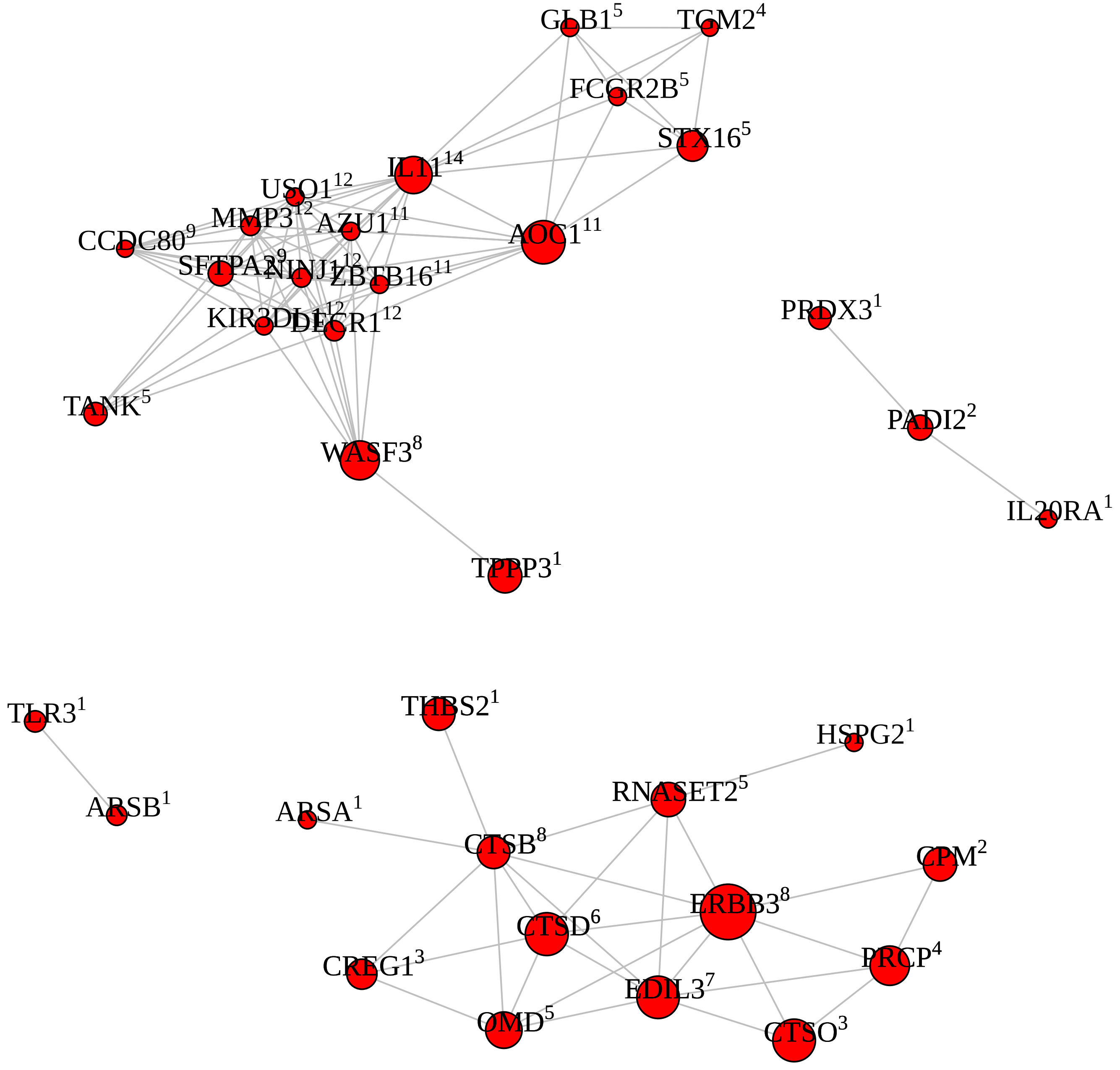

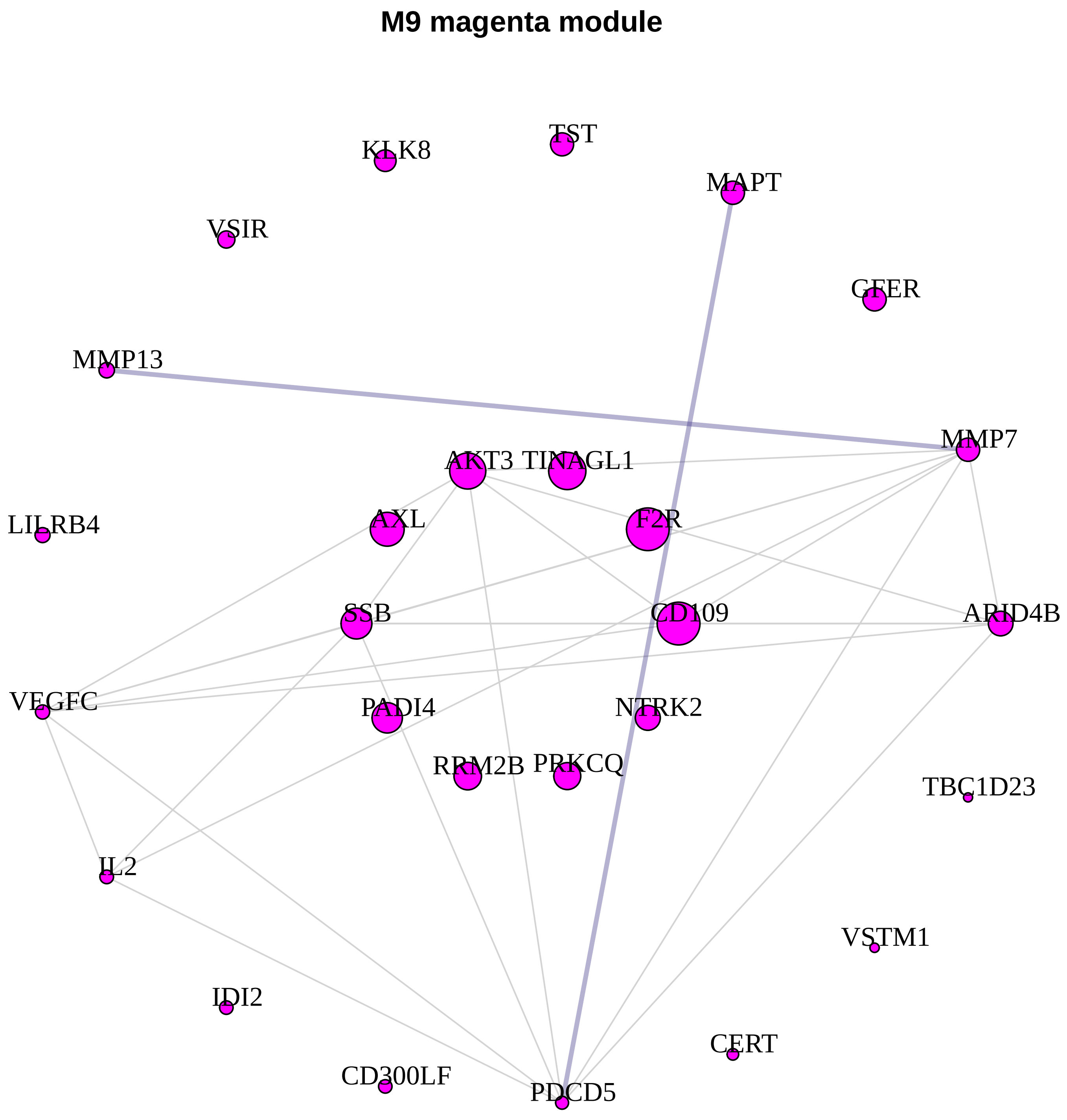

M9 magenta module
KLK8
TST
MAPT
VSIR
GFER
MMP13
MMP7
AKT3
TINAGL1
LILRB4
AXL
F2R
SSB
CD109
ARID4B
VEGFC
PADI4
NTRK2
RRM2B
PRKCQ
TBC1D23
IL2
VSTM1
IDI2
CERT
CD300LF
PDCD5


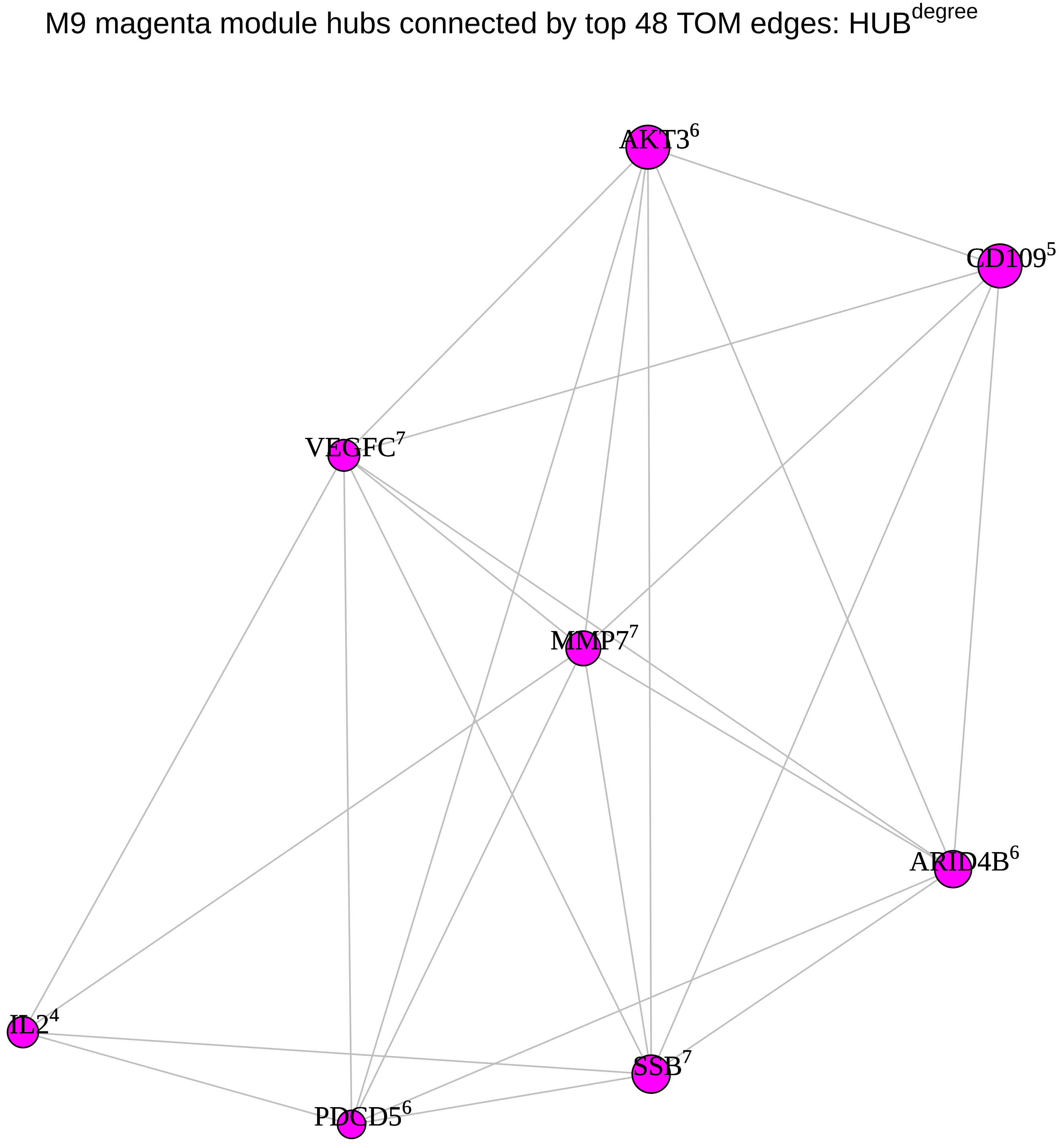

M9 magenta module hubs connected by top 48 TOM edges: HUB$^{degree}$
AKT3$^{6}$
CD109$^{5}$
VEGFC$^{7}$
MMP7$^{7}$
ARID4B$^{6}$
IL2$^{4}$
SSB$^{7}$
PDCD5$^{6}$

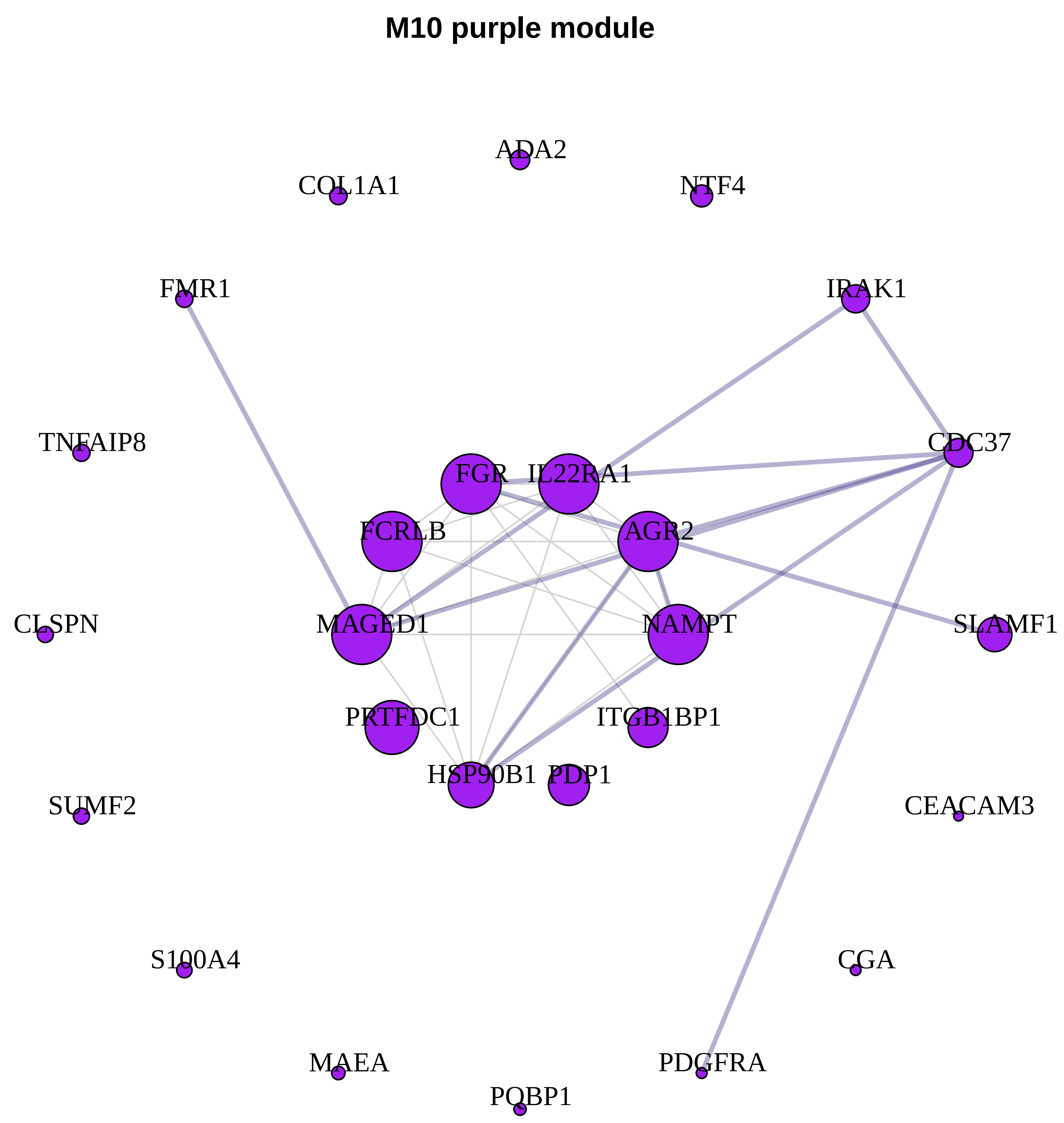
M10 purple module
ADA2
COL1A1
NTF4
FMR1
IRAK1
TNFAIP8
CDC37
FGR
IL22RA1
FCRLB
AGR2
CLSPN
MAGED1
NAMPT
SLAMF1
PRTFDC1
ITGB1BP1
HSP90B1
PDP1
SUMF2
CEACAM3
S100A4
CGA
MAEA
PDGFRA
PQBP1

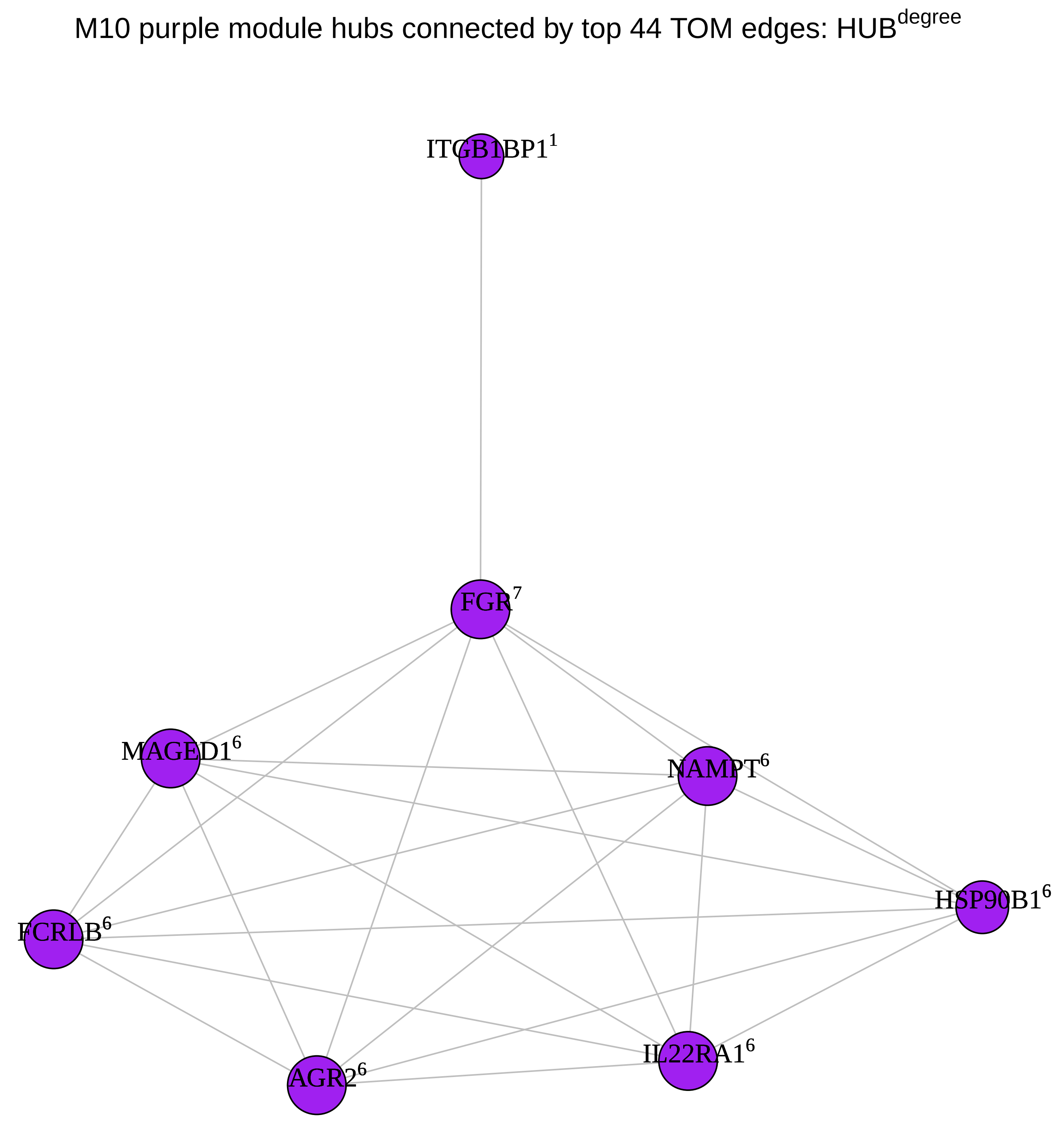
M10 purple module hubs connected by top 44 TOM edges: HUB^degree
ITGB1BP1^1
FGR^7
MAGED1^6
NAMPT^6
HSP90B1^6
FCRLB^6
IL22RA1^6
AGR2^6
