## Supplementary figures and images for "Maintenance of chronic neuroinflammation in multiple sclerosis via interferon signaling and CD8 T cell-mediated cytotoxicity"

### Supplementary File 7

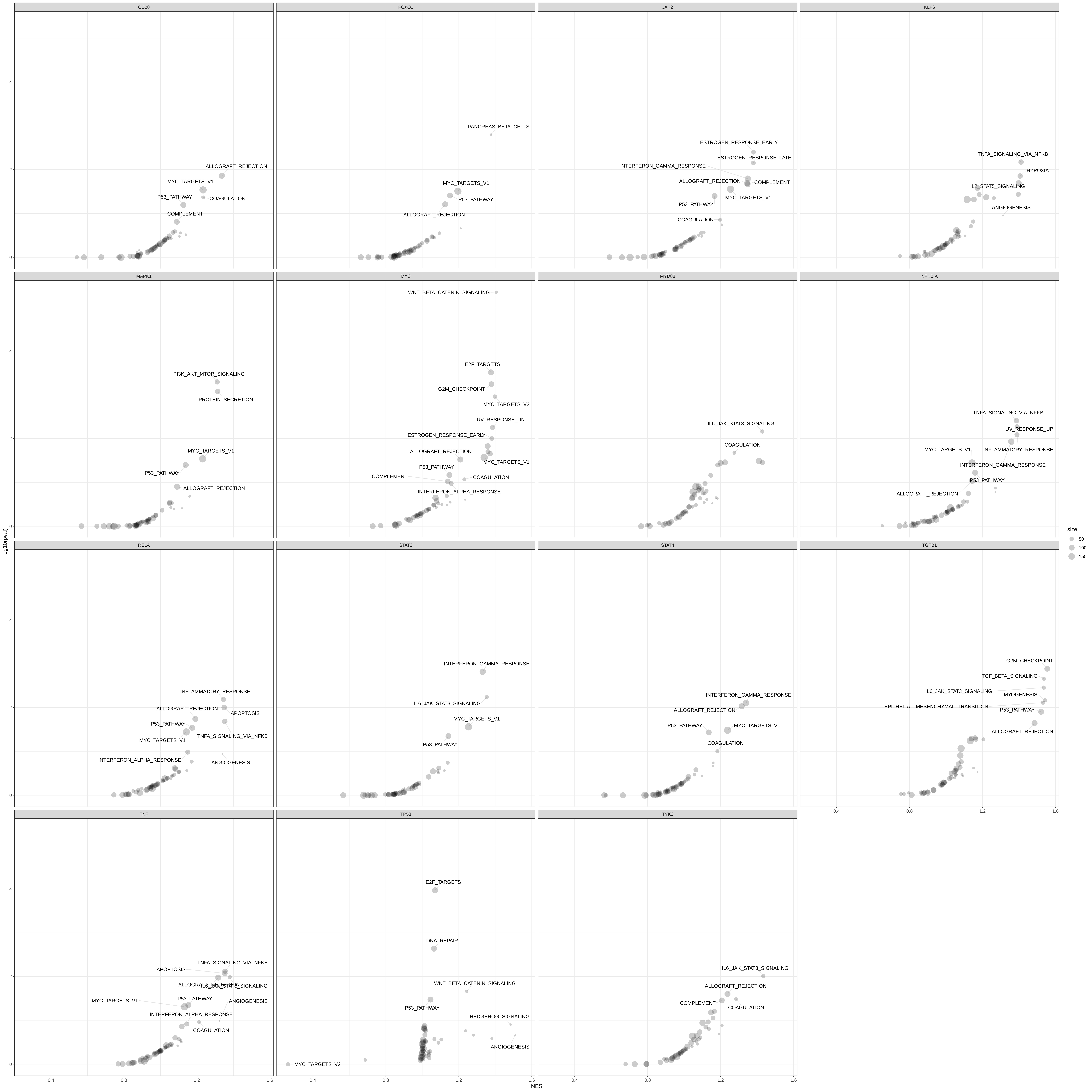
